# Genome-wide screen identifies curli amyloid fibril as a bacterial component promoting host neurodegeneration

**DOI:** 10.1101/2021.03.22.436366

**Authors:** Chenyin Wang, Chun Yin Lau, Fuqiang Ma, Chaogu Zheng

**Author notes:** Correspondence (C.Z.).

## Abstract

Growing evidence indicate that gut microbiota play a critical role in regulating the progression of neurodegenerative diseases, such as Parkinson’s disease (PD). The molecular mechanism underlying such microbe-host interaction is unclear. In this study, by feeding *C. elegans* expressing human α-syn with *E. coli* knockout mutants, we conducted a genome-wide screen to identify bacterial genes that promote host neurodegeneration. The screen yielded 38 genes that fall into several genetic pathways, including curli formation, lipopolysaccharide assembly, adenosylcobalamin biosynthesis among others. We then focused on the curli amyloid fibril and found that genetically deleting or pharmacologically inhibiting the curli major subunit CsgA in *E. coli* reduced α-syn-induced neuronal death, restored mitochondrial health, and improved neuronal functions. CsgA secreted by the bacteria colocalized with α-syn inside neurons and promoted α-syn aggregation through cross-seeding. Similarly, curli also promoted neurodegeneration in *C. elegans* models of AD, ALS, and HD and in human neuroblastoma cells.

## Introduction

Neurodegenerative diseases are characterized by protein misfolding and aggregation, leading to the formation of amyloid fibril enriched in β-sheet structures. Such protein aggregates trigger proteotoxicity, overwhelm the chaperone and degradation machineries, and eventually cause neuronal death (Douglas and Dillin, 2010). For example, Parkinson’s disease (PD) is associated with the intracellular aggregation of α-synuclein (α-syn) into Lewy bodies and Lewy neurites, which causes the degeneration of mostly dopaminergic (DA) neurons in the substantia nigra (Poewe et al., 2017). The loss of DA neurons leads to decreased dopamine signaling in the striatum, which results in impaired motor functions in PD patients. At the molecular level, α-syn is a small (140 amino acid) protein made of three domains: an N-terminal domain, a non-Aβ component (NAC) domain that is the fibrilization core, and a C-terminal region. Missense mutations in the N-terminal domain, such as A30P, G46K, and A53T, result in autosomal dominant familial PD by producing α-syn mutant proteins that are more prone to misfolding and aggregation than the wild-type α-syn (Stefanis, 2012). Mutant α-syn proteins form toxic β-sheet-like oligomers that cause mitochondrial dysfunctions, oxidative stress, disruption in calcium homeostasis, and neuroinflammation, which all lead to neurodegeneration (Poewe et al., 2017). Effective therapeutic intervention that prevent α-syn aggregation is currently missing.

Studies in the last few years have suggested that the gut microbiota may play an important role in the pathogenesis of neurodegenerative diseases (Quigley, 2017). For example, antibiotic treatment ameliorate the pathophysiology of PD mice and microbial recolonization after the treatment restored the PD symptoms (Sampson et al., 2016). Colonization of α-syn-overexpressing mice with microbiota from PD patients enhanced the physical impairments, compared to transplantation of microbiota from healthy human donors (Sampson et al., 2016). In addition to animal models, clinical studies have also provided evidence for a microbiota-gut-brain link in PD. Gastrointestinal dysfunction was frequently found in PD patients (Quigley, 1996) and infection with *Helicobacter pylori* has been linked with disease severity and progression (Tan et al., 2015). Sequencing of the fecal samples of PD patients revealed changes in the gut bacterial composition (e.g. increased *Lactobacillaceae*) compared to healthy individuals (Barichella et al., 2019; Hopfner et al., 2017). Similar to PD, gut bacteria in mouse models of Alzheimer’s disease (AD) promoted amyloid pathology (Harach et al., 2017) and altered gut microbiome composition was also observed in AD patients (Cattaneo et al., 2017).

Despite the growing connection between the disturbed gut microbiota and the development of neurodegenerative diseases, mechanistic understanding of the communication between the bacteria and the nervous system is limited. Most theories focus on the neurodegenerative effects of the systemic inflammation and neuroinflammation caused by the abnormal microbiota. Whether bacteria proteins or metabolites can directly act on the host neurons to module the progression of neurodegeneration induced by α-syn or Aβ proteotoxicity is unclear. This limitation is largely due to the lack of a simple model that allows systematic tests of individual bacterial components for their neuronal effects in promoting degeneration.

To address the problem, we employed a *Caenorhabditis elegans* model of PD that expressed the human α-syn proteins in *C. elegans* neurons to investigate the mechanisms of microbial regulation on PD. Because *C. elegans* use bacteria as their natural diet and can be easily cultivated under axenic or monoaxenic conditions, it has emerged as an important organism to model microbe-host interaction. In fact, previous studies found that alteration of the bacterial genome affected the development, metabolism, and behaviour of *C. elegans* (Watson et al., 2014; Zhang et al., 2019).

In this study, we screened all non-essential *E. coli* genes for their effects on PD pathogenesis by feeding individual *E. coli* knockout mutants to PD *C. elegans* and assessing the severity of neurodegeneration. This screen identified 38 *E. coli* genes whose deletion led to amelioration of PD symptoms. These genes fall into distinct genetic pathways, including curli formation, lipopolysaccharide (LPS) production, lysozyme inhibition, biosynthesis of adenosylcobalamin, and oxidative stress response. These results suggested that diverse bacteria components could promote neurodegeneration. As an example, we next focused on the role of bacterial curli amyloid fibril on PD and found that deleting the curli genes *csgA* and *csgB* in the *E. coli* genome reduced α-syn-induced cell death, restored mitochondrial health, and improved neuronal functions. Using antibody staining and biochemical analysis, we showed that CsgA promoted α-syn aggregation and removing curli in the bacteria diet enabled proteasome-dependent degradation of α-syn in neurons. Although previous studies observed the cross-seeding between curli and α-syn *in vitro*, we provided direct evidence to show that bacteria-derived CsgA colocalized with α-syn in *C. elegans* neurons at a single-cell resolution. More importantly, we extended our findings into *C. elegans* models of AD, Amyotrophic lateral sclerosis (ALS), and Huntington’s disease (HD), and into human neuroblastoma SH-SY5Y cells. Overall, our studies indicate that bacteria components, such as curli, can have direct neurodegenerative effects by promoting protein aggregation.

## Results

### A genome-wide screen identified *E. coli* genes that promote human α-synuclein-induced neurodegeneration in *C. elegans*

To systematically identify bacterial genes that contribute to neurodegeneration in the host, we employed *C. elegans* transgenic animals that express the human α-syn in all neurons and screened for *E. coli* K12 knockout mutants that could ameliorate *C. elegans* neurodegenerative phenotypes when fed to the animals. Pan-neuronal expression (using the *aex-3* promoter) of the pro-aggregating human α-syn A53T mutants but not the wild-type α-syn led to degeneration of the motor neurons, causing uncoordinated (Unc) movements in both larva and adults (Lakso et al., 2003). The penetrance of this Unc phenotype is 100% in animals carrying the *aex-3p::α-syn(A53T)* transgene and fed with the wild-type *E. coli* K12 strain. By screening the 3985 K12 knockout mutants in the Keio collection (Baba et al., 2006), we identified 380 *E. coli* mutants that restored normal locomotion (non-Unc) in at least 25% of the PD animals in the first round (Figure 1A). These 380 positive clones were subjected to three more rounds of locomotion screens, and we obtained 172 mutants that led to consistent recovery of locomotion in ≥ 25% of the PD animals on average (Figure 1B). We then subjected the 172 positive clones to a visual screen for the suppression of neuronal death using a dopaminergic (DA) neuronal marker. When fed with wild-type K12, only ~10% of the PD animals carrying the *aex-3p::α-syn(A53T)* transgene had two visible ADE neurons labelled by *dat-1p::GFP* at day 2 adult stage (Figure 1C). By screening the above 172 clones for three repeated rounds, we identified 104 *E. coli* mutants that resulted in the survival of two ADE neurons in ≥ 25% of the animals on average.

**Figure 1.**
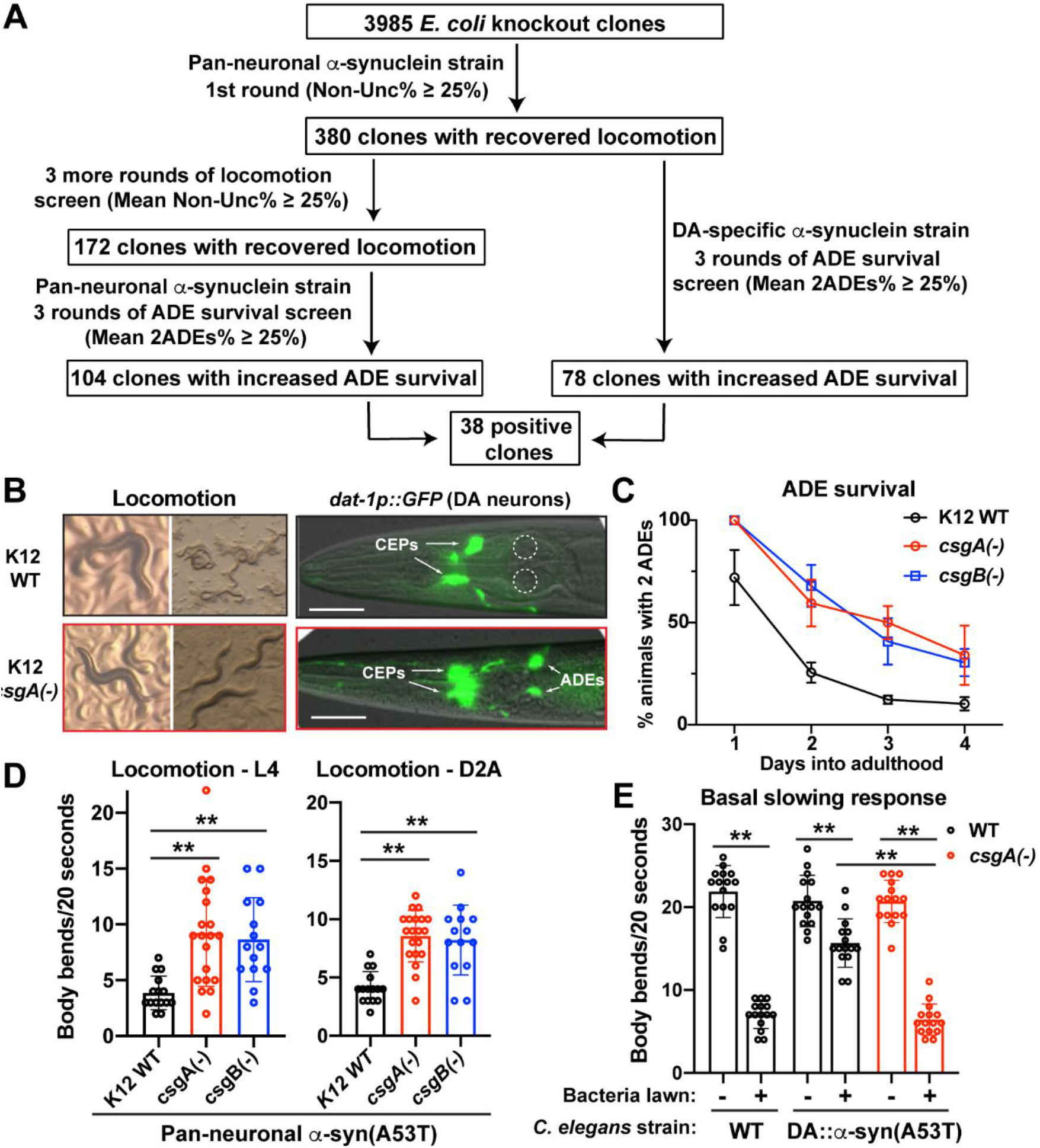
Genome-wide screen for pro-neurodegenerative genes in *E. coli*. (A) The flowchart of the screen using UM10 *unkIs7[aex-3p::α-syn(A53T), dat-1p::gfp]* and UM6 *unkIs9 [dat-1p::α-syn(A53T), dat-1p::gfp]* strains and the Keio library. (B) Representative images of uncoordinated movement and ADE neurodegeneration in UM10 animals fed with wild-type (WT) K12 *E. coli* and the restored locomotion and intact ADE neurons in animals fed with *csgA*(-) K12. (C) Percentage of UM10 animals with 2 ADE neurons when fed with WT, *csgA*(-) and *csgB*(-) K12 on various days into adulthood. (D) Locomotion rate of UM10 animals at L4 stage and day-two-adult stage when fed with K12 WT, *csgA*(-) and *csgB*(-). Mean ± SD were shown and each dot represents one animal assayed. Double asterisks indicate statistical significance (*p* < 0.01) in multiple comparisons using ANOVA analysis followed by a Tukey HSD post-hoc test. The same applies to all other figures. (E) Locomotion rate of wild-type N2 and UM6 animals on and off the bacteria lawn. UM6 animals exhibited impairment in food-induced basal slowing response when fed with WT K12, and the slowing response is restored when UM6 animals were fed with *csgA(-)* K12.

To avoid any bias associated with specific genetic background, we conducted a separate visual screen using an independent PD model, in which the human α-syn A53T mutant protein was expressed from the DA-specific *dat-1* promoter. We fed the 380 first round positive *E. coli* knockout mutants to animals carrying the *dat-1p::α-syn(A53T)* transgene. Through three rounds of screens, we identified 78 *E. coli* mutants that led to the survival of two ADE neurons in ≥ 25% in these DA-specific degenerative model (Figure 1A). By overlapping these 78 positive clones with the 104 positive hits found using the pan-neuronal model, we obtained the final 38 *E. coli* mutants that significantly inhibited neurodegeneration induced by human α-syn A53T mutants.

We categorized the 38 *E. coli* genes that contribute to neurodegeneration based on their functions and several microbial genetic pathways emerged (Table 1). For example, we identified genes responsible for the formation of curli amyloid fibril (*csgA* and *csgB*), the production and assembly of LPS (*lapA*, *lapB*, *lpcA*, *rfe*, and *pldA*), biosynthesis of adenosylcobalamin (*cobS*, *btuR*, and *eutT*), inhibition of eukaryotic lysozyme (*ivy* and *ydhA*), as well as genes involved in oxidative stress response, energy homeostasis, membrane transport, and other functions. Our systematic screen revealed extensive microbe-host interactions, through which bacterial molecules promote α-syn proteotoxicity and neurodegeneration in the host. In this study, we focus on the mechanisms by which bacterial curli promotes neurodegeneration.

**Table 1.** Deletion of 38 *E. coli* genes led to amelioration of α-syn-induced neurodegeneration.

| Bacterial clones | Pan-neuronal α-syn (A53T) | | DA-specific α-syn (A53T) | Gene function |
| --- | --- | --- | --- | --- |
|  | Non-Unc% | 2ADEs% | 2ADEs% |  |
| K12 WT | 0% | 6±8% | 11±10% | |
| **Curli amyloid fibril formation** |  |  |  |  |
| *csgA*(-) | 33±4% | 50±7% | 50±7% | Major subunit of curlin |
| *csgB*(-) | 33±8% | 42±6% | 42±12% | Minor subunit of curlin |
| **LPS production and assembly** |  |  |  |  |
| *lapA*(-) | 35±9% | 28±8% | 35±19% | Lipopolysaccharide assembly protein A |
| *lapB*(-) | 41±5% | 35±7% | 42±6% | Lipopolysaccharide assembly protein B |
| *lpcA*(-) | 35±9% | 42±13% | 33±14% | Sedoheptulose 7-phosphate isomerase; LPS biosynthesis |
| *rfe*(-) | 29±16% | 45±15% | 38±8% | Synthesis of ECA and lipopolysaccharide O-side chains |
| *pldA*(-) | 28±10% | 43±19% | 30±4% | Outer membrane phospholipase A; mlaA-dependent LPS hyperproduction |
| **Adenosyl-cobalamin biosynthesis** |  |  |  |  |
| *cobS*(-) | 31±5% | 35±7% | 33±6% | Cobalamin 5'-phosphate synthase |
| *btuR*(-) | 29±4% | 27±6% | 35±11% | Cobinamide/cobalamin adenosyltransferase |
| *eutT*(-) | 30±5% | 40±15% | 37±10% | Putative ethanolamine utilization cobalamin adenosyltransferase |
| **Inhibitors of eukaryotic lysozyme** |  |  |  |  |
| *ivy*(-) | 45±27% | 43±12% | 42±12% | Inhibitor of vertebrate lysozyme |
| *ydhA*(-) | 30±4% | 43±18% | 32±14% | Inhibitor of c-type lysozyme, putative lipoprotein |
| **Oxidative stress response** |  |  |  |  |
| *gpmI*(-) | 43±17% | 37±10% | 57±17% | 2,3-bisphosphoglycerate-independent phosphoglycerate mutase |
| *yaaA*(-) | 43±11% | 45±15% | 28±15% | Peroxide stress resistance protein |
| *sodA*(-) | 35±8% | 38±15% | 30±15% | Superoxide dismutase (Mn) |
| *msrA*(-) | 38±8% | 38±12% | 30±8% | Peptide methionine sulfoxide reductase |
| *nrfA*(-) | 26±10% | 27±2% | 35±11% | Cytochrome c nitrite reductase subunit |
| *yfiD*(-) | 31±5% | 35±7% | 32±2% | Stress-induced alternate pyruvate formate-lyase subunit |
| **Metabolism and energy homeostasis** |  |  |  |  |
| *pck*(-) | 39±7% | 37±9% | 48±6% | Phosphoenolpyruvate carboxykinase; gluconeogenesis |
| *tpiA*(-) | 26±2% | 27±5% | 43±2% | Triose-phosphate isomerase; glycolysis and gluconeogenesis |
| *ldhA*(-) | 39±16% | 25±4% | 27±14% | D-lactate dehydrogenase |
| *lldD*(-) | 29±6% | 31±5% | 30±8% | L-lactate dehydrogenase |
| *tdh*(-) | 31±8% | 32±6% | 32±6% | Threonine dehydrogenase; major catabolic pathway for threonine utilization |
| *cysQ*(-) | 41±10% | 38±6% | 28±8% | 3'(2'),5'-bisphosphate nucleotidase |
| *nrdD*(-) | 26±10% | 38±2% | 27±6% | Anaerobic ribonucleoside-triphosphate reductase; DNA synthesis |
| *betA*(-) | 25±13% | 50±16% | 33±12% | Choline dehydrogenase |
| **Transporter** |  |  |  |  |
| *yjcD*(-) | 31±8% | 33±10% | 38±2% | Guanine/hypoxanthine transporter |
| *mdtC*(-) | 30±4% | 43±14% | 37±8% | Multidrug efflux pump RND permease subunit |
| *ybbY*(-) | 29±4% | 40±15% | 32±5% | Putative purine transporter |
| *tolQ*(-) | 30±6% | 50±11% | 25±12% | An inner membrane component of the Tol-Pal system |
| **Transcription factors and DNA regulation** |  |  |  |  |
| *yebC*(-) | 33±7% | 40±16% | 38±6% | Putative transcriptional regulator |
| *ybbS*(-) | 30±6% | 42±16% | 43±2% | DNA-binding transcriptional activator |
| *rdgC*(-) | 26±5% | 43±15% | 30±14% | Nucleoid-associated protein RdgC |
| **Others** |  |  |  |  |
| *ppiD*(-) | 41±17% | 37±16% | 42±10% | Periplasmic folding chaperone |
| *baeS*(-) | 31±4% | 40±8% | 35±21% | Sensory histidine kinase |
| *ycaC*(-) | 31±4% | 38±15% | 42±5% | Putative hydrolase |
| *ybjQ*(-) | 30±13% | 32±6% | 43±15% | Putative heavy metal binding protein |
| *yfeY*(-) | 29±2% | 45±15% | 27±17% | DUF1131 domain-containing lipoprotein |

### Curli subunits CsgA and CsgB are required for α-syn-induced degenerative phenotypes

Among the top hits in our screen are *csgA* and *csgB*, which code for the major and minor curli subunits. Curli are amyloid fibril secreted by certain enterobacterial strains, such as *E. coli* and Salmonella, and are important for biofilm formation (Evans and Chapman, 2014). Deletion of *csgA* or *csgB* in *E. coli* K12 fed to *C. elegans* resulted in significantly improved motor functions (measured as the number of body bends per 20 seconds) in animals expressing α-syn(A53T) pan-neuronally (Figure 1D) and largely restored dopaminergic neuron functions (measured as food-induced basal slowing response) in animals expressing α-syn(A53T) specifically in DA neurons (Figure 1E). These results suggest that *E. coli* curli promotes α-syn-induced neurodegeneration in *C. elegans*.

Congo red can stain curli amyloid fibril. As expected, deletion of *csgA* and *csgB* led to the complete loss of Congo red staining (Figure 2A). We also stained the other 36 *E. coli* mutants identified from our screen and found that *tolQ(-)*, *lpcA(-)*, *lapA(-)*, and *lapB(-)* mutants showed weak Congo red staining, indicating reduced curli production. These four genes were previously found to be associated with curli biogenesis (Smith et al., 2017), so their effects in promoting neurodegeneration may be partly connected to their roles in enhancing curli production. In contrast, the rest 32 mutants showed largely wild-type level of staining (Figure S1A).

**Figure 2.**
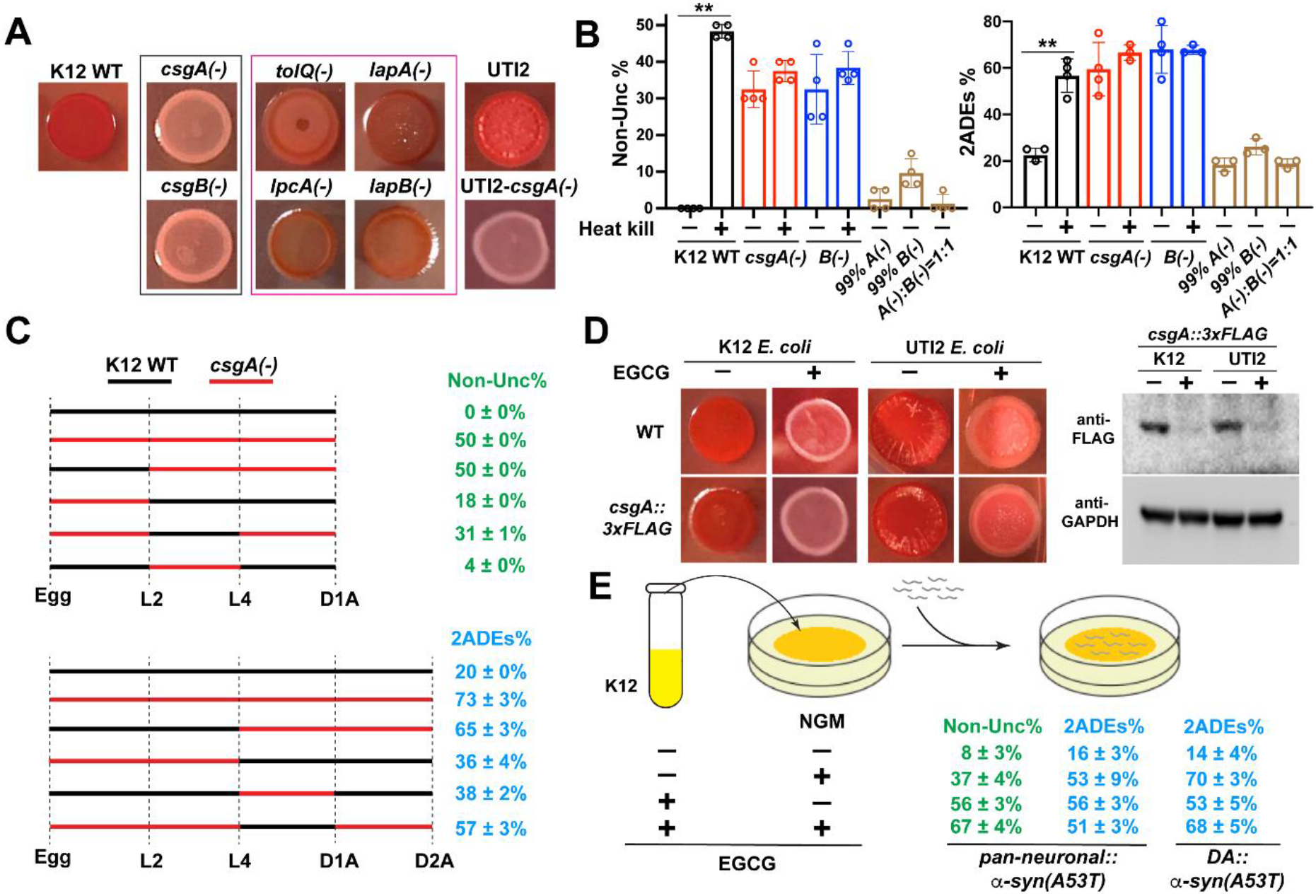
Bacterial curli production promotes α-syn-induced neurodegeneration. (A) WT and mutant K12 *E. coli* and WT and *csgA(-)* UTI2 *E. coli* were grown on Congo Red indicator plates at 25 C for two days to visualize curli production. Curli subunit mutants are in black box and mutants that were found in our screen and also showed reduced curli production are in pink box. (B) Percentage of UM10 *unkIs7[aex-3p::α-syn(A53T), dat-1p::gfp]* animals with Non-Unc phenotype at L4 stage or 2 ADE neurons at day-two-adult stage when fed with either heat-killed WT, *csgA*(-), and *csgB*(-) K12 or a mixture of WT with *csgA(-)* or *csgB(-)* K12 at 1:99 ratio (indicated as 99% A(-) or B(-), respectively) or a mixture of *csgA(-)* and *csgB(-)* at 1:1 ratio (indicated as A(-):B(-)=1:1). Mean ± SD were shown and each dot represents one independent experiment with 20 to 30 animals scored. (C) Percentage of UM10 animals with Non-Unc phenotype and two intact ADE neurons when fed with either K12 WT or *csgA(-)* at different developmental stages. The dotted lines indicate the timing of diet switch at specific stages from egg to day-one-adults (D1A) or day-two-adults (D2A). (D) WT K12 and UTI2 *E. coli* and their derivative containing the *csgA::3xFLAG* genomic edits were grown on Congo red indicator plates with or without 200 μg/ml EGCG. Western blot of the bacteria lysate using anti-FLAG antibodies were used to confirm the inhibition of CsgA expression by EGCG. Anti-GAPDH blotting was used as loading control. (E) Percentage of UM10 animals with Non-Unc phenotype and two intact ADE neurons or the percentage of UM6 *unkIs9 [dat-1p::α-syn(A53T), dat-1p::gfp]* animals with two ADEs when grown on NGM plates that contained 200 μg/ml EGCG or empty vehicle and seeded with EGCG-treated or untreated WT K12. For the treatment of K12 with EGCG, a single colony was cultured with LB medium containing 200 μg/ml EGCG overnight prior to seeding on NGM plate. The mean ± SD of three independent experiments (25 animals were sc ored for each experiment) is shown.

To support that curli functions as structured protein fibrils in *C. elegans*, we heat-killed the K12 wild-type bacteria to denature all proteins and found that the heat-kill phenocopied the *csgA(-)* and *csgB(-)* deletion in suppressing α-syn-induced locomotion defects and ADE degeneration (Figure 2B). Importantly, heat-kill did not further enhance the effects of curli deletion. Interestingly, we found that mixing the wild-type K12 with *csgA(-)* or *csgB(-)* bacteria strongly suppressed the neuroprotective effects of the mutants, suggesting that a small amount of curli may be enough to trigger neurodegeneration (Figure 2B). Moreover, mixing *csgA(-)* with *csgB(-)* also eliminated their neuroprotective effects (Figure 2B), likely because the CsgA proteins produced by *csgB(-)* mutants can bind to CsgB produced by *csgA(-)* mutants. Such cross-seeding allows the assembly of curli fibril and was previously observed (Evans and Chapman, 2014).

Next, we switched the diet of the PD animals between wild-type K12 and *csgA(-)* mutants at different developmental stages and found that the α-syn-induced locomotion defects and ADE degeneration is largely associated with post-L4 and adult consumption of the curli-producing bacteria (Figure 2C). Whether the animals were exposed to curli in larval development did not affect much on neurodegeneration. Since L4 and adult animals consumed more food than younger animals, the amount of curli uptake may be associated with the severity of the degenerative phenotypes.

### Pharmacological inhibition of *csgA* expression suppresses neurodegeneration

In addition to genetic inactivation of curli subunits, we also used pharmacological agents to inhibit curli production and tested the effects on neurodegeneration. Epigallocatechin gallate (EGCG) is a polyphenol extracted from green tea and showed a strong effect in inhibiting biofilm formation by impairing curli assembly in *E. coli* (Serra et al., 2016). We confirmed that EGCG treatment completely eliminated Congo red staining signal (Figure 2D). To measure endogenous CsgA levels, we initially engineered the *csgA* locus to insert a C-terminal mCherry, but the resulted *csgA::mCherry* fusion completely blocked curli production (Figure S1B), suggesting that fusing large fluorescent proteins (~28 kDa for mCherry) with small amyloidogenic proteins (~15 kDa for CsgA) may interfere with fibril formation. We then inserted a small 3×FLAG tag at the *csgA* locus, and the resulted *K12-csgA::3xFLAG* strains produced normal levels of curli and formed biofilm as the wild-type K12. As expected, EGCG strongly inhibited endogenous *csgA* expression detected by the FLAG tag (Figure 2D).

Because EGCG is also known to have neuroprotective effects in PD models (Xu et al., 2016), we set out to assess the contribution of its bacterial effects in neuroprotection. To our surprise, treating the K12 bacteria diet with EGCG alone is sufficient to create a strong inhibition of α-syn-induced neurodegeneration (Figure 2E). The inhibitory effects was almost indistinguishable with animals that also received the EGCG treatment in addition to being fed with the EGCG-treated diet. This result suggests that, at least in the *C. elegans* PD model, the neuroprotective effects of EGCG may be largely due to its activities in inhibiting bacterial curli production.

### Bacterial curli promotes α-syn aggregation

Because both CsgA and α-syn are enriched in β-sheet structures, we next examined whether CsgA promotes α-syn aggregation *in vivo*. First, using a transgenic strain with muscle-specific expression of α-syn::YFP fusion, we found the α-syn aggregation in the form of fluorescent puncta is strongly inhibited when the animals were fed with *csgA(-)* or *csgB(-)* bacteria (Figure 3A). Conversely, when the animals were fed with a uropathogenic *E. coli* strain (UTI2) that has high levels of curli production (Figure 2A), much more α-syn aggregates were observed than in animals grown on wild-type K12 (Figure 3A). As expected, *csgA(-)* knockout in the UTI2 strain significantly reduced the number of the aggregates. Thus, not only curli promotes α-syn aggregation but also the amount of curli consumed by the animals correlates with the severity of α-syn aggregation.

**Figure 3.**
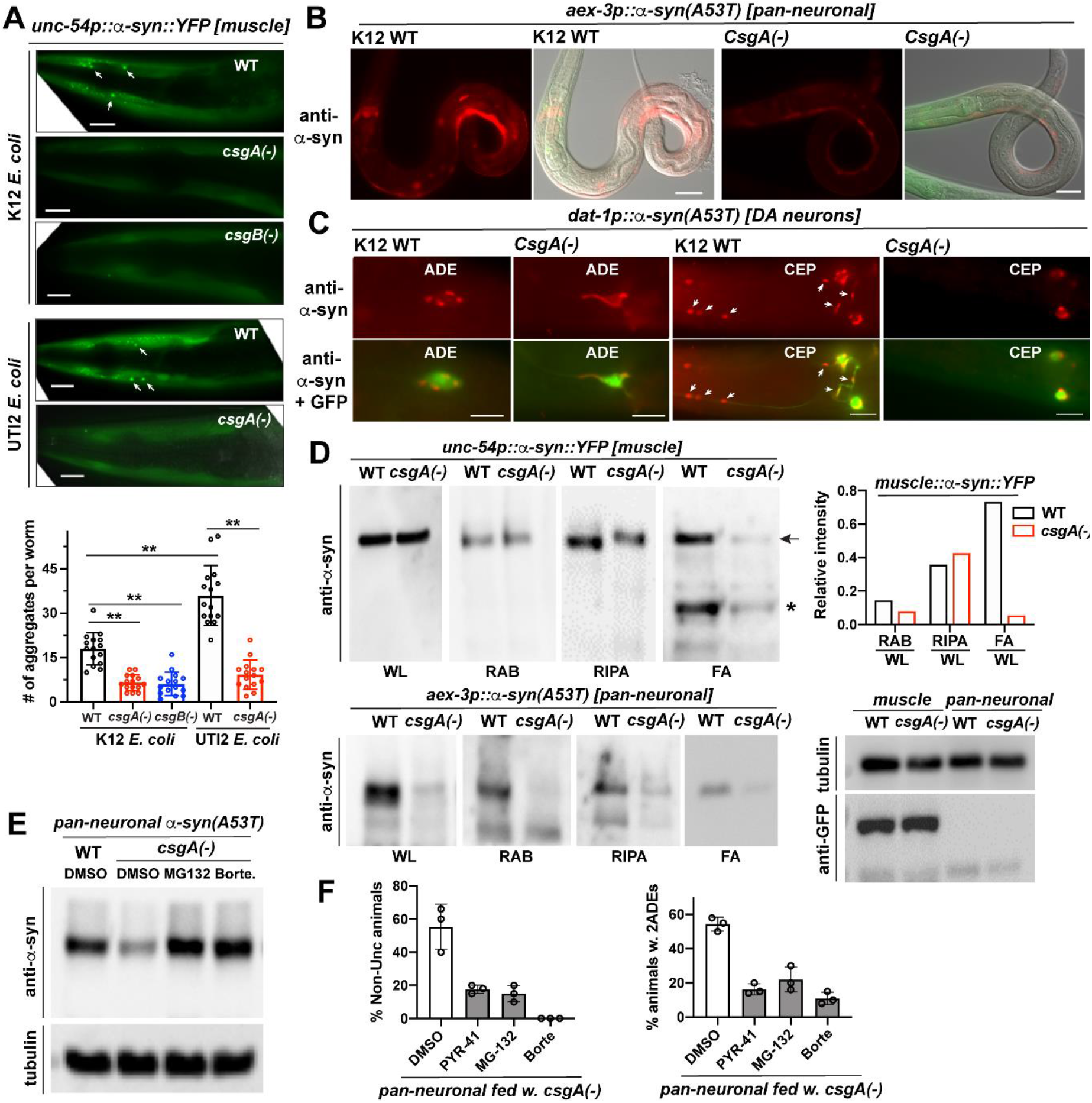
Bacterial curli promotes α-syn aggregation. (A) Representative images of α-syn::YFP aggregates in the muscle of NL5901 *pkIs2386[unc-54p::α-synuclein::YFP; unc-119(+)]* animals fed with different bacteria. Aggregates at the head region of day one adults were shown. Scale bar = 20 μm. For image quantification, the number of fluorescent aggregates in the same head area were quantified manually for fifteen worms per group. Mean ± SD were shown. (B) Anti-α-syn antibody staining of UM10 *unkIs7[aex-3p::α-syn(A53T), dat-1p::gfp]* animals at L2 stage fed with WT or *csgA*(-) K12. Scale bar = 20 μm. (C) Anti-α-syn staining showed the α-syn aggregates (arrows) in the DA neurons of day two adults in UM6 *unkIs9 [dat-1p::α-syn(A53T), dat-1p::gfp]* animals. Animals fed with *csgA*(-) K12 showed less aggregation. GFP expressed from the *dat-1* promoter labels the DA neurons. Scale bar=10 μm. (D) Sequential fractionation of the lysate of NL5901 and UM10 animals fed with WT or *csgA*(-) K12 and different fractions were blotted by anti-α-syn antibodies in western blot assays. Relative intensity of different fractions was quantified using ImageJ and normalized to whole animal lysate (WL). Anti-tubulin and anti-GFP blotting were used as internal controls. Arrow and asterisk indicate intact and degraded α-syn::YFP, respectively. (E) Western blot of α-syn and tubulin in the lysate of UM10 animals fed with WT K12 and treated with proteasome inhibitors MG132 (11 μM) and Bortezomib (13 μM). (F) Percentage of UM10 animals with Non-Unc phenotype and two intact ADE neurons under the treatment of ubiquitination inhibitor PYR-41 (1.4 mM) or proteasome inhibitors MG132 (11 μM) and Bortezomib (13 μM). The mean ± SD of three independent experiments (20-30 animals were scored for each experiment) is shown.

Because the α-syn::YFP fusion may not accurately reflect the aggregation pattern of α-syn, we next directly stained α-syn expressed pan-neuronally or specifically in DA neurons using anti-α-syn antibodies. We found that the amount of α-syn proteins detected in *aex-3p::α-syn(A53T)* animals fed with *csgA(-)* bacteria was greatly reduced compared to the animals fed with wild-type K12 (Figure 3B). In animals carrying the *dat-1p::α-syn(A53T)* transgene and fed with wild-type K12, α-syn proteins were found as discrete aggregates in both the cytoplasm and axons of DA neurons. Feeding with *csgA(-)* bacteria, however, removed most of the aggregates and led to a diffusive pattern of α-syn (Figure 3C).

Biochemical analysis supported the immunofluorescence results. After sequentially fractionizing the lysate of animals expressing α-syn::YFP in the muscle, we found that the amount of α-syn::YFP was markedly reduced in the insoluble fraction (FA) in animals fed with *csgA(-)* K12 compared to the animals fed with wild-type K12 (Figure 3D). In contrast, α-syn::YFP level in the high-salt soluble (RAB) and detergent soluble (RIPA) fraction did not show much difference. In animals expressing α-syn(A53T) pan-neuronally, however, feeding with *csgA(-)* K12 led to the downregulation of the whole animal lysate and all three fractions (Figure 3D). This downregulation likely occurred at the protein level, because the α-syn mRNA level did not show any difference in animals fed with wild-type and *csgA(-)* K12 (Figure S2A) and treatment with proteasome inhibitors, such as MG132 and Bortezomib, restored the protein level of α-syn in *aex-3p::α-syn(A53T)* animals fed with *csgA(-)* K12 (Figure 3E). These data support that CsgA derived from the bacteria promotes the formation of insoluble α-syn aggregates that are not accessible by the proteasomal machinery. Importantly, treatment with ubiquitination or proteasome inhibitors could largely block the neuroprotective effects of deleting *csgA* in the bacteria (Figure 3F). Thus, in the absence of bacterial curli, neurons were capable of handling the pro-aggregating α-syn(A53T) proteins through the ubiquitination-proteasome system. Curli-induced α-syn aggregation may exacerbate the proteotoxic stress and overwhelm the proteasome system, leading to neurodegeneration.

### CsgA colocalizes with α-syn in muscles and neurons

We next tested whether bacteria curli can get into *C. elegans* tissues to promote α-syn aggregation through cross-seeding *in vivo*. According the theory of cross-seeding (Morales et al., 2013), we hypothesizes that CsgA serves as the seed to nucleate α-syn. As in the K12 strain, we engineered the *csgA* locus to fuse a C-terminal 3×FLAG tag in the UTI2 *E. coli* strain, which produce high levels of curli, and then feed this bacteria to animals expressing α-syn::YFP in the muscle. By staining the FLAG tag, we observed the colocalization of CsgA with α-syn::YFP puncta (Figure 4A). Strikingly, CsgA appeared to be located in the center of the α-syn::YFP aggregates, which is consistent with the notion that curli helps nucleate α-syn. To avoid possible complication with the α-syn::YFP fusion, we directly visualize the colocalization of α-syn and CsgA through immunofluorescence double staining in animals expressing α-syn(A53T) in the muscle (Figure 4B). In this case, CsgA and α-syn signal appeared to overlap completely.

**Figure 4.**
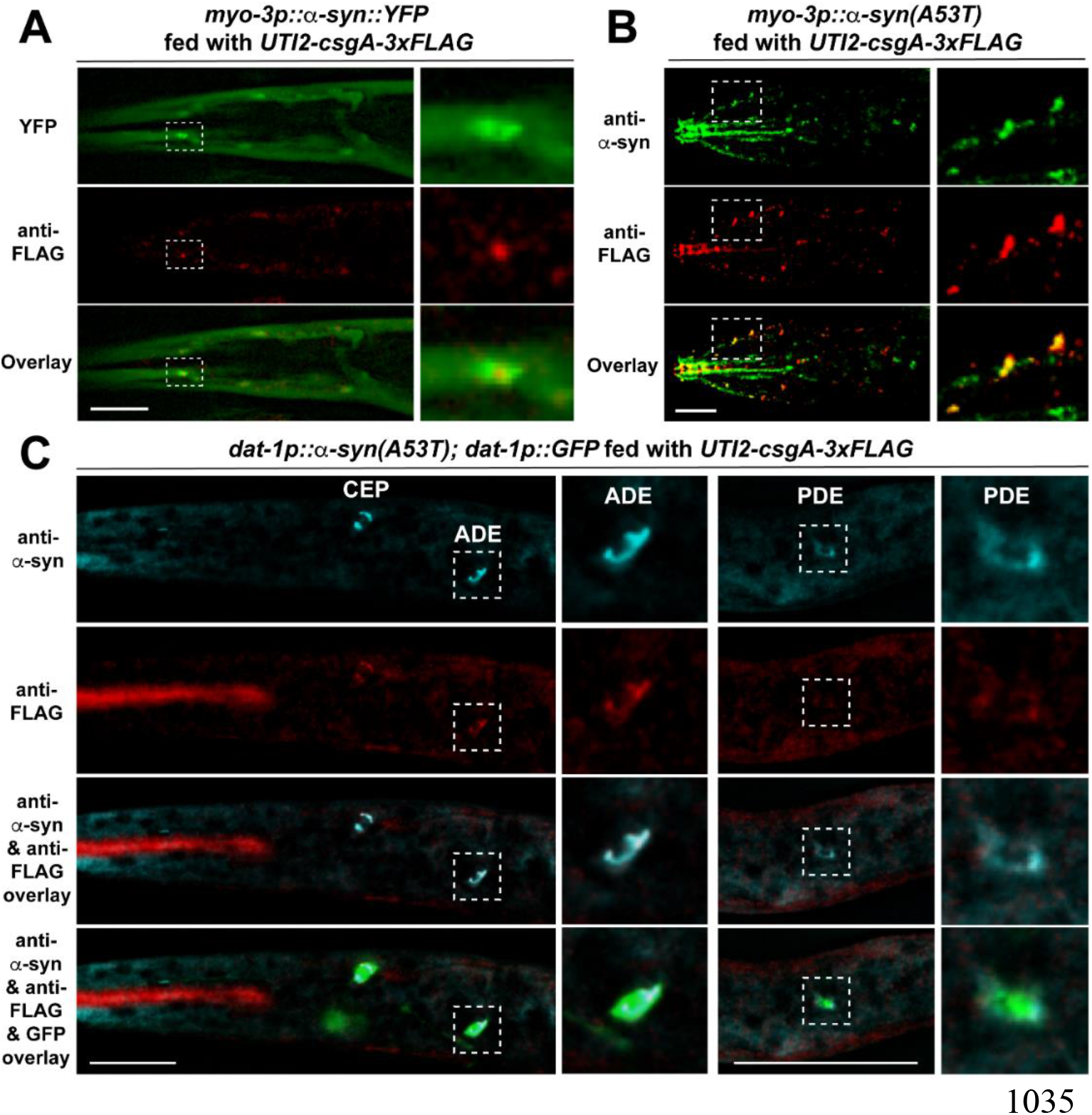
*CsgA* colocalizes with α-syn. (A) Day five adults of NL5901 *pkIs2386[unc-54p::α-synuclein::YFP; unc-119(+)]* animals fed with *UTI2-csgA-3xFLAG* bacteria were stained with anti-FLAG (red) antibodies. Insets show the enlarged region outline by the dashed box and indicate the colocalization of CsgA and α-syn. (B) Day five adults of CGZ512 *unkEx109[myo-3p::α-syn(A53T); dpy-5(+)]* animals fed with *UTI2-csgA-3xFLAG* bacteria were stained with anti-α-syn (green) and anti-FLAG (red) antibodies. Insets are enlarged images of the boxed regions. (C) Day one adults of UM6 *unkIs9[dat-1p::α-syn(A53T), dat-1p::gfp]* animals fed with *UTI2-csgA-3xFLAG* bacteria were stained with both anti-α-syn (cyan) and anti-FLAG (red) antibodies. GFP signal indicates the position of the DA neurons. Colocalization of CsgA and α-syn were observed in all three types of DA neurons, CEP, ADE, and PDE neurons. Inserts are enlarged images of the boxed region showing the colocalization in ADE and PDE neurons. Scale bar = 20 μm in all panels. Images were processed using the Leica THUNDER imaging system. The raw images can be found in Figure S5.

Colocalization of CsgA with α-syn was also observed in dopaminergic neurons of animals carrying the *dat-1p::α-syn(A53T)* transgene and fed with *UTI-2-csgA-3xFLAG* bacteria (Figure 4C). Importantly, colocalization was not only found in the CEP and ADE neurons, which are adjacent to the pharynx that grinds up the bacteria, but also found in PDE neurons that are located in the posterior half of the body. Thus, the CsgA proteins appeared to be transported inside the *C. elegans* (Figure 4C). Interestingly, CsgA signals were not observed in normal DA neurons that do not express α-syn, suggesting that the retention of bacterial curli may also depends on α-syn aggregation (Figure S3). These results support the mutual cross-seeding between CsgA and α-syn.

As controls for the above immunofluorescence experiments, we also fed PD animals with the wild-type UTI2 bacteria without the CsgA::3×FLAG fusion and did not observe any anti-FLAG signals (Figure S4). Deconvolution were used to analyse the above imaging data to increase clarity and to remove out-of-focus light. The raw unprocessed images showed similar colocalization patterns (Figure S5).

### Curli promotes α-syn-induced mitochondrial dysfunction and energy failure

We next conducted transcriptomics studies to investigate what molecular aspects of the α-syn neurodegenerative pathology can be rescued by deleting CsgA in the bacteria. Through RNA sequencing, we found 1274 genes downregulated (fold change > 2; adjusted *p* < 0.05) in animals expressing α-syn(A53T) pan-neuronally compared to wild-type animals, when both fed with the wild-type K12. Gene ontology analysis found that genes that function in the mitochondria and genes that regulate metabolic processes and energy production were enriched in the 1274 downregulated genes, which indicated that α-syn aggregation disrupted mitochondrial functions. Among these 1274 genes, 168 genes were upregulated when *aex-3p::α-syn(A53T)* animals were fed with *csgA(-)* K12 bacteria (Figure 5A and Table S1-2). Importantly, 84% (168/199) of the genes that were upregulated by feeding with *csgA(-)* K12 were genes downregulated by α-syn(A53T), and no significant transcriptomic changes were found between the wild-type animals fed with wild-type and *csgA(-)* K12 bacteria. Thus, promoting α-syn aggregation is likely the only activity of curli in the PD animals; and it does not seem to affect other aspects of normal *C. elegans* physiology.

**Figure 5.**
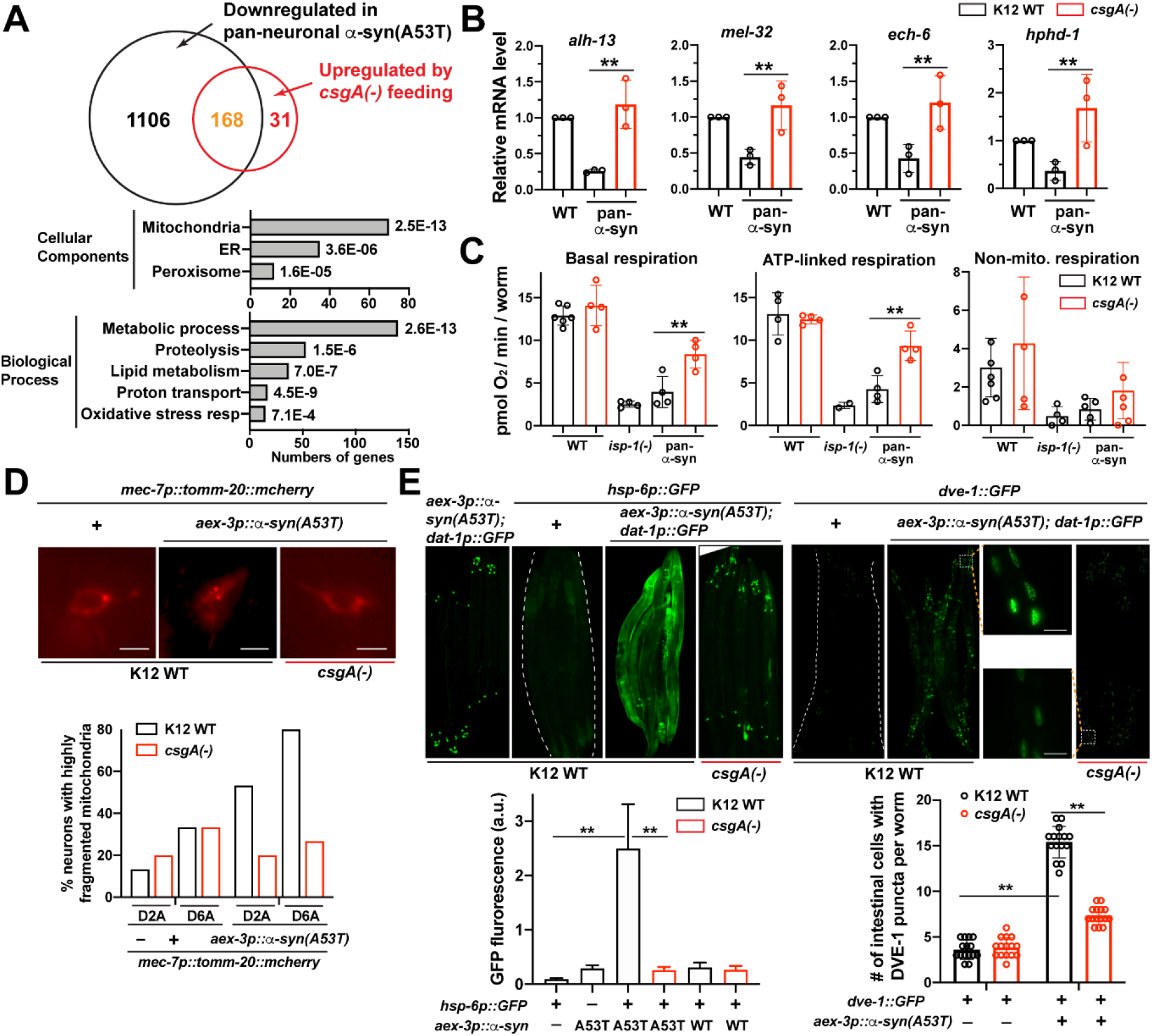
Bacterial curli promotes α-syn-induced mitochondrial dysfunction. (A) Venn diagram of genes that are downregulated in UM10 *unkIs7[aex-3p::*α*-syn(A53T), dat-1p::gfp]* animals compared to N2 control and genes that are upregulated in UM10 animals fed with *csgA(-)* K12 compared to animals fed with WT K12. Gene ontology analysis for down-regulated genes between N2 and UM10 using DAVID. (B) RT-qPCR measurement of mRNA level of mitochondrial genes *alh-13, mel-32, ech-6* and *hphd-1* in day one adults of UM10 animals fed with WT or *csgA(-)* K12 bacteria. Three biological replicates were performed and mean ± SD were shown. (C) Basal respiration, ATP-linked respiration, and Non-mitochondrial respiration (measured as oxygen consumption rate) of day one adults of N2, MQ887 *isp-1(qm150)*, and UM10 animals fed with WT or *csgA*(-) K12 bacteria. Representative results of 6~18 repeats for each condition were shown as mean ± SD. 70 to 160 animals were added to each microplate well. (D) Representative images of the ALM neurons in CGZ833 *unkIs7; twnEx8[mec-7p::tomm-20::mCherry; myo-2p::GFP]* animals fed with WT or *csgA*(-) K12. Scale bar=2 μm. Quantification shows the percentages of neurons with highly fragmented mitochondria in the soma in day two and day six adults. At least 20 animals were examined. (E) Representative images of *zcIs13[hsp-6::GFP]* and *zcIs39[dve-1p::GFP]* reporter expression in animals carrying *unkIs7[aex-3p::*α*-syn(A53T), dat-1p::gfp]* and fed with WT or *csgA*(-) K12 bacteria. Insets are enlarged images of the boxed region showing nuclear localization of DVE-1. Scale bar=25 μm. Mean ± SD were shown for the quantification of the *hsp-6p::GFP* intensity and the number of intestinal cells showing DVE-1::GFP nuclear puncta (20 animals were analysed for each experiment).

Focusing on the effects of curli on the mitochondria, we confirmed the downregulation of seven mitochondrial genes (*alh-13*, *acdh-1*, *bcat-1*, *ech-6*, *hach-1*, *hpdh-1*, and *mel-32*) in PD animals and their restored expression upon feeding with *csgA(-)* bacteria using RT-qPCR (Figure 5B and S2B). Some of these genes regulate mitochondrial cellular respiration. For example, *acdh-1* (a short/branched chain acyl-CoA dehydrogenase) and *ech-6* (a short chain enoyl-CoA hydratase) are involved in mitochondrial fatty acid β-oxidation, and *bcat-1*codes for a branched-chain amino acid (BCAA) aminotransferase that initiates the catabolism of BCAAs. Both β-oxidation and BCAA breakdown generate acetyl-CoA, which feeds into the tricarboxylic acid (TCA) cycle to produce NADH and FADH_2_, which are then supplied for the electron transport chain to produce ATP (Nolfi-Donegan et al., 2020). *hpdh-1* (a hydroxyacid-oxoacid transhydrogenase) directly functions in the TCA cycle and *mel-32* codes for a serine hydroxymethyltransferase, which is essential for maintaining mitochondrial respiration (Lucas et al., 2018). Interestingly, knockdown of *bcat-1* was found to promote neurodegeneration in PD models (Yao et al., 2018). In addition, the expression of lipid elongases (*elo-5*, *elo-6*, and *elo-9*) and acyl-coA oxidase (*acox-2* and *F08A8.4*), which are also known genetic modifiers of PD (Chen and Burgoyne, 2012; Lee et al., 2011), were downregulated in *aex-3p::α-syn(A53T)* animals and recovered upon feeding with *csgA(-)* bacteria (Table S1-2).

To test whether the alteration in genetic programs associated with mitochondrial activities and metabolism led to defects in energy production, we measured oxygen consumption rates in *C. elegans* using the Agilent Seahorse XFe24 analyser. We found that animals expressing α-syn(A53T) pan-neuronally had much lower basal and ATP-linked respiration than the wild-type animals. Feeding with *csgA(-)* bacteria strongly rescued cellular respiration (Figure 5C). These results support that CsgA-induced α-syn aggregation caused mitochondrial dysfunction and energy failure, leading to neuronal cell death.

We also visualized mitochondrial morphology using a strain expressing *tomm-20::mcherry* fusion in the touch receptor neurons. When fed with wild-type K12, the expression of α-syn(A53T) led to fragmentation of the mitochondria, whereas feeding with *csgA(-)* K12 could largely rescue the morphological defects of the mitochondria (Figure 5D). A well-known mitochondrial response to proteotoxic stress is the activation of mitoUPR (mitochondrial unfolded protein response) pathways (Anderson and Haynes, 2020). In *C. elegans*, mitochondrial stress in neurons can trigger mitoUPR in intestine through inter-tissue signalling (Zhang et al., 2018). As expected, we observed the activation of mitoUPR markers *hsp-6::GFP* and *dve-1::GFP* in the intestine of animals with pan-neuronal expression of α-syn(A53T); and the mitoUPR response was not engaged when the animals were fed with *csgA(-)* bacteria (Figure 5E). As controls, we found that the mitoUPR markers were not activated by the expression of wild-type α-syn and the endoplasmic reticulum (ER) UPR marker *hsp-4::GFP* were not activated by either wild-type or A53T α-syn proteins (data not shown). In fact, our transcriptomic analysis found that ER UPR genes (e.g. *hsp-3*, *apy-1*, and eight others) were enriched in genes downregulated by α-syn(A53T) overexpression (Table S1). Overall, our results indicate that bacterial curli is indispensable for the disruption of mitochondrial health by α-syn proteotoxicity.

### Bacterial curli promotes neurodegeneration induced by diverse protein aggregates in ALS, AD, and HD models

In addition to α-syn in PD models, we also examined whether CsgA promoted the neurodegeneration caused by other protein aggregates, e.g. SOD1 in Amyotrophic lateral sclerosis (ALS), amyloid β in Alzheimer’s disease (AD), and huntingtin in Huntington’s disease (HD). Using a *C. elegans* ALS model that express human SOD1(G85R)::YFP pan-neuronally (Wang et al., 2009), we found that, when the animals were fed with *csgA(-)* K12, the perinuclear accumulation of SOD1(G85R)::YFP aggregates largely disappeared in ALM neurons and the bright discrete puncta appeared more diffused and less aggregated in the ventral nerve cord motor neurons (Figure 6A). Thus, CsgA may promote SOD1 aggregation.

**Figure 6.**
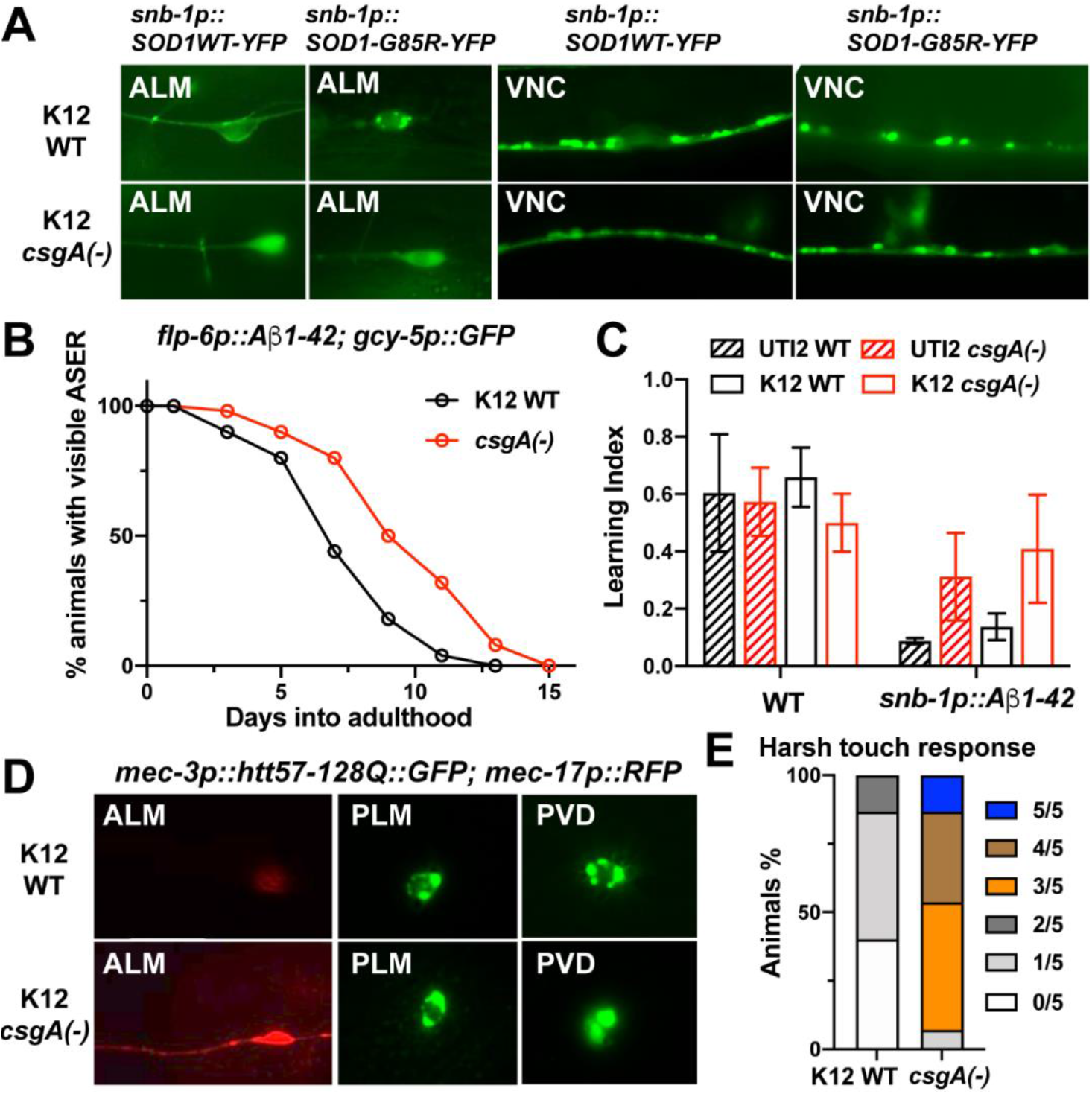
Bacterial curli promotes neurodegeneration in C. elegans models of ALS, AD, and HD. (A) Representative images of ALM neurons and ventral nerve cord (VNC) neurons in ALS strains carrying the transgene *snb-1p::SOD1(G85R)::YFP* or *snb-1p::SOD1(WT)::YFP* and fed with WT or *csgA*(-) K12. At least thirty day two adults were assessed. (B) The percentage of animals with visible ASER neurons in FDX25 *sesIs25[flp-6p::Aβ1-42; gcy-5p::GFP; rol-6(D)]* animals fed with WT or *csgA*(-) K12. For each condition, 50 animals were scored. (C) Learning index of day one adults of CL2355 *smg-1(cc546) dvIs50 [snb-1p::Aβ1-42::3’UTR; mtl-2::GFP]* animals grown on WT and *csgA(-)* K12 and WT and *csgA(-)* UTI2 bacteria in an associative learning assay. Two to four hundred animals were used in each experiment; 3 biological replicates and 3 technical replicates were performed. Mean±SD is shown. (D) Morphological changes of the ALM neurons and the alteration of Huntingtin (Htt) aggregation pattern in PLM and PVD neurons in TU6295 *uIs115[mec-17p::TagRFP]; igIs5[mec-3p::htt57-128Q::GFP; lin-15(+)]* animals fed with *csgA(-)* K12 compared to animals fed with WT K12. Twenty to thirty day one adults were assessed for each condition. (E) Harsh touch sensitivity of TU6295 animals fed with WT or *csgA*(-) K12 bacteria.

For AD models, we employed two *C. elegans* strains that expressed Aβ1-42 either in a few pairs of sensory neurons (using *flp-6* promoter) (Melentijevic et al., 2017) or pan-neuronally (using *snb-1* promoter) (Wu et al., 2006). In the first model, we observed increased ASE neuron survival when the animals were fed with *csgA(-)* K12 instead of wild-type K12 (Figure 6B). In the second case, we observed restored butanone associative learning upon the feeding with *csgA(-)* bacteria (Figure 6C). Thus, eliminating bacteria curli could partially suppress Aβ-induced neurodegeneration.

For HD models, we fed the animals expressing htt57-128Q::GFP fusion in the mechanosensory neurons (using *mec-3* promoter) with either wild-type or *csgA(-)* K12. The discrete perinuclear clusters of htt57-128Q::GFP signals became more diffused in animals fed with *csgA(-)* K12 and the degeneration of ALM neurons were also suppressed (Figure 6D). As functional output, we tested harsh touch sensed by the PVD neurons and found the percentage of response increased dramatically in the HD animals fed with *csgA(-)* K12, suggesting improved PVD functions (Figure 6E). Nevertheless, we did not observe significant improvement in gentle touch response mediated by ALM and PLM neurons upon feeding with *csgA(-)* K12.

The above data expanded the pro-neurodegenerative role of bacteria curli and suggested that CsgA may cross-seed not only α-syn but also a wide range of other aggregation-prone proteins, including SOD1, Aβ, and polyQ-expanded huntingtin. Targeting CsgA may be generally effective in reducing neurodegeneration in many neurodegenerative diseases.

### CsgA-derived amyloidogenic peptides cross-seed α-syn and induce neuronal death in human cells

Finally, we tested the *in vivo* cross-seeding of CsgA and α-syn in human neuroblastoma SH-SY5Y cells. After transfecting the SH-SY5Y cells with plasmids expressing α-syn wild-type or A53T, we treated the cells with a CsgA-derived amyloidogenic hexapeptides (N’-QYGGNN-C’) or a non-amyloidogenic control (N’-QYGGNA-C’) (Tukel et al., 2009). We found that the CsgA amyloidogenic peptides significantly enhanced α-syn expression and aggregation in the SH-SY5Y cells (Figure 7A and B) and expression of α-syn or EGFP::α-syn fusion facilitated the accumulation of rhodamine-conjugated CsgA peptides (Figure 7C and S6A). In fact, without the expression of α-syn, the peptides cannot be retained in the SH-SY5Y cells. Thus, the cross-seeding and mutual facilitation of aggregation between CsgA and α-syn observed in *C. elegans* also occurred in human cells.

**Figure 7.**
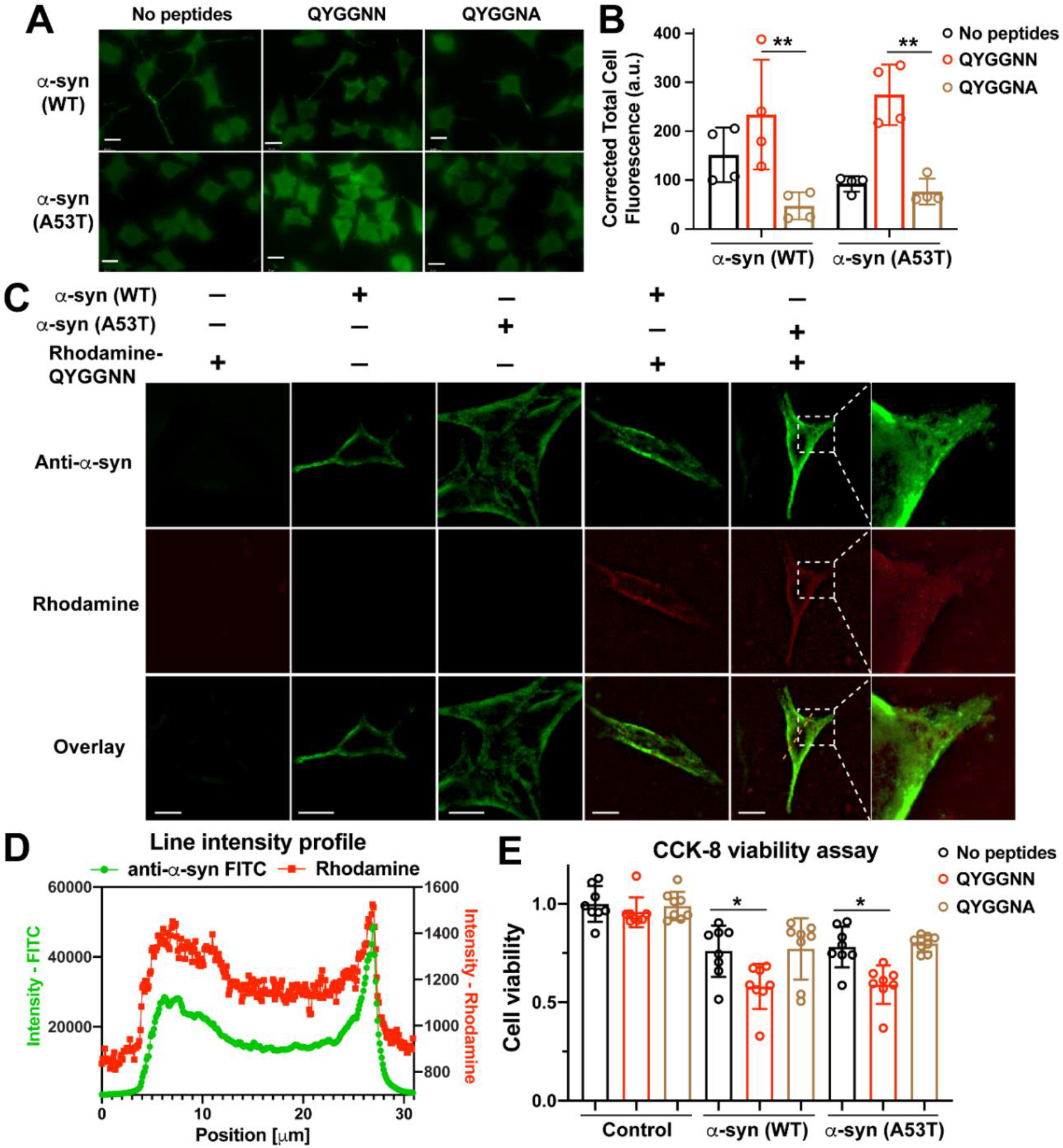
*CsgA*-derived amyloidogenic peptides cross-seed α-syn and induce neuronal death in human cells. (A) Representative anti-α-syn immunofluorescent images of SH-SY5Y cells transfected with α-syn(WT or A53T)-expressing constructs and then treated with *CsgA*-derived amyloidogenic hexapeptides, non-amyloidogenic control or empty vehicle. Scale bar = 20 μm in all panels in this Figure. (B) Corrected total cell fluorescence (CTCF shown as mean±SD) from the experiments shown in (A). Amyloidogenic peptides significantly enhanced α-syn expression and accumulation in the SH-SY5Y cells. (C) SH-SY5Y cells transfected with α-syn(WT or A53T)-expressing constructs and treated with Rhodamine-conjugated QYGGNN peptides (red) were then stained with anti-α-syn antibodies (green) to show colocalization of CsgA-derived peptides and α-syn. Insets are enlarged images of the boxed regions. Images were processed using the Leica THUNDER imaging system. The raw images can be found in Figure S6B. (D) Intensity profile for the orange dashed line in (C). (E) Representative results of CCK-8 cell viability assays using α-syn(WT or A53T)-expressing SH-SY5Y cells treated with amyloidogenic hexapeptides, non-amyloidogenic control or empty vehicle. Mean±SD is shown and single asterisk indicates *p* < 0.05 in a one-way ANOVA followed by a Tukey’s post-hoc test.

Using the rhodamine-QYGGNN peptide and antibody staining against α-syn, we directly observed the colocalization of CsgA-derived peptides with both α-syn wild-type and A53T proteins in SH-SY5Y cells (Figure 7C-D and S6B). Both α-syn and CsgA appeared to be enriched in the periphery of the cells. α-syn(A53T) also showed stronger signal than wild-type α-syn in the presence of CsgA peptides. To assess neuronal cell death, we carried out a cell viability assay on SH-SY5Y cells transfected with α-syn-expressing constructs and treated with the CsgA peptides. We found that the amyloidogenic QYGGNN but not the control peptide strongly exacerbated cell death induced by α-syn proteins (Figure 7E). Treatment of the QYGGNN peptide alone in cells that express no α-syn did not affect cell survival. Thus, bacterial curli may enhance the α-syn aggregates-induced degeneration of human neurons.

## Discussion

### Genome-wide screen reveals pro-neurodegenerative factors in bacteria

In this study, we conducted a genome-wide screen for pro-neurodegenerative factors in bacteria and identified 38 *E. coli* genes that promote α-syn-induced neurodegeneration in a *C. elegans* model of PD. The design of our screen ensured low level of false positives, since two independent PD models were used and two independent phenotypes (locomotion defects caused by motor neuron degeneration and the loss of fluorescently labelled dopaminergic neurons) were scored (Figure 1A). Nevertheless, we expect relatively high false negative rate, since our screen strategy biased towards the positive hits of the beginning rounds. Despite that, the final 38 positive *E. coli* genes clearly converged on several genetic pathways that likely play important roles in mediating bacteria-host interactions in PD pathogenesis. Besides the curli genes, *csgA* and *csgB*, which are the focus of this study, we also identified five genes involved in LPS production and assembly, three genes involved in the biosynthesis of adenosylcobalamine, two genes that code for inhibitors of eukaryotic lysozymes, six genes involved in oxidative stress response, and eight genes involved in metabolism and energy homeostasis, among others. These results not only confirmed some previous hypotheses about the gut microbiota-brain interactions in PD but also offered new insights into the process.

For example, intact LPS triggers innate immunity in both *C. elegans* and humans and the resulted neuroinflammatory response has detrimental effects on PD pathology (Aballay et al., 2003; Kelly et al., 2014). Consistent with this notion, we observed that the disruption of LPS assembly in *E. coli* mutants, such as *lapA(-)* and *lapB(-)*, alleviated neurodegeneration in *C. elegans*. The receptor for LPS in C*. elegans* is, however, unclear, since the presumptive Toll-like receptor signalling pathway does not mediate LPS toxicity (Aballay et al., 2003). Thus, LPS may promote neurodegeneration through diverse molecule mechanisms in different organisms.

Deletions of *E. coli* genes (*cobS*, *btuR*, and *eutT*) involved in the biosynthesis of adenosyl-cobalamin (AdoCbl), one active form of vitamin B12, also suppressed neurodegeneration in *C. elegans*. Clinical studies suggested that vitamin B12 insufficiency may be a contributing factor for the cognitive impairment and rapid disease progression in PD (McCarter et al., 2019). So, our results were quite unexpected. One hypothesis is that blocking AboCbl synthesis in *E. coli* might lead to increased production of methylcobalamin (MeCbl), another form of B12 that may be beneficial for preventing neurodegeneration. An alternative hypothesis is that AboCbl-dependent metabolic pathways in *C. elegans* might promote degeneration. In either case, given that vitamin B12 is synthesized by the gut bacteria in humans, the *C. elegans* model can be instrumental to understand the roles of AdoCbl and MeCbl in the microbial regulation of neuronal health.

Among the other positive hits, genes (e.g. *sodA, yaaA*, *msrA*, and *nrfA*) that help *E. coli* cope with oxidative stress, genes (*ldhA* and *lldD*) that mediate the conversion of lactate into pyruvate, and genes (*pck* and *tpiA*) that mediate key steps in gluconeogenesis also promoted α-syn-induced neurodegeneration. Although the mechanisms by which these bacterial genes regulate host neurodegeneration await further investigation, our systematic screen successfully identified several possible routes of communication between the bacteria and host neurons.

### Cross-seeding between bacteria curli and pathologically aggregated proteins in neurons

Cross-seeding refers to the process that oligomers composed by one type of misfolded proteins can promote the polymerization of another (Morales et al., 2013). Previous studies have observed the cross-seeding between Aβ and other misfolded proteins, including prion, tau, and α-syn (Mandal et al., 2006; Morales et al., 2010). Bacterial curli, made of the major subunit CsgA and the minor subunit CsgB, is a type of amyloid fibril with cross β-sheet structure. Since curli fibers from different bacterial species are able to cross-seed *in vitro* and facilitate multispecies biofilm formation (Zhou et al., 2012), it has been hypothesized that curli secreted by the gut bacteria may also cross-seed with Aβ or α-syn and thus promote neurodegeneration. Direct evidence for this hypothesis was missing until recently.

While we are conducting this study, Sampson et al. (2020) reported the *in vitro* cross-seeding between and CsgA and α-syn and showed that purified CsgA accelerates α-syn fibrilization. To extend their *in vitro* studies, we directly visualized the colocalization of CsgA and α-syn *in vivo* in *C. elegans* neurons and human neuroblastoma cells at single-cell resolution. Importantly, previous studies mostly used Congo red staining to visualize bacteria curli, which may be problematic because Congo red stains not only curli but many other types of amyloid deposition (Ho et al., 2014). In this study, we tagged CsgA with a FLAG-tag and were able to clearly demonstrate the presence of curli and their colocalization with α-syn inside neurons. Moreover, we also showed that the cross-seeding is bidirectional. In both *C. elegans* neurons and human SH-SY5Y cells, CsgA promoted α-syn aggregation and α-syn facilitated the retention of CsgA. Similar bidirectional interaction was also observed between Aβ and prions and α-syn (Morales et al., 2009).

In addition to the cross-seeding between curli and α-syn, we also provided evidence to show that CsgA promoted neurodegeneration caused by the aggregation of Aβ, SOD1, and polyQ-expanded Huntingtin in *C. elegans* models of AD, ALS, and HD, respectively. Therefore, the bacteria-secreted curli may have detrimental effects in a range of neurodegenerative disorders. Targeting curli production in the gut may represent a general therapeutic approach to prevent or slow down the progression of the protein aggregation diseases. In this study, we showed that blocking curli production in bacteria using EGCG, a green tea extract, has remarkable effects in preventing neurodegeneration, suggesting that pharmacological inhibition of curli may be an effective treatment for PD and likely other neurodegenerative diseases. It is encouraging that in a recent study using a mice PD model, Sampson *et al.* (2020) found that colonization of mouse gut with curli-producing *E. coli* also promoted α-syn pathology in the brain and exacerbated motor impairment. Thus, the effects of curli on neurodegeneration appeared to be consistent across different disease models in diverse organisms.

### Bacteria-host interactions regulate neuronal mitochondrial functions

Mitochondrial dysfunction is a hallmark of PD pathogenesis and α-syn aggregation and mitochondrial damage exacerbate each other through a vicious cycle (Poewe et al., 2017). Consistent with this idea, our transcriptomic studies found that α-syn(A53T) overexpression downregulated genes that function in mitochondria, lipid metabolism, and ATP production. Eliminating bacterial curli restored the expression of a subset of these genes, which may be the key regulatory points in the metabolic network disrupted in PD. For example, mitochondrial genes *acdh-1*, *bcat-1*, *ech-6*, and *hpdh-1*, which code for various enzymes critical in BCAA catabolism, fatty acid β-oxidation, or TCA cycle, were reactivated in PD animals fed with the *csgA(-)* bacteria. This correction in transcriptional program is accompanied by restored mitochondrial morphology, blocked mitoUPR activation, and revived cellular respiration. Thus, removing curli from the bacteria reduced α-syn proteotoxicity and rescued neurons from mitochondrial dysfunction and energy failure, which may be the key in preventing neuronal death.

Supporting the regulation of key metabolic genes in PD, *bcat-1* was recently identified as a PD-associated gene (Yao et al., 2018). Human BCAT-1 is highly expressed in the substantia nigra of healthy individuals, and its expression is significantly diminished in PD patients; knockdown of *bcat-1* in *C. elegans* neurons recapitulated aging phenotypes and enhanced α-syn-induced neurodegeneration. Since BCAT-1 is the aminotransferase that catalyzes the initial step of BCAA breakdown, downregulation of *bcat-1* leads to not only reduced energy production but also increased BCAA levels, which appeared to correlate with disease severity in PD patients (Luan et al., 2015). In another example, impaired fatty acid β-oxidation is associated with the energy crisis in PD and the levels of short chain 3-hydroxyacyl-CoA dehydrogenase (SCHAD), a key enzyme in β-oxidation, are significantly reduced in the ventral midbrain of both PD mice and PD patients (Przedborski et al., 2004). Overexpression of SCHAD mitigates the impairment of oxidative phosphorylation and ATP production in PD models (Tieu et al., 2004). Our work found that *acdh-1* (a short/branched chain acyl-CoA dehydrogenase, homolog of human ACADSB) and *ech-6* (a short chain enoyl-CoA hydratase, homolog of human ECHS1), two other essential enzymes in β-oxidation, were also downregulated in PD but recovered after eliminating bacteria curli. Restoration of the expression of these metabolic genes may be critical for restoring mitochondrial respiration and energy production at the cellular level.

Furthermore, we found that in the absence of curli, the reduced α-syn aggregation could be coped with by the ubiquitination-proteasome system, which protect the neurons from toxic oligomers-induced mitochondrial dysfunction and from cell death. In fact, our transcriptomic analysis found that many proteolytic genes (e.g. aspartyl protease *asp-8*, prolyl carboxypeptidase *pcp-4*, metalloprotease *nep-22*, etc) were downregulated in PD animals and recovered when feeding with *csgA(-)* bacteria (Figure 5A and Table S1-2). The reactivation of the proteolytic machineries after removing curli may have allowed the neurons to curb α-syn aggregation and maintain the oligomers at low levels. Nevertheless, only 168 (13%) of the 1274 genes downregulated in PD animals were rescued after deleting curli in the *E. coli*, suggesting that many transcriptional change may still be induced by low levels of α-syn aggregates independent of curli.

## STAR Methods

### ❖ *C. elegans* Strains

*Caenorhabditis elegans* wild-type (N2) and mutant strains were maintained at 20°C as previously described (Brenner, 1974). PD-related strains UM{Schneider, 2012 #60}9 *unkIs11[dat-1p::GFP]*, UM10 *unkIs7[aex-3p::α-syn(A53T), dat-1p::gfp]*, UM11 *unkIs8[aex-3p::α-syn(WT), dat-1p::gfp]*, UM3 *unkIs10 [dat-1p::α-syn(WT), dat-1p::gfp]*, and UM6 *unkIs9 [dat-1p::α-syn(A53T), dat-1p::gfp]* were generous gifts from Garry Wong at the University of Macau. ALS strains carrying the transgenes *snb-1p::SOD1(G85R)::YFP]* and *snb-1p::SOD1(WT)::YFP* were kindly provided by Jiou Wang at Johns Hopkins University. The AD strains carrying *sesIs25 [flp-6p::Abeta1-42; gcy-5p::GFP;rol-6]* were kindly provided by Monica Driscoll at Rutgers University. CGZ512 *dpy-5(e907); unkEx109[myo-3p::α-syn(A53T); dpy-5(+)]* were generated in this study through microinjections. The *myo-3p::α-syn(A53T)* constructs were created by swapping the *dat-1* promoter in the *dat-1p::α-syn(A53T)* construct provided by Garry Wong with a 2.5 kb *myo-3* promoter.

Another PD strain NL5901 *pkIs2386[unc-54p::α-synuclein::YFP; unc-119(+)]*, the AD strain CL2355 *smg-1(cc546); dvIs50[pCL45 (snb-1::Aβ1-42::3’UTR(long); mtl-2:: GFP]*, the HD strain ID5 *igIs5 [mec-3p::htt57-128Q::GFP; lin-15(+)]*, the mitochondrial marker CLP215 *twnEx8[mec-7p::tomm-20::mCherry; myo-2p::GFP]*, and the mitoUPR reporters SJ4100 *zcIs13[hsp-6::GFP]* and SJ4197 *zcIs39 [dve-1p::dve-1::GFP]* were provided by the Caenorhabditis Genetics Center.

### ❖ Keio library screens, *E. coli* genetic engineering, and Congo red staining

The Keio library (Baba et al., 2006) were purchased from the Dharmocon (Colorado, United States). *E. coli* knockout clones from the library were grown overnight at 37°C in LB medium with 50 μg/ml kanamycin in 96-deep well plates. 20 μL of the overnight culture was seeded onto NGM agar-containing 96-well plates. For locomotion screens, about 20 synchronized first-stage larva (L1) of the UM10 strain were added into each well and the animals were grown for 48 hours at 20°C before scored for the penetrance of non-Unc phenotype at the L4 stage. *E. coli* mutants that caused at least 25% of the animals in the well to be non-Unc were considered positive hits. Animals fed with the parental wild-type K12 (BW25113) served as the negative control. For the ADE survival screen, bacterial culture was seeded on a 5 cm petri dish containing NGM and about 20 L1 animals of either UM10 or UM6 strains were added to the plate. Animals were screened as day-two adults for the percentage of animals showing two ADE neurons that clearly expressed GFP and showed normal cell morphology.

To delete *csgA* gene in the uropathogenic *E. coli* strain UTI2 (provided by Dr. Aixin Yan at the University of Hong Kong) and insert DNA fragment coding the 3xFLAG tag in the endogenous *csgA* locus in K12 and UTI2, we performed lambda red recombineering using a previously described method (Datsenko and Wanner, 2000). For deleting *csgA*, we used primers carrying ~50 bp *csgA*-flanking sequences to amplify a chloramphenicol (Cm) resistance gene. For inserting 3xFLAG, we amplified the Cm-resistance gene using primers containing both the FLAG tag and flanking sequences. Primer sequences can be found in the supplemental materials. *E. coli* K12 or UTI2 transformed with the helper plasmids pKD46 and pTNT was grown until OD600 reaches 0.4-0.6 in the presence of L-Arabinose at 30°C and washed with ice-cold 10% glycerol to make electrocompetent cells, which were then mixed with the purified PCR products and electroporated at 1.8 kV. Cells were then plated on LB agar plates containing Cm. Colonies were verified by colony PCR and Sanger sequencing.

For Congo red staining, the indicator plates were made of YESCA (1 g yeast extract, 10 g casamino acids, and 20 g agar per liter) media with 50 μg/mL Congo Red and 10 μg/mL Coomassie Brilliant Blue. 10 μl of overnight bacteria culture was seeded onto plates and cultured at 25°C for 48 h prior to assessing the curli production.

### ❖ Biochemical analysis

To measure the level of α-syn in high-salt soluble, detergent soluble and insoluble fractions, sequential fractionation was performed according to previously described methods (Fatouros et al., 2012). Briefly, synchronized worms were washed off NGM plates with M9 buffer. Dead animals and bacteria were removed by flotation on a 30% sucrose solution. The entire extraction procedure was carried out on ice and centrifugation steps were performed at 4°C except for the last step with 30% FA, which was performed at room temperature. To extract different fractions, worm pellets were directly resuspended in an equal amount (w/v) of high-salt RAB buffer [100 mM 2-(N-morpholino) ethanesulfonic acid (MES), 1 mM EGTA, 0.5 mM MgSO4, 20 mM NaF] and then were lysed by sonication (6 × 10 s, 10 s break). Homogenates were centrifuged at 40,000×g for 40 min. The supernatant constitutes the RAB fraction. The pellet was re-extracted with 1 M sucrose in RAB buffer and centrifuged for 20 min at 40,000×g, and the supernatant was discarded. The pellet was then extracted with RIPA buffer (150 mM NaCl, 1% Nonidet P-40, 0.5% deoxycholate, 0.1% SDS, 50 mM Tris, pH 8.0) and centrifuged at 40,000×g for 20 min. The supernatant is the RIPA fraction. The pellet, after a brief washing with RIPA buffer, was extracted with 30% FA and centrifuged at 13,000×g for 15 min. The supernatant is the FA fraction. All buffers contained the Roche cOmplete Protease Inhibitor cocktail and 0.5 mM PMSF (Sigma-Aldrich). The pH of FA fraction was adjusted by 5M NaOH. The RAB, RIPA and FA fraction were loaded to SDS-PAGE gel and probed with anti-α-synuclein (ab27766, Abcam; 1:1000 dilution) and anti-tubulin (ab76286, Abcam; 1:2000 dilution) antibodies.

### ❖ RNA-seq and RT-qPCR

Total RNA from *C. elegans* was extracted using TRIzol reagent (Thermo Fisher). Samples were sent to BGI (Beijing Genome Institutes) Hong Kong for standard library construction and pair-end sequencing. Around 20 million reads were obtained for each sample and the reads were aligned to the C. elegans genome (WS235) using STAR 2.7. To identify genes differentially expressed, the transcript counts were analyzed using DESeq2, and genes with false discovery rate–corrected *p* values (q values) below 0.05 and fold change above 2 or below 0.5 were identified. Gene enrichment analysis was then performed on the genes showing significant expression change using the DAVID functional annotation tool 6.8. The raw RNA-seq data can be accessed through https://www.ncbi.nlm.nih.gov/geo/query/acc.cgi?acc=GSE169204 in GEO database.

For transcriptional analysis, cDNA was reversed transcribed from total RNA using PrimeScript RT reagent kit (Takara) from synchronized L4 animals. Quantitative real-time PCR was performed using a TB Green Premix Ex Taq kit (Takara) in CFX96 real-time PCR machine (BioRad). Values were normalized to the internal control *tba-1*. All data shown represent the average of three biologically independent replicates. The qPCR primers are listed in supplementary Table S3.

### ❖ Antibody staining and fluorescent imaging

The antibody staining was performed using the Ruvkun protocol previously described (Finney and Ruvkun, 1990). *C. elegans* were fixed in fixation buffer with 2% formaldehyde in liquid nitrogen and several freeze-and-thaw cycles were carried out to break the cuticle. To detect synuclein and FLAG simultaneously, the mouse monoclonal anti-α-syn (ab27766, Abcam; 1:500 dilution) and rabbit polyclonal anti-FLAG antibodies (F7425, Sigma; 1:500 dilution) were added to the worm suspension and the incubation were carried out overnight on a rotating platform. Alexa 488-conjugated goat anti-mouse IgG antibody (115-545-003, Jackson Lab; 1:1000 dilution) and rhodamine-conjugated goat anti-rabbit IgG antibody (111-025-003, Jackson Lab; 1:1000) were used as secondary antibodies. After washing, worms were mounted on agarose pads for visualization.

Fluorescent imaging was done on a Leica DMi8 inverted microscope equipped with a Leica DFC7000 GT or K5 monochrome camera. Measurements were made using the Leica Application Suite X (3.7.0.20979) software. The Leica THUNDER deconvolution imaging system was used to process the colocalization images to increase the clarity of the pictures and remove out-of-focus light. The raw images without processing can be found in the supplemental materials.

### ❖ Human cell culture, transfection, and cell viability assay

Human neuroblastoma SH-SY5Y cells were maintained in DMEM supplemented with Fetal Bovine Serum (10% v/v), penicillin (100 IU/mL) and streptomycin (100 μg/mL) in humidified 5% CO2 at 37°C. The plasmids pHM6-α-syn(WT) (#40824), pHM6-α-syn(A53T) (#40825), EGFP-α-syn(WT) (#40822), EGFP-α-syn(A53T) (#40823) used for transfection were obtained from Addgene. 0.5 μg DNA was combined with lipofectamine reagent (Invitrogen Inc., USA) for 30 min in serum-free DMEM medium and the transfection mixture was incubated with the cells for 6 h. The DMEM medium containing 20% FBS and synthetic peptides (QYGGNN and QYGGNA synthesized by Ontores Biotechnology, Shanghai, China) were then added to the cells without removing the transfection mixture.

For staining of the SH-SY5Y cells, the cells were seeded on coverslip coated with poly-D-lysine and then transfected and treated with peptides. Cells were then fixed in 4% paraformaldehyde, washed with PBST, blocked in 4% BSA blocking solution, and then stained with anti-α-syn mouse monoclonal antibody (ab27766, Abcam) overnight. After washing with PBST for 3 times, cells on the coverslip were incubated with FITC-conjugated goat-anti-mouse secondary antibody diluted 1:5000 in blocking solution. After washing, the coverslip was mounted and imaged. The fluorescence intensity was analysed using ImageJ (Schneider et al., 2012) and corrected total cell fluorescence (CTCF) was calculated as CTCF = Integrated Intensity – (Area of selected cell × Mean background fluorescence).

Cell viability was determined using the cell counting kit 8 (CCK-8, Abcam). Transfected α-syn-overexpressing SH-SY5Y cells were cultured with or without synthetic peptides in 96-well plates at a density of 1×10 ^4^ cells/well and grown for 24 h. 10 μL of CCK-8 solution was added to each well and the absorbance was measured at 460 nm on a plate reader (SpectraFluor, Tecan) after 1 h.

### ❖ *C. elegans* behavioral assays and statistical analysis

The basal slowing response assay was performed according to previous methods (Sawin et al., 2000). Animals were grown from eggs on different bacteria diet. Well-fed young adults were transferred to unseeded 6 cm NGM plate or plates with regular OP50. Worms were allowed to acclimate to the assay plates for 5 minutes, and then the number of body bends/20 seconds was counted for each animal. For diet switching assays, worms were washed with M9 twice and then incubated in M9 with ampicillin and kanamycin for 1 hour before the switch to remove residual bacteria on the surface. For the harsh touch assays, the anterior half of the animal was touched by a platinum wire (20.3 μm in diameter, THOMAS) in a top-down manner. Each worm was touched 5 times with a 2-minutes inter-trial interval. A positive touch response resulted in animals backing away from the touch.

The butanone associative learning assay was performed according to previous methods (Kauffman et al., 2011). Young adults were washed in M9 and collected by gravity sedimentation in a 15 mL conical tube to remove bacteria. Some of the animals were transferred immediately to the chemotaxis plate for assaying for the naïve condition. The rest was starved in M9 for 1h. After the starvation, animals were conditioned for 1h in a OP50 seeded NGM plate with 2 μL streaks of 10% butanone diluted in absolute ethanol on the lid. The conditioned worms were washed with M9 and transferred to the chemotaxis plate. For the chemotaxis assay, 10 cm unseeded NGM plates were used 629 and three circles (1 cm in diameter of 1 cm) on the bottom and both sides of the plate were marked. Animals were place onto the bottom spot. 1 μL of 1 M sodium azide and 1 μL of 0.01% of butanone were added onto the left spot, and sodium azide and pure ethanol control were added onto the right spot. After one hour, the number of worms located in the butanone and ethanol spots, as well as at the original bottom spots were counted. Chemotaxis index (CI) and Learning index (LI) were calculated as Chemotaxis index (CI): [N (butanone) − N (ethanol)] / [Total – N (origin)]; Learning index (LI): CI (butanone) – CI (naïve). Each chemotaxis assay was performed in technical triplicate and contained about 200–400 worms. Each bacteria diet treatment was performed in three biological replicates.

For mitochondrial cellular respiration assay, oxygen consumption rate (OCR) was measured using the Agilent Seahorse XFe24 technology (Luz et al., 2015). Briefly, the sensor cartridges were hydrated in Seahorse calibration buffer at 25°C one night before the experiment. Day-one adults were washed with complete K-medium and suspended in 525 μL of unbuffered EPA water. About 50-100 animals were added into each well of the Agilent Seahorse microplate. To achieve the equivalent volume after compound injection, we loaded 75 μL of 160 μM DCCD (8×), 225 μM FCCP (9× concentrated) and 100 mM sodium azide (10×) for Port A, B and C. The basal respiration was measured first in every well, followed by injection of FCCP in half of the wells to uncouple mitochondria or DCCD in the other half of the wells to inhibit ATP synthesis. The sodium azide was injected at the end of the assay in every well to completely block mitochondrial respiration. The number of measurements for basal respiration, DCCD response, FCCP response and sodium azide response were set to 8, 14, 8 and 4, respectively. All respiration parameters were normalized to the number of animals per well. ATP-linked respiration is calculated by subtracting DCCD response from the basal OCR. Non-mitochondrial respiration is defined as the OCR after the azide treatment.

All quantitative data were shown as mean ± SD. For statistical analysis, we used one-way ANOVA followed by a Tukey HSD post-hoc test to compare different treatments in a multiple comparison. Two-tailed Student’s t-test was used to compare two groups. Differences were considered significant at *p* < 0.05. Double asterisks in figures indicate *p* < 0.01. All statistical analysis was carried out using GraphPad Prism 8.0 software (GraphPad Software, San Diego, CA, USA).

## Supporting information

key resources table

## Acknowledgements

We thank Garry Wong, Jiou Wang, and Monica Driscoll for sharing strains. We also thank Dr. Aixin Yan, Dr. Jetty Lee, and Prof. Xiang Li at the University of Hong Kong for sharing equipment and reagents. Some strains used in this study were provided by the Caenorhabditis Genetics Center, which is funded by the NIH Office of Research Infrastructure Programs (P40 OD010440). This work was supported by grants from the Research Grant Council of Hong Kong (ECS 27104219 and CRF C7026-20G to C.Z.), the Food and Health Bureau of Hong Kong (HMRF 07183186 to C.Z.), and the University of Hong Kong (seed fund 201910159087 to C.Z.).

## Author Contributions

C.W. performed all experiments with technical help from C.Z. in microinjection. C.Y.L and F.M. performed bioinformatic analysis on the RNA-seq data. C.Z. and C.W. prepared the figures and wrote the paper. C.Z. conceived the project, secured the funding, and supervised the experiments.

## Notes

### Competing Interest Statement

The authors have declared no competing interest.

