## supplementary materials for "Genome-wide screen identifies curli amyloid fibril as a bacterial component promoting host neurodegeneration"

**Figure S1. Congo red staining of bacteria related to Figure 2.**
**Figure S2. RT-qPCR analysis of mitochondrial genes in PD animals, related to Figure 5.**
**Figure S3. CsgA retention by DA neurons depends on the expression of α-syn, related to Figure 4.**
**Figure S4. Control experiments for the colocalization between CsgA and α-syn, related to Figure 4.**
**Figure S5. Unprocessed images of CsgA colocalization with α-syn in *C. elegans* muscle and neurons, related to Figure 4.**
**Figure S6. CsgA-derived amyloidogenic peptides were retained in SH-SY5Y cells expressing α-syn, related to Figure 7.**

**Table S1. Genes differently expressed between N2 and UM10 animals at L4 stage and fed with the WT K12 diet, related to Figure 5. The raw RNA-seq data can be accessed through https://www.ncbi.nlm.nih.gov/geo/query/acc.cgi?acc=GSE169204 in GEO database.**
**Table S2. Genes differently expressed between UM10 animals at L4 stage and fed with WT and *csgA(-)* K12 diet, related to Figure 5. The raw RNA-seq data can be accessed through https://www.ncbi.nlm.nih.gov/geo/query/acc.cgi?acc=GSE169204 in GEO database.**
**Table S3. Oligonucleotides used in this study, related to Star Methods.**

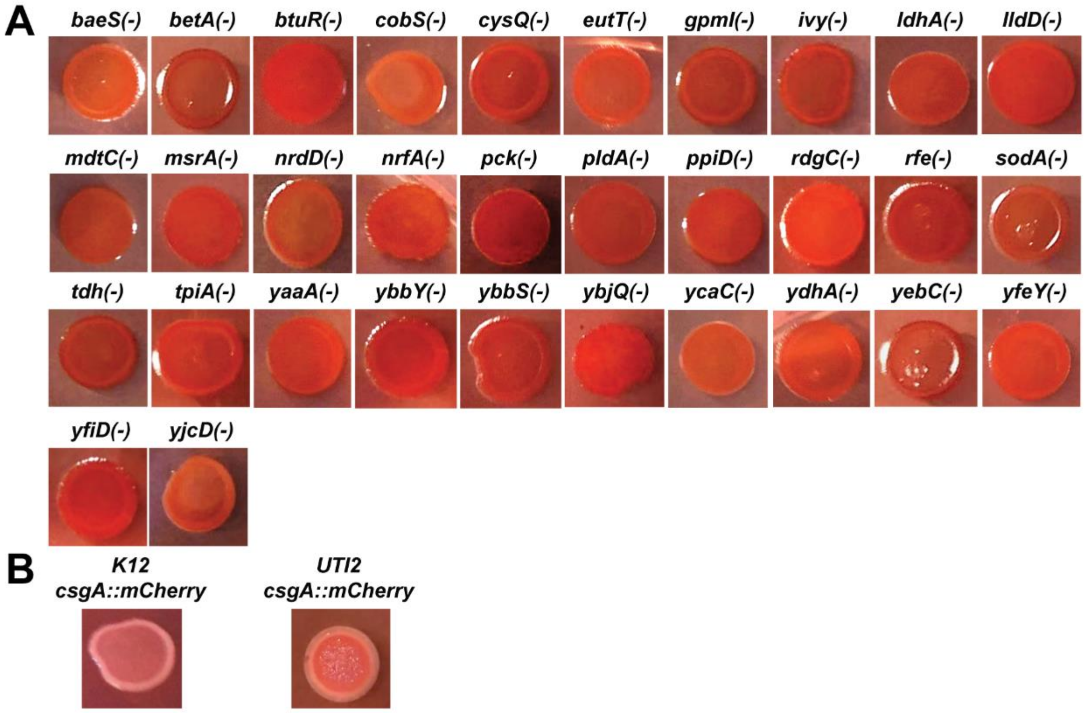


**Figure S1. Congo red staining of bacteria related to Figure 2.** (A) 32 mutant bacteria were grown in CR indicator plate at 25 degree for 2 days. (B) The congo red staining images of *K12-csgA::mCherry* and *UTI2-csgA:: mCherry* bacteria.

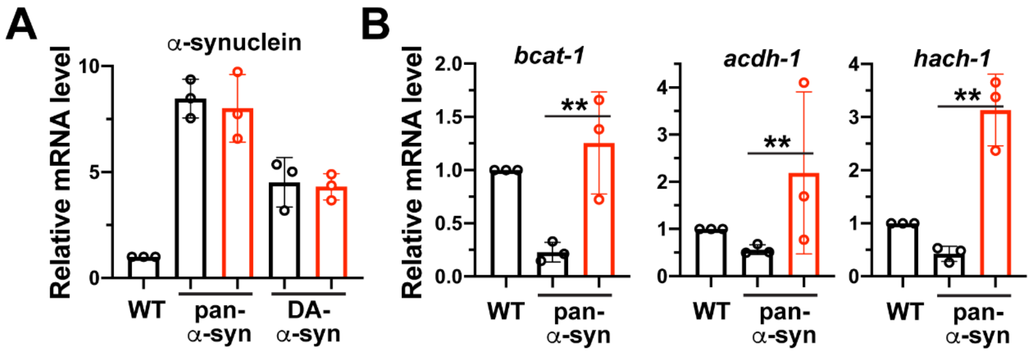


**Figure S2. RT-qPCR analysis of mitochondrial genes in PD animals, related to Figure 5.** (A) Transcription levels of *α-syn* in UM10 *unkIs7[aex-3p::α-syn(A53T), dat-1p::gfp]* and UM6 *unkIs9 [dat-1p::α -syn(A53T), dat-1p::gfp]* animals fed with WT or *csgA(-)* K12. Three independent reverse transcription and quantitative PCR (RT-qPCR) experiments were performed. (B) mRNA levels of *bcat-1*, *acdh-1*, and *hach-1* in UM10 animals fed with WT or *csgA(-)* K12. The data were presented as mean ± SD, followed with one way ANOVA *post-hoc* Tukey's analysis. Double asterisks indicate $p < 0.01$. The same applies to all supplemental figures.

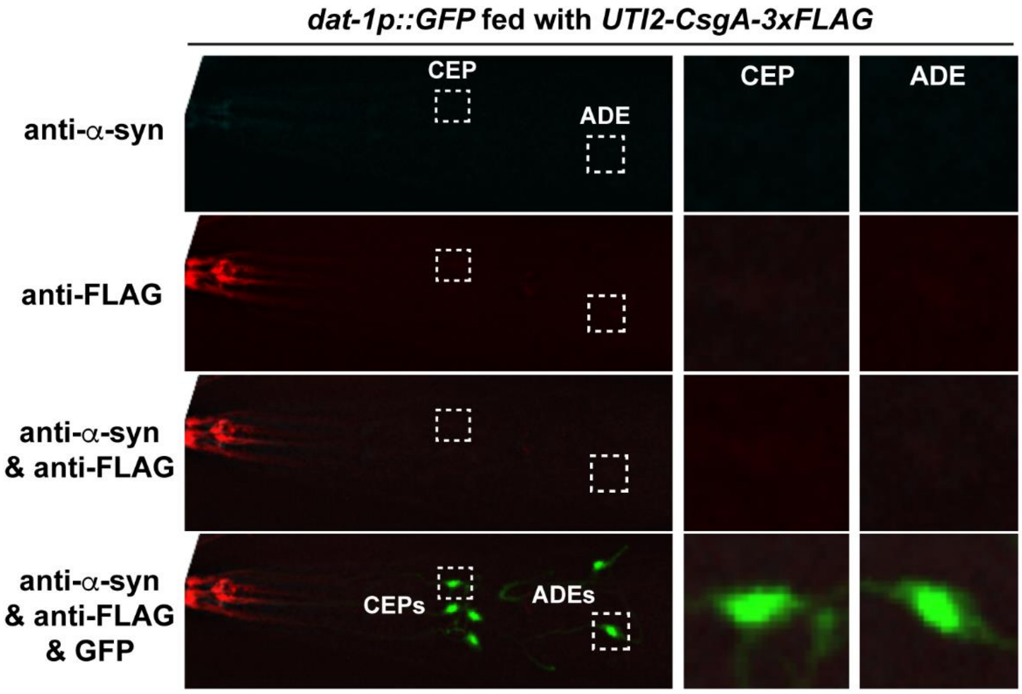


**Figure S3. CsgA retention by DA neurons depends on the expression of α-syn, related to Figure 4.** UM9 *unkIs11[dat-1p::GFP]* animals fed with *UTI2-csgA-3xFLAG* bacteria were stained with both anti-α-syn (cyan) and anti-FLAG (red) antibodies. GFP indicated the position of DA neurons. Inserts are enlarged images of the boxed region showing a representative CEP and ADE neuron, respectively. No anti-α-syn and anti-FLAG signals were detected in the DA neurons.

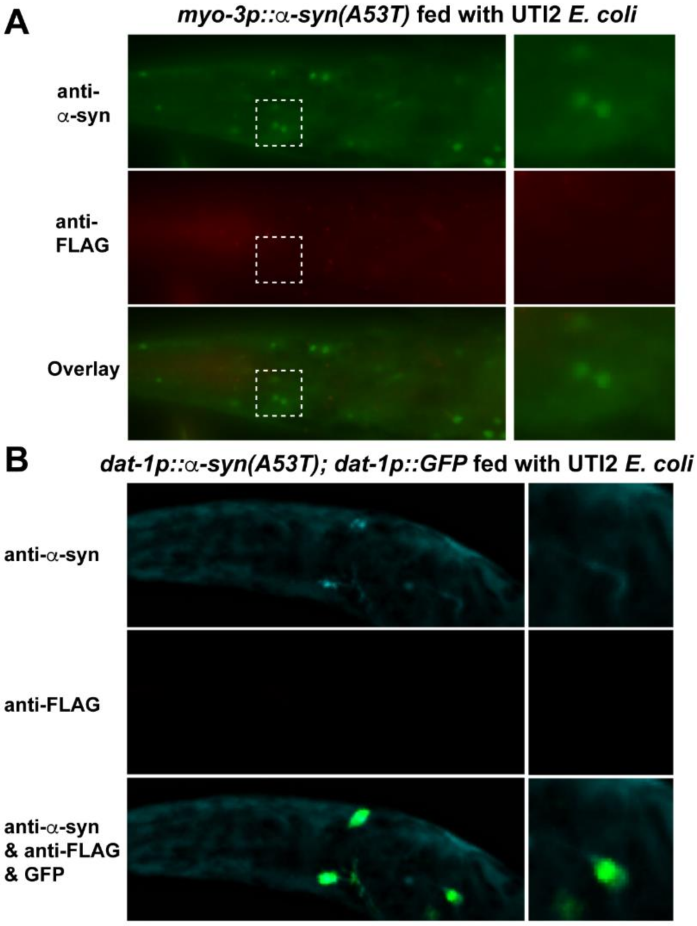


**Figure S4. Control experiments for the colocalization between CsgA and α-syn, related to Figure 4.** (A) CGZ512 *dpy-5(e907); unkEx109[myo-3p::α-syn(A53T); dpy-5(+)]* animals fed with UTI2 bacteria were stained with anti-α-syn (green) and anti-FLAG (red) antibodies. Insets are enlarged images of the boxed regions. No FLAG signals were detected. (B) UM6 *unkIs9 [dat-1p::α -syn(A53T), dat-1p::gfp]* animals fed with UTI2 bacteria were stained with anti-α-syn (cyan) and anti-FLAG (red) antibodies. Insets are enlarged images of the boxed regions. GFP labels the DA neurons, in which no FLAG signal was detected.

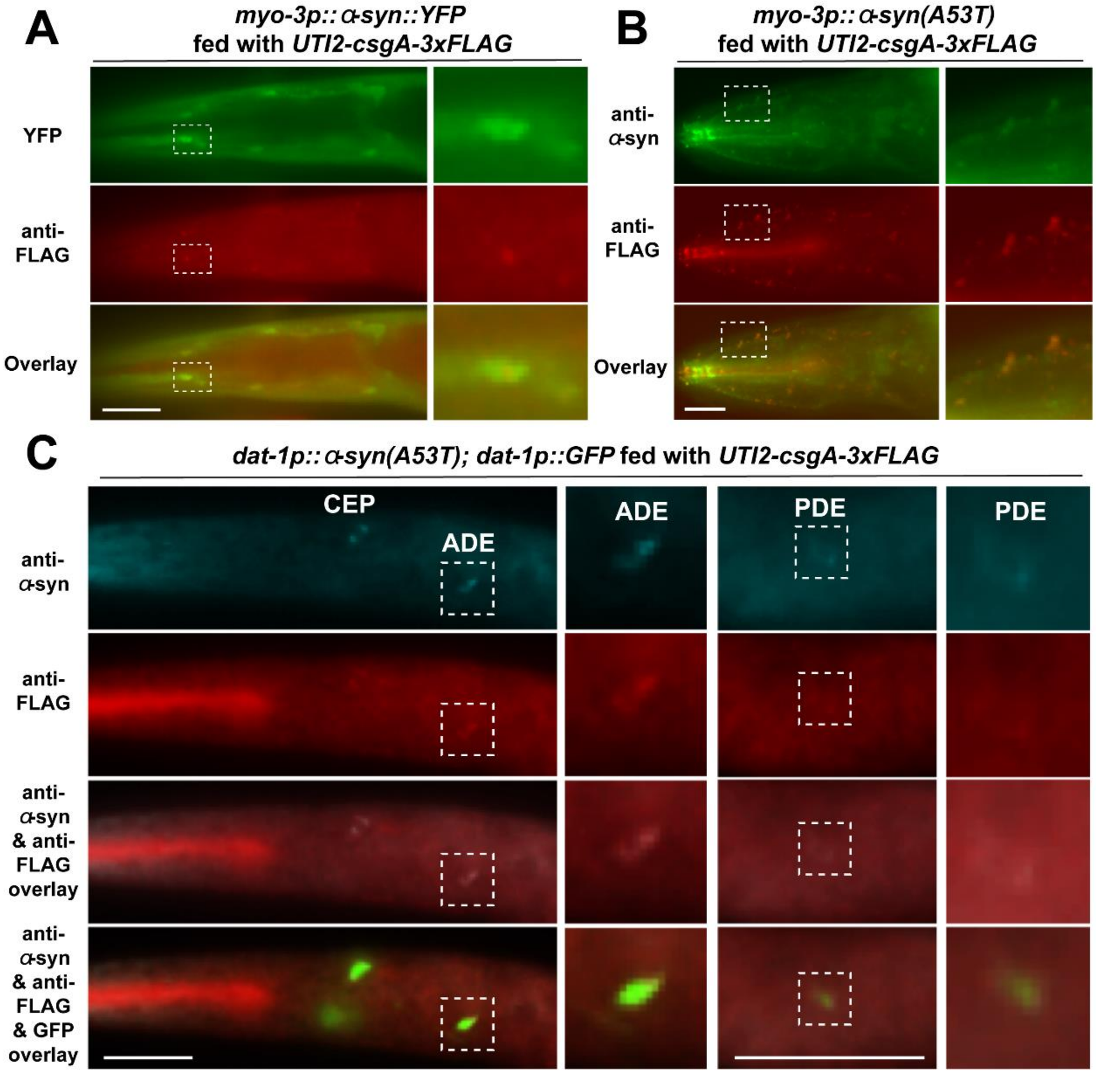


**Figure S5. Unprocessed images of CsgA colocalization with α-syn in *C. elegans* muscle and neurons, related to Figure 4.** (A) NL5901 *pkIs2386[unc-54p::α-syn::YFP; unc-119(+)]* animals fed with *UTI2-csgA-3xFLAG* bacteria were stained with anti-FLAG (red) antibodies. (B) CGZ512 *unkEx109[myo-3p::α-syn(A53T); dpy-5(+)]* animals fed with *UTI2-csgA-3xFLAG* bacteria were stained with anti-α-syn (green) and anti-FLAG (red) antibodies. (C) UM6 *unkIs9[dat-1p::α-syn(A53T), dat-1p::gfp]* animals fed with *UTI2-csgA-3xFLAG* bacteria were stained with both anti-α-syn (cyan) and anti-FLAG (red) antibodies. Scale bar = 20 μm. Insets are enlarged images of the boxed regions. Images were taken using a Leica DMi8 microscope without the THUNDER deconvolution processing. The deconvoluted images are shown in Figure 4.

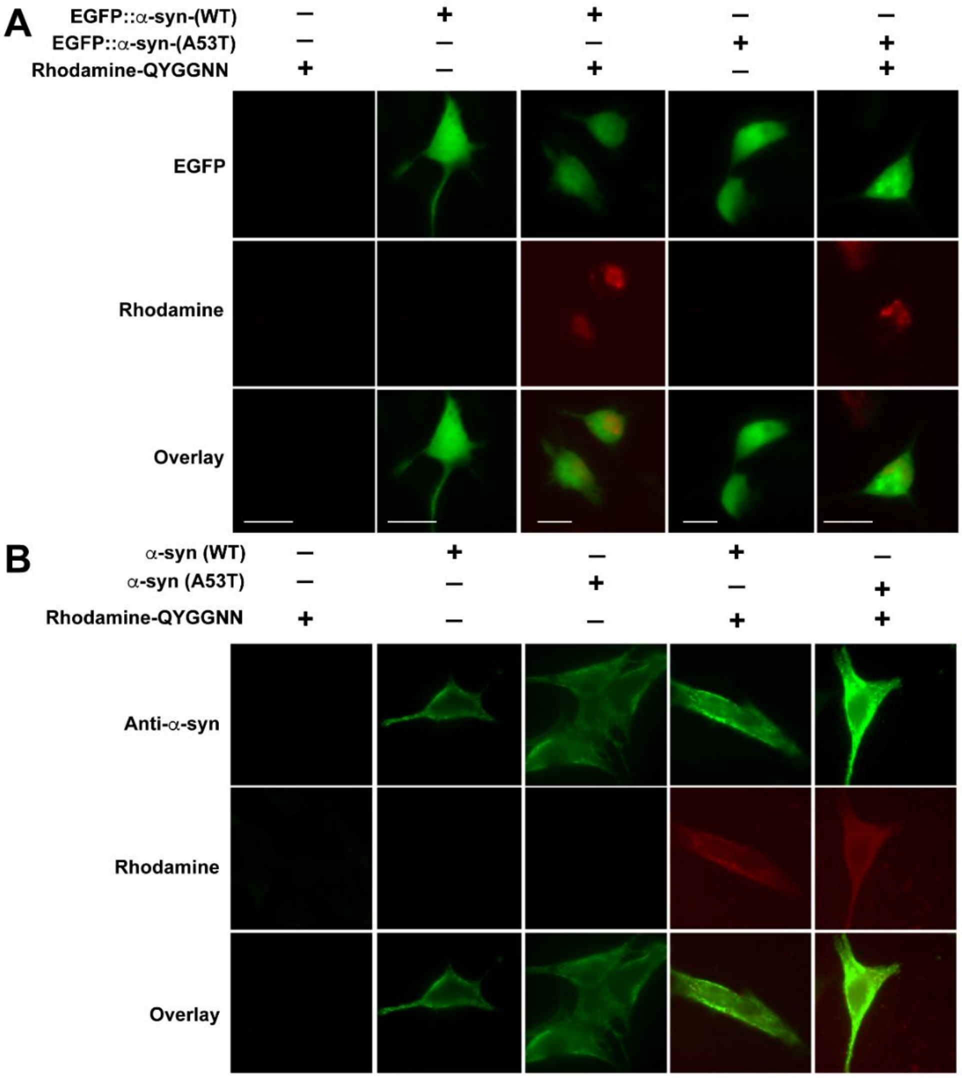


**Figure S6. CsgA-derived amyloidogenic peptides were retained in SH-SY5Y cells expressing α-syn, related to Figure 7.** (A) SH-SY5Y cells were transfected with constructs expressing EGFP::α-syn(WT or A53T) and then treated with Rhodamine-conjugated QYGGNN peptides. 24 hours after the transfection, the cells were imaged, and Rhodamine signal was only observed in cells expressing α-syn. (B) Unprocessed anti-α-syn (green) staining images of SH-SY5Y cells transfected with α-syn(WT or A53T)-expressing constructs and then treated with Rhodamine-conjugated QYGGNN peptides. The THUNDER-deconvoluted images is shown in Figure 7C.
