## supplementary table 1 for "Genome-wide screen identifies curli amyloid fibril as a bacterial component promoting host neurodegeneration"

| Gene | BaseMean of N2 fed with K12 WT | BaseMean of aex-3p::α-syn(A53T) fed with K12 WT | log2Fold Change | lfcSE | p value | adjusted p value |
|---|---|---|---|---|---|---|
| *W01A11.1* | 6.89 | 1515.48 | 6.94 | 0.54 | 3.14E-37 | 2.40E-33 |
| *T28D9.4* | 7.70 | 1565.65 | 6.85 | 0.54 | 7.20E-37 | 3.66E-33 |
| *mtl-1* | 10.70 | 1606.37 | 6.46 | 0.54 | 9.06E-33 | 3.46E-29 |
| *tts-1* | 276.95 | 33197.24 | 6.11 | 0.58 | 2.74E-26 | 5.97E-23 |
| *T23G5.11* | 0.22 | 120.91 | 5.60 | 0.79 | 1.35E-12 | 3.82E-10 |
| *K08D12.7* | 1.65 | 204.44 | 5.36 | 0.72 | 7.09E-14 | 2.58E-11 |
| *ilys-3* | 14.88 | 880.61 | 5.26 | 0.55 | 2.46E-21 | 3.41E-18 |
| *col-36* | 2.70 | 220.29 | 5.11 | 0.69 | 1.34E-13 | 4.46E-11 |
| *pals-14* | 2.91 | 197.82 | 5.03 | 0.65 | 1.15E-14 | 4.76E-12 |
| *col-40* | 6.23 | 384.80 | 4.88 | 0.67 | 4.62E-13 | 1.44E-10 |
| *C50F7.5* | 78.31 | 3540.93 | 4.85 | 0.59 | 1.27E-16 | 8.11E-14 |
| *fbxa-99* | 0.22 | 71.25 | 4.74 | 0.84 | 1.37E-08 | 1.36E-06 |
| *Y53F4B.8* | 1.81 | 121.59 | 4.63 | 0.72 | 1.33E-10 | 2.17E-08 |
| *D1086.2* | 2.70 | 143.34 | 4.63 | 0.69 | 2.08E-11 | 4.30E-09 |
| *T27A1.2* | 2.16 | 136.93 | 4.60 | 0.73 | 3.56E-10 | 5.23E-08 |
| *ugt-18* | 1.73 | 98.99 | 4.47 | 0.73 | 7.86E-10 | 1.08E-07 |
| *MTCE.7* | 343.45 | 9654.28 | 4.45 | 0.48 | 1.58E-20 | 1.86E-17 |
| *fbxa-163* | 2.26 | 126.20 | 4.40 | 0.74 | 3.24E-09 | 3.84E-07 |
| *W02D7.11* | 0.58 | 93.77 | 4.34 | 0.84 | 2.61E-07 | 1.70E-05 |
| *C54F6.5* | 4.78 | 157.77 | 4.26 | 0.64 | 2.71E-11 | 5.53E-09 |
| *M163.11* | 0.58 | 67.94 | 4.22 | 0.83 | 3.73E-07 | 2.28E-05 |
| *col-2* | 1.18 | 71.91 | 4.20 | 0.77 | 5.43E-08 | 4.46E-06 |
| *F56A4.3* | 0.58 | 52.94 | 4.11 | 0.82 | 5.24E-07 | 3.04E-05 |
| *rpr-1* | 1.18 | 64.55 | 4.09 | 0.78 | 1.31E-07 | 9.37E-06 |
| *hsp-12.6* | 6.36 | 163.38 | 4.03 | 0.62 | 6.25E-11 | 1.19E-08 |
| *Y39H10A.1* | 1.27 | 65.66 | 3.94 | 0.79 | 5.57E-07 | 3.20E-05 |
| *col-37* | 0.22 | 35.70 | 3.91 | 0.87 | 6.81E-06 | 0.0002453 |
| *Y37F4.15* | 0.22 | 33.76 | 3.86 | 0.87 | 9.69E-06 | 0.0003251 |
| *MTCE.33* | 237.60 | 3985.51 | 3.78 | 0.47 | 6.30E-16 | 3.44E-13 |
| *oac-12* | 1.72 | 90.41 | 3.78 | 0.83 | 4.96E-06 | 0.0001925 |
| *pals-6* | 6.36 | 157.38 | 3.76 | 0.70 | 7.01E-08 | 5.51E-06 |
| *col-51* | 0.22 | 35.33 | 3.74 | 0.88 | 2.13E-05 | 0.0005874 |
| *fbxa-48* | 0.58 | 43.88 | 3.72 | 0.85 | 1.24E-05 | 0.0003837 |
| *sri-11* | 1.18 | 48.24 | 3.69 | 0.80 | 3.57E-06 | 0.0001466 |
| *col-95* | 125.59 | 2203.27 | 3.69 | 0.56 | 6.36E-11 | 1.20E-08 |
| *sod-5* | 1.18 | 55.77 | 3.62 | 0.83 | 1.19E-05 | 0.0003766 |
| *egas-3* | 3.31 | 75.63 | 3.58 | 0.72 | 6.12E-07 | 3.50E-05 |
| *fbxa-30* | 1.83 | 54.84 | 3.57 | 0.77 | 3.65E-06 | 0.0001494 |
| *tbb-6* | 122.35 | 1749.57 | 3.51 | 0.51 | 3.99E-12 | 9.98E-10 |
| *hsp-16.49* | 4.86 | 134.87 | 3.51 | 0.79 | 7.76E-06 | 0.000271 |
| *F19H6.6* | 8.07 | 187.61 | 3.51 | 0.75 | 2.76E-06 | 0.0001198 |
| *col-102* | 1.09 | 42.09 | 3.49 | 0.83 | 2.28E-05 | 0.0006162 |
| *C45H4.14* | 0.58 | 32.96 | 3.48 | 0.85 | 4.57E-05 | 0.0010892 |
| *F02H6.3* | 6.88 | 119.23 | 3.47 | 0.65 | 1.11E-07 | 8.23E-06 |
| *Y110A2AL.2* | 0.22 | 26.41 | 3.46 | 0.89 | 9.77E-05 | 0.0019849 |
| *clec-60* | 99.39 | 1429.96 | 3.46 | 0.55 | 3.01E-10 | 4.61E-08 |
| *nnt-1* | 95.44 | 1306.19 | 3.45 | 0.51 | 1.57E-11 | 3.47E-09 |
| *F47B8.4* | 4.46 | 87.34 | 3.41 | 0.71 | 1.57E-06 | 7.51E-05 |
| *C29F7.10* | 2.39 | 57.04 | 3.40 | 0.77 | 9.73E-06 | 0.0003251 |
| *F36F12.2* | 1.08 | 41.70 | 3.38 | 0.84 | 5.81E-05 | 0.0013106 |
| *ech-9* | 4.72 | 116.55 | 3.38 | 0.80 | 2.19E-05 | 0.0005996 |

| | | | | | | |
|---|---|---|---|---|---|---|
| *fbxa-158* | 1.84 | 48.26 | 3.37 | 0.79 | 1.87E-05 | 0.0005302 |
| *oac-14* | 26.16 | 340.40 | 3.36 | 0.52 | 1.34E-10 | 2.17E-08 |
| *F59H6.5* | 0.22 | 23.52 | 3.34 | 0.89 | 0.00018 | 0.0032718 |
| *saeg-2* | 230.54 | 2701.74 | 3.30 | 0.47 | 1.55E-12 | 4.23E-10 |
| *far-7* | 29.84 | 362.85 | 3.28 | 0.52 | 3.43E-10 | 5.13E-08 |
| *srg-31* | 19.06 | 278.15 | 3.26 | 0.65 | 6.24E-07 | 3.53E-05 |
| *ilys-2* | 9.74 | 155.62 | 3.23 | 0.70 | 4.31E-06 | 0.0001716 |
| *grl-23* | 0.22 | 23.51 | 3.22 | 0.90 | 0.000348 | 0.0054795 |
| *T28A11.19* | 9.21 | 128.60 | 3.21 | 0.65 | 7.65E-07 | 4.17E-05 |
| *F49H6.5* | 1.29 | 38.26 | 3.21 | 0.82 | 9.76E-05 | 0.0019849 |
| *R07C12.1* | 4.11 | 71.11 | 3.19 | 0.75 | 2.12E-05 | 0.0005869 |
| *C54F6.15* | 0.22 | 21.12 | 3.17 | 0.90 | 0.000415 | 0.0062747 |
| *col-158* | 5.85 | 86.60 | 3.15 | 0.70 | 7.66E-06 | 0.0002698 |
| *nep-14* | 3.63 | 61.14 | 3.14 | 0.76 | 3.27E-05 | 0.0008347 |
| *Y58A7A.4* | 2.92 | 59.56 | 3.14 | 0.80 | 8.08E-05 | 0.0017093 |
| *F15B9.6* | 24.28 | 363.05 | 3.13 | 0.72 | 1.47E-05 | 0.0004421 |
| *fbxa-161* | 0.22 | 21.77 | 3.12 | 0.90 | 0.000535 | 0.0075106 |
| *C18D11.6* | 5.85 | 114.64 | 3.10 | 0.80 | 0.000106 | 0.0021272 |
| *cyp-37B1* | 130.01 | 1451.10 | 3.09 | 0.57 | 7.41E-08 | 5.77E-06 |
| *Y54G2A.57* | 0.62 | 29.42 | 3.09 | 0.87 | 0.000384 | 0.0058959 |
| *ZK829.9* | 66.43 | 695.59 | 3.07 | 0.53 | 6.93E-09 | 7.45E-07 |
| *R04D3.2* | 18.60 | 206.79 | 3.07 | 0.58 | 1.48E-07 | 1.04E-05 |
| *fkh-4* | 0.22 | 19.51 | 3.06 | 0.90 | 0.000711 | 0.0095346 |
| *grl-20* | 3.44 | 52.47 | 3.04 | 0.75 | 4.70E-05 | 0.0011136 |
| *C13A2.12* | 0.58 | 25.55 | 3.03 | 0.88 | 0.000592 | 0.0081822 |
| *F17A2.13* | 0.58 | 23.54 | 3.00 | 0.88 | 0.000623 | 0.0085305 |
| *sea-1* | 9.63 | 118.29 | 3.00 | 0.69 | 1.23E-05 | 0.0003818 |
| *F10C2.7* | 2.92 | 45.71 | 2.99 | 0.76 | 9.18E-05 | 0.0018978 |
| *F54F7.6* | 7.69 | 89.66 | 2.98 | 0.66 | 6.79E-06 | 0.0002452 |
| *flh-3* | 15.92 | 168.05 | 2.97 | 0.60 | 8.10E-07 | 4.35E-05 |
| *D1086.1* | 2.62 | 43.57 | 2.97 | 0.79 | 0.000185 | 0.0033485 |
| *fipr-24* | 4.02 | 66.38 | 2.95 | 0.79 | 0.000175 | 0.0032023 |
| *ceh-83* | 14.23 | 148.90 | 2.95 | 0.61 | 1.23E-06 | 6.20E-05 |
| *fbxa-170* | 8.79 | 96.52 | 2.94 | 0.64 | 4.54E-06 | 0.0001792 |
| *pals-32* | 16.63 | 170.30 | 2.94 | 0.60 | 8.02E-07 | 4.34E-05 |
| *C49F5.6* | 30.69 | 293.39 | 2.94 | 0.54 | 6.25E-08 | 5.02E-06 |
| *K09E3.7* | 14.69 | 154.72 | 2.93 | 0.63 | 2.78E-06 | 0.0001202 |
| *F23D12.2* | 9.13 | 104.39 | 2.93 | 0.68 | 1.45E-05 | 0.0004381 |
| *skr-14* | 2.25 | 37.54 | 2.91 | 0.80 | 0.000273 | 0.0045336 |
| *Y48A6B.8* | 11.21 | 133.65 | 2.88 | 0.72 | 5.87E-05 | 0.0013181 |
| *F26G5.1* | 26.31 | 257.21 | 2.87 | 0.61 | 2.41E-06 | 0.0001066 |
| *cdc-25.3* | 22.20 | 208.03 | 2.87 | 0.57 | 6.15E-07 | 3.50E-05 |
| *meg-4* | 36.03 | 326.34 | 2.85 | 0.56 | 3.78E-07 | 2.31E-05 |
| *arrd-11* | 2.25 | 35.69 | 2.83 | 0.81 | 0.000455 | 0.0066844 |
| *sip-1* | 2352.38 | 18997.70 | 2.82 | 0.45 | 4.02E-10 | 5.84E-08 |
| *fbxa-36* | 4.34 | 59.80 | 2.81 | 0.78 | 0.000314 | 0.0050562 |
| *C08E8.4* | 58.81 | 476.96 | 2.79 | 0.49 | 1.02E-08 | 1.06E-06 |
| *C17E7.4* | 40.29 | 336.70 | 2.77 | 0.54 | 2.39E-07 | 1.58E-05 |
| *pals-39* | 20.07 | 172.07 | 2.76 | 0.57 | 1.09E-06 | 5.54E-05 |
| *skr-7* | 6.06 | 66.15 | 2.76 | 0.72 | 0.000113 | 0.0022419 |
| *arrd-2* | 1.59 | 29.06 | 2.76 | 0.85 | 0.001099 | 0.0133679 |
| *clec-222* | 4.47 | 50.30 | 2.74 | 0.74 | 0.000207 | 0.0036401 |
| *F54D5.5* | 38.10 | 314.95 | 2.74 | 0.55 | 6.57E-07 | 3.69E-05 |
| *slc-36.5* | 6.36 | 68.51 | 2.73 | 0.73 | 0.000175 | 0.0031914 |
| *R09A8.2* | 13.50 | 120.36 | 2.72 | 0.63 | 1.41E-05 | 0.0004305 |
| *cup-16* | 61.09 | 475.53 | 2.72 | 0.50 | 6.49E-08 | 5.16E-06 |

| | | | | | | |
|---|---|---|---|---|---|---|
| *Y17D7C.3* | 43.25 | 359.89 | 2.72 | 0.57 | 2.28E-06 | 0.0001016 |
| *R09F10.8* | 108.94 | 832.15 | 2.71 | 0.48 | 2.12E-08 | 2.01E-06 |
| *Y110A2AL.3* | 2.10 | 31.88 | 2.71 | 0.83 | 0.001065 | 0.013042 |
| *W07B8.4* | 5.54 | 58.57 | 2.70 | 0.72 | 0.000188 | 0.003381 |
| *F36F12.1* | 5.15 | 61.17 | 2.70 | 0.77 | 0.000473 | 0.0068806 |
| *hsp-16.48* | 6.10 | 82.73 | 2.70 | 0.81 | 0.000824 | 0.0107387 |
| *Y53F4B.62* | 0.58 | 23.23 | 2.69 | 0.90 | 0.002821 | 0.0269446 |
| *sdz-28* | 1.19 | 23.91 | 2.69 | 0.86 | 0.001718 | 0.018716 |
| *col-135* | 98.13 | 729.65 | 2.68 | 0.48 | 1.74E-08 | 1.70E-06 |
| *F43D2.2* | 19.00 | 160.01 | 2.68 | 0.61 | 1.26E-05 | 0.0003915 |
| *fbxa-164* | 0.22 | 15.70 | 2.67 | 0.91 | 0.003505 | 0.031731 |
| *cpg-1* | 3154.72 | 23757.41 | 2.66 | 0.51 | 1.78E-07 | 1.21E-05 |
| *F15E6.10* | 0.22 | 16.00 | 2.66 | 0.91 | 0.00353 | 0.0318486 |
| *K03A11.1* | 2.56 | 35.76 | 2.66 | 0.81 | 0.001037 | 0.0127397 |
| *btb-6* | 3.62 | 43.21 | 2.66 | 0.78 | 0.000665 | 0.0090073 |
| *memi-2* | 98.19 | 738.71 | 2.65 | 0.52 | 3.04E-07 | 1.93E-05 |
| *F14D7.2* | 60.52 | 465.35 | 2.65 | 0.54 | 1.03E-06 | 5.31E-05 |
| *Y49F6C.8* | 11.51 | 95.48 | 2.63 | 0.62 | 2.42E-05 | 0.0006437 |
| *ZK177.1* | 27.87 | 211.39 | 2.63 | 0.55 | 1.39E-06 | 6.85E-05 |
| *tag-276* | 2.59 | 34.18 | 2.63 | 0.82 | 0.00129 | 0.0150917 |
| *col-44* | 1.32 | 26.17 | 2.63 | 0.86 | 0.002303 | 0.0234702 |
| *ZK813.2* | 256.73 | 1832.40 | 2.62 | 0.48 | 4.40E-08 | 3.80E-06 |
| *Y17D7C.2* | 4.13 | 43.82 | 2.62 | 0.75 | 0.000494 | 0.0071068 |
| *T01G5.7* | 5.96 | 58.45 | 2.62 | 0.73 | 0.000321 | 0.0051245 |
| *Y47H10A.3* | 1.07 | 21.86 | 2.62 | 0.87 | 0.002724 | 0.0263466 |
| *meg-3* | 63.60 | 480.10 | 2.61 | 0.56 | 3.36E-06 | 0.0001405 |
| *grl-25* | 4.52 | 48.90 | 2.60 | 0.76 | 0.000647 | 0.0087929 |
| *ZK1240.1* | 4.58 | 56.85 | 2.60 | 0.81 | 0.001262 | 0.0148214 |
| *T12D8.5* | 246.17 | 1847.57 | 2.60 | 0.56 | 4.22E-06 | 0.0001687 |
| *nas-10* | 2.42 | 43.76 | 2.59 | 0.87 | 0.002817 | 0.026937 |
| *R04D3.3* | 141.11 | 1022.07 | 2.59 | 0.53 | 1.06E-06 | 5.43E-05 |
| *F55G1.7* | 12.49 | 102.25 | 2.59 | 0.64 | 5.59E-05 | 0.0012814 |
| *meg-1* | 208.23 | 1508.82 | 2.59 | 0.54 | 1.37E-06 | 6.82E-05 |
| *let-99* | 292.18 | 2093.73 | 2.58 | 0.53 | 8.70E-07 | 4.63E-05 |
| *ztf-25* | 10.11 | 85.32 | 2.58 | 0.67 | 0.000111 | 0.0022111 |
| *gcy-17* | 1.58 | 25.34 | 2.57 | 0.86 | 0.002682 | 0.0260369 |
| *rgs-11* | 6.72 | 62.84 | 2.57 | 0.73 | 0.000426 | 0.0063619 |
| *R11A5.3* | 16.76 | 146.31 | 2.57 | 0.69 | 0.000214 | 0.0037315 |
| *pals-26* | 9.80 | 79.72 | 2.57 | 0.65 | 7.64E-05 | 0.0016258 |
| *fbxa-166* | 3.44 | 40.35 | 2.56 | 0.80 | 0.001416 | 0.0161136 |
| *C31G12.1* | 1.87 | 28.27 | 2.56 | 0.84 | 0.002401 | 0.0241602 |
| *ZC443.3* | 128.54 | 876.78 | 2.56 | 0.48 | 1.12E-07 | 8.23E-06 |
| *tag-52* | 70.78 | 504.82 | 2.56 | 0.54 | 2.11E-06 | 9.52E-05 |
| *F22F7.4* | 1.19 | 23.00 | 2.56 | 0.87 | 0.003305 | 0.030381 |
| *clec-17* | 29.29 | 222.94 | 2.56 | 0.60 | 2.09E-05 | 0.0005801 |
| *ZC308.4* | 77.80 | 540.47 | 2.56 | 0.51 | 4.81E-07 | 2.85E-05 |
| *arrd-24* | 8.22 | 70.92 | 2.56 | 0.69 | 0.000191 | 0.0034183 |
| *nhr-2* | 19.09 | 143.91 | 2.56 | 0.60 | 1.85E-05 | 0.0005244 |
| *cnc-11* | 2.92 | 34.21 | 2.55 | 0.80 | 0.001449 | 0.0164126 |
| *tsp-2* | 9.30 | 80.74 | 2.55 | 0.70 | 0.00027 | 0.0044904 |
| *M01B2.13* | 0.58 | 18.04 | 2.55 | 0.90 | 0.004688 | 0.0391511 |
| *R03H10.6* | 25.84 | 195.18 | 2.55 | 0.61 | 2.88E-05 | 0.0007502 |
| *memi-1* | 157.65 | 1063.32 | 2.54 | 0.49 | 2.07E-07 | 1.39E-05 |
| *col-50* | 2.57 | 33.10 | 2.54 | 0.82 | 0.002043 | 0.0214059 |
| *Y110A2AL.9* | 6.24 | 56.98 | 2.53 | 0.73 | 0.000555 | 0.0077578 |
| *ssp-37* | 15.42 | 117.51 | 2.53 | 0.62 | 4.94E-05 | 0.0011554 |

| | | | | | | |
|---|---|---|---|---|---|---|
| *F14H3.6* | 224.89 | 1482.23 | 2.53 | 0.47 | 5.71E-08 | 4.66E-06 |
| *clx-1* | 6.24 | 61.06 | 2.53 | 0.75 | 0.000785 | 0.0102989 |
| *rgs-9* | 22.45 | 161.02 | 2.52 | 0.58 | 1.55E-05 | 0.0004572 |
| *linc-84* | 1.07 | 20.44 | 2.51 | 0.88 | 0.004295 | 0.036693 |
| *F47B10.9* | 44.54 | 355.16 | 2.51 | 0.67 | 0.000173 | 0.0031734 |
| *ZK355.3* | 0.62 | 19.28 | 2.51 | 0.90 | 0.00506 | 0.0413048 |
| *hsp-12.3* | 65.36 | 483.70 | 2.51 | 0.61 | 4.36E-05 | 0.0010571 |
| *F11A5.15* | 0.22 | 13.85 | 2.51 | 0.91 | 0.006073 | 0.0470094 |
| *clec-87* | 1173.00 | 7655.96 | 2.51 | 0.48 | 1.41E-07 | 1.00E-05 |
| *fbxc-37* | 0.22 | 14.89 | 2.50 | 0.92 | 0.006218 | 0.047722 |
| *F58F12.5* | 1.19 | 21.34 | 2.50 | 0.87 | 0.004075 | 0.0353235 |
| *F08F3.6* | 39.15 | 266.81 | 2.50 | 0.54 | 3.46E-06 | 0.0001429 |
| *T24D1.3* | 255.55 | 1677.79 | 2.50 | 0.49 | 3.80E-07 | 2.31E-05 |
| *F19H6.4* | 25.89 | 181.60 | 2.50 | 0.57 | 1.17E-05 | 0.0003723 |
| *cpg-2* | 3358.66 | 22030.36 | 2.50 | 0.49 | 3.91E-07 | 2.37E-05 |
| *F14F9.2* | 0.22 | 15.89 | 2.50 | 0.92 | 0.006333 | 0.0483643 |
| *fbxa-21* | 15.71 | 125.17 | 2.48 | 0.68 | 0.000264 | 0.0044141 |
| *K10G4.3* | 61.18 | 452.18 | 2.48 | 0.63 | 9.31E-05 | 0.0019161 |
| *W05F2.3* | 1300.41 | 8154.29 | 2.47 | 0.46 | 1.07E-07 | 7.99E-06 |
| *tbx-33* | 8.62 | 69.81 | 2.46 | 0.70 | 0.000457 | 0.0067076 |
| *Y54G2A.40* | 20.68 | 169.58 | 2.46 | 0.71 | 0.000543 | 0.0076004 |
| *C01G6.3* | 70.71 | 464.91 | 2.46 | 0.53 | 3.76E-06 | 0.0001528 |
| *M151.7* | 41.11 | 286.39 | 2.46 | 0.60 | 3.93E-05 | 0.0009751 |
| *ZK355.8* | 9.29 | 70.39 | 2.46 | 0.66 | 0.0002 | 0.0035535 |
| *B0281.5* | 2.92 | 32.42 | 2.46 | 0.81 | 0.002518 | 0.0249605 |
| *T06D4.1* | 42.74 | 277.41 | 2.46 | 0.52 | 2.47E-06 | 0.0001088 |
| *C06G3.3* | 11.75 | 92.49 | 2.45 | 0.69 | 0.000385 | 0.0059075 |
| *C53B7.3* | 80.70 | 502.35 | 2.45 | 0.47 | 2.38E-07 | 1.58E-05 |
| *F14H3.3* | 9.59 | 75.02 | 2.44 | 0.69 | 0.000433 | 0.006445 |
| *his-25* | 5.86 | 48.86 | 2.44 | 0.72 | 0.000729 | 0.0097093 |
| *F02H6.4* | 23.67 | 158.82 | 2.44 | 0.57 | 2.16E-05 | 0.0005944 |
| *F02H6.2* | 20.79 | 139.13 | 2.43 | 0.58 | 2.47E-05 | 0.0006523 |
| *F14D2.8* | 18.43 | 135.72 | 2.43 | 0.66 | 0.000259 | 0.0043441 |
| *pals-33* | 3.62 | 35.69 | 2.42 | 0.79 | 0.002247 | 0.0230182 |
| *F52G3.6* | 26.89 | 199.58 | 2.42 | 0.68 | 0.00035 | 0.0054957 |
| *fbxb-7* | 2.24 | 26.33 | 2.41 | 0.84 | 0.003968 | 0.0347109 |
| *F43C11.7* | 83.66 | 533.46 | 2.40 | 0.55 | 1.15E-05 | 0.0003651 |
| *E02H4.6* | 21.54 | 139.83 | 2.40 | 0.57 | 2.28E-05 | 0.0006162 |
| *C09B8.5* | 1.57 | 22.74 | 2.40 | 0.87 | 0.005762 | 0.0453191 |
| *ent-5* | 29.64 | 191.10 | 2.40 | 0.57 | 2.27E-05 | 0.0006162 |
| *F43G6.10* | 33.48 | 215.63 | 2.40 | 0.57 | 2.35E-05 | 0.0006294 |
| *egg-4* | 215.73 | 1305.84 | 2.40 | 0.49 | 9.30E-07 | 4.91E-05 |
| *F55B12.10* | 13.93 | 97.44 | 2.40 | 0.64 | 0.000172 | 0.0031541 |
| *cpg-3* | 436.66 | 2630.43 | 2.39 | 0.49 | 9.71E-07 | 5.05E-05 |
| *ptr-22* | 4.65 | 40.78 | 2.39 | 0.76 | 0.001775 | 0.0192054 |
| *ZK813.7* | 577.88 | 3458.65 | 2.39 | 0.48 | 6.37E-07 | 3.59E-05 |
| *C04B4.2* | 83.87 | 511.32 | 2.39 | 0.51 | 2.99E-06 | 0.0001284 |
| *egg-5* | 390.26 | 2306.90 | 2.38 | 0.47 | 3.96E-07 | 2.39E-05 |
| *F02E8.4* | 107.31 | 652.13 | 2.38 | 0.51 | 3.18E-06 | 0.0001343 |
| *ttr-39* | 38.29 | 238.38 | 2.38 | 0.54 | 1.10E-05 | 0.0003558 |
| *M151.3* | 9.57 | 69.09 | 2.38 | 0.68 | 0.000433 | 0.0064494 |
| *R09D1.12* | 2.08 | 25.15 | 2.38 | 0.85 | 0.005299 | 0.0425749 |
| *fkh-3* | 5.35 | 43.85 | 2.38 | 0.74 | 0.001362 | 0.0156591 |
| *era-1* | 312.54 | 1884.63 | 2.37 | 0.51 | 3.42E-06 | 0.0001419 |
| *K04C1.5* | 50.85 | 312.35 | 2.37 | 0.54 | 1.01E-05 | 0.0003326 |
| *tos-1* | 2634.45 | 16211.89 | 2.37 | 0.54 | 1.33E-05 | 0.0004075 |

| | | | | | | |
|---|---|---|---|---|---|---|
| *fbxa-127* | 5.45 | 44.80 | 2.36 | 0.75 | 0.001759 | 0.0190761 |
| *abf-5* | 16.42 | 115.79 | 2.35 | 0.68 | 0.000537 | 0.007515 |
| *F53F8.3* | 24.34 | 153.75 | 2.35 | 0.58 | 5.84E-05 | 0.0013132 |
| *pos-1* | 1312.68 | 7638.18 | 2.35 | 0.48 | 1.20E-06 | 6.09E-05 |
| *str-183* | 2.40 | 25.82 | 2.35 | 0.83 | 0.004628 | 0.0387702 |
| *puf-5* | 1174.58 | 6870.41 | 2.35 | 0.49 | 1.86E-06 | 8.54E-05 |
| *B0416.4* | 5.45 | 44.37 | 2.35 | 0.75 | 0.001869 | 0.0199232 |
| *ttr-23* | 9.79 | 71.20 | 2.35 | 0.70 | 0.000751 | 0.0099278 |
| *T10H9.8* | 25.16 | 161.24 | 2.34 | 0.60 | 9.42E-05 | 0.0019311 |
| *memi-3* | 195.21 | 1130.50 | 2.34 | 0.49 | 1.56E-06 | 7.50E-05 |
| *inx-2* | 21.85 | 143.01 | 2.34 | 0.63 | 0.000204 | 0.0035892 |
| *ZC53.1* | 15.90 | 103.46 | 2.33 | 0.62 | 0.000188 | 0.0033781 |
| *EEED8.3* | 101.49 | 589.12 | 2.33 | 0.50 | 3.06E-06 | 0.00013 |
| *W06D11.3* | 23.10 | 145.94 | 2.33 | 0.60 | 0.000115 | 0.0022687 |
| *try-1* | 48.28 | 287.30 | 2.32 | 0.54 | 1.78E-05 | 0.0005098 |
| *T10C6.7* | 34.21 | 209.01 | 2.32 | 0.57 | 5.35E-05 | 0.0012363 |
| *F31F6.2* | 16.15 | 102.72 | 2.32 | 0.61 | 0.000159 | 0.0029429 |
| *C50E3.11* | 28.45 | 170.67 | 2.30 | 0.57 | 4.82E-05 | 0.0011358 |
| *rrf-2* | 61.26 | 365.35 | 2.30 | 0.57 | 4.70E-05 | 0.0011136 |
| *dct-1* | 172.47 | 1025.80 | 2.30 | 0.57 | 4.85E-05 | 0.0011416 |
| *meg-2* | 38.04 | 228.12 | 2.30 | 0.58 | 6.80E-05 | 0.0014805 |
| *skr-15* | 2.75 | 26.75 | 2.30 | 0.82 | 0.005385 | 0.0430428 |
| *mesp-1* | 449.84 | 2472.51 | 2.29 | 0.46 | 6.18E-07 | 3.51E-05 |
| *gln-6* | 833.31 | 4615.72 | 2.29 | 0.47 | 1.39E-06 | 6.85E-05 |
| *clec-88* | 444.39 | 2464.12 | 2.29 | 0.48 | 1.66E-06 | 7.94E-05 |
| *F01G4.4* | 606.55 | 3531.39 | 2.29 | 0.55 | 2.86E-05 | 0.0007477 |
| *spn-4* | 1862.05 | 10446.60 | 2.29 | 0.50 | 4.03E-06 | 0.0001618 |
| *C05C10.5* | 466.95 | 2653.24 | 2.29 | 0.52 | 9.11E-06 | 0.0003085 |
| *F40G12.11* | 32.77 | 193.59 | 2.28 | 0.57 | 6.70E-05 | 0.0014634 |
| *egg-2* | 226.10 | 1256.04 | 2.28 | 0.49 | 4.13E-06 | 0.0001653 |
| *lsy-27* | 27.86 | 162.40 | 2.28 | 0.56 | 4.48E-05 | 0.0010767 |
| *mex-6* | 833.17 | 4658.85 | 2.28 | 0.50 | 6.47E-06 | 0.0002386 |
| *C01G8.1* | 507.58 | 2816.91 | 2.28 | 0.50 | 4.37E-06 | 0.0001738 |
| *C27C12.3* | 42.67 | 252.58 | 2.27 | 0.58 | 9.21E-05 | 0.0019027 |
| *puf-7* | 452.42 | 2538.08 | 2.27 | 0.52 | 1.12E-05 | 0.0003616 |
| *R05G9.3* | 12.73 | 82.21 | 2.27 | 0.66 | 0.000588 | 0.008132 |
| *puf-6* | 319.83 | 1786.57 | 2.26 | 0.52 | 1.39E-05 | 0.000426 |
| *T12G3.6* | 136.86 | 738.53 | 2.26 | 0.47 | 1.86E-06 | 8.54E-05 |
| *gyg-2* | 526.63 | 2858.28 | 2.25 | 0.49 | 3.34E-06 | 0.0001402 |
| *W02D9.6* | 88.56 | 487.82 | 2.25 | 0.51 | 1.24E-05 | 0.0003837 |
| *F58G11.3* | 352.01 | 1944.22 | 2.25 | 0.52 | 1.54E-05 | 0.0004572 |
| *C50E3.13* | 19.02 | 115.14 | 2.25 | 0.62 | 0.000319 | 0.0051035 |
| *trcs-1* | 492.97 | 2650.52 | 2.25 | 0.48 | 3.05E-06 | 0.00013 |
| *F53B2.8* | 369.92 | 2093.56 | 2.24 | 0.55 | 5.10E-05 | 0.0011877 |
| *Y75D11A.3* | 21.04 | 128.33 | 2.24 | 0.64 | 0.000417 | 0.0062881 |
| *oma-2* | 1050.31 | 5670.70 | 2.24 | 0.50 | 6.87E-06 | 0.0002469 |
| *cbd-1* | 4201.90 | 22920.64 | 2.23 | 0.52 | 1.55E-05 | 0.0004572 |
| *C49F5.3* | 26.56 | 152.38 | 2.23 | 0.58 | 0.000129 | 0.0024869 |
| *pzf-1* | 15.42 | 93.16 | 2.23 | 0.63 | 0.000423 | 0.0063362 |
| *egg-1* | 906.51 | 4829.56 | 2.23 | 0.49 | 4.79E-06 | 0.0001868 |
| *chs-1* | 463.81 | 2554.84 | 2.23 | 0.54 | 3.65E-05 | 0.0009175 |
| *F55G11.4* | 89.26 | 492.35 | 2.22 | 0.54 | 3.89E-05 | 0.000966 |
| *F02D10.6* | 6.64 | 47.17 | 2.22 | 0.74 | 0.002816 | 0.026937 |
| *nos-2* | 225.58 | 1184.65 | 2.22 | 0.47 | 2.57E-06 | 0.0001124 |
| *ZC239.22* | 34.18 | 194.06 | 2.22 | 0.58 | 0.000128 | 0.0024789 |
| *T10C6.10* | 29.41 | 164.63 | 2.22 | 0.56 | 8.29E-05 | 0.0017463 |

| | | | | | | |
|---|---|---|---|---|---|---|
| *mex-1* | 1028.31 | 5423.51 | 2.21 | 0.49 | 7.36E-06 | 0.0002608 |
| *Y54G2A.36* | 38.96 | 223.38 | 2.21 | 0.60 | 0.000241 | 0.0041099 |
| *mei-2* | 338.84 | 1768.87 | 2.21 | 0.48 | 3.71E-06 | 0.0001509 |
| *W02D7.5* | 14.66 | 92.06 | 2.20 | 0.68 | 0.001308 | 0.0152508 |
| *cex-1* | 377.48 | 2200.72 | 2.19 | 0.63 | 0.000506 | 0.0072026 |
| *F32D1.7* | 194.18 | 1019.37 | 2.19 | 0.51 | 1.64E-05 | 0.0004777 |
| *tbx-9* | 35.91 | 192.52 | 2.18 | 0.55 | 6.65E-05 | 0.0014544 |
| *flh-1* | 215.94 | 1138.18 | 2.17 | 0.53 | 4.91E-05 | 0.0011508 |
| *W01F3.2* | 397.96 | 2042.15 | 2.17 | 0.49 | 1.13E-05 | 0.0003641 |
| *spsb-2* | 141.37 | 726.21 | 2.17 | 0.50 | 1.31E-05 | 0.0004025 |
| *D1086.7* | 251.56 | 1254.43 | 2.17 | 0.45 | 1.43E-06 | 6.98E-05 |
| *Y39B6A.10* | 25.45 | 142.22 | 2.17 | 0.61 | 0.000371 | 0.0057493 |
| *lea-1* | 3197.69 | 16067.50 | 2.17 | 0.47 | 3.39E-06 | 0.0001409 |
| *cdr-4* | 653.74 | 3641.85 | 2.17 | 0.61 | 0.000361 | 0.0056345 |
| *F43G6.7* | 50.47 | 263.77 | 2.17 | 0.53 | 4.42E-05 | 0.0010653 |
| *C44H9.5* | 120.96 | 688.54 | 2.15 | 0.64 | 0.000808 | 0.010576 |
| *perm-1* | 364.45 | 1836.19 | 2.15 | 0.49 | 1.42E-05 | 0.0004331 |
| *grl-9* | 5.36 | 37.17 | 2.14 | 0.76 | 0.0049 | 0.0404123 |
| *rme-2* | 805.50 | 4063.95 | 2.14 | 0.51 | 2.30E-05 | 0.0006188 |
| *daf-18* | 763.19 | 3871.41 | 2.14 | 0.52 | 3.28E-05 | 0.0008347 |
| *Y53F4B.45* | 631.90 | 3198.47 | 2.14 | 0.52 | 3.45E-05 | 0.0008707 |
| *hsp-43* | 879.47 | 4482.85 | 2.13 | 0.53 | 5.17E-05 | 0.0012009 |
| *H29C22.1* | 30.95 | 169.86 | 2.13 | 0.62 | 0.000526 | 0.0074075 |
| *ttr-50* | 14.19 | 80.41 | 2.13 | 0.65 | 0.000955 | 0.0119451 |
| *puf-11* | 858.85 | 4202.33 | 2.13 | 0.47 | 5.81E-06 | 0.000219 |
| *fbxa-69* | 16.87 | 93.56 | 2.12 | 0.64 | 0.000884 | 0.0113153 |
| *fbxa-192* | 42.64 | 221.09 | 2.12 | 0.57 | 0.000203 | 0.0035797 |
| *T23F2.4* | 8.58 | 51.73 | 2.11 | 0.70 | 0.002627 | 0.025643 |
| *tbx-36* | 30.46 | 159.38 | 2.11 | 0.58 | 0.000278 | 0.004583 |
| *F14F9.8* | 41.19 | 207.87 | 2.11 | 0.54 | 8.10E-05 | 0.0017097 |
| *cyb-2.1* | 325.71 | 1574.82 | 2.11 | 0.47 | 8.87E-06 | 0.0003029 |
| *gln-5* | 436.58 | 2118.92 | 2.11 | 0.48 | 1.21E-05 | 0.0003784 |
| *C17E7.9* | 13.95 | 76.69 | 2.10 | 0.64 | 0.000993 | 0.0123199 |
| *T05F1.2* | 241.76 | 1192.70 | 2.10 | 0.51 | 4.33E-05 | 0.0010529 |
| *F12E12.1* | 39.41 | 200.85 | 2.10 | 0.56 | 0.000177 | 0.0032247 |
| *puf-3* | 1084.99 | 5180.15 | 2.10 | 0.47 | 8.22E-06 | 0.0002846 |
| *pie-1* | 387.61 | 1858.48 | 2.09 | 0.48 | 1.41E-05 | 0.000431 |
| *C25F9.11* | 197.81 | 1048.40 | 2.09 | 0.62 | 0.000691 | 0.0092951 |
| *gfat-2* | 763.13 | 3710.24 | 2.09 | 0.51 | 3.90E-05 | 0.0009688 |
| *aakg-4* | 83.92 | 430.47 | 2.09 | 0.58 | 0.000339 | 0.005369 |
| *44258* | 242.32 | 1162.15 | 2.09 | 0.49 | 1.95E-05 | 0.0005467 |
| *cyb-2.2* | 506.70 | 2420.27 | 2.09 | 0.48 | 1.51E-05 | 0.0004506 |
| *ZK1053.4* | 7.40 | 46.03 | 2.08 | 0.74 | 0.004909 | 0.0404645 |
| *cyp-31A5* | 88.31 | 432.09 | 2.08 | 0.53 | 7.34E-05 | 0.0015687 |
| *fbxa-215* | 311.94 | 1479.79 | 2.08 | 0.48 | 1.44E-05 | 0.0004381 |
| *rnh-1.3* | 76.69 | 390.93 | 2.08 | 0.58 | 0.000357 | 0.0055815 |
| *fbxc-32* | 21.29 | 109.27 | 2.08 | 0.59 | 0.000423 | 0.0063362 |
| *Y48G1A.2* | 107.38 | 513.86 | 2.08 | 0.50 | 2.79E-05 | 0.0007318 |
| *F31B9.3* | 41.22 | 203.12 | 2.08 | 0.54 | 0.000127 | 0.0024571 |
| *F41G3.10* | 60.50 | 305.59 | 2.08 | 0.58 | 0.00032 | 0.0051245 |
| *lys-3* | 8.83 | 53.20 | 2.07 | 0.73 | 0.004339 | 0.0369781 |
| *B0462.5* | 24.46 | 139.18 | 2.07 | 0.69 | 0.002576 | 0.025318 |
| *mom-2* | 194.78 | 944.79 | 2.07 | 0.52 | 7.54E-05 | 0.0016076 |
| *W06D11.1* | 21.51 | 109.62 | 2.07 | 0.59 | 0.000424 | 0.0063506 |
| *inx-3* | 74.50 | 382.97 | 2.07 | 0.60 | 0.000589 | 0.00814 |
| *Y46H3C.7* | 19.85 | 107.38 | 2.06 | 0.66 | 0.001786 | 0.0192672 |

| | | | | | | |
|---|---|---|---|---|---|---|
| *C10G8.4* | 54.39 | 265.47 | 2.06 | 0.55 | 0.000186 | 0.0033632 |
| *R09A8.1* | 15.17 | 79.61 | 2.05 | 0.64 | 0.001243 | 0.0146488 |
| *Y73B6BL.44* | 28.33 | 138.79 | 2.04 | 0.57 | 0.00031 | 0.0050018 |
| *F14H3.4* | 23.62 | 119.73 | 2.04 | 0.61 | 0.00083 | 0.0108069 |
| *C06H2.7* | 34.08 | 165.37 | 2.04 | 0.56 | 0.000254 | 0.0042831 |
| *cpar-1* | 177.85 | 817.88 | 2.04 | 0.48 | 2.17E-05 | 0.000596 |
| *anr-32* | 9.54 | 56.84 | 2.04 | 0.74 | 0.00579 | 0.0454382 |
| *ZK813.1* | 446.35 | 2020.94 | 2.03 | 0.46 | 8.82E-06 | 0.0003019 |
| *T21C9.13* | 127.16 | 583.90 | 2.03 | 0.49 | 4.04E-05 | 0.0009987 |
| *nasp-2* | 1034.57 | 4709.00 | 2.02 | 0.48 | 2.41E-05 | 0.0006413 |
| *dod-22* | 41.29 | 193.92 | 2.02 | 0.53 | 0.000132 | 0.0025276 |
| *mex-5* | 2545.47 | 11933.06 | 2.02 | 0.53 | 0.000131 | 0.002514 |
| *fbxa-24* | 34.05 | 162.53 | 2.02 | 0.56 | 0.00027 | 0.0044904 |
| *B0304.4* | 43.59 | 203.78 | 2.02 | 0.53 | 0.000131 | 0.0025013 |
| *F31F6.1* | 8.61 | 48.48 | 2.02 | 0.72 | 0.004836 | 0.0400375 |
| *clec-91* | 307.28 | 1389.02 | 2.02 | 0.48 | 2.37E-05 | 0.0006357 |
| *szy-4* | 350.46 | 1621.23 | 2.01 | 0.52 | 0.000103 | 0.0020672 |
| *M60.4* | 1693.70 | 7749.92 | 2.01 | 0.50 | 6.27E-05 | 0.0013844 |
| *ceh-49* | 26.05 | 126.62 | 2.01 | 0.59 | 0.000639 | 0.0087132 |
| *clec-70* | 24.53 | 118.20 | 2.01 | 0.58 | 0.000565 | 0.00786 |
| *T04D3.5* | 33.36 | 164.79 | 2.01 | 0.61 | 0.001098 | 0.0133635 |
| *F18A1.7* | 658.29 | 2922.38 | 2.01 | 0.46 | 1.20E-05 | 0.0003777 |
| *ceh-39* | 57.26 | 271.16 | 2.01 | 0.56 | 0.000373 | 0.005757 |
| *T02G6.5* | 9.04 | 51.52 | 2.01 | 0.73 | 0.005714 | 0.0450341 |
| *B0507.6* | 10.52 | 55.72 | 2.00 | 0.68 | 0.00311 | 0.0289974 |
| *ZC239.14* | 14.93 | 76.29 | 2.00 | 0.65 | 0.00202 | 0.0212069 |
| *W06B4.1* | 360.50 | 1603.78 | 2.00 | 0.47 | 1.99E-05 | 0.0005566 |
| *srw-86* | 80.80 | 425.40 | 2.00 | 0.68 | 0.003302 | 0.0303789 |
| *F11A5.9* | 87.41 | 406.16 | 1.99 | 0.55 | 0.000306 | 0.0049517 |
| *T01C3.3* | 204.74 | 901.84 | 1.99 | 0.47 | 1.89E-05 | 0.0005343 |
| *mes-1* | 40.30 | 195.84 | 1.99 | 0.62 | 0.001253 | 0.0147457 |
| *F57A10.4* | 16.63 | 83.26 | 1.98 | 0.64 | 0.002095 | 0.0217703 |
| *cav-1* | 8.31 | 46.73 | 1.98 | 0.73 | 0.006643 | 0.0499542 |
| *srz-99* | 7.81 | 43.67 | 1.98 | 0.73 | 0.006524 | 0.049359 |
| *sodh-1* | 895.73 | 4338.18 | 1.98 | 0.62 | 0.001307 | 0.0152484 |
| *T22D1.5* | 317.66 | 1379.18 | 1.97 | 0.46 | 1.96E-05 | 0.0005476 |
| *egg-3* | 222.76 | 982.83 | 1.97 | 0.49 | 6.31E-05 | 0.001389 |
| *thn-1* | 12.53 | 66.55 | 1.97 | 0.70 | 0.004694 | 0.0391554 |
| *Y4C6A.3* | 95.39 | 427.34 | 1.96 | 0.53 | 0.000204 | 0.0035926 |
| *C27F2.7* | 41.25 | 192.61 | 1.96 | 0.59 | 0.000927 | 0.011681 |
| *ubc-25* | 1826.88 | 7730.03 | 1.95 | 0.44 | 8.01E-06 | 0.000278 |
| *lys-7* | 345.57 | 1550.01 | 1.95 | 0.54 | 0.000337 | 0.0053415 |
| *sod-3* | 32.57 | 148.87 | 1.95 | 0.57 | 0.000683 | 0.0092116 |
| *gna-2* | 204.59 | 882.15 | 1.95 | 0.49 | 6.28E-05 | 0.0013844 |
| *R02F2.4* | 122.63 | 532.83 | 1.94 | 0.50 | 0.000114 | 0.0022616 |
| *ule-2* | 287.73 | 1223.86 | 1.94 | 0.47 | 3.71E-05 | 0.0009299 |
| *T13F2.6* | 399.53 | 1732.35 | 1.94 | 0.51 | 0.000142 | 0.0026729 |
| *W09G12.7* | 21.81 | 102.09 | 1.94 | 0.61 | 0.001569 | 0.017528 |
| *T16G12.8* | 15.41 | 74.68 | 1.94 | 0.65 | 0.002894 | 0.0275044 |
| *pcm-1* | 44.18 | 213.69 | 1.94 | 0.65 | 0.002849 | 0.0271847 |
| *T24E12.1* | 9.53 | 49.70 | 1.93 | 0.71 | 0.006567 | 0.0495284 |
| *bir-2* | 216.09 | 926.93 | 1.93 | 0.50 | 0.000117 | 0.0023035 |
| *oma-1* | 720.27 | 3134.34 | 1.93 | 0.53 | 0.000265 | 0.0044141 |
| *F22E5.17* | 30.58 | 136.50 | 1.93 | 0.56 | 0.000642 | 0.0087421 |
| *ceh-91* | 163.90 | 701.44 | 1.92 | 0.51 | 0.000193 | 0.0034378 |
| *F14H3.5* | 11.27 | 55.29 | 1.91 | 0.68 | 0.004674 | 0.0390741 |

| | | | | | | |
|---|---|---|---|---|---|---|
| *wdr-5.3* | 383.49 | 1619.99 | 1.91 | 0.49 | 0.000106 | 0.0021218 |
| *Y51H7C.3* | 45.09 | 197.10 | 1.91 | 0.55 | 0.000493 | 0.0070949 |
| *rmd-1* | 1064.15 | 4405.83 | 1.91 | 0.46 | 3.50E-05 | 0.0008829 |
| *gst-24* | 84.90 | 419.61 | 1.91 | 0.69 | 0.005648 | 0.0446024 |
| *F13C5.1* | 38.94 | 194.98 | 1.91 | 0.70 | 0.006544 | 0.0494035 |
| *F10E9.12* | 51.36 | 242.31 | 1.90 | 0.65 | 0.003515 | 0.0317838 |
| *C25H3.10* | 37.67 | 162.81 | 1.90 | 0.55 | 0.000512 | 0.0072698 |
| *mex-3* | 1669.34 | 6940.34 | 1.90 | 0.49 | 9.49E-05 | 0.0019407 |
| *DY3.8* | 133.96 | 556.48 | 1.90 | 0.49 | 0.000101 | 0.002049 |
| *kgb-2* | 57.43 | 250.32 | 1.89 | 0.57 | 0.000833 | 0.010824 |
| *Y47H10A.5* | 74.63 | 335.40 | 1.89 | 0.61 | 0.001896 | 0.0201306 |
| *allo-1* | 970.08 | 3970.46 | 1.89 | 0.47 | 5.39E-05 | 0.0012451 |
| *T25B9.8* | 52.09 | 221.04 | 1.89 | 0.53 | 0.0004 | 0.0060877 |
| *aptf-3* | 170.10 | 711.87 | 1.89 | 0.51 | 0.00022 | 0.0038267 |
| *K08H2.3* | 13.95 | 65.22 | 1.89 | 0.65 | 0.003785 | 0.0335101 |
| *cyp-31A3* | 149.90 | 622.80 | 1.88 | 0.50 | 0.00019 | 0.0034055 |
| *puf-10* | 241.54 | 1022.04 | 1.88 | 0.54 | 0.00045 | 0.0066262 |
| *inx-22* | 221.63 | 923.24 | 1.88 | 0.51 | 0.000247 | 0.0042031 |
| *gld-3* | 758.95 | 3220.13 | 1.88 | 0.55 | 0.000628 | 0.0085746 |
| *ekl-7* | 235.28 | 975.45 | 1.88 | 0.51 | 0.000259 | 0.0043378 |
| *fbxa-162* | 13.72 | 64.37 | 1.87 | 0.66 | 0.00474 | 0.0394545 |
| *B0222.5* | 458.37 | 1911.19 | 1.87 | 0.53 | 0.000371 | 0.0057421 |
| *C50E3.12* | 14.68 | 67.27 | 1.87 | 0.65 | 0.003991 | 0.0347906 |
| *F23F1.6* | 62.23 | 256.50 | 1.87 | 0.51 | 0.000276 | 0.0045638 |
| *W02D9.7* | 149.28 | 612.54 | 1.87 | 0.51 | 0.000238 | 0.0040614 |
| *C16C8.12* | 19.47 | 87.70 | 1.86 | 0.64 | 0.003572 | 0.0321146 |
| *C14B1.9* | 333.44 | 1337.44 | 1.86 | 0.48 | 9.38E-05 | 0.0019245 |
| *D1086.6* | 599.17 | 2365.90 | 1.86 | 0.44 | 2.88E-05 | 0.0007502 |
| *clec-82* | 170.23 | 711.67 | 1.86 | 0.55 | 0.000725 | 0.0096739 |
| *maph-1.2* | 942.98 | 3805.84 | 1.86 | 0.49 | 0.000162 | 0.0029914 |
| *T04D3.1* | 15.70 | 72.35 | 1.85 | 0.67 | 0.005441 | 0.0433926 |
| *cyp-34A9* | 46.88 | 206.26 | 1.85 | 0.62 | 0.002937 | 0.0277965 |
| *skpo-1* | 363.24 | 1444.83 | 1.84 | 0.48 | 0.00013 | 0.0024947 |
| *clec-146* | 52.32 | 215.02 | 1.84 | 0.55 | 0.000782 | 0.010279 |
| *C06A5.8* | 79.33 | 315.22 | 1.83 | 0.49 | 0.000207 | 0.0036386 |
| *ced-3* | 741.01 | 2964.59 | 1.83 | 0.51 | 0.000357 | 0.0055827 |
| *B0393.6* | 482.91 | 1922.67 | 1.82 | 0.51 | 0.000354 | 0.0055374 |
| *Y71F9AL.8* | 141.95 | 557.87 | 1.82 | 0.49 | 0.000182 | 0.00329 |
| *ZK637.6* | 91.69 | 368.26 | 1.82 | 0.53 | 0.000649 | 0.0088031 |
| *ZK813.3* | 363.96 | 1418.23 | 1.81 | 0.49 | 0.000204 | 0.0035892 |
| *bath-19* | 27.14 | 118.20 | 1.81 | 0.65 | 0.005175 | 0.0419316 |
| *suco-1* | 1423.94 | 5838.42 | 1.81 | 0.58 | 0.00169 | 0.0184437 |
| *M60.7* | 39.01 | 160.24 | 1.80 | 0.58 | 0.002053 | 0.0214669 |
| *Y51F10.2* | 595.43 | 2297.80 | 1.80 | 0.48 | 0.000191 | 0.0034177 |
| *tps-1* | 510.35 | 1950.85 | 1.80 | 0.46 | 0.000101 | 0.002049 |
| *Y11D7A.7* | 24.37 | 101.30 | 1.80 | 0.60 | 0.002771 | 0.0266401 |
| *clec-223* | 23.55 | 99.10 | 1.80 | 0.62 | 0.00365 | 0.0327327 |
| *cht-3* | 217.32 | 841.30 | 1.79 | 0.50 | 0.000349 | 0.0054872 |
| *gst-15* | 31.14 | 127.93 | 1.79 | 0.59 | 0.002594 | 0.0254188 |
| *F33E11.2* | 277.65 | 1057.22 | 1.79 | 0.47 | 0.000146 | 0.0027327 |
| *dut-1* | 479.36 | 1821.12 | 1.79 | 0.47 | 0.000127 | 0.0024571 |
| *C08F11.13* | 448.32 | 1869.48 | 1.79 | 0.62 | 0.003669 | 0.0328332 |
| *csa-1* | 19.70 | 85.66 | 1.79 | 0.66 | 0.006634 | 0.0499356 |
| *M01H9.3* | 3276.37 | 12748.39 | 1.79 | 0.52 | 0.000594 | 0.0081926 |
| *B0416.7* | 22.22 | 93.71 | 1.79 | 0.63 | 0.004584 | 0.0385091 |
| *C31H1.8* | 380.23 | 1454.52 | 1.78 | 0.49 | 0.000289 | 0.0047254 |

| | | | | | | |
|---|---|---|---|---|---|---|
| *F54D10.7* | 22.85 | 95.73 | 1.78 | 0.63 | 0.004435 | 0.037573 |
| *hpo-15* | 1179.22 | 4550.33 | 1.78 | 0.51 | 0.000498 | 0.0071416 |
| *math-38* | 42.60 | 176.84 | 1.78 | 0.62 | 0.004088 | 0.0353752 |
| *lido-12* | 79.53 | 305.73 | 1.78 | 0.51 | 0.000481 | 0.0069645 |
| *F18G5.6* | 547.68 | 2106.34 | 1.78 | 0.51 | 0.000503 | 0.0071872 |
| *Y54G9A.5* | 110.58 | 425.54 | 1.78 | 0.51 | 0.000528 | 0.0074247 |
| *W05H9.1* | 2667.86 | 10468.32 | 1.78 | 0.55 | 0.001195 | 0.0142113 |
| *trt-1* | 99.21 | 384.90 | 1.77 | 0.53 | 0.000903 | 0.0114958 |
| *C44B9.3* | 167.05 | 631.65 | 1.77 | 0.49 | 0.00027 | 0.0044904 |
| *aptf-2* | 36.47 | 143.31 | 1.77 | 0.55 | 0.001404 | 0.0159995 |
| *Y68A4A.13* | 27.37 | 110.49 | 1.77 | 0.59 | 0.002795 | 0.0268016 |
| *bath-36* | 28.80 | 115.52 | 1.77 | 0.59 | 0.002514 | 0.0249521 |
| *F41B4.3* | 60.11 | 252.75 | 1.77 | 0.64 | 0.005997 | 0.0466119 |
| *str-176* | 105.73 | 417.94 | 1.77 | 0.57 | 0.001857 | 0.0198212 |
| *fbxc-50* | 85.10 | 323.44 | 1.77 | 0.50 | 0.000447 | 0.0066048 |
| *abf-2* | 23.83 | 96.78 | 1.76 | 0.61 | 0.003854 | 0.0339702 |
| *K11D12.13* | 48.96 | 191.91 | 1.76 | 0.56 | 0.001807 | 0.0194349 |
| *H21P03.2* | 316.28 | 1174.02 | 1.76 | 0.47 | 0.000198 | 0.0035199 |
| *alg-5* | 251.85 | 958.45 | 1.75 | 0.53 | 0.000864 | 0.0111103 |
| *K03H1.7* | 25.99 | 104.48 | 1.75 | 0.61 | 0.004079 | 0.0353277 |
| *F48E3.4* | 237.66 | 886.67 | 1.75 | 0.49 | 0.000367 | 0.0057119 |
| *swt-1* | 250.63 | 942.78 | 1.75 | 0.51 | 0.000621 | 0.0085183 |
| *fbxa-182* | 33.78 | 135.64 | 1.75 | 0.61 | 0.004389 | 0.0372657 |
| *cyb-3* | 2579.01 | 9473.50 | 1.74 | 0.47 | 0.000235 | 0.0040179 |
| *gpd-1* | 729.42 | 2654.17 | 1.74 | 0.46 | 0.000149 | 0.0027819 |
| *wee-1.3* | 948.52 | 3524.04 | 1.74 | 0.51 | 0.000603 | 0.0082797 |
| *F47B8.3* | 67.63 | 262.31 | 1.73 | 0.58 | 0.002692 | 0.0261107 |
| *F13A7.11* | 48.83 | 184.96 | 1.73 | 0.54 | 0.001366 | 0.0156897 |
| *F59A6.5* | 266.78 | 1018.66 | 1.73 | 0.56 | 0.001827 | 0.019611 |
| *C27D9.1* | 264.82 | 988.55 | 1.73 | 0.52 | 0.000857 | 0.011043 |
| *W03C9.2* | 219.49 | 808.47 | 1.73 | 0.49 | 0.000459 | 0.006708 |
| *Y43E12A.3* | 61.22 | 235.78 | 1.73 | 0.57 | 0.002511 | 0.0249362 |
| *F11E6.7* | 399.43 | 1492.41 | 1.73 | 0.52 | 0.000969 | 0.0120808 |
| *K05C4.4* | 40.40 | 158.65 | 1.73 | 0.60 | 0.004287 | 0.0366449 |
| *glc-1* | 41.79 | 167.70 | 1.73 | 0.63 | 0.006408 | 0.0487844 |
| *R05H10.1* | 34.21 | 134.10 | 1.73 | 0.60 | 0.004057 | 0.0352291 |
| *K02F6.7* | 81.22 | 318.54 | 1.73 | 0.61 | 0.004363 | 0.0371342 |
| *kca-1* | 323.55 | 1171.37 | 1.72 | 0.47 | 0.000255 | 0.0042998 |
| *F54E12.2* | 876.53 | 3225.99 | 1.72 | 0.51 | 0.000855 | 0.0110362 |
| *EEED8.14* | 51.53 | 193.96 | 1.72 | 0.55 | 0.001964 | 0.0207163 |
| *Y54G2A.11* | 1089.03 | 3976.66 | 1.71 | 0.50 | 0.000668 | 0.0090409 |
| *dgtr-1* | 103.33 | 376.94 | 1.71 | 0.51 | 0.000717 | 0.0096065 |
| *C56A3.4* | 231.13 | 848.80 | 1.71 | 0.52 | 0.001003 | 0.0124136 |
| *pqn-82* | 160.85 | 589.29 | 1.71 | 0.52 | 0.000972 | 0.0121124 |
| *Y71F9AL.2* | 81.37 | 298.67 | 1.70 | 0.53 | 0.001391 | 0.0158959 |
| *F55H12.2* | 125.10 | 483.40 | 1.70 | 0.61 | 0.005494 | 0.0436613 |
| *E01G4.5* | 38.16 | 141.51 | 1.70 | 0.55 | 0.002124 | 0.0219514 |
| *Y42H9B.3* | 185.09 | 658.77 | 1.70 | 0.48 | 0.000383 | 0.0058882 |
| *C37C3.9* | 264.46 | 941.27 | 1.70 | 0.48 | 0.000405 | 0.0061618 |
| *lgg-2* | 2174.10 | 7905.04 | 1.69 | 0.53 | 0.001322 | 0.0153594 |
| *ttr-17* | 1176.00 | 4232.32 | 1.69 | 0.51 | 0.000886 | 0.0113313 |
| *hrde-1* | 2549.38 | 9299.95 | 1.69 | 0.54 | 0.001587 | 0.0176709 |
| *neg-1* | 271.45 | 966.11 | 1.69 | 0.49 | 0.000601 | 0.0082724 |
| *cpg-4* | 263.30 | 951.63 | 1.69 | 0.52 | 0.001297 | 0.0151598 |
| *C39D10.7* | 3122.51 | 11081.87 | 1.69 | 0.49 | 0.000557 | 0.0077664 |
| *F17C11.11* | 256.38 | 922.43 | 1.69 | 0.52 | 0.001076 | 0.0131699 |

| | | | | | | |
|---|---|---|---|---|---|---|
| *kel-20* | 155.33 | 562.10 | 1.69 | 0.53 | 0.001428 | 0.0162302 |
| *cdc-7* | 174.51 | 624.24 | 1.68 | 0.51 | 0.001001 | 0.0123884 |
| *Y54G2A.12* | 51.86 | 192.47 | 1.68 | 0.58 | 0.003512 | 0.0317755 |
| *apx-1* | 388.73 | 1407.66 | 1.68 | 0.54 | 0.0018 | 0.0193925 |
| *vps-51* | 351.68 | 1240.44 | 1.68 | 0.49 | 0.0006 | 0.0082596 |
| *Y105C5A.8* | 81.74 | 304.32 | 1.68 | 0.58 | 0.00408 | 0.0353277 |
| *C08F8.3* | 219.53 | 782.94 | 1.68 | 0.51 | 0.001085 | 0.0132521 |
| *Y38E10A.22* | 864.70 | 3148.02 | 1.68 | 0.55 | 0.002306 | 0.0234851 |
| *cyp-31A2* | 185.37 | 656.83 | 1.67 | 0.50 | 0.000895 | 0.0114213 |
| *Y56A3A.33* | 33.24 | 124.54 | 1.67 | 0.60 | 0.005312 | 0.042634 |
| *clec-147* | 36.45 | 133.10 | 1.67 | 0.56 | 0.00291 | 0.0276075 |
| *C02F12.8* | 92.79 | 331.07 | 1.67 | 0.53 | 0.00149 | 0.0167655 |
| *D1081.7* | 1024.83 | 3645.71 | 1.67 | 0.52 | 0.001341 | 0.0154997 |
| *F47G3.4* | 65.42 | 232.90 | 1.67 | 0.52 | 0.001382 | 0.0158302 |
| *hop-1* | 101.95 | 358.11 | 1.67 | 0.50 | 0.000853 | 0.0110137 |
| *C30G7.4* | 60.74 | 224.97 | 1.66 | 0.59 | 0.004752 | 0.0394904 |
| *tth-1* | 1013.31 | 3455.41 | 1.66 | 0.45 | 0.000209 | 0.003663 |
| *duo-1* | 171.54 | 610.81 | 1.65 | 0.55 | 0.002662 | 0.0258805 |
| *ctl-3* | 45.20 | 159.87 | 1.65 | 0.54 | 0.002106 | 0.0218435 |
| *glp-1* | 622.37 | 2206.36 | 1.64 | 0.55 | 0.002635 | 0.0256982 |
| *glh-2* | 416.46 | 1464.33 | 1.64 | 0.53 | 0.002064 | 0.0215541 |
| *inx-8* | 275.83 | 937.78 | 1.64 | 0.47 | 0.000456 | 0.0067065 |
| *evl-18* | 267.30 | 922.16 | 1.64 | 0.50 | 0.001117 | 0.0135091 |
| *K04G7.1* | 266.87 | 910.47 | 1.64 | 0.48 | 0.000622 | 0.0085188 |
| *C30F12.4* | 369.23 | 1252.44 | 1.64 | 0.47 | 0.00046 | 0.0067229 |
| *T25B2.2* | 141.00 | 478.20 | 1.64 | 0.47 | 0.000506 | 0.0072026 |
| *math-27* | 352.56 | 1274.57 | 1.63 | 0.59 | 0.005861 | 0.0458339 |
| *ZK616.5* | 501.12 | 1710.45 | 1.63 | 0.49 | 0.000911 | 0.0115583 |
| *zim-2* | 149.60 | 531.96 | 1.63 | 0.57 | 0.00412 | 0.0356093 |
| *lin-41* | 1713.54 | 5785.44 | 1.63 | 0.47 | 0.000499 | 0.0071501 |
| *eri-7* | 100.60 | 349.75 | 1.63 | 0.53 | 0.002178 | 0.0223785 |
| *rnp-8* | 1021.66 | 3493.53 | 1.63 | 0.50 | 0.001107 | 0.0134445 |
| *san-1* | 85.39 | 291.99 | 1.63 | 0.50 | 0.001202 | 0.0142692 |
| *F35C11.5* | 293.20 | 992.35 | 1.62 | 0.49 | 0.00084 | 0.0108985 |
| *F53A2.3* | 37.22 | 130.19 | 1.62 | 0.55 | 0.003169 | 0.0293909 |
| *Y39E4B.5* | 193.45 | 655.27 | 1.62 | 0.49 | 0.000943 | 0.0118283 |
| *btb-19* | 100.07 | 342.64 | 1.62 | 0.51 | 0.001636 | 0.0180075 |
| *ogr-2* | 62.79 | 222.61 | 1.62 | 0.58 | 0.005035 | 0.0412114 |
| *gla-3* | 656.24 | 2231.12 | 1.62 | 0.50 | 0.00126 | 0.0148099 |
| *C09E7.4* | 100.01 | 348.61 | 1.61 | 0.56 | 0.003761 | 0.0333584 |
| *T04D3.8* | 37.51 | 130.64 | 1.61 | 0.56 | 0.003669 | 0.0328332 |
| *set-22* | 163.83 | 553.51 | 1.61 | 0.50 | 0.001279 | 0.015 |
| *H05L14.2* | 272.34 | 939.97 | 1.61 | 0.54 | 0.003002 | 0.0282671 |
| *mcm-3* | 744.81 | 2503.33 | 1.61 | 0.49 | 0.00112 | 0.0135199 |
| *F56C9.3* | 528.95 | 1783.41 | 1.61 | 0.51 | 0.001554 | 0.0173932 |
| *ptc-1* | 2727.03 | 9140.56 | 1.60 | 0.50 | 0.001361 | 0.0156591 |
| *ddo-3* | 185.61 | 615.84 | 1.60 | 0.48 | 0.000877 | 0.0112362 |
| *mcm-5* | 554.05 | 1838.75 | 1.60 | 0.48 | 0.000906 | 0.0115157 |
| *F26A1.1* | 182.43 | 603.64 | 1.60 | 0.48 | 0.000832 | 0.0108134 |
| *F09F7.6* | 134.26 | 455.77 | 1.60 | 0.54 | 0.002916 | 0.0276437 |
| *efk-1* | 785.43 | 2569.01 | 1.59 | 0.46 | 0.000501 | 0.0071558 |
| *F41G4.8* | 32.82 | 114.13 | 1.59 | 0.58 | 0.006055 | 0.0468962 |
| *thn-2* | 215.56 | 720.98 | 1.59 | 0.51 | 0.001896 | 0.0201306 |
| *B0001.7* | 281.22 | 954.90 | 1.59 | 0.54 | 0.003351 | 0.0306277 |
| *acly-2* | 996.09 | 3286.88 | 1.59 | 0.48 | 0.001018 | 0.0125789 |
| *T04G9.7* | 161.02 | 531.35 | 1.59 | 0.48 | 0.001026 | 0.0126367 |

| | | | | | | |
|---|---|---|---|---|---|---|
| *T16G12.4* | 66.66 | 224.76 | 1.59 | 0.53 | 0.002875 | 0.0273483 |
| *C14B1.3* | 244.54 | 831.57 | 1.59 | 0.55 | 0.003902 | 0.0343074 |
| *lrr-1* | 202.84 | 667.90 | 1.59 | 0.49 | 0.001118 | 0.0135091 |
| *B0393.3* | 459.92 | 1512.95 | 1.59 | 0.49 | 0.001189 | 0.0141519 |
| *deps-1* | 782.05 | 2564.65 | 1.58 | 0.49 | 0.001208 | 0.0143108 |
| *rad-26* | 545.06 | 1848.28 | 1.58 | 0.55 | 0.004386 | 0.0372546 |
| *hcp-3* | 317.83 | 1039.42 | 1.58 | 0.48 | 0.001101 | 0.013379 |
| *F54B8.4* | 47.14 | 161.95 | 1.58 | 0.58 | 0.006345 | 0.0483801 |
| *hil-1* | 48.70 | 163.98 | 1.58 | 0.55 | 0.003945 | 0.0345318 |
| *Y75B12B.1* | 467.44 | 1511.87 | 1.58 | 0.47 | 0.000734 | 0.0097578 |
| *zen-4* | 500.33 | 1671.50 | 1.57 | 0.54 | 0.003357 | 0.0306277 |
| *lin-3* | 148.88 | 495.54 | 1.57 | 0.53 | 0.003035 | 0.0284836 |
| *C44B7.5* | 311.96 | 1003.99 | 1.57 | 0.47 | 0.000772 | 0.0101683 |
| *rtel-1* | 300.38 | 988.11 | 1.57 | 0.52 | 0.002395 | 0.0241281 |
| *T24C4.5* | 121.71 | 398.70 | 1.57 | 0.51 | 0.002087 | 0.0217423 |
| *inx-9* | 123.16 | 398.72 | 1.57 | 0.48 | 0.001226 | 0.0144835 |
| *rskn-1* | 713.85 | 2276.08 | 1.56 | 0.45 | 0.000516 | 0.0072995 |
| *sdz-27* | 202.17 | 649.94 | 1.56 | 0.47 | 0.000927 | 0.011681 |
| *ect-2* | 755.84 | 2474.58 | 1.56 | 0.51 | 0.002365 | 0.0239577 |
| *nop-1* | 320.97 | 1053.11 | 1.56 | 0.52 | 0.002582 | 0.0253475 |
| *lab-2* | 120.17 | 395.62 | 1.56 | 0.53 | 0.00296 | 0.027968 |
| *C32F10.4* | 1058.93 | 3466.30 | 1.56 | 0.51 | 0.002389 | 0.0241259 |
| *F45F2.11* | 478.81 | 1572.59 | 1.56 | 0.53 | 0.003036 | 0.0284836 |
| *nos-1* | 145.89 | 472.41 | 1.56 | 0.50 | 0.001726 | 0.01878 |
| *mcm-7* | 1291.87 | 4129.17 | 1.56 | 0.47 | 0.000874 | 0.0112101 |
| *C14B1.2* | 216.73 | 692.01 | 1.55 | 0.48 | 0.001111 | 0.0134629 |
| *bath-30* | 127.65 | 409.83 | 1.55 | 0.49 | 0.001518 | 0.0170315 |
| *C16C8.4* | 108.75 | 350.87 | 1.55 | 0.50 | 0.002048 | 0.0214244 |
| *Y65A5A.2* | 217.72 | 702.78 | 1.55 | 0.51 | 0.002263 | 0.023153 |
| *C50B6.3* | 324.03 | 1041.25 | 1.55 | 0.50 | 0.00188 | 0.0200004 |
| *elpc-4* | 196.48 | 625.54 | 1.55 | 0.48 | 0.001328 | 0.0154187 |
| *xnd-1* | 822.32 | 2649.99 | 1.54 | 0.51 | 0.002531 | 0.0250344 |
| *egg-6* | 1277.86 | 4086.83 | 1.54 | 0.49 | 0.001803 | 0.0194122 |
| *Y54F10AM.11* | 54.14 | 176.97 | 1.54 | 0.54 | 0.004409 | 0.0373742 |
| *W03D2.6* | 173.52 | 553.40 | 1.54 | 0.50 | 0.002016 | 0.021179 |
| *mre-11* | 316.26 | 1012.16 | 1.54 | 0.51 | 0.002416 | 0.0242282 |
| *Y18D10A.11* | 675.19 | 2138.58 | 1.54 | 0.48 | 0.001485 | 0.0167387 |
| *lig-1* | 505.19 | 1603.15 | 1.54 | 0.49 | 0.001721 | 0.0187334 |
| *cbp-2* | 195.55 | 615.82 | 1.53 | 0.47 | 0.00123 | 0.0145239 |
| *scc-1* | 292.53 | 933.01 | 1.53 | 0.51 | 0.002391 | 0.0241259 |
| *spo-11* | 120.69 | 391.22 | 1.53 | 0.54 | 0.004571 | 0.0384363 |
| *Y37D8A.19* | 308.74 | 980.64 | 1.53 | 0.50 | 0.002167 | 0.0222935 |
| *gpd-4* | 483.12 | 1515.22 | 1.53 | 0.48 | 0.001351 | 0.0155646 |
| *mut-2* | 177.33 | 568.66 | 1.53 | 0.53 | 0.003765 | 0.0333584 |
| *ule-5* | 812.40 | 2578.63 | 1.53 | 0.51 | 0.00256 | 0.0252696 |
| *skpt-1* | 454.22 | 1426.50 | 1.53 | 0.48 | 0.001582 | 0.0176398 |
| *pal-1* | 365.76 | 1138.92 | 1.53 | 0.46 | 0.000995 | 0.0123401 |
| *sun-1* | 322.16 | 1030.36 | 1.52 | 0.53 | 0.003799 | 0.0336084 |
| *plk-3* | 1188.45 | 3781.82 | 1.52 | 0.52 | 0.003169 | 0.0293909 |
| *syp-5* | 271.83 | 872.78 | 1.52 | 0.54 | 0.005177 | 0.0419316 |
| *fem-3* | 145.26 | 462.21 | 1.52 | 0.53 | 0.004161 | 0.0358422 |
| *B0511.19* | 148.17 | 464.97 | 1.52 | 0.50 | 0.002425 | 0.0242689 |
| *F56F11.4* | 177.84 | 558.24 | 1.51 | 0.50 | 0.002597 | 0.0254301 |
| *zif-1* | 589.78 | 1820.77 | 1.51 | 0.47 | 0.001199 | 0.0142419 |
| *hcp-1* | 1000.62 | 3154.22 | 1.51 | 0.52 | 0.004097 | 0.0354322 |
| *D1054.10* | 906.36 | 2796.34 | 1.51 | 0.48 | 0.001586 | 0.0176692 |

| | | | | | | |
|---|---|---|---|---|---|---|
| *unc-61* | 586.60 | 1810.28 | 1.50 | 0.48 | 0.001924 | 0.0203794 |
| *C42C1.8* | 244.09 | 771.50 | 1.50 | 0.54 | 0.005318 | 0.042634 |
| *C17G1.2* | 103.82 | 319.92 | 1.50 | 0.49 | 0.002088 | 0.0217423 |
| *simr-1* | 690.00 | 2169.95 | 1.50 | 0.53 | 0.004953 | 0.0407386 |
| *catp-3* | 353.28 | 1111.80 | 1.49 | 0.54 | 0.005771 | 0.0453416 |
| *orc-5* | 262.30 | 803.03 | 1.49 | 0.48 | 0.001955 | 0.0206337 |
| *Y47G6A.14* | 94.21 | 291.75 | 1.49 | 0.51 | 0.003355 | 0.0306277 |
| *Y65B4BL.4* | 244.21 | 755.06 | 1.49 | 0.51 | 0.003168 | 0.0293909 |
| *F55B11.2* | 524.81 | 1592.30 | 1.49 | 0.46 | 0.001281 | 0.0150126 |
| *Y62F5A.12* | 97.60 | 302.92 | 1.49 | 0.52 | 0.003923 | 0.0343986 |
| *D1086.10* | 4514.42 | 13605.36 | 1.49 | 0.45 | 0.000857 | 0.011043 |
| *dsb-1* | 370.16 | 1129.97 | 1.49 | 0.48 | 0.002109 | 0.0218435 |
| *cnc-4* | 188.76 | 591.87 | 1.49 | 0.54 | 0.006377 | 0.0485769 |
| *nasp-1* | 532.49 | 1609.27 | 1.48 | 0.47 | 0.001517 | 0.0170221 |
| *Y95D11A.3* | 125.79 | 383.11 | 1.48 | 0.49 | 0.002732 | 0.0263764 |
| *C04F12.1* | 405.23 | 1227.58 | 1.48 | 0.48 | 0.002151 | 0.0221841 |
| *F57C2.4* | 65.09 | 201.41 | 1.48 | 0.53 | 0.005266 | 0.042351 |
| *R11H6.4* | 85.93 | 263.30 | 1.48 | 0.51 | 0.003814 | 0.0336932 |
| *com-1* | 222.46 | 689.84 | 1.48 | 0.54 | 0.006011 | 0.0466713 |
| *puf-8* | 634.92 | 1909.23 | 1.48 | 0.47 | 0.001686 | 0.018429 |
| *wago-1* | 1858.22 | 5693.81 | 1.48 | 0.51 | 0.004134 | 0.0356744 |
| *Y59E9AL.3* | 66.53 | 203.73 | 1.48 | 0.51 | 0.004046 | 0.0351893 |
| *fog-2* | 118.36 | 361.44 | 1.47 | 0.51 | 0.003868 | 0.0340667 |
| *B0238.9* | 479.55 | 1475.54 | 1.47 | 0.53 | 0.005251 | 0.0422794 |
| *atg-13* | 248.45 | 752.07 | 1.47 | 0.50 | 0.003066 | 0.0286583 |
| *klp-15* | 684.59 | 2052.32 | 1.47 | 0.48 | 0.002031 | 0.0213116 |
| *epg-4* | 499.01 | 1502.66 | 1.47 | 0.49 | 0.002618 | 0.0255665 |
| *F10E9.7* | 155.45 | 466.82 | 1.46 | 0.49 | 0.00274 | 0.026421 |
| *F55B11.3* | 181.66 | 553.91 | 1.46 | 0.53 | 0.00538 | 0.0430417 |
| *pgl-3* | 787.19 | 2407.28 | 1.46 | 0.53 | 0.006115 | 0.0472509 |
| *ZK1307.9* | 235.81 | 699.54 | 1.46 | 0.46 | 0.001599 | 0.0177661 |
| *lem-3* | 378.16 | 1129.63 | 1.46 | 0.48 | 0.002416 | 0.0242282 |
| *Y71F9AL.10* | 404.82 | 1207.98 | 1.46 | 0.48 | 0.002406 | 0.0241907 |
| *panl-2* | 387.94 | 1158.26 | 1.46 | 0.49 | 0.002665 | 0.0258977 |
| *ZK384.7* | 79.74 | 239.86 | 1.46 | 0.50 | 0.003885 | 0.0341814 |
| *vha-18* | 172.84 | 516.13 | 1.45 | 0.49 | 0.00329 | 0.0303225 |
| *C29A12.1* | 212.70 | 631.72 | 1.45 | 0.48 | 0.002609 | 0.0255175 |
| *ZC373.2* | 394.94 | 1189.46 | 1.45 | 0.52 | 0.004841 | 0.0400539 |
| *T20D4.11* | 151.91 | 455.28 | 1.45 | 0.51 | 0.004396 | 0.0373052 |
| *F28H6.4* | 146.49 | 438.02 | 1.45 | 0.50 | 0.004052 | 0.0352227 |
| *dnj-26* | 76.26 | 230.37 | 1.45 | 0.53 | 0.006023 | 0.0467455 |
| *bath-41* | 334.17 | 985.36 | 1.45 | 0.47 | 0.002113 | 0.0218627 |
| *Y58A7A.3* | 237.96 | 712.82 | 1.45 | 0.51 | 0.004526 | 0.0381477 |
| *klp-16* | 558.68 | 1640.03 | 1.45 | 0.46 | 0.001671 | 0.0183014 |
| *him-10* | 207.11 | 620.44 | 1.45 | 0.52 | 0.005039 | 0.0412165 |
| *ttr-26* | 79.07 | 237.53 | 1.44 | 0.52 | 0.00591 | 0.0461245 |
| *maph-1.3* | 660.80 | 1943.71 | 1.44 | 0.48 | 0.002421 | 0.0242607 |
| *sumv-1* | 584.45 | 1729.96 | 1.44 | 0.49 | 0.003359 | 0.0306277 |
| *Y55D9A.2* | 169.54 | 504.93 | 1.44 | 0.51 | 0.004987 | 0.0408864 |
| *adal-1* | 391.82 | 1140.50 | 1.43 | 0.47 | 0.002164 | 0.0222751 |
| *cey-2* | 4914.41 | 14197.32 | 1.43 | 0.45 | 0.001489 | 0.0167639 |
| *F25H2.12* | 137.75 | 400.99 | 1.43 | 0.47 | 0.002523 | 0.0249896 |
| *best-20* | 83.59 | 247.47 | 1.43 | 0.51 | 0.005465 | 0.0435536 |
| *C14C11.2* | 206.13 | 607.30 | 1.43 | 0.50 | 0.00463 | 0.0387702 |
| *dvc-1* | 274.94 | 803.13 | 1.43 | 0.48 | 0.003131 | 0.0291252 |
| *R02F11.4* | 293.76 | 854.82 | 1.43 | 0.47 | 0.002594 | 0.0254188 |

| | | | | | | |
|---|---|---|---|---|---|---|
| *F17E9.4* | 275.96 | 796.64 | 1.43 | 0.45 | 0.001603 | 0.0177827 |
| *F56C9.6* | 796.12 | 2321.65 | 1.43 | 0.48 | 0.002989 | 0.0281827 |
| *rei-1* | 387.11 | 1114.97 | 1.43 | 0.45 | 0.001456 | 0.0164777 |
| *F38E9.1* | 239.00 | 695.33 | 1.43 | 0.48 | 0.003041 | 0.0285156 |
| *D1086.11* | 512.31 | 1491.39 | 1.42 | 0.49 | 0.003691 | 0.0329303 |
| *klp-18* | 807.78 | 2359.31 | 1.42 | 0.50 | 0.004319 | 0.0368341 |
| *inx-14* | 311.44 | 917.60 | 1.42 | 0.52 | 0.006145 | 0.0473362 |
| *zim-1* | 268.83 | 790.49 | 1.42 | 0.51 | 0.00576 | 0.0453191 |
| *fut-3* | 222.67 | 642.47 | 1.42 | 0.47 | 0.002427 | 0.0242819 |
| *lab-1* | 86.49 | 253.17 | 1.42 | 0.51 | 0.005073 | 0.0413662 |
| *sas-5* | 377.23 | 1094.51 | 1.42 | 0.49 | 0.003988 | 0.0347882 |
| *T28H10.3* | 1196.39 | 3510.98 | 1.42 | 0.52 | 0.006484 | 0.0491474 |
| *Y57G11B.5* | 285.97 | 827.57 | 1.41 | 0.49 | 0.003764 | 0.0333584 |
| *D2030.2* | 741.42 | 2148.82 | 1.41 | 0.50 | 0.004369 | 0.0371342 |
| *lact-3* | 370.56 | 1057.01 | 1.41 | 0.46 | 0.00194 | 0.0205229 |
| *asp-12* | 159.64 | 461.54 | 1.41 | 0.49 | 0.00402 | 0.035003 |
| *Y110A2AR.1* | 294.11 | 849.04 | 1.41 | 0.49 | 0.003802 | 0.0336084 |
| *F32E10.5* | 82.46 | 240.84 | 1.41 | 0.52 | 0.006295 | 0.0481632 |
| *rnf-1* | 223.21 | 647.94 | 1.41 | 0.50 | 0.005141 | 0.0417432 |
| *R06F6.12* | 191.20 | 549.77 | 1.41 | 0.48 | 0.003332 | 0.0305665 |
| *rsa-1* | 286.33 | 820.68 | 1.40 | 0.48 | 0.003639 | 0.0326539 |
| *hpr-17* | 117.95 | 341.69 | 1.40 | 0.51 | 0.006191 | 0.047559 |
| *44440* | 619.39 | 1780.70 | 1.40 | 0.50 | 0.004612 | 0.0386836 |
| *taf-8* | 257.65 | 738.24 | 1.40 | 0.49 | 0.004132 | 0.0356744 |
| *lem-4* | 244.92 | 699.23 | 1.40 | 0.48 | 0.003499 | 0.0317142 |
| *cec-7* | 240.92 | 687.72 | 1.40 | 0.48 | 0.003749 | 0.0333339 |
| *Y5F2A.4* | 267.16 | 757.83 | 1.40 | 0.47 | 0.002866 | 0.0272894 |
| *nyn-2* | 244.17 | 694.80 | 1.40 | 0.48 | 0.00356 | 0.0320619 |
| *C13G5.2* | 152.09 | 432.57 | 1.40 | 0.48 | 0.003518 | 0.0317931 |
| *snpc-1.2* | 412.46 | 1180.31 | 1.39 | 0.50 | 0.005091 | 0.0414677 |
| *frm-10* | 506.89 | 1453.69 | 1.39 | 0.50 | 0.005594 | 0.0442915 |
| *T04A8.8* | 380.56 | 1075.60 | 1.39 | 0.47 | 0.002898 | 0.0275239 |
| *Y71F9AL.4* | 151.85 | 435.17 | 1.39 | 0.50 | 0.00575 | 0.0452887 |
| *polk-1* | 156.60 | 447.57 | 1.39 | 0.50 | 0.005432 | 0.0433464 |
| *gbas-1* | 175.36 | 498.68 | 1.38 | 0.50 | 0.005853 | 0.0457937 |
| *cku-80* | 240.17 | 680.32 | 1.38 | 0.49 | 0.005065 | 0.0413209 |
| *Y62H9A.5* | 778.00 | 2204.92 | 1.38 | 0.49 | 0.005176 | 0.0419316 |
| *srgp-1* | 1347.09 | 3769.58 | 1.38 | 0.46 | 0.00286 | 0.0272506 |
| *mina-1* | 489.78 | 1392.49 | 1.38 | 0.51 | 0.00629 | 0.0481528 |
| *ptr-8* | 308.35 | 868.93 | 1.38 | 0.49 | 0.004532 | 0.0381693 |
| *imp-3* | 326.58 | 907.08 | 1.37 | 0.46 | 0.002806 | 0.0268904 |
| *ule-3* | 1959.29 | 5504.58 | 1.37 | 0.49 | 0.005245 | 0.0422733 |
| *klp-19* | 1059.09 | 2953.87 | 1.37 | 0.47 | 0.003762 | 0.0333584 |
| *sld-2* | 104.28 | 294.34 | 1.37 | 0.51 | 0.006604 | 0.0497605 |
| *perm-4* | 5308.76 | 14660.49 | 1.37 | 0.45 | 0.002374 | 0.0240181 |
| *ndc-80* | 265.02 | 745.28 | 1.37 | 0.50 | 0.006214 | 0.0477134 |
| *dsh-2* | 257.64 | 720.81 | 1.37 | 0.49 | 0.005002 | 0.0409847 |
| *ari-1.4* | 165.91 | 464.16 | 1.37 | 0.49 | 0.005244 | 0.0422733 |
| *chk-1* | 295.12 | 819.00 | 1.37 | 0.47 | 0.003929 | 0.0344294 |
| *col-143* | 2693.55 | 7457.06 | 1.36 | 0.47 | 0.003731 | 0.0332202 |
| *gfat-1* | 2579.27 | 7120.20 | 1.36 | 0.46 | 0.003299 | 0.0303734 |
| *F20D1.3* | 3155.79 | 8799.36 | 1.36 | 0.49 | 0.005696 | 0.0449365 |
| *T23B3.2* | 538.56 | 1486.87 | 1.36 | 0.47 | 0.003984 | 0.0347877 |
| *lin-15B* | 324.57 | 895.82 | 1.36 | 0.48 | 0.004629 | 0.0387702 |
| *ima-1* | 303.38 | 839.88 | 1.35 | 0.50 | 0.006459 | 0.0489925 |
| *glh-3* | 258.32 | 712.48 | 1.35 | 0.49 | 0.005813 | 0.0455983 |

| | | | | | | |
|---|---|---|---|---|---|---|
| *tasp-1* | 409.30 | 1123.99 | 1.35 | 0.48 | 0.004972 | 0.0408442 |
| *wrm-1* | 435.40 | 1201.95 | 1.35 | 0.50 | 0.006443 | 0.0489832 |
| *44287* | 539.89 | 1473.61 | 1.34 | 0.48 | 0.005615 | 0.0444099 |
| *W05F2.6* | 454.97 | 1227.74 | 1.33 | 0.47 | 0.004191 | 0.0360659 |
| *T07C4.3* | 1147.77 | 3088.72 | 1.33 | 0.47 | 0.005044 | 0.0412166 |
| *mcm-6* | 616.81 | 1662.40 | 1.32 | 0.48 | 0.00596 | 0.046444 |
| *F46F11.8* | 278.48 | 742.53 | 1.32 | 0.47 | 0.004694 | 0.0391554 |
| *F44B9.8* | 301.14 | 805.17 | 1.32 | 0.47 | 0.005477 | 0.0435929 |
| *ikke-1* | 998.24 | 2653.08 | 1.31 | 0.46 | 0.0046 | 0.0386055 |
| *C48B4.7* | 127.84 | 341.04 | 1.31 | 0.48 | 0.006448 | 0.0489925 |
| *tag-196* | 824.05 | 2169.98 | 1.30 | 0.45 | 0.004057 | 0.0352291 |
| *usip-1* | 755.87 | 1993.49 | 1.30 | 0.46 | 0.004633 | 0.0387774 |
| *mrp-4* | 916.23 | 2401.57 | 1.30 | 0.45 | 0.003802 | 0.0336084 |
| *B0513.4* | 228.20 | 602.70 | 1.30 | 0.48 | 0.006524 | 0.049359 |
| *msa-1* | 465.04 | 1223.92 | 1.30 | 0.47 | 0.005701 | 0.0449485 |
| *cya-1* | 532.11 | 1398.31 | 1.29 | 0.47 | 0.006288 | 0.0481528 |
| *pup-2* | 2321.31 | 6021.19 | 1.29 | 0.44 | 0.003414 | 0.0310539 |
| *psf-3* | 249.32 | 652.42 | 1.29 | 0.47 | 0.006306 | 0.0482006 |
| *picc-1* | 453.92 | 1183.09 | 1.29 | 0.46 | 0.005329 | 0.0426776 |
| *pigv-1* | 570.13 | 1485.45 | 1.29 | 0.46 | 0.005317 | 0.042634 |
| *spas-1* | 403.58 | 1051.34 | 1.28 | 0.47 | 0.006525 | 0.049359 |
| *cey-3* | 1638.92 | 4223.49 | 1.28 | 0.45 | 0.004304 | 0.0367482 |
| *M01F1.9* | 517.95 | 1330.22 | 1.28 | 0.44 | 0.003904 | 0.0343127 |
| *F21D5.1* | 553.25 | 1423.84 | 1.27 | 0.46 | 0.005873 | 0.0458835 |
| *F39G3.3* | 786.32 | 2024.99 | 1.27 | 0.46 | 0.006185 | 0.0475503 |
| *C35B1.4* | 473.97 | 1220.64 | 1.27 | 0.47 | 0.006461 | 0.0489925 |
| *col-119* | 9554.95 | 24431.76 | 1.27 | 0.45 | 0.004439 | 0.0375853 |
| *K04C2.3* | 403.73 | 1027.70 | 1.26 | 0.45 | 0.004826 | 0.0399929 |
| *kup-1* | 473.66 | 1205.90 | 1.26 | 0.45 | 0.005594 | 0.0442915 |
| *mel-26* | 1040.27 | 2608.60 | 1.25 | 0.44 | 0.004242 | 0.0362738 |
| *zyg-11* | 654.90 | 1635.33 | 1.24 | 0.44 | 0.004856 | 0.0401308 |
| *fsn-1* | 516.27 | 1288.61 | 1.24 | 0.45 | 0.005767 | 0.0453301 |
| *F25H2.6* | 601.56 | 1489.84 | 1.23 | 0.44 | 0.005114 | 0.0415716 |
| *F57C2.5* | 2044.73 | 842.70 | -1.20 | 0.44 | 0.006249 | 0.0479075 |
| *Y71H2AM.11* | 1413.70 | 579.43 | -1.21 | 0.44 | 0.006311 | 0.0482134 |
| *dnj-20* | 19724.70 | 8071.62 | -1.21 | 0.43 | 0.00506 | 0.0413048 |
| *K11H3.3* | 2658.89 | 1060.72 | -1.24 | 0.46 | 0.006536 | 0.049395 |
| *Y51F10.7* | 4576.99 | 1824.84 | -1.24 | 0.45 | 0.006142 | 0.0473362 |
| *C14B9.2* | 2352.56 | 940.00 | -1.24 | 0.44 | 0.004803 | 0.0398483 |
| *adsl-1* | 2948.74 | 1173.64 | -1.24 | 0.45 | 0.005957 | 0.0464408 |
| *myrf-1* | 4026.74 | 1603.48 | -1.25 | 0.44 | 0.004707 | 0.0392453 |
| *fipr-21* | 3180.35 | 1267.73 | -1.25 | 0.44 | 0.004366 | 0.0371342 |
| *sym-2* | 1232.65 | 489.99 | -1.25 | 0.44 | 0.004534 | 0.0381693 |
| *col-104* | 18494.40 | 7288.75 | -1.25 | 0.46 | 0.00654 | 0.0493967 |
| *erm-1* | 15370.51 | 6089.42 | -1.25 | 0.44 | 0.004233 | 0.0362738 |
| *col-14* | 21701.96 | 8543.33 | -1.26 | 0.45 | 0.005557 | 0.044069 |
| *F09E5.3* | 1171.51 | 461.88 | -1.26 | 0.45 | 0.005008 | 0.0410156 |
| *C55B7.3* | 548.69 | 215.60 | -1.26 | 0.46 | 0.006009 | 0.0466713 |
| *trak-1* | 3912.13 | 1541.75 | -1.26 | 0.44 | 0.004536 | 0.0381693 |
| *ifd-2* | 1695.86 | 664.99 | -1.26 | 0.46 | 0.006116 | 0.0472509 |
| *C53D5.5* | 1069.59 | 420.56 | -1.26 | 0.45 | 0.005107 | 0.0415367 |
| *nucb-1* | 2801.73 | 1105.23 | -1.26 | 0.44 | 0.003845 | 0.0339305 |
| *cgr-1* | 1505.67 | 588.81 | -1.27 | 0.45 | 0.004483 | 0.0378481 |
| *gly-6* | 2030.08 | 788.26 | -1.27 | 0.47 | 0.006587 | 0.0496577 |
| *ncs-2* | 13591.73 | 5271.20 | -1.27 | 0.47 | 0.00664 | 0.0499542 |
| *F19H8.2* | 889.08 | 346.42 | -1.27 | 0.45 | 0.004241 | 0.0362738 |

| | | | | | | |
|---|---|---|---|---|---|---|
| *nra-4* | 5461.50 | 2132.05 | -1.28 | 0.43 | 0.003022 | 0.0284225 |
| *Y11D7A.9* | 720.76 | 278.30 | -1.28 | 0.46 | 0.005981 | 0.0465369 |
| *Y54F10AM.5* | 2479.58 | 956.53 | -1.28 | 0.47 | 0.006054 | 0.0468962 |
| *elo-3* | 3305.46 | 1283.57 | -1.28 | 0.44 | 0.003972 | 0.0347306 |
| *nkb-3* | 8407.89 | 3246.13 | -1.28 | 0.46 | 0.005172 | 0.0419316 |
| *vha-19* | 14638.35 | 5664.92 | -1.28 | 0.45 | 0.00446 | 0.0377115 |
| *col-115* | 779.66 | 301.78 | -1.28 | 0.45 | 0.004348 | 0.0370378 |
| *gpi-1* | 5540.87 | 2122.34 | -1.29 | 0.47 | 0.00564 | 0.0445648 |
| *tts-2* | 8805.34 | 3390.49 | -1.29 | 0.44 | 0.003504 | 0.031731 |
| *zipt-7.2* | 3034.60 | 1154.84 | -1.30 | 0.47 | 0.00532 | 0.042634 |
| *vgln-1* | 45018.24 | 17271.61 | -1.30 | 0.44 | 0.003176 | 0.0294343 |
| *Y53F4B.27* | 696.22 | 263.26 | -1.30 | 0.48 | 0.006186 | 0.0475503 |
| *C10G8.8* | 3408.71 | 1306.26 | -1.30 | 0.43 | 0.002617 | 0.0255665 |
| *cisd-3.2* | 3215.58 | 1216.07 | -1.30 | 0.47 | 0.005821 | 0.04564 |
| *atgp-2* | 1050.42 | 398.34 | -1.30 | 0.46 | 0.004975 | 0.0408442 |
| *ctb-1* | 52916.35 | 20234.22 | -1.30 | 0.43 | 0.002516 | 0.0249521 |
| *T04F8.2* | 768.74 | 290.85 | -1.31 | 0.45 | 0.003934 | 0.0344522 |
| *blmp-1* | 1941.61 | 734.84 | -1.31 | 0.45 | 0.003693 | 0.0329303 |
| *egl-30* | 4373.10 | 1638.90 | -1.31 | 0.48 | 0.006141 | 0.0473362 |
| *tre-4* | 316.47 | 118.46 | -1.31 | 0.48 | 0.006457 | 0.0489925 |
| *pdi-2* | 82890.79 | 31018.28 | -1.31 | 0.48 | 0.006529 | 0.0493618 |
| *gba-3* | 342.77 | 128.26 | -1.32 | 0.47 | 0.005506 | 0.0437277 |
| *tram-1* | 5853.50 | 2219.86 | -1.32 | 0.43 | 0.002278 | 0.0232744 |
| *F53F4.16* | 1769.31 | 662.96 | -1.32 | 0.46 | 0.004029 | 0.035064 |
| *ampd-1* | 8415.32 | 3129.20 | -1.33 | 0.47 | 0.0049 | 0.0404123 |
| *C14F11.6* | 749.75 | 280.72 | -1.33 | 0.45 | 0.003222 | 0.0297484 |
| *nhr-68* | 1694.85 | 629.15 | -1.33 | 0.47 | 0.005245 | 0.0422733 |
| *pqn-35* | 676.76 | 251.00 | -1.33 | 0.48 | 0.005424 | 0.043305 |
| *bed-3* | 1176.95 | 439.80 | -1.33 | 0.45 | 0.003366 | 0.030656 |
| *T19B10.2* | 9729.10 | 3642.92 | -1.33 | 0.44 | 0.002705 | 0.0262014 |
| *catp-5* | 1203.13 | 447.46 | -1.33 | 0.46 | 0.004134 | 0.0356744 |
| *scav-1* | 2669.97 | 985.18 | -1.33 | 0.48 | 0.005841 | 0.045726 |
| *gale-1* | 2619.06 | 979.39 | -1.33 | 0.44 | 0.002683 | 0.0260369 |
| *nduf-6* | 1418.28 | 523.85 | -1.33 | 0.47 | 0.004852 | 0.0401256 |
| *Y94H6A.8* | 6656.75 | 2447.52 | -1.33 | 0.48 | 0.005865 | 0.0458389 |
| *pgk-1* | 4573.65 | 1680.97 | -1.34 | 0.48 | 0.005492 | 0.0436613 |
| *cgt-2* | 317.25 | 116.26 | -1.34 | 0.49 | 0.006055 | 0.0468962 |
| *Y43C5A.2* | 5289.45 | 1943.08 | -1.34 | 0.48 | 0.005178 | 0.0419316 |
| *T16G1.2* | 925.28 | 338.56 | -1.34 | 0.48 | 0.005552 | 0.0440486 |
| *T19A5.3* | 2331.74 | 865.68 | -1.34 | 0.44 | 0.002215 | 0.0227226 |
| *mab-31* | 2965.78 | 1085.81 | -1.34 | 0.48 | 0.004977 | 0.0408442 |
| *col-48* | 9971.42 | 3705.73 | -1.34 | 0.43 | 0.001715 | 0.0187012 |
| *smf-3* | 275.53 | 100.41 | -1.35 | 0.49 | 0.005532 | 0.0439142 |
| *dif-1* | 3009.43 | 1093.75 | -1.35 | 0.49 | 0.005755 | 0.0453091 |
| *dpy-5* | 25000.03 | 9144.30 | -1.35 | 0.47 | 0.004157 | 0.0358283 |
| *Y119D3B.13* | 302.70 | 109.52 | -1.36 | 0.49 | 0.005205 | 0.0420371 |
| *ctsa-4.2* | 1318.30 | 479.29 | -1.36 | 0.46 | 0.002836 | 0.0270733 |
| *Y38H6C.16* | 269.69 | 96.49 | -1.36 | 0.50 | 0.006164 | 0.0474409 |
| *acly-1* | 6825.19 | 2471.92 | -1.36 | 0.46 | 0.003352 | 0.0306277 |
| *eps-8* | 6461.19 | 2323.04 | -1.36 | 0.48 | 0.004799 | 0.0398336 |
| *T25C12.3* | 37401.43 | 13515.74 | -1.36 | 0.47 | 0.003655 | 0.0327619 |
| *lep-5* | 11203.03 | 4012.36 | -1.37 | 0.49 | 0.005263 | 0.042351 |
| *Y56A3A.19* | 3634.64 | 1298.39 | -1.37 | 0.50 | 0.005777 | 0.0453669 |
| *C13C12.2* | 364.28 | 130.56 | -1.37 | 0.49 | 0.004918 | 0.0405058 |
| *Y69A2AR.18* | 43708.18 | 15647.34 | -1.37 | 0.49 | 0.005191 | 0.0419916 |
| *cyp-25A2* | 576.70 | 208.70 | -1.37 | 0.46 | 0.002771 | 0.0266401 |

| | | | | | | |
|---|---|---|---|---|---|---|
| *cyp-33A1* | 269.44 | 96.32 | -1.37 | 0.49 | 0.004888 | 0.0403546 |
| *ent-2* | 3386.95 | 1209.57 | -1.37 | 0.49 | 0.004721 | 0.0393222 |
| *ZC434.9* | 1709.20 | 615.11 | -1.38 | 0.46 | 0.002731 | 0.0263764 |
| *C03H12.1* | 991.25 | 357.00 | -1.38 | 0.45 | 0.002352 | 0.0238518 |
| *cox-4* | 35531.89 | 12533.32 | -1.38 | 0.50 | 0.006116 | 0.0472509 |
| *mab-10* | 2085.34 | 751.19 | -1.38 | 0.44 | 0.001884 | 0.0200296 |
| *hmit-1.3* | 805.63 | 284.71 | -1.38 | 0.49 | 0.005251 | 0.0422794 |
| *lap-1* | 4473.08 | 1575.32 | -1.38 | 0.50 | 0.00597 | 0.0464883 |
| *T01D1.4* | 1369.43 | 492.15 | -1.38 | 0.44 | 0.001784 | 0.0192551 |
| *F54A3.5* | 6259.12 | 2211.50 | -1.39 | 0.49 | 0.004491 | 0.0378933 |
| *R12C12.9* | 563.86 | 199.42 | -1.39 | 0.48 | 0.00424 | 0.0362738 |
| *M03F4.6* | 716.63 | 253.89 | -1.39 | 0.48 | 0.003734 | 0.0332202 |
| *C34C6.3* | 247.56 | 86.97 | -1.39 | 0.50 | 0.005288 | 0.0425081 |
| *col-90* | 2721.52 | 968.54 | -1.39 | 0.46 | 0.002711 | 0.0262394 |
| *R53.4* | 28706.50 | 10085.48 | -1.39 | 0.49 | 0.004642 | 0.0388298 |
| *C09F9.2* | 1031.68 | 367.88 | -1.39 | 0.45 | 0.002006 | 0.0211045 |
| *F31F4.1* | 373.40 | 131.66 | -1.39 | 0.48 | 0.003676 | 0.0328558 |
| *dpy-13* | 54790.61 | 19258.67 | -1.39 | 0.49 | 0.004138 | 0.0356867 |
| *PDB1.1* | 1244.00 | 439.77 | -1.39 | 0.47 | 0.002931 | 0.0277746 |
| *chch-3* | 4391.78 | 1556.48 | -1.40 | 0.46 | 0.002331 | 0.0236697 |
| *cox-6B* | 16193.39 | 5640.37 | -1.40 | 0.50 | 0.005488 | 0.0436567 |
| *D2023.4* | 769.08 | 270.33 | -1.40 | 0.47 | 0.003142 | 0.0291931 |
| *bus-19* | 1411.48 | 499.41 | -1.40 | 0.45 | 0.001826 | 0.019611 |
| *slc-17.1* | 454.27 | 157.40 | -1.40 | 0.50 | 0.005332 | 0.0426776 |
| *Y73B3A.18* | 1438.65 | 499.13 | -1.40 | 0.50 | 0.004815 | 0.0399261 |
| *cts-1* | 39889.11 | 13988.08 | -1.40 | 0.47 | 0.002752 | 0.026517 |
| *Y45G12B.3* | 1396.55 | 488.26 | -1.41 | 0.48 | 0.003178 | 0.0294406 |
| *D2023.1* | 645.33 | 224.25 | -1.41 | 0.49 | 0.004232 | 0.0362738 |
| *kdp-1* | 12733.50 | 4377.34 | -1.41 | 0.52 | 0.006442 | 0.0489832 |
| *tin-13* | 1225.87 | 423.68 | -1.41 | 0.50 | 0.004684 | 0.0391389 |
| *F01D5.6* | 692.70 | 238.73 | -1.41 | 0.50 | 0.004941 | 0.040657 |
| *clec-57* | 370.46 | 127.48 | -1.41 | 0.50 | 0.005104 | 0.0415312 |
| *ugt-64* | 783.87 | 272.78 | -1.41 | 0.48 | 0.002961 | 0.027968 |
| *cyp-42A1* | 415.75 | 144.80 | -1.41 | 0.47 | 0.002736 | 0.0264025 |
| *tomm-22* | 2723.13 | 944.80 | -1.41 | 0.48 | 0.003306 | 0.030381 |
| *D2092.4* | 2898.20 | 1016.95 | -1.41 | 0.45 | 0.001664 | 0.0182319 |
| *ugt-17* | 857.41 | 294.09 | -1.41 | 0.51 | 0.0052 | 0.0420213 |
| *K08D12.3* | 27598.80 | 9572.14 | -1.42 | 0.48 | 0.002974 | 0.028057 |
| *eef-1A.2* | 80166.32 | 27253.00 | -1.42 | 0.52 | 0.006115 | 0.0472509 |
| *rhy-1* | 337.41 | 114.59 | -1.42 | 0.52 | 0.006341 | 0.0483752 |
| *W03G11.4* | 1177.10 | 405.33 | -1.42 | 0.49 | 0.00357 | 0.0321094 |
| *F20G2.3* | 705.19 | 243.64 | -1.42 | 0.48 | 0.002762 | 0.0265807 |
| *srf-3* | 659.75 | 229.53 | -1.42 | 0.45 | 0.001656 | 0.0181785 |
| *T06A1.5* | 926.21 | 318.89 | -1.43 | 0.48 | 0.002813 | 0.026937 |
| *F19G12.10* | 587.19 | 201.79 | -1.43 | 0.48 | 0.002936 | 0.0277965 |
| *ard-1* | 6843.91 | 2327.83 | -1.43 | 0.50 | 0.004722 | 0.0393222 |
| *let-754* | 3102.97 | 1073.44 | -1.43 | 0.46 | 0.001979 | 0.0208449 |
| *C53B4.3* | 289.63 | 99.30 | -1.43 | 0.48 | 0.003196 | 0.0295706 |
| *arf-1.2* | 20235.15 | 7034.76 | -1.43 | 0.44 | 0.001237 | 0.0145979 |
| *sec-22* | 705.04 | 242.30 | -1.43 | 0.47 | 0.002563 | 0.0252728 |
| *col-76* | 2058.52 | 710.27 | -1.43 | 0.46 | 0.001843 | 0.0197477 |
| *gpx-1* | 363.78 | 122.45 | -1.43 | 0.52 | 0.00562 | 0.0444245 |
| *W04G3.5* | 2892.79 | 985.18 | -1.43 | 0.49 | 0.003468 | 0.0314559 |
| *zyx-1* | 20308.91 | 6988.49 | -1.44 | 0.46 | 0.001746 | 0.0189728 |
| *cyn-1* | 3230.59 | 1091.83 | -1.44 | 0.50 | 0.003853 | 0.0339702 |
| *tag-275* | 290.34 | 97.45 | -1.44 | 0.51 | 0.005041 | 0.0412165 |

| | | | | | | |
|---|---|---|---|---|---|---|
| *C26B9.3* | 1571.15 | 540.38 | -1.44 | 0.45 | 0.001337 | 0.0154705 |
| *cdr-6* | 2163.20 | 734.28 | -1.44 | 0.48 | 0.002818 | 0.026937 |
| *T20H4.5* | 4957.07 | 1679.93 | -1.44 | 0.49 | 0.003035 | 0.0284836 |
| *pitp-1* | 1568.86 | 538.10 | -1.44 | 0.45 | 0.001369 | 0.0157079 |
| *F20D1.1* | 2585.70 | 880.39 | -1.44 | 0.47 | 0.002156 | 0.022222 |
| *R07E4.3* | 864.79 | 293.38 | -1.44 | 0.48 | 0.00256 | 0.0252696 |
| *Y45G5AM.6* | 646.58 | 214.30 | -1.45 | 0.53 | 0.006241 | 0.0478737 |
| *act-3* | 44453.48 | 15239.27 | -1.45 | 0.45 | 0.00116 | 0.0139047 |
| *dpy-4* | 66152.47 | 21996.46 | -1.45 | 0.52 | 0.005471 | 0.043562 |
| *ifd-1* | 1333.08 | 441.08 | -1.45 | 0.53 | 0.006337 | 0.0483669 |
| *aass-1* | 1983.35 | 672.09 | -1.45 | 0.47 | 0.002093 | 0.0217703 |
| *sft-4* | 6392.72 | 2180.96 | -1.45 | 0.45 | 0.001322 | 0.0153594 |
| *gbb-2* | 259.27 | 85.75 | -1.45 | 0.53 | 0.005913 | 0.0461246 |
| *serp-1.1* | 33267.28 | 11104.67 | -1.45 | 0.50 | 0.004 | 0.0348542 |
| *romo-1* | 2759.45 | 923.78 | -1.45 | 0.50 | 0.003341 | 0.0305949 |
| *arr-1* | 877.80 | 296.99 | -1.45 | 0.47 | 0.001857 | 0.0198212 |
| *gei-13* | 1598.38 | 542.59 | -1.46 | 0.46 | 0.001402 | 0.0159808 |
| *asg-2* | 5571.51 | 1863.74 | -1.46 | 0.49 | 0.003138 | 0.0291721 |
| *cld-9* | 616.97 | 204.68 | -1.46 | 0.51 | 0.004208 | 0.0361865 |
| *Y34B4A.9* | 8821.76 | 2921.05 | -1.46 | 0.51 | 0.004502 | 0.0379727 |
| *T27A10.6* | 3210.55 | 1076.98 | -1.46 | 0.48 | 0.002482 | 0.0247026 |
| *cnnm-3* | 477.49 | 160.15 | -1.46 | 0.48 | 0.00244 | 0.0243732 |
| *casc-3* | 4545.33 | 1532.62 | -1.46 | 0.47 | 0.001774 | 0.0192042 |
| *dhs-20* | 339.13 | 112.65 | -1.46 | 0.50 | 0.003734 | 0.0332202 |
| *tmem-135* | 513.83 | 172.66 | -1.46 | 0.48 | 0.002133 | 0.0220171 |
| *ifc-2* | 18951.27 | 6295.91 | -1.46 | 0.50 | 0.003678 | 0.0328559 |
| *sptl-3* | 1904.47 | 642.45 | -1.46 | 0.46 | 0.001651 | 0.0181422 |
| *R151.2* | 6299.40 | 2147.52 | -1.46 | 0.43 | 0.000773 | 0.0101683 |
| *har-1* | 10789.20 | 3586.12 | -1.46 | 0.50 | 0.003434 | 0.0311672 |
| *mans-2* | 2163.84 | 719.49 | -1.46 | 0.50 | 0.003356 | 0.0306277 |
| *col-58* | 1008.97 | 329.34 | -1.46 | 0.54 | 0.006513 | 0.0493413 |
| *C26C6.9* | 821.85 | 276.38 | -1.46 | 0.47 | 0.001797 | 0.0193769 |
| *R102.2* | 443.44 | 145.47 | -1.46 | 0.53 | 0.005467 | 0.0435536 |
| *kel-8* | 700.62 | 234.36 | -1.46 | 0.48 | 0.002394 | 0.0241281 |
| *rpl-41.1* | 1555.33 | 514.50 | -1.46 | 0.50 | 0.003665 | 0.0328332 |
| *F15E6.6* | 711.73 | 236.71 | -1.46 | 0.49 | 0.002785 | 0.0267533 |
| *pdhb-1* | 8740.52 | 2946.80 | -1.47 | 0.45 | 0.001153 | 0.0138468 |
| *F56A8.3* | 6352.49 | 2155.68 | -1.47 | 0.43 | 0.00068 | 0.0091885 |
| *dim-1* | 33429.91 | 11122.15 | -1.47 | 0.48 | 0.002444 | 0.0243973 |
| *C07D8.6* | 8293.49 | 2690.29 | -1.47 | 0.54 | 0.006548 | 0.0494073 |
| *his-24* | 1079.40 | 364.55 | -1.47 | 0.44 | 0.000917 | 0.0116139 |
| *T11G6.4* | 358.48 | 117.70 | -1.47 | 0.51 | 0.004182 | 0.0360015 |
| *Y71H2AR.1* | 975.31 | 327.95 | -1.47 | 0.45 | 0.00115 | 0.0138314 |
| *ndx-2* | 383.19 | 125.69 | -1.47 | 0.51 | 0.003917 | 0.0343603 |
| *slc-36.2* | 902.23 | 302.49 | -1.47 | 0.46 | 0.001216 | 0.0143926 |
| *col-71* | 29532.33 | 9762.16 | -1.47 | 0.49 | 0.00264 | 0.0257303 |
| *F23H12.5* | 1191.25 | 387.73 | -1.47 | 0.52 | 0.004859 | 0.0401332 |
| *idhb-1* | 4484.30 | 1504.10 | -1.48 | 0.45 | 0.000934 | 0.011736 |
| *F22H10.3* | 4876.72 | 1578.60 | -1.48 | 0.53 | 0.005398 | 0.043118 |
| *Y87G2A.2* | 335.31 | 107.72 | -1.48 | 0.54 | 0.006623 | 0.0498776 |
| *nuo-6* | 3988.44 | 1307.45 | -1.48 | 0.50 | 0.003112 | 0.0289983 |
| *W05B5.1* | 8628.71 | 2785.95 | -1.48 | 0.53 | 0.005186 | 0.0419716 |
| *H18N23.2* | 6574.94 | 2174.18 | -1.49 | 0.47 | 0.001449 | 0.0164126 |
| *mev-1* | 10356.16 | 3372.39 | -1.49 | 0.50 | 0.002959 | 0.027968 |
| *ZC250.4* | 1948.36 | 641.54 | -1.49 | 0.47 | 0.001571 | 0.0175413 |
| *F13B6.3* | 2818.04 | 915.78 | -1.49 | 0.50 | 0.003046 | 0.0285259 |

| | | | | | | |
|---|---|---|---|---|---|---|
| *C05G5.1* | 331.32 | 107.52 | -1.49 | 0.50 | 0.003181 | 0.0294482 |
| *vha-2* | 59386.17 | 19442.57 | -1.49 | 0.48 | 0.002068 | 0.0215721 |
| *ugt-49* | 688.94 | 225.76 | -1.49 | 0.48 | 0.001942 | 0.0205321 |
| *set-20* | 190.69 | 61.29 | -1.49 | 0.52 | 0.0044 | 0.0373144 |
| *zig-12* | 12049.74 | 3938.91 | -1.49 | 0.49 | 0.002118 | 0.0219048 |
| *cpr-5* | 23856.19 | 7698.54 | -1.49 | 0.51 | 0.003733 | 0.0332202 |
| *slcr-46.3* | 514.81 | 169.78 | -1.49 | 0.46 | 0.001254 | 0.014753 |
| *iff-2* | 53254.03 | 17334.08 | -1.49 | 0.49 | 0.002571 | 0.0253063 |
| *T10H10.2* | 672.48 | 221.24 | -1.49 | 0.47 | 0.00143 | 0.0162457 |
| *F42A10.9* | 779.62 | 251.83 | -1.49 | 0.51 | 0.003433 | 0.0311672 |
| *Y63D3A.7* | 913.09 | 293.95 | -1.49 | 0.51 | 0.003693 | 0.0329303 |
| *daf-22* | 5244.37 | 1659.96 | -1.49 | 0.55 | 0.00646 | 0.0489925 |
| *aco-1* | 9294.71 | 3047.69 | -1.49 | 0.47 | 0.001498 | 0.0168423 |
| *dpy-11* | 8821.94 | 2921.41 | -1.49 | 0.44 | 0.00075 | 0.0099278 |
| *F11C1.1* | 308.02 | 99.96 | -1.49 | 0.49 | 0.00251 | 0.0249362 |
| *T04B8.5* | 1931.43 | 638.34 | -1.50 | 0.45 | 0.000841 | 0.0109035 |
| *subs-4* | 257.85 | 83.05 | -1.50 | 0.51 | 0.003354 | 0.0306277 |
| *F23A7.4* | 1365.90 | 439.87 | -1.50 | 0.51 | 0.003338 | 0.030589 |
| *clec-160* | 304.62 | 97.41 | -1.50 | 0.52 | 0.003883 | 0.0341795 |
| *Y105C5B.9* | 617.28 | 199.92 | -1.50 | 0.49 | 0.002255 | 0.0230832 |
| *H10E21.4* | 1714.11 | 557.24 | -1.50 | 0.48 | 0.001755 | 0.0190589 |
| *C06G1.1* | 1643.22 | 517.66 | -1.50 | 0.55 | 0.006034 | 0.0468034 |
| *Y45G12C.1* | 209.15 | 66.57 | -1.50 | 0.53 | 0.00423 | 0.0362738 |
| *T02H6.11* | 16785.57 | 5384.08 | -1.50 | 0.51 | 0.003049 | 0.0285337 |
| *F58E6.13* | 1098.89 | 355.27 | -1.50 | 0.49 | 0.002098 | 0.0217861 |
| *dhs-28* | 10963.17 | 3485.35 | -1.50 | 0.53 | 0.004229 | 0.0362738 |
| *scav-6* | 647.95 | 211.57 | -1.50 | 0.46 | 0.001163 | 0.0139357 |
| *mlt-9* | 3494.48 | 1104.54 | -1.51 | 0.53 | 0.004835 | 0.0400375 |
| *hil-3* | 1739.04 | 569.48 | -1.51 | 0.45 | 0.00074 | 0.0098195 |
| *C56C10.4* | 319.44 | 99.85 | -1.51 | 0.55 | 0.006166 | 0.0474409 |
| *hsp-3* | 47840.08 | 15438.38 | -1.51 | 0.48 | 0.001624 | 0.0179524 |
| *Y25C1A.13* | 1380.79 | 438.18 | -1.51 | 0.51 | 0.003211 | 0.0296742 |
| *prx-3* | 753.60 | 239.14 | -1.51 | 0.51 | 0.003025 | 0.0284382 |
| *cth-2* | 11572.99 | 3724.65 | -1.52 | 0.48 | 0.00153 | 0.0171391 |
| *immt-1* | 3805.78 | 1232.15 | -1.52 | 0.46 | 0.001029 | 0.0126564 |
| *R07B1.11* | 177.01 | 55.23 | -1.52 | 0.54 | 0.005095 | 0.0414783 |
| *nhr-31* | 2656.09 | 824.76 | -1.52 | 0.55 | 0.005908 | 0.0461245 |
| *gst-13* | 1381.46 | 438.85 | -1.52 | 0.50 | 0.002594 | 0.0254188 |
| *gst-42* | 3256.49 | 1047.63 | -1.52 | 0.47 | 0.00132 | 0.0153568 |
| *Y106G6H.14* | 580.51 | 185.57 | -1.52 | 0.48 | 0.001678 | 0.0183622 |
| *ugt-50* | 1816.01 | 585.75 | -1.52 | 0.46 | 0.000871 | 0.0111868 |
| *Y53F4B.25* | 462.51 | 146.29 | -1.52 | 0.50 | 0.00241 | 0.0242014 |
| *H11E01.3* | 1265.79 | 398.89 | -1.53 | 0.51 | 0.002654 | 0.0258387 |
| *erv-46* | 869.69 | 280.07 | -1.53 | 0.45 | 0.000756 | 0.009997 |
| *K02E10.4* | 611.29 | 192.86 | -1.53 | 0.50 | 0.002339 | 0.0237441 |
| *W10C8.5* | 2431.81 | 773.21 | -1.53 | 0.48 | 0.001509 | 0.0169541 |
| *C29F7.3* | 1629.27 | 506.35 | -1.53 | 0.53 | 0.003839 | 0.0338928 |
| *cpl-1* | 38072.14 | 12055.04 | -1.53 | 0.49 | 0.00176 | 0.0190761 |
| *col-107* | 38070.46 | 11915.13 | -1.53 | 0.51 | 0.002875 | 0.0273483 |
| *tfg-1* | 12318.01 | 3953.38 | -1.53 | 0.45 | 0.000725 | 0.0096739 |
| *Y32G9B.1* | 1110.82 | 351.33 | -1.53 | 0.49 | 0.001622 | 0.0179367 |
| *nrf-6* | 2687.41 | 857.98 | -1.53 | 0.46 | 0.000935 | 0.0117382 |
| *F07A11.4* | 1629.57 | 519.77 | -1.53 | 0.47 | 0.000984 | 0.0122252 |
| *tag-96* | 260.50 | 80.45 | -1.53 | 0.54 | 0.004223 | 0.0362738 |
| *wah-1* | 4507.76 | 1449.67 | -1.54 | 0.44 | 0.000489 | 0.007059 |
| *asb-2* | 26186.21 | 8243.33 | -1.54 | 0.49 | 0.001854 | 0.0198212 |

| | | | | | | |
|---|---|---|---|---|---|---|
| *ins-27* | 2403.90 | 745.91 | -1.54 | 0.52 | 0.003311 | 0.0304111 |
| *cav-2* | 595.15 | 185.84 | -1.54 | 0.51 | 0.00257 | 0.0253059 |
| *fip-5* | 464.57 | 145.23 | -1.54 | 0.50 | 0.002309 | 0.0234966 |
| *atp-1* | 134845.99 | 42744.42 | -1.54 | 0.47 | 0.001141 | 0.0137427 |
| *F29B9.11* | 13063.96 | 4074.00 | -1.54 | 0.51 | 0.002459 | 0.0245209 |
| *C44F1.1* | 378.06 | 117.11 | -1.54 | 0.52 | 0.003131 | 0.0291252 |
| *aldo-1* | 12009.30 | 3751.28 | -1.54 | 0.50 | 0.002106 | 0.0218435 |
| *bre-1* | 998.68 | 314.51 | -1.54 | 0.48 | 0.001333 | 0.015436 |
| *Y73E7A.3* | 338.97 | 105.22 | -1.54 | 0.51 | 0.002524 | 0.0249896 |
| *Y47D7A.13* | 18273.76 | 5610.39 | -1.55 | 0.53 | 0.00343 | 0.0311625 |
| *C17F4.7* | 71113.63 | 22433.72 | -1.55 | 0.47 | 0.000905 | 0.0115066 |
| *B0250.5* | 3827.84 | 1193.83 | -1.55 | 0.49 | 0.001634 | 0.0179978 |
| *col-91* | 7000.80 | 2130.84 | -1.55 | 0.54 | 0.004241 | 0.0362738 |
| *haf-4* | 3677.50 | 1159.94 | -1.55 | 0.46 | 0.000737 | 0.0097849 |
| *unc-95* | 1884.50 | 581.00 | -1.55 | 0.51 | 0.002407 | 0.0241921 |
| *LLC1.2* | 4705.56 | 1423.72 | -1.55 | 0.55 | 0.004591 | 0.0385487 |
| *Y34B4A.10* | 1306.47 | 414.33 | -1.55 | 0.44 | 0.000416 | 0.0062747 |
| *Y53F4B.39* | 1695.49 | 529.41 | -1.56 | 0.48 | 0.001138 | 0.0137212 |
| *gas-1* | 6994.39 | 2162.93 | -1.56 | 0.50 | 0.001782 | 0.0192551 |
| *hpo-18* | 3826.71 | 1185.78 | -1.56 | 0.49 | 0.001596 | 0.0177402 |
| *mocs-1* | 251.24 | 77.25 | -1.56 | 0.51 | 0.002242 | 0.0229818 |
| *Y49E10.18* | 2054.05 | 628.53 | -1.56 | 0.52 | 0.002729 | 0.0263764 |
| *B0261.8* | 490.08 | 147.33 | -1.56 | 0.55 | 0.004743 | 0.0394545 |
| *shw-3* | 147.48 | 44.59 | -1.56 | 0.54 | 0.00391 | 0.034325 |
| *ptr-12* | 656.43 | 197.86 | -1.56 | 0.55 | 0.004235 | 0.0362738 |
| *F35E12.6* | 8148.72 | 2489.28 | -1.56 | 0.52 | 0.002569 | 0.0253059 |
| *col-60* | 8105.28 | 2481.83 | -1.56 | 0.51 | 0.002193 | 0.0225175 |
| *F35F10.5* | 257.86 | 79.19 | -1.56 | 0.50 | 0.001837 | 0.0196899 |
| *pdi-6* | 11740.11 | 3650.27 | -1.56 | 0.47 | 0.000913 | 0.0115758 |
| *F30F8.5* | 445.75 | 134.96 | -1.56 | 0.53 | 0.003082 | 0.028788 |
| *Y66H1A.5* | 1322.70 | 399.41 | -1.57 | 0.53 | 0.003291 | 0.0303225 |
| *tpi-1* | 16771.55 | 5208.19 | -1.57 | 0.47 | 0.000924 | 0.0116712 |
| *vha-14* | 12988.37 | 4038.50 | -1.57 | 0.47 | 0.000844 | 0.0109167 |
| *aqp-10* | 2484.16 | 760.30 | -1.57 | 0.49 | 0.001484 | 0.0167387 |
| *cutl-15* | 706.76 | 216.78 | -1.58 | 0.49 | 0.001168 | 0.0139793 |
| *R03E9.2* | 268.55 | 81.65 | -1.58 | 0.50 | 0.001611 | 0.0178492 |
| *zip-3* | 3466.75 | 1061.44 | -1.58 | 0.48 | 0.001083 | 0.013243 |
| *sqt-1* | 16333.38 | 4990.86 | -1.58 | 0.49 | 0.001172 | 0.014007 |
| *vdac-1* | 49061.08 | 14883.25 | -1.58 | 0.50 | 0.001605 | 0.0177894 |
| *hyl-2* | 3155.99 | 978.51 | -1.58 | 0.44 | 0.000375 | 0.0057708 |
| *F21D5.3* | 557.85 | 168.75 | -1.58 | 0.50 | 0.001656 | 0.0181785 |
| *bckd-1B* | 3768.90 | 1164.31 | -1.58 | 0.45 | 0.000452 | 0.0066485 |
| *C40H1.2* | 355.44 | 108.61 | -1.58 | 0.48 | 0.000901 | 0.0114867 |
| *C18E9.4* | 2929.04 | 890.39 | -1.59 | 0.48 | 0.001064 | 0.013042 |
| *E04F6.15* | 141.46 | 41.61 | -1.59 | 0.55 | 0.003908 | 0.0343215 |
| *slcr-46.1* | 449.55 | 135.31 | -1.59 | 0.51 | 0.001663 | 0.0182319 |
| *col-125* | 61934.25 | 18401.92 | -1.59 | 0.53 | 0.002787 | 0.0267554 |
| *W09D6.5* | 7886.13 | 2385.87 | -1.59 | 0.49 | 0.001298 | 0.0151598 |
| *col-182* | 1109.19 | 329.66 | -1.59 | 0.53 | 0.002648 | 0.0258001 |
| *Y66A7A.7* | 650.31 | 192.97 | -1.59 | 0.53 | 0.002547 | 0.0251861 |
| *K12H4.5* | 5587.58 | 1692.72 | -1.59 | 0.48 | 0.000966 | 0.0120585 |
| *col-144* | 71989.19 | 21400.59 | -1.60 | 0.52 | 0.002047 | 0.0214244 |
| *clec-49* | 1582.55 | 467.87 | -1.60 | 0.53 | 0.002435 | 0.0243403 |
| *R17.3* | 577.93 | 172.86 | -1.60 | 0.50 | 0.001466 | 0.0165734 |
| *lipl-6* | 290.56 | 87.06 | -1.60 | 0.50 | 0.001346 | 0.0155259 |
| *nep-17* | 37369.72 | 11294.07 | -1.60 | 0.48 | 0.000843 | 0.0109167 |

| | | | | | | |
|---|---|---|---|---|---|---|
| *let-653* | 809.58 | 245.14 | -1.60 | 0.47 | 0.00075 | 0.0099278 |
| *pgp-1* | 845.58 | 257.63 | -1.60 | 0.46 | 0.000459 | 0.006708 |
| *hum-4* | 2693.49 | 827.71 | -1.60 | 0.43 | 0.000225 | 0.0038937 |
| *unc-7* | 474.85 | 141.99 | -1.60 | 0.50 | 0.001319 | 0.0153559 |
| *col-65* | 4911.97 | 1466.73 | -1.60 | 0.50 | 0.001358 | 0.0156327 |
| *C54D2.2* | 318.52 | 93.68 | -1.60 | 0.53 | 0.002453 | 0.0244708 |
| *phy-2* | 1288.63 | 386.99 | -1.61 | 0.48 | 0.000812 | 0.0106125 |
| *K10B3.1* | 1290.90 | 385.76 | -1.61 | 0.49 | 0.000922 | 0.01166 |
| *C14H10.3* | 1855.86 | 565.82 | -1.61 | 0.44 | 0.000218 | 0.0037939 |
| *col-117* | 1550.50 | 457.57 | -1.61 | 0.51 | 0.00158 | 0.0176336 |
| *F44B9.2* | 1632.70 | 489.05 | -1.61 | 0.48 | 0.000718 | 0.0096115 |
| *pgph-2* | 3912.41 | 1133.53 | -1.61 | 0.54 | 0.003045 | 0.0285259 |
| *Y54F10AM.8* | 8297.59 | 2496.10 | -1.61 | 0.46 | 0.000523 | 0.0073726 |
| *mrps-33* | 1186.03 | 353.02 | -1.61 | 0.49 | 0.000933 | 0.011736 |
| *vrp-1* | 974.08 | 293.37 | -1.61 | 0.46 | 0.000449 | 0.0066242 |
| *C18H9.6* | 297.16 | 86.99 | -1.61 | 0.52 | 0.001916 | 0.0203141 |
| *clec-173* | 1556.50 | 446.58 | -1.62 | 0.56 | 0.003674 | 0.0328558 |
| *hpo-36* | 475.74 | 142.56 | -1.62 | 0.47 | 0.000558 | 0.0077712 |
| *nlp-24* | 4484.06 | 1312.53 | -1.62 | 0.51 | 0.001629 | 0.0179578 |
| *cest-1.1* | 213.08 | 61.00 | -1.62 | 0.55 | 0.003313 | 0.0304111 |
| *ZK809.8* | 2329.27 | 688.38 | -1.62 | 0.49 | 0.000927 | 0.011681 |
| *ptr-24* | 2153.88 | 648.39 | -1.62 | 0.44 | 0.000243 | 0.0041351 |
| *T19D12.1* | 5486.43 | 1644.20 | -1.62 | 0.45 | 0.000345 | 0.0054367 |
| *F36H9.5* | 616.08 | 183.60 | -1.62 | 0.47 | 0.000482 | 0.006975 |
| *C01H6.8* | 163.54 | 47.30 | -1.63 | 0.53 | 0.002109 | 0.0218435 |
| *clec-50* | 13237.35 | 3861.99 | -1.63 | 0.51 | 0.001389 | 0.0158845 |
| *lin-10* | 1939.35 | 573.15 | -1.63 | 0.48 | 0.000705 | 0.0094712 |
| *ivd-1* | 6525.41 | 1933.84 | -1.63 | 0.47 | 0.000586 | 0.0081252 |
| *nhr-76* | 690.01 | 204.76 | -1.63 | 0.47 | 0.000516 | 0.0072943 |
| *col-33* | 163.12 | 45.25 | -1.63 | 0.59 | 0.005972 | 0.0464883 |
| *F52G2.3* | 562.57 | 167.24 | -1.63 | 0.46 | 0.000445 | 0.0065802 |
| *sem-2* | 1816.76 | 538.32 | -1.63 | 0.47 | 0.000513 | 0.0072704 |
| *hog-1* | 187.30 | 54.06 | -1.63 | 0.52 | 0.00185 | 0.0197894 |
| *acds-10* | 1598.52 | 454.79 | -1.63 | 0.55 | 0.002854 | 0.0272104 |
| *ugt-29* | 570.81 | 167.48 | -1.63 | 0.49 | 0.000776 | 0.0101966 |
| *cysl-1* | 1498.74 | 434.67 | -1.63 | 0.51 | 0.001347 | 0.015532 |
| *vha-1* | 13817.57 | 4037.74 | -1.64 | 0.49 | 0.000888 | 0.0113408 |
| *cpr-4* | 7888.32 | 2315.73 | -1.64 | 0.48 | 0.000688 | 0.0092663 |
| *lips-9* | 1141.37 | 337.32 | -1.64 | 0.46 | 0.000426 | 0.0063619 |
| *F39E9.1* | 355.33 | 100.47 | -1.64 | 0.55 | 0.002992 | 0.0281938 |
| *F46C8.8* | 3017.68 | 847.23 | -1.64 | 0.56 | 0.003556 | 0.0320456 |
| *cest-2.1* | 266.45 | 77.20 | -1.64 | 0.50 | 0.001062 | 0.0130301 |
| *Y22D7AL.10* | 10448.40 | 3063.91 | -1.64 | 0.47 | 0.000478 | 0.0069393 |
| *Y53G8B.1* | 781.82 | 220.13 | -1.64 | 0.55 | 0.002793 | 0.0268013 |
| *C49F8.3* | 2037.01 | 603.44 | -1.65 | 0.44 | 0.0002 | 0.0035466 |
| *col-97* | 28744.27 | 8475.04 | -1.65 | 0.45 | 0.00025 | 0.0042378 |
| *lbp-3* | 3224.15 | 926.13 | -1.65 | 0.51 | 0.001175 | 0.0140199 |
| *unc-52* | 17273.23 | 5072.22 | -1.65 | 0.46 | 0.000327 | 0.0052186 |
| *H03E18.1* | 1903.46 | 531.45 | -1.65 | 0.56 | 0.0031 | 0.0289263 |
| *B0393.5* | 645.57 | 185.27 | -1.65 | 0.50 | 0.000965 | 0.0120497 |
| *ZK180.5* | 40061.56 | 11673.78 | -1.65 | 0.47 | 0.000405 | 0.0061618 |
| *Y73F4A.1* | 275.23 | 79.37 | -1.65 | 0.49 | 0.000737 | 0.0097849 |
| *C27A2.12* | 142.92 | 40.07 | -1.66 | 0.54 | 0.002126 | 0.0219566 |
| *F58B4.5* | 963.25 | 269.01 | -1.66 | 0.55 | 0.002383 | 0.0240931 |
| *txdc-12.2* | 3449.43 | 983.11 | -1.66 | 0.51 | 0.00109 | 0.0133012 |
| *tatn-1* | 12390.55 | 3487.55 | -1.66 | 0.53 | 0.001783 | 0.0192551 |

| | | | | | | |
|---|---|---|---|---|---|---|
| *C01F1.3* | 544.29 | 153.56 | -1.66 | 0.53 | 0.001628 | 0.0179524 |
| *K09C4.5* | 315.29 | 90.82 | -1.66 | 0.48 | 0.000595 | 0.008202 |
| *cuti-1* | 400.63 | 114.44 | -1.66 | 0.50 | 0.000903 | 0.0114958 |
| *T20D3.2* | 15171.77 | 4316.88 | -1.66 | 0.51 | 0.001084 | 0.013243 |
| *T04F3.4* | 316.00 | 90.14 | -1.66 | 0.50 | 0.000934 | 0.011736 |
| *apy-1* | 2243.33 | 651.12 | -1.66 | 0.46 | 0.000337 | 0.0053415 |
| *D1086.3* | 1306.68 | 369.36 | -1.67 | 0.51 | 0.001191 | 0.0141696 |
| *dbt-1* | 5599.76 | 1638.61 | -1.67 | 0.44 | 0.000133 | 0.0025357 |
| *fpn-1.1* | 747.32 | 214.24 | -1.67 | 0.48 | 0.000557 | 0.0077702 |
| *F35E12.10* | 1740.89 | 498.62 | -1.67 | 0.48 | 0.000517 | 0.0073038 |
| *nlp-20* | 3641.32 | 1006.54 | -1.67 | 0.55 | 0.002365 | 0.0239577 |
| *paqr-3* | 325.13 | 93.04 | -1.67 | 0.48 | 0.000534 | 0.0075104 |
| *tnt-4* | 319.02 | 84.71 | -1.67 | 0.61 | 0.006146 | 0.0473362 |
| *col-3* | 1958.37 | 554.39 | -1.67 | 0.50 | 0.000926 | 0.011681 |
| *dao-3* | 2578.05 | 736.09 | -1.67 | 0.49 | 0.000587 | 0.0081268 |
| *vha-15* | 17486.47 | 5026.05 | -1.67 | 0.47 | 0.000392 | 0.0059835 |
| *B0035.13* | 3257.58 | 892.90 | -1.67 | 0.56 | 0.00291 | 0.0276075 |
| *clec-51* | 685.08 | 194.63 | -1.67 | 0.49 | 0.000702 | 0.0094327 |
| *T02D1.8* | 1234.05 | 348.94 | -1.67 | 0.50 | 0.000824 | 0.0107387 |
| *F09B12.3* | 9445.48 | 2720.81 | -1.67 | 0.46 | 0.00027 | 0.0044904 |
| *F41F3.3* | 73758.83 | 20791.47 | -1.67 | 0.51 | 0.000921 | 0.011653 |
| *C32H11.4* | 346.24 | 96.47 | -1.68 | 0.52 | 0.0014 | 0.0159808 |
| *lys-4* | 10112.87 | 2797.58 | -1.68 | 0.54 | 0.001761 | 0.0190769 |
| *elo-1* | 20229.70 | 5694.75 | -1.68 | 0.50 | 0.000861 | 0.0110779 |
| *skpo-2* | 1184.90 | 338.42 | -1.68 | 0.47 | 0.00037 | 0.0057378 |
| *exc-15* | 13934.41 | 3798.80 | -1.68 | 0.56 | 0.002481 | 0.0247026 |
| *T05E7.1* | 623.67 | 171.04 | -1.69 | 0.54 | 0.001858 | 0.0198214 |
| *tomm-7* | 275.20 | 74.93 | -1.69 | 0.55 | 0.002295 | 0.023398 |
| *T05C3.6* | 875.19 | 249.32 | -1.69 | 0.46 | 0.00025 | 0.0042354 |
| *fbn-1* | 1646.12 | 432.82 | -1.69 | 0.60 | 0.005076 | 0.0413673 |
| *ptc-3* | 4078.06 | 1168.22 | -1.69 | 0.45 | 0.000146 | 0.002747 |
| *col-128* | 186.69 | 50.64 | -1.69 | 0.55 | 0.002279 | 0.0232747 |
| *T13C2.3* | 521.43 | 141.90 | -1.69 | 0.55 | 0.002038 | 0.0213675 |
| *ckb-4* | 446.23 | 123.93 | -1.69 | 0.51 | 0.000887 | 0.0113408 |
| *gln-3* | 12579.24 | 3467.20 | -1.69 | 0.52 | 0.001222 | 0.0144473 |
| *gpdh-1* | 1142.09 | 316.03 | -1.69 | 0.52 | 0.001026 | 0.0126367 |
| *F33A8.7* | 1199.40 | 339.21 | -1.69 | 0.47 | 0.000307 | 0.0049679 |
| *gcst-1* | 5346.54 | 1509.01 | -1.69 | 0.47 | 0.000329 | 0.0052372 |
| *tsn-1* | 30098.52 | 8608.97 | -1.70 | 0.44 | 0.000107 | 0.0021376 |
| *C18E9.5* | 1403.51 | 395.88 | -1.70 | 0.47 | 0.000293 | 0.0047865 |
| *pgrn-1* | 2005.43 | 554.49 | -1.70 | 0.51 | 0.00087 | 0.0111777 |
| *K09E2.3* | 3165.33 | 861.24 | -1.70 | 0.54 | 0.001641 | 0.018052 |
| *mpst-3* | 1124.51 | 308.98 | -1.70 | 0.52 | 0.00102 | 0.0125902 |
| *R07B7.10* | 237.93 | 65.31 | -1.70 | 0.52 | 0.000947 | 0.0118642 |
| *col-7* | 2606.85 | 690.24 | -1.70 | 0.58 | 0.003099 | 0.0289263 |
| *acox-1.1* | 2799.61 | 730.09 | -1.70 | 0.60 | 0.004367 | 0.0371342 |
| *F22B8.7* | 759.27 | 211.48 | -1.71 | 0.48 | 0.000373 | 0.0057619 |
| *T19B4.3* | 1820.55 | 507.91 | -1.71 | 0.48 | 0.000333 | 0.0052914 |
| *Y47G6A.33* | 397.90 | 110.70 | -1.71 | 0.48 | 0.000367 | 0.0057094 |
| *Y39A1A.21* | 574.39 | 159.31 | -1.71 | 0.48 | 0.0004 | 0.0060877 |
| *ldh-1* | 2354.34 | 641.09 | -1.71 | 0.52 | 0.000977 | 0.0121591 |
| *osm-8* | 996.11 | 260.10 | -1.71 | 0.58 | 0.003382 | 0.0307893 |
| *agxt-1* | 1342.61 | 372.07 | -1.71 | 0.48 | 0.000362 | 0.0056394 |
| *Y73C8B.3* | 902.53 | 243.87 | -1.71 | 0.53 | 0.001184 | 0.0141235 |
| *ZK652.8* | 498.23 | 133.91 | -1.71 | 0.54 | 0.001419 | 0.0161389 |
| *cept-1* | 1235.47 | 330.73 | -1.72 | 0.54 | 0.001555 | 0.0173932 |

| | | | | | | |
|---|---|---|---|---|---|---|
| *cdh-10* | 1122.43 | 306.82 | -1.72 | 0.50 | 0.00066 | 0.0089437 |
| *rol-6* | 6561.81 | 1850.14 | -1.72 | 0.43 | 6.93E-05 | 0.0014999 |
| *D2005.6* | 121.59 | 31.87 | -1.72 | 0.57 | 0.002661 | 0.0258805 |
| *mxt-1* | 1019.82 | 283.52 | -1.72 | 0.46 | 0.000192 | 0.0034278 |
| *ddp-1* | 3646.99 | 996.53 | -1.72 | 0.50 | 0.000535 | 0.0075127 |
| *T03G6.1* | 667.51 | 179.01 | -1.72 | 0.53 | 0.001206 | 0.014306 |
| *F25B5.3* | 12424.97 | 3399.82 | -1.72 | 0.49 | 0.000492 | 0.0070949 |
| *ZC434.3* | 487.52 | 129.35 | -1.72 | 0.55 | 0.001625 | 0.0179524 |
| *rhr-1* | 5286.92 | 1385.93 | -1.72 | 0.57 | 0.002314 | 0.0235281 |
| *fut-8* | 807.07 | 223.98 | -1.73 | 0.45 | 0.00013 | 0.0024903 |
| *cyp-35C1* | 509.82 | 137.13 | -1.73 | 0.52 | 0.000811 | 0.0106057 |
| *fah-1* | 8979.96 | 2411.19 | -1.73 | 0.52 | 0.000815 | 0.0106366 |
| *sqt-3* | 40967.60 | 11136.61 | -1.73 | 0.49 | 0.000444 | 0.006571 |
| *acs-5* | 4316.73 | 1184.14 | -1.74 | 0.47 | 0.000196 | 0.0034828 |
| *T27E9.2* | 2740.80 | 739.80 | -1.74 | 0.50 | 0.00051 | 0.0072401 |
| *fard-1* | 2423.69 | 656.38 | -1.74 | 0.49 | 0.000374 | 0.0057639 |
| *Y39E4B.6* | 1963.99 | 529.97 | -1.74 | 0.49 | 0.00043 | 0.0064116 |
| *mth-1* | 521.90 | 139.99 | -1.74 | 0.51 | 0.000584 | 0.0080986 |
| *clec-238* | 74.31 | 18.35 | -1.74 | 0.63 | 0.00559 | 0.0442915 |
| *mec-17* | 622.45 | 162.93 | -1.74 | 0.54 | 0.001375 | 0.0157644 |
| *ncx-6* | 177.68 | 46.64 | -1.74 | 0.54 | 0.001207 | 0.014306 |
| *K12C11.1* | 4342.65 | 1161.32 | -1.75 | 0.50 | 0.000463 | 0.0067598 |
| *C15C7.5* | 9818.63 | 2652.50 | -1.75 | 0.48 | 0.000256 | 0.0043071 |
| *fbp-1* | 4826.65 | 1298.31 | -1.75 | 0.49 | 0.000312 | 0.0050303 |
| *vha-4* | 13595.91 | 3675.51 | -1.75 | 0.47 | 0.000222 | 0.0038535 |
| *math-4* | 1407.31 | 364.18 | -1.75 | 0.55 | 0.001592 | 0.0177134 |
| *pes-9* | 7105.95 | 1911.29 | -1.75 | 0.48 | 0.000296 | 0.0048265 |
| *sdha-1* | 8341.61 | 2246.88 | -1.75 | 0.48 | 0.000257 | 0.0043229 |
| *C07A9.9* | 362.58 | 97.53 | -1.75 | 0.48 | 0.000251 | 0.0042528 |
| *lbp-7* | 699.30 | 184.50 | -1.76 | 0.51 | 0.000647 | 0.0087929 |
| *T25B9.9* | 7520.75 | 2036.63 | -1.76 | 0.46 | 0.000138 | 0.0025999 |
| *col-38* | 33627.98 | 8941.20 | -1.76 | 0.50 | 0.000419 | 0.0063036 |
| *slc-25A18.1* | 1857.78 | 492.25 | -1.76 | 0.50 | 0.000418 | 0.0062929 |
| *mai-2* | 10901.35 | 2888.22 | -1.76 | 0.50 | 0.000413 | 0.006265 |
| *strm-1* | 75.45 | 18.10 | -1.76 | 0.64 | 0.005988 | 0.0465616 |
| *C52G5.2* | 2235.26 | 599.98 | -1.76 | 0.47 | 0.000171 | 0.0031328 |
| *C05D12.1* | 356.13 | 94.70 | -1.76 | 0.49 | 0.000297 | 0.0048283 |
| *col-159* | 61957.08 | 15952.78 | -1.77 | 0.54 | 0.001146 | 0.0137903 |
| *cox-6C* | 6483.77 | 1691.68 | -1.77 | 0.52 | 0.000682 | 0.009204 |
| *dpy-20* | 219.21 | 55.84 | -1.77 | 0.56 | 0.00153 | 0.0171391 |
| *clec-169* | 112.59 | 27.83 | -1.77 | 0.60 | 0.003054 | 0.0285612 |
| *H42K12.3* | 5110.41 | 1308.27 | -1.77 | 0.55 | 0.00117 | 0.0139976 |
| *Y51A2D.18* | 378.69 | 95.73 | -1.77 | 0.56 | 0.001688 | 0.018429 |
| *col-69* | 65.64 | 15.71 | -1.77 | 0.63 | 0.0052 | 0.0420213 |
| *ugt-7* | 79.46 | 19.40 | -1.77 | 0.61 | 0.003628 | 0.0325759 |
| *F58H1.2* | 238.77 | 59.67 | -1.77 | 0.58 | 0.002178 | 0.0223785 |
| *T10E9.3* | 1666.11 | 447.38 | -1.78 | 0.45 | 6.56E-05 | 0.0014394 |
| *nlp-30* | 529.92 | 139.76 | -1.78 | 0.48 | 0.000227 | 0.003905 |
| *anmt-2* | 610.02 | 160.26 | -1.78 | 0.49 | 0.000277 | 0.0045792 |
| *gst-5* | 1198.76 | 314.98 | -1.78 | 0.48 | 0.000229 | 0.0039222 |
| *ptr-11* | 2100.76 | 561.08 | -1.78 | 0.45 | 7.03E-05 | 0.0015138 |
| *ZK185.3* | 301.36 | 71.84 | -1.78 | 0.63 | 0.004573 | 0.0384363 |
| *swt-5* | 356.37 | 92.54 | -1.78 | 0.50 | 0.000388 | 0.0059345 |
| *vwa-8* | 3183.88 | 853.31 | -1.79 | 0.43 | 3.82E-05 | 0.0009523 |
| *W02F12.2* | 174.54 | 44.22 | -1.79 | 0.54 | 0.001019 | 0.012579 |
| *C01G10.9* | 729.58 | 189.58 | -1.79 | 0.50 | 0.000346 | 0.0054555 |

| | | | | | | |
|---|---|---|---|---|---|---|
| *F56F10.1* | 3393.11 | 891.95 | -1.79 | 0.48 | 0.000177 | 0.0032232 |
| *bus-4* | 920.69 | 242.50 | -1.79 | 0.47 | 0.000154 | 0.0028585 |
| *clik-1* | 81904.50 | 21190.71 | -1.79 | 0.51 | 0.000416 | 0.0062782 |
| *T24D5.2* | 536.09 | 138.03 | -1.79 | 0.51 | 0.000498 | 0.0071416 |
| *Y18D10A.23* | 997.99 | 246.36 | -1.79 | 0.58 | 0.002069 | 0.0215721 |
| *R07E3.6* | 1924.26 | 480.67 | -1.79 | 0.56 | 0.00148 | 0.016709 |
| *col-175* | 11598.23 | 3082.93 | -1.79 | 0.45 | 6.08E-05 | 0.0013499 |
| *R09H10.5* | 6286.09 | 1680.92 | -1.79 | 0.43 | 3.35E-05 | 0.0008496 |
| *acbp-1* | 13128.58 | 3339.66 | -1.79 | 0.53 | 0.000732 | 0.0097381 |
| *Y43F4A.1* | 654.19 | 170.42 | -1.79 | 0.48 | 0.000214 | 0.00373 |
| *aat-6* | 690.17 | 171.01 | -1.79 | 0.57 | 0.001616 | 0.0178855 |
| *bli-6* | 44792.14 | 11167.86 | -1.80 | 0.56 | 0.001287 | 0.0150705 |
| *mdh-2* | 40013.18 | 10392.08 | -1.80 | 0.49 | 0.000221 | 0.0038443 |
| *F53C11.3* | 2124.53 | 547.50 | -1.80 | 0.50 | 0.000309 | 0.0049901 |
| *pgp-6* | 398.68 | 94.38 | -1.80 | 0.62 | 0.003975 | 0.0347344 |
| *ent-4* | 535.95 | 122.82 | -1.80 | 0.66 | 0.0063 | 0.0481799 |
| *noah-2* | 5624.72 | 1456.08 | -1.80 | 0.48 | 0.000196 | 0.0034933 |
| *F54H5.14* | 148.53 | 36.31 | -1.80 | 0.58 | 0.001814 | 0.0195014 |
| *dhcr-24* | 1100.64 | 289.34 | -1.80 | 0.45 | 5.75E-05 | 0.0013008 |
| *F07A11.5* | 800.94 | 201.93 | -1.80 | 0.53 | 0.000643 | 0.0087421 |
| *R12C12.10* | 71.70 | 16.89 | -1.81 | 0.62 | 0.003754 | 0.0333553 |
| *clec-170* | 620.30 | 147.44 | -1.81 | 0.61 | 0.003223 | 0.0297484 |
| *nlp-36* | 8007.78 | 1973.82 | -1.81 | 0.56 | 0.001331 | 0.0154277 |
| *lgc-34* | 836.43 | 213.46 | -1.81 | 0.50 | 0.000344 | 0.0054306 |
| *Y38A10A.2* | 326.94 | 83.73 | -1.81 | 0.50 | 0.000274 | 0.004545 |
| *clec-56* | 1424.80 | 369.86 | -1.81 | 0.47 | 0.000118 | 0.0023071 |
| *hrg-6* | 105.77 | 25.36 | -1.81 | 0.60 | 0.002507 | 0.0249306 |
| *F22B8.4* | 242.55 | 55.93 | -1.81 | 0.64 | 0.004985 | 0.0408864 |
| *hhat-1* | 716.15 | 178.85 | -1.81 | 0.54 | 0.000768 | 0.0101248 |
| *C33G3.4* | 395.95 | 101.50 | -1.81 | 0.49 | 0.000226 | 0.0038942 |
| *cdh-7* | 533.86 | 134.60 | -1.81 | 0.52 | 0.000513 | 0.0072704 |
| *C44B7.11* | 866.28 | 226.46 | -1.81 | 0.45 | 5.45E-05 | 0.0012558 |
| *Y71H2AM.14* | 138.72 | 34.02 | -1.81 | 0.56 | 0.001342 | 0.0154997 |
| *ptr-14* | 976.26 | 252.56 | -1.81 | 0.47 | 0.000107 | 0.0021438 |
| *kat-1* | 6124.34 | 1571.38 | -1.81 | 0.48 | 0.000173 | 0.0031734 |
| *mtp-18* | 333.91 | 83.01 | -1.82 | 0.54 | 0.000721 | 0.0096387 |
| *grd-5* | 10660.10 | 2750.25 | -1.82 | 0.47 | 0.000108 | 0.0021531 |
| *F42A10.7* | 7585.84 | 1859.84 | -1.82 | 0.55 | 0.000982 | 0.0122032 |
| *F46G10.4* | 633.22 | 160.72 | -1.82 | 0.49 | 0.000211 | 0.0036835 |
| *F18F11.4* | 151.48 | 36.62 | -1.82 | 0.57 | 0.001386 | 0.0158544 |
| *noah-1* | 5591.10 | 1404.99 | -1.83 | 0.51 | 0.000318 | 0.0051022 |
| *F33D4.6* | 5762.45 | 1401.11 | -1.83 | 0.56 | 0.001108 | 0.0134472 |
| *F32D1.11* | 292.09 | 71.01 | -1.83 | 0.56 | 0.001094 | 0.0133237 |
| *F54D5.12* | 3430.93 | 882.69 | -1.83 | 0.46 | 5.94E-05 | 0.0013256 |
| *T09B4.8* | 3037.41 | 753.22 | -1.83 | 0.53 | 0.000509 | 0.0072386 |
| *C26B9.5* | 4516.93 | 1114.07 | -1.83 | 0.53 | 0.000627 | 0.0085618 |
| *bus-17* | 137.02 | 31.79 | -1.83 | 0.62 | 0.003128 | 0.0291252 |
| *mct-4* | 3011.82 | 778.02 | -1.83 | 0.45 | 3.97E-05 | 0.0009832 |
| *W10G11.19* | 725.12 | 182.63 | -1.83 | 0.50 | 0.000223 | 0.0038539 |
| *Y51H7C.1* | 4224.49 | 1051.37 | -1.83 | 0.52 | 0.000388 | 0.0059388 |
| *C17C3.1* | 848.99 | 214.96 | -1.83 | 0.48 | 0.000137 | 0.0025854 |
| *ugt-6* | 1174.40 | 298.07 | -1.83 | 0.48 | 0.000116 | 0.0022753 |
| *erd-2.1* | 3576.31 | 913.60 | -1.83 | 0.46 | 6.84E-05 | 0.001488 |
| *clec-150* | 9091.97 | 2228.67 | -1.83 | 0.54 | 0.00064 | 0.0087234 |
| *madf-5* | 689.64 | 175.73 | -1.84 | 0.46 | 7.19E-05 | 0.0015443 |
| *C50D2.6* | 137.86 | 33.32 | -1.84 | 0.56 | 0.000951 | 0.0119062 |

| | | | | | | |
|---|---|---|---|---|---|---|
| *dhs-3* | 2749.20 | 701.74 | -1.84 | 0.46 | 5.75E-05 | 0.0013008 |
| *C48B6.10* | 2135.06 | 546.35 | -1.84 | 0.45 | 4.35E-05 | 0.0010568 |
| *frpr-5* | 178.27 | 43.78 | -1.84 | 0.53 | 0.000459 | 0.006708 |
| *nas-27* | 593.07 | 129.25 | -1.84 | 0.67 | 0.006143 | 0.0473362 |
| *ads-1* | 8045.37 | 1982.91 | -1.84 | 0.52 | 0.000372 | 0.0057495 |
| *atf-2* | 474.41 | 118.10 | -1.84 | 0.50 | 0.000223 | 0.0038539 |
| *gfi-1* | 12037.15 | 3083.59 | -1.85 | 0.44 | 2.51E-05 | 0.00066 |
| *col-169* | 8736.45 | 2204.62 | -1.85 | 0.47 | 8.27E-05 | 0.0017432 |
| *Y65B4BL.1* | 3486.97 | 885.20 | -1.85 | 0.45 | 4.78E-05 | 0.0011285 |
| *Y17D7B.7* | 920.83 | 227.00 | -1.85 | 0.51 | 0.000305 | 0.0049433 |
| *col-168* | 7298.85 | 1841.08 | -1.85 | 0.47 | 7.71E-05 | 0.0016399 |
| *cuc-1* | 2089.57 | 522.47 | -1.85 | 0.48 | 0.000129 | 0.0024895 |
| *poml-2* | 153.78 | 35.76 | -1.85 | 0.60 | 0.001877 | 0.0199836 |
| *T24C12.3* | 746.86 | 179.14 | -1.85 | 0.55 | 0.00079 | 0.0103572 |
| *ZK622.4* | 85.95 | 19.43 | -1.85 | 0.63 | 0.003203 | 0.0296111 |
| *ugt-30* | 227.49 | 51.87 | -1.85 | 0.62 | 0.002578 | 0.0253236 |
| *cyp-25A1* | 137.56 | 32.75 | -1.86 | 0.56 | 0.000845 | 0.0109222 |
| *ger-1* | 913.84 | 230.05 | -1.86 | 0.46 | 4.55E-05 | 0.0010876 |
| *clec-48* | 4529.81 | 1071.26 | -1.86 | 0.56 | 0.000999 | 0.012378 |
| *C14A4.9* | 887.08 | 205.60 | -1.86 | 0.59 | 0.0016 | 0.0177661 |
| *T08G2.2* | 170.31 | 39.95 | -1.86 | 0.57 | 0.001173 | 0.014007 |
| *W01A8.6* | 290.71 | 66.51 | -1.86 | 0.60 | 0.002055 | 0.0214726 |
| *spl-1* | 2545.47 | 637.37 | -1.86 | 0.46 | 4.74E-05 | 0.0011207 |
| *nhx-2* | 1815.58 | 453.40 | -1.87 | 0.46 | 4.94E-05 | 0.0011554 |
| *hacd-1* | 8324.70 | 1994.21 | -1.87 | 0.53 | 0.000474 | 0.0068857 |
| *F37A4.3* | 68.26 | 15.12 | -1.87 | 0.64 | 0.003267 | 0.0301375 |
| *C35A11.4* | 394.29 | 86.90 | -1.87 | 0.64 | 0.003564 | 0.032076 |
| *cbl-1* | 544.38 | 133.05 | -1.87 | 0.50 | 0.000176 | 0.0032112 |
| *F22H10.6* | 510.14 | 115.75 | -1.87 | 0.61 | 0.002009 | 0.0211247 |
| *ndx-3* | 449.79 | 109.72 | -1.87 | 0.50 | 0.00016 | 0.0029487 |
| *K08D8.3* | 791.73 | 188.63 | -1.88 | 0.53 | 0.000435 | 0.0064683 |
| *pfas-1* | 3902.20 | 979.13 | -1.88 | 0.43 | 1.50E-05 | 0.0004489 |
| *F54D5.15* | 84.19 | 17.52 | -1.88 | 0.69 | 0.00628 | 0.048123 |
| *cest-31* | 356.08 | 84.95 | -1.88 | 0.52 | 0.000329 | 0.0052372 |
| *nduo-4* | 8394.95 | 2087.82 | -1.88 | 0.44 | 2.29E-05 | 0.0006182 |
| *C34E7.3* | 115.15 | 26.18 | -1.88 | 0.59 | 0.001488 | 0.0167585 |
| *R102.11* | 77.16 | 17.15 | -1.88 | 0.62 | 0.002268 | 0.0231848 |
| *vha-8* | 33596.18 | 8240.53 | -1.88 | 0.47 | 5.70E-05 | 0.0013003 |
| *C50A2.3* | 212.19 | 50.08 | -1.89 | 0.53 | 0.000421 | 0.0063225 |
| *cutl-18* | 251.77 | 60.11 | -1.89 | 0.52 | 0.000253 | 0.0042823 |
| *F32H5.1* | 651.77 | 159.50 | -1.89 | 0.47 | 5.96E-05 | 0.0013271 |
| *ctsa-3.1* | 4068.23 | 965.92 | -1.89 | 0.52 | 0.000277 | 0.0045694 |
| *pmt-2* | 41052.79 | 9733.83 | -1.89 | 0.52 | 0.000288 | 0.0047221 |
| *sfxn-1.5* | 1236.46 | 291.16 | -1.89 | 0.53 | 0.000369 | 0.0057313 |
| *F14B8.6* | 1699.58 | 404.01 | -1.89 | 0.52 | 0.000241 | 0.0041099 |
| *bckd-1A* | 6652.75 | 1615.50 | -1.89 | 0.48 | 6.94E-05 | 0.0014999 |
| *dpyd-1* | 4342.38 | 1032.65 | -1.89 | 0.51 | 0.000228 | 0.0039064 |
| *amt-4* | 1916.21 | 425.03 | -1.89 | 0.61 | 0.00193 | 0.0204309 |
| *col-63* | 17907.71 | 4182.18 | -1.90 | 0.54 | 0.00042 | 0.0063107 |
| *R57.2* | 334.85 | 80.04 | -1.90 | 0.49 | 0.000121 | 0.0023694 |
| *F09F9.2* | 933.00 | 212.84 | -1.90 | 0.57 | 0.000828 | 0.0107839 |
| *C18B2.3* | 5267.75 | 1234.36 | -1.90 | 0.53 | 0.000323 | 0.00515 |
| *mrps-28* | 2170.46 | 521.37 | -1.90 | 0.48 | 8.73E-05 | 0.001825 |
| *Y18H1A.9* | 481.39 | 114.75 | -1.90 | 0.50 | 0.000136 | 0.0025757 |
| *amx-3* | 228.12 | 53.62 | -1.90 | 0.52 | 0.000256 | 0.0043099 |
| *grd-14* | 4817.34 | 1126.45 | -1.90 | 0.53 | 0.000309 | 0.0049901 |

| | | | | | | |
|---|---|---|---|---|---|---|
| *C03B1.5* | 84.26 | 17.53 | -1.90 | 0.67 | 0.004311 | 0.0367862 |
| *alh-12* | 3700.67 | 864.85 | -1.91 | 0.53 | 0.000289 | 0.0047254 |
| *vha-10* | 22738.98 | 5478.55 | -1.91 | 0.47 | 5.22E-05 | 0.0012105 |
| *F57F4.4* | 20933.33 | 5140.30 | -1.91 | 0.43 | 9.05E-06 | 0.0003079 |
| *Y116F11A.6* | 150.30 | 32.27 | -1.91 | 0.63 | 0.002608 | 0.0255175 |
| *EGAP9.3* | 265.03 | 62.28 | -1.91 | 0.51 | 0.000201 | 0.0035598 |
| *ppat-1* | 601.99 | 145.59 | -1.91 | 0.46 | 3.27E-05 | 0.0008347 |
| *C35B1.5* | 4037.02 | 955.76 | -1.91 | 0.50 | 0.000129 | 0.0024895 |
| *Y22D7AL.11* | 211.03 | 46.67 | -1.91 | 0.60 | 0.001438 | 0.0163137 |
| *F01D5.10* | 323.95 | 75.31 | -1.91 | 0.53 | 0.000297 | 0.0048283 |
| *hrg-1* | 626.04 | 133.60 | -1.91 | 0.64 | 0.002694 | 0.0261107 |
| *dhs-2* | 395.91 | 93.70 | -1.91 | 0.50 | 0.000112 | 0.002222 |
| *C15H9.9* | 4455.11 | 1053.31 | -1.91 | 0.50 | 0.000118 | 0.0023109 |
| *F59B10.5* | 684.22 | 161.19 | -1.91 | 0.50 | 0.000134 | 0.0025455 |
| *clec-218* | 605.37 | 144.46 | -1.92 | 0.48 | 5.74E-05 | 0.0013008 |
| *cest-7* | 145.26 | 32.97 | -1.92 | 0.56 | 0.000602 | 0.0082724 |
| *F33A8.10* | 126.52 | 28.32 | -1.92 | 0.57 | 0.000843 | 0.0109167 |
| *H06H21.8* | 953.65 | 225.31 | -1.92 | 0.49 | 9.30E-05 | 0.001916 |
| *cest-6* | 260.39 | 61.17 | -1.92 | 0.50 | 0.000114 | 0.0022616 |
| *lbp-5* | 1451.38 | 332.20 | -1.92 | 0.54 | 0.000377 | 0.0058023 |
| *grl-7* | 3832.82 | 884.61 | -1.92 | 0.53 | 0.000263 | 0.0044023 |
| *F56C9.7* | 6105.03 | 1388.25 | -1.92 | 0.55 | 0.000443 | 0.0065659 |
| *F46G11.4* | 106.19 | 22.21 | -1.92 | 0.65 | 0.002968 | 0.0280147 |
| *col-73* | 54253.25 | 12973.53 | -1.92 | 0.46 | 2.80E-05 | 0.0007349 |
| *E01G4.6* | 25149.79 | 5546.45 | -1.93 | 0.58 | 0.000978 | 0.0121625 |
| *mtrr-1* | 4167.18 | 1001.13 | -1.93 | 0.44 | 1.14E-05 | 0.0003649 |
| *umps-1* | 4115.07 | 946.67 | -1.93 | 0.52 | 0.000208 | 0.0036444 |
| *art-1* | 15055.99 | 3517.15 | -1.93 | 0.49 | 8.04E-05 | 0.0017016 |
| *Y54F10AM.6* | 616.53 | 143.14 | -1.94 | 0.50 | 0.000103 | 0.0020647 |
| *pyk-2* | 1142.26 | 269.48 | -1.94 | 0.47 | 3.21E-05 | 0.0008221 |
| *T08B1.1* | 2708.90 | 625.75 | -1.94 | 0.50 | 0.000108 | 0.0021531 |
| *hprt-1* | 3804.77 | 886.92 | -1.94 | 0.48 | 6.12E-05 | 0.0013553 |
| *Y47G6A.15* | 4608.97 | 1065.26 | -1.94 | 0.50 | 0.000101 | 0.002049 |
| *vha-3* | 8977.89 | 2135.12 | -1.94 | 0.44 | 1.13E-05 | 0.0003641 |
| *rol-8* | 15221.79 | 3581.00 | -1.94 | 0.46 | 2.90E-05 | 0.0007535 |
| *pde-6* | 2089.52 | 492.42 | -1.94 | 0.46 | 2.47E-05 | 0.0006523 |
| *W02B12.1* | 1006.11 | 227.58 | -1.94 | 0.53 | 0.000257 | 0.0043128 |
| *col-180* | 6807.64 | 1536.47 | -1.94 | 0.54 | 0.000281 | 0.0046239 |
| *idh-1* | 9203.60 | 2150.43 | -1.95 | 0.47 | 4.11E-05 | 0.0010104 |
| *col-167* | 32536.80 | 7501.93 | -1.95 | 0.50 | 8.71E-05 | 0.001825 |
| *nas-4* | 74.86 | 14.34 | -1.95 | 0.71 | 0.005825 | 0.04565 |
| *mct-6* | 4929.39 | 1118.79 | -1.95 | 0.52 | 0.000193 | 0.0034378 |
| *C23H5.8* | 1802.28 | 411.81 | -1.95 | 0.51 | 0.000137 | 0.0025923 |
| *trap-4* | 25703.84 | 6002.82 | -1.95 | 0.47 | 3.38E-05 | 0.0008548 |
| *C25F6.7* | 505.56 | 110.93 | -1.95 | 0.57 | 0.00061 | 0.0083787 |
| *T01C8.3* | 73.67 | 14.23 | -1.95 | 0.70 | 0.005123 | 0.0416215 |
| *acaa-2* | 6830.35 | 1576.39 | -1.95 | 0.49 | 5.64E-05 | 0.0012892 |
| *vha-13* | 38013.19 | 8799.23 | -1.96 | 0.48 | 4.31E-05 | 0.00105 |
| *B0272.4* | 1179.02 | 259.10 | -1.96 | 0.56 | 0.000464 | 0.0067606 |
| *nas-37* | 1371.13 | 315.50 | -1.96 | 0.48 | 4.51E-05 | 0.00108 |
| *ZC434.10* | 209.96 | 44.09 | -1.96 | 0.61 | 0.001329 | 0.0154188 |
| *C52B11.5* | 536.41 | 122.36 | -1.97 | 0.49 | 6.27E-05 | 0.0013844 |
| *cpr-8* | 632.00 | 144.05 | -1.97 | 0.49 | 5.75E-05 | 0.0013008 |
| *C41H7.2* | 113.01 | 23.81 | -1.97 | 0.60 | 0.001025 | 0.0126363 |
| *ptr-16* | 653.04 | 150.59 | -1.97 | 0.46 | 2.00E-05 | 0.0005566 |
| *F23C8.5* | 10100.51 | 2241.94 | -1.97 | 0.53 | 0.000181 | 0.003278 |

| | | | | | | |
|---|---|---|---|---|---|---|
| *H23N18.5* | 1538.16 | 353.06 | -1.97 | 0.47 | 2.38E-05 | 0.0006358 |
| *pqn-32* | 9793.60 | 2209.01 | -1.98 | 0.50 | 7.14E-05 | 0.0015352 |
| *sucl-1* | 7580.53 | 1714.16 | -1.98 | 0.49 | 5.92E-05 | 0.0013244 |
| *ZK6.8* | 929.33 | 213.01 | -1.98 | 0.46 | 1.80E-05 | 0.0005119 |
| *C25B8.8* | 69.36 | 14.05 | -1.98 | 0.63 | 0.001744 | 0.0189607 |
| *C30G12.4* | 172.57 | 34.24 | -1.98 | 0.65 | 0.002289 | 0.0233524 |
| *B0454.8* | 198.87 | 39.76 | -1.99 | 0.64 | 0.001986 | 0.0209114 |
| *Y34B4A.6* | 16739.93 | 3688.13 | -1.99 | 0.52 | 0.000144 | 0.0027125 |
| *F19C6.5* | 411.29 | 90.29 | -1.99 | 0.53 | 0.000164 | 0.0030147 |
| *mam-2* | 470.40 | 106.45 | -1.99 | 0.48 | 2.94E-05 | 0.0007628 |
| *F28A12.3* | 53.38 | 10.18 | -1.99 | 0.68 | 0.003527 | 0.0318486 |
| *folt-1* | 165.71 | 35.39 | -1.99 | 0.57 | 0.000447 | 0.0066048 |
| *F13E9.13* | 240.59 | 53.36 | -1.99 | 0.51 | 9.02E-05 | 0.0018743 |
| *col-17* | 25934.66 | 5960.43 | -1.99 | 0.44 | 5.56E-06 | 0.0002113 |
| *R09H10.6* | 51.43 | 9.33 | -1.99 | 0.72 | 0.005321 | 0.042634 |
| *grl-5* | 3544.15 | 766.92 | -1.99 | 0.54 | 0.000238 | 0.0040614 |
| *ctsa-2* | 19878.76 | 4547.77 | -1.99 | 0.45 | 7.74E-06 | 0.000271 |
| *rft-1* | 249.13 | 54.80 | -2.00 | 0.51 | 9.16E-05 | 0.001897 |
| *lon-8* | 2263.79 | 510.47 | -2.00 | 0.46 | 1.64E-05 | 0.0004777 |
| *acdh-9* | 3481.91 | 773.78 | -2.00 | 0.49 | 4.49E-05 | 0.0010776 |
| *C18A11.3* | 1601.77 | 350.78 | -2.00 | 0.51 | 8.44E-05 | 0.0017726 |
| *W10G11.3* | 79.82 | 15.60 | -2.01 | 0.65 | 0.001967 | 0.0207319 |
| *R102.4* | 1311.85 | 286.23 | -2.01 | 0.51 | 9.23E-05 | 0.0019044 |
| *F11E6.9* | 167.08 | 34.15 | -2.01 | 0.60 | 0.000807 | 0.0105702 |
| *lips-14* | 53.64 | 9.81 | -2.01 | 0.70 | 0.004062 | 0.035251 |
| *wrt-2* | 2298.38 | 492.60 | -2.01 | 0.54 | 0.000184 | 0.0033216 |
| *K02B12.9* | 122.79 | 23.87 | -2.01 | 0.65 | 0.001954 | 0.0206337 |
| *F36G3.2* | 543.18 | 111.22 | -2.01 | 0.60 | 0.000726 | 0.0096782 |
| *C45G7.4* | 332.91 | 70.28 | -2.01 | 0.55 | 0.000284 | 0.0046589 |
| *R09H10.3* | 1556.21 | 332.90 | -2.02 | 0.53 | 0.000153 | 0.002849 |
| *agmo-1* | 1054.23 | 234.87 | -2.02 | 0.46 | 1.20E-05 | 0.0003766 |
| *oatr-1* | 8363.75 | 1851.90 | -2.02 | 0.47 | 1.78E-05 | 0.0005088 |
| *ndg-4* | 1874.17 | 413.56 | -2.02 | 0.48 | 2.26E-05 | 0.0006148 |
| *haf-9* | 3868.26 | 868.56 | -2.02 | 0.44 | 4.65E-06 | 0.000183 |
| *suca-1* | 8175.53 | 1777.22 | -2.02 | 0.50 | 5.61E-05 | 0.0012828 |
| *Y38E10A.14* | 403.56 | 81.60 | -2.02 | 0.60 | 0.000767 | 0.0101211 |
| *cah-3* | 5993.88 | 1246.77 | -2.02 | 0.57 | 0.000352 | 0.0055188 |
| *C34E7.4* | 4277.37 | 956.93 | -2.02 | 0.45 | 6.09E-06 | 0.0002268 |
| *mab-7* | 721.67 | 160.58 | -2.02 | 0.46 | 9.11E-06 | 0.0003085 |
| *cut-2* | 25745.89 | 5316.80 | -2.02 | 0.57 | 0.000414 | 0.0062706 |
| *C05D12.3* | 2952.39 | 662.45 | -2.03 | 0.44 | 3.50E-06 | 0.0001441 |
| *nkat-3* | 1838.74 | 398.38 | -2.03 | 0.50 | 4.97E-05 | 0.0011606 |
| *col-138* | 21675.27 | 4769.71 | -2.03 | 0.47 | 1.73E-05 | 0.0005012 |
| *T05H10.3* | 65.60 | 11.20 | -2.03 | 0.73 | 0.005612 | 0.0444099 |
| *col-154* | 14932.65 | 3173.96 | -2.03 | 0.53 | 0.000118 | 0.0023164 |
| *acbp-3* | 1449.70 | 316.78 | -2.03 | 0.48 | 2.39E-05 | 0.0006358 |
| *ram-2* | 40067.96 | 8658.47 | -2.03 | 0.50 | 4.07E-05 | 0.0010044 |
| *sfxn-2* | 353.20 | 75.89 | -2.03 | 0.50 | 5.46E-05 | 0.0012579 |
| *F52F10.2* | 143.14 | 29.26 | -2.03 | 0.58 | 0.000406 | 0.0061735 |
| *T06A4.1* | 194.07 | 39.69 | -2.04 | 0.57 | 0.000395 | 0.0060308 |
| *C15C7.7* | 807.82 | 173.15 | -2.04 | 0.51 | 5.71E-05 | 0.0013008 |
| *T02G5.7* | 5237.35 | 1128.97 | -2.04 | 0.49 | 3.81E-05 | 0.0009519 |
| *C06H5.6* | 1239.06 | 269.37 | -2.04 | 0.47 | 1.55E-05 | 0.0004572 |
| *samt-1* | 1109.44 | 244.28 | -2.04 | 0.45 | 4.80E-06 | 0.000187 |
| *bus-1* | 141.45 | 28.94 | -2.05 | 0.56 | 0.000279 | 0.0045982 |
| *ZK829.3* | 78.20 | 15.23 | -2.05 | 0.62 | 0.00091 | 0.0115565 |

| | | | | | | |
|---|---|---|---|---|---|---|
| *cut-6* | 108.89 | 21.83 | -2.05 | 0.58 | 0.000457 | 0.0067076 |
| *clec-10* | 792.94 | 159.76 | -2.05 | 0.58 | 0.000369 | 0.0057359 |
| *mpc-1* | 1859.50 | 396.33 | -2.05 | 0.49 | 3.29E-05 | 0.0008354 |
| *suro-1* | 1950.37 | 427.65 | -2.05 | 0.44 | 3.05E-06 | 0.00013 |
| *ugt-48* | 846.99 | 170.28 | -2.06 | 0.58 | 0.000351 | 0.0055074 |
| *C52D10.3* | 1674.10 | 362.28 | -2.06 | 0.46 | 8.66E-06 | 0.0002979 |
| *ZK287.4* | 196.76 | 33.48 | -2.06 | 0.72 | 0.004444 | 0.0376035 |
| *spp-4* | 2234.25 | 474.41 | -2.06 | 0.49 | 3.13E-05 | 0.0008067 |
| *R12E2.6* | 252.41 | 50.76 | -2.06 | 0.57 | 0.000315 | 0.0050581 |
| *H31G24.1* | 48.66 | 8.09 | -2.06 | 0.73 | 0.004752 | 0.0394904 |
| *ZC449.1* | 128.58 | 24.11 | -2.06 | 0.64 | 0.001341 | 0.0154997 |
| *mab-3* | 173.41 | 34.01 | -2.07 | 0.60 | 0.000536 | 0.007515 |
| *lpr-1* | 190.44 | 38.25 | -2.07 | 0.56 | 0.000227 | 0.003905 |
| *ugt-26* | 476.00 | 95.62 | -2.07 | 0.56 | 0.000226 | 0.0038942 |
| *txdc-12.1* | 4982.73 | 1083.98 | -2.07 | 0.43 | 1.44E-06 | 6.99E-05 |
| *pyr-1* | 4499.38 | 960.78 | -2.08 | 0.46 | 6.54E-06 | 0.0002398 |
| *col-77* | 35101.31 | 7373.28 | -2.08 | 0.49 | 2.10E-05 | 0.0005818 |
| *sqt-2* | 10836.86 | 2327.06 | -2.08 | 0.44 | 2.71E-06 | 0.0001178 |
| *F27D4.1* | 10629.96 | 2190.46 | -2.08 | 0.51 | 4.65E-05 | 0.0011057 |
| *C07E3.10* | 279.68 | 56.77 | -2.09 | 0.53 | 8.73E-05 | 0.001825 |
| *gsto-1* | 1080.19 | 222.59 | -2.09 | 0.51 | 4.09E-05 | 0.0010065 |
| *Y39G8B.1* | 4123.05 | 860.78 | -2.09 | 0.48 | 1.52E-05 | 0.0004541 |
| *gst-38* | 356.45 | 69.85 | -2.09 | 0.57 | 0.000275 | 0.0045592 |
| *F58H10.1* | 436.91 | 91.69 | -2.09 | 0.47 | 9.73E-06 | 0.0003251 |
| *clec-76* | 73.95 | 13.12 | -2.09 | 0.67 | 0.001835 | 0.0196854 |
| *F55F3.4* | 76.83 | 14.37 | -2.09 | 0.62 | 0.000762 | 0.0100561 |
| *ptr-4* | 2125.85 | 445.24 | -2.10 | 0.47 | 7.67E-06 | 0.0002698 |
| *C29F3.7* | 4137.37 | 867.36 | -2.10 | 0.47 | 6.60E-06 | 0.0002406 |
| *F15H10.8* | 298.99 | 60.37 | -2.10 | 0.53 | 6.74E-05 | 0.0014697 |
| *M02H5.8* | 1977.55 | 402.51 | -2.10 | 0.51 | 4.56E-05 | 0.0010891 |
| *R07E3.2* | 59.26 | 9.32 | -2.10 | 0.75 | 0.004972 | 0.0408442 |
| *gst-35* | 241.89 | 45.83 | -2.10 | 0.60 | 0.00052 | 0.0073361 |
| *dpy-2* | 1681.52 | 350.67 | -2.10 | 0.47 | 9.16E-06 | 0.0003094 |
| *ptr-18* | 1909.95 | 403.46 | -2.10 | 0.45 | 2.63E-06 | 0.0001148 |
| *Y51A2D.8* | 86.79 | 15.45 | -2.10 | 0.66 | 0.001437 | 0.0163124 |
| *F31D4.8* | 255.20 | 51.46 | -2.10 | 0.52 | 5.75E-05 | 0.0013008 |
| *zmp-2* | 490.27 | 96.85 | -2.10 | 0.55 | 0.000133 | 0.0025357 |
| *rol-1* | 38616.59 | 7740.57 | -2.11 | 0.53 | 6.32E-05 | 0.001389 |
| *F10D2.10* | 158.97 | 30.25 | -2.11 | 0.59 | 0.000377 | 0.0058023 |
| *Y71H10B.1* | 7736.92 | 1577.63 | -2.11 | 0.50 | 2.49E-05 | 0.0006569 |
| *F13H8.5* | 8838.43 | 1844.67 | -2.11 | 0.46 | 3.99E-06 | 0.000161 |
| *gcsh-2* | 3689.89 | 751.71 | -2.11 | 0.50 | 2.16E-05 | 0.0005944 |
| *T26G10.5* | 85.90 | 13.96 | -2.11 | 0.72 | 0.003362 | 0.0306434 |
| *wrt-9* | 1658.42 | 348.01 | -2.11 | 0.44 | 1.72E-06 | 8.16E-05 |
| *cnc-3* | 456.44 | 92.72 | -2.11 | 0.50 | 2.11E-05 | 0.0005839 |
| *ckc-1* | 1447.84 | 282.80 | -2.12 | 0.55 | 0.000128 | 0.0024777 |
| *T05C1.3* | 133.99 | 25.91 | -2.12 | 0.56 | 0.000164 | 0.0030147 |
| *C01B9.1* | 257.20 | 50.58 | -2.12 | 0.54 | 8.44E-05 | 0.0017726 |
| *got-1.2* | 3605.51 | 727.01 | -2.12 | 0.50 | 2.25E-05 | 0.0006146 |
| *B0272.3* | 2746.33 | 549.10 | -2.12 | 0.51 | 3.28E-05 | 0.0008352 |
| *col-49* | 10576.66 | 2139.49 | -2.12 | 0.49 | 1.64E-05 | 0.0004777 |
| *lon-1* | 1616.47 | 332.42 | -2.13 | 0.46 | 4.38E-06 | 0.0001738 |
| *T13C5.3* | 56.78 | 9.48 | -2.13 | 0.69 | 0.002161 | 0.0222597 |
| *ZC123.1* | 1807.11 | 363.50 | -2.13 | 0.50 | 1.81E-05 | 0.0005158 |
| *T15B7.10* | 66.68 | 11.69 | -2.13 | 0.66 | 0.001154 | 0.0138468 |
| *ahcy-1* | 147523.81 | 29844.01 | -2.13 | 0.48 | 1.05E-05 | 0.0003427 |

| | | | | | | |
|---|---|---|---|---|---|---|
| *C17E7.13* | 46.71 | 7.66 | -2.13 | 0.70 | 0.002391 | 0.0241259 |
| *C31C9.2* | 2906.79 | 587.10 | -2.13 | 0.48 | 1.09E-05 | 0.0003551 |
| *abu-12* | 2901.50 | 601.85 | -2.13 | 0.43 | 8.10E-07 | 4.35E-05 |
| *F53B1.4* | 6342.82 | 1303.44 | -2.13 | 0.45 | 2.17E-06 | 9.75E-05 |
| *C44B7.7* | 452.32 | 87.63 | -2.13 | 0.54 | 8.85E-05 | 0.0018432 |
| *dpy-10* | 1316.96 | 262.06 | -2.14 | 0.50 | 2.28E-05 | 0.0006162 |
| *R07B1.6* | 42.00 | 6.28 | -2.14 | 0.75 | 0.004474 | 0.0377917 |
| *F55G11.2* | 115.94 | 20.94 | -2.15 | 0.61 | 0.000475 | 0.0068921 |
| *C15F1.2* | 362.20 | 70.39 | -2.15 | 0.53 | 4.25E-05 | 0.0010376 |
| *K11H12.11* | 47.78 | 7.30 | -2.15 | 0.74 | 0.003529 | 0.0318486 |
| *K02E2.8* | 175.43 | 32.39 | -2.15 | 0.59 | 0.000241 | 0.0041099 |
| *C41H7.2* | 146.17 | 26.61 | -2.15 | 0.60 | 0.000344 | 0.0054341 |
| *mltn-12* | 182.44 | 33.33 | -2.16 | 0.59 | 0.000285 | 0.0046708 |
| *mec-7* | 2376.24 | 443.94 | -2.16 | 0.57 | 0.000141 | 0.0026631 |
| *slc-25A10* | 4181.98 | 815.40 | -2.16 | 0.51 | 2.14E-05 | 0.0005908 |
| *lpr-5* | 3147.12 | 618.12 | -2.16 | 0.50 | 1.28E-05 | 0.0003948 |
| *K03B8.6* | 1257.13 | 240.78 | -2.16 | 0.53 | 4.89E-05 | 0.001148 |
| *Y105E8A.13* | 317.06 | 62.47 | -2.16 | 0.49 | 9.83E-06 | 0.0003263 |
| *lbp-6* | 9765.19 | 1872.08 | -2.16 | 0.53 | 4.39E-05 | 0.0010628 |
| *C15C6.1* | 814.38 | 161.02 | -2.17 | 0.48 | 5.87E-06 | 0.0002198 |
| *cutl-28* | 191.66 | 35.88 | -2.17 | 0.56 | 9.67E-05 | 0.0019711 |
| *W08E12.6* | 45.97 | 6.71 | -2.17 | 0.75 | 0.003988 | 0.0347882 |
| *bcat-1* | 13524.51 | 2666.87 | -2.17 | 0.48 | 6.45E-06 | 0.0002382 |
| *smd-1* | 37009.10 | 6994.11 | -2.17 | 0.54 | 6.59E-05 | 0.0014431 |
| *acn-1* | 4446.83 | 890.60 | -2.17 | 0.45 | 1.54E-06 | 7.42E-05 |
| *Y38H8A.1* | 265.64 | 51.43 | -2.17 | 0.51 | 1.90E-05 | 0.000535 |
| *cah-5* | 644.36 | 110.79 | -2.17 | 0.64 | 0.000714 | 0.0095707 |
| *nlp-68* | 113.36 | 19.91 | -2.17 | 0.62 | 0.000464 | 0.0067606 |
| *F35C12.3* | 130.18 | 23.91 | -2.18 | 0.57 | 0.000115 | 0.0022687 |
| *asp-13* | 16063.80 | 3077.67 | -2.18 | 0.51 | 1.80E-05 | 0.0005119 |
| *sptl-2* | 1103.63 | 211.80 | -2.18 | 0.50 | 1.47E-05 | 0.0004426 |
| *F14D7.6* | 529.68 | 96.70 | -2.19 | 0.57 | 0.000126 | 0.0024431 |
| *W09G12.10* | 247.17 | 47.18 | -2.19 | 0.51 | 1.75E-05 | 0.0005038 |
| *zipt-2.3* | 152.87 | 24.48 | -2.19 | 0.69 | 0.001401 | 0.0159808 |
| *ugt-62* | 2486.56 | 453.78 | -2.19 | 0.56 | 9.62E-05 | 0.0019639 |
| *linc-40* | 136.93 | 23.51 | -2.20 | 0.63 | 0.000443 | 0.0065659 |
| *ZK154.1* | 1056.05 | 204.65 | -2.20 | 0.47 | 3.43E-06 | 0.0001419 |
| *ugt-16* | 246.70 | 46.52 | -2.20 | 0.51 | 1.97E-05 | 0.0005518 |
| *grl-15* | 1944.61 | 367.34 | -2.20 | 0.51 | 1.78E-05 | 0.0005088 |
| *spp-17* | 5622.45 | 1088.22 | -2.20 | 0.47 | 2.32E-06 | 0.0001032 |
| *T13F3.6* | 880.40 | 161.32 | -2.20 | 0.55 | 5.83E-05 | 0.0013132 |
| *ugt-46* | 2202.61 | 432.62 | -2.20 | 0.44 | 4.94E-07 | 2.89E-05 |
| *F37C4.6* | 1948.23 | 361.50 | -2.21 | 0.53 | 3.20E-05 | 0.0008221 |
| *mtl-2* | 206.74 | 36.89 | -2.21 | 0.58 | 0.000129 | 0.0024895 |
| *C12D12.1* | 24185.47 | 4553.28 | -2.21 | 0.50 | 1.11E-05 | 0.0003582 |
| *T26C5.2* | 861.52 | 152.71 | -2.21 | 0.58 | 0.000134 | 0.0025455 |
| *fkb-3* | 2708.08 | 520.33 | -2.21 | 0.47 | 2.09E-06 | 9.49E-05 |
| *Y106G6H.1* | 2627.61 | 472.89 | -2.22 | 0.56 | 6.95E-05 | 0.0014999 |
| *col-130* | 19698.84 | 3691.74 | -2.22 | 0.50 | 9.91E-06 | 0.000328 |
| *trpp-12* | 614.54 | 116.19 | -2.22 | 0.49 | 5.11E-06 | 0.0001966 |
| *dod-19* | 10181.97 | 1843.79 | -2.22 | 0.55 | 4.98E-05 | 0.0011606 |
| *sur-5* | 4754.79 | 853.87 | -2.22 | 0.56 | 6.86E-05 | 0.0014894 |
| *let-767* | 19805.00 | 3737.96 | -2.22 | 0.49 | 5.11E-06 | 0.0001966 |
| *B0554.4* | 72.30 | 11.26 | -2.22 | 0.69 | 0.001187 | 0.014142 |
| *dpy-9* | 1366.70 | 260.22 | -2.23 | 0.47 | 1.75E-06 | 8.26E-05 |
| *B0365.9* | 483.70 | 89.27 | -2.23 | 0.51 | 1.48E-05 | 0.0004452 |

| | | | | | | |
|---|---|---|---|---|---|---|
| *clec-75* | 75.70 | 11.57 | -2.23 | 0.70 | 0.001377 | 0.0157807 |
| *ech-7* | 3716.57 | 683.73 | -2.23 | 0.52 | 1.66E-05 | 0.0004816 |
| *aqp-11* | 2869.19 | 535.94 | -2.23 | 0.49 | 6.62E-06 | 0.0002406 |
| *ldp-1* | 15928.80 | 2942.59 | -2.23 | 0.51 | 1.19E-05 | 0.0003759 |
| *F54B11.10* | 41.43 | 5.77 | -2.23 | 0.75 | 0.002761 | 0.0265807 |
| *F19C7.1* | 22882.08 | 4150.99 | -2.23 | 0.53 | 2.99E-05 | 0.0007748 |
| *T28C12.6* | 75.24 | 11.08 | -2.23 | 0.72 | 0.001874 | 0.0199592 |
| *C35A5.5* | 43.99 | 5.83 | -2.24 | 0.77 | 0.003547 | 0.0319812 |
| *K02D3.2* | 50.46 | 7.46 | -2.24 | 0.71 | 0.001627 | 0.0179524 |
| *irg-4* | 1148.84 | 193.19 | -2.24 | 0.61 | 0.00025 | 0.0042354 |
| *ZC513.2* | 157.25 | 27.44 | -2.24 | 0.57 | 8.79E-05 | 0.0018337 |
| *grl-16* | 33111.24 | 6202.88 | -2.24 | 0.47 | 1.96E-06 | 8.93E-05 |
| *F46F3.3* | 69.77 | 11.15 | -2.25 | 0.65 | 0.000578 | 0.0080327 |
| *K01D12.5* | 221.52 | 29.76 | -2.25 | 0.76 | 0.003016 | 0.0283809 |
| *Y119D3B.21* | 12611.60 | 2274.60 | -2.25 | 0.52 | 1.32E-05 | 0.0004065 |
| *Y61A9LA.7* | 38.87 | 4.87 | -2.26 | 0.78 | 0.003763 | 0.0333584 |
| *F47B8.5* | 84.93 | 13.83 | -2.26 | 0.62 | 0.000293 | 0.0047865 |
| *C26B2.8* | 69.59 | 9.49 | -2.26 | 0.75 | 0.002399 | 0.0241545 |
| *Y18H1A.8* | 29.46 | 3.26 | -2.26 | 0.82 | 0.005679 | 0.0448218 |
| *F28H7.3* | 5722.85 | 1001.49 | -2.26 | 0.55 | 3.75E-05 | 0.0009392 |
| *M03B6.3* | 534.78 | 94.50 | -2.26 | 0.53 | 2.29E-05 | 0.0006177 |
| *C30H6.5* | 1067.02 | 188.00 | -2.27 | 0.54 | 2.46E-05 | 0.0006514 |
| *T11B7.2* | 353.12 | 62.66 | -2.27 | 0.53 | 1.67E-05 | 0.000486 |
| *C49F5.9* | 63.20 | 9.70 | -2.27 | 0.67 | 0.000676 | 0.0091462 |
| *col-46* | 343.56 | 61.90 | -2.27 | 0.50 | 6.13E-06 | 0.0002277 |
| *paic-1* | 3378.96 | 627.00 | -2.27 | 0.45 | 5.57E-07 | 3.20E-05 |
| *dpy-7* | 2074.05 | 376.76 | -2.27 | 0.49 | 3.31E-06 | 0.0001391 |
| *col-155* | 12386.35 | 2191.06 | -2.28 | 0.52 | 1.21E-05 | 0.0003784 |
| *F35D2.1* | 120.31 | 18.70 | -2.28 | 0.65 | 0.000498 | 0.0071416 |
| *gst-27* | 5608.91 | 985.19 | -2.28 | 0.53 | 1.59E-05 | 0.0004682 |
| *Y71F9B.9* | 141.29 | 23.59 | -2.28 | 0.59 | 0.0001 | 0.00203 |
| *asns-1* | 683.05 | 111.91 | -2.28 | 0.60 | 0.00016 | 0.0029561 |
| *T05C12.4* | 68.33 | 9.17 | -2.28 | 0.74 | 0.002095 | 0.0217703 |
| *gst-6* | 907.61 | 154.07 | -2.29 | 0.56 | 4.61E-05 | 0.0010975 |
| *igcm-1* | 150.21 | 22.72 | -2.29 | 0.67 | 0.000577 | 0.0080196 |
| *F58G6.3* | 494.44 | 77.61 | -2.29 | 0.64 | 0.000317 | 0.0050888 |
| *sec-61.G* | 21629.84 | 3881.16 | -2.29 | 0.48 | 1.72E-06 | 8.16E-05 |
| *T14E8.4* | 54.41 | 6.49 | -2.30 | 0.79 | 0.003595 | 0.0323024 |
| *col-162* | 14614.24 | 2573.85 | -2.30 | 0.51 | 5.77E-06 | 0.0002179 |
| *R04B5.6* | 37.40 | 4.73 | -2.30 | 0.76 | 0.002575 | 0.025318 |
| *grd-12* | 81.79 | 12.39 | -2.30 | 0.66 | 0.000505 | 0.0072016 |
| *phat-5* | 40.54 | 4.30 | -2.30 | 0.82 | 0.005217 | 0.0421145 |
| *T10B5.10* | 524.72 | 93.93 | -2.30 | 0.47 | 1.11E-06 | 5.63E-05 |
| *elo-2* | 8094.70 | 1288.00 | -2.30 | 0.62 | 0.000187 | 0.0033756 |
| *col-99* | 795.85 | 143.34 | -2.31 | 0.46 | 5.48E-07 | 3.17E-05 |
| *K11H12.4* | 433.41 | 70.04 | -2.31 | 0.60 | 0.000124 | 0.0024168 |
| *C02E7.7* | 11735.67 | 2028.18 | -2.31 | 0.52 | 9.96E-06 | 0.0003289 |
| *trap-3* | 26068.45 | 4614.15 | -2.31 | 0.48 | 1.74E-06 | 8.22E-05 |
| *H20E11.2* | 28.24 | 3.13 | -2.31 | 0.80 | 0.004069 | 0.0352889 |
| *vha-11* | 15193.20 | 2749.21 | -2.31 | 0.44 | 1.91E-07 | 1.30E-05 |
| *lpr-3* | 7454.59 | 1337.80 | -2.31 | 0.46 | 4.19E-07 | 2.51E-05 |
| *F20G2.2* | 1965.53 | 325.76 | -2.32 | 0.56 | 3.84E-05 | 0.0009568 |
| *pcp-2* | 3137.43 | 542.53 | -2.32 | 0.51 | 4.76E-06 | 0.0001868 |
| *K08C7.1* | 335.92 | 57.08 | -2.32 | 0.53 | 1.19E-05 | 0.0003766 |
| *F41F3.8* | 305.38 | 52.95 | -2.32 | 0.50 | 3.38E-06 | 0.0001409 |
| *C39B5.5* | 2031.01 | 352.47 | -2.32 | 0.50 | 3.18E-06 | 0.0001343 |

| | | | | | | |
|---|---|---|---|---|---|---|
| *ZC334.7* | 28.35 | 2.75 | -2.33 | 0.84 | 0.005386 | 0.0430428 |
| *bas-1* | 876.17 | 148.59 | -2.33 | 0.53 | 9.79E-06 | 0.0003262 |
| *M05D6.9* | 93.09 | 14.69 | -2.33 | 0.60 | 0.000119 | 0.0023241 |
| *gldc-1* | 7179.31 | 1248.75 | -2.33 | 0.49 | 1.80E-06 | 8.41E-05 |
| *dhs-14* | 558.81 | 94.81 | -2.33 | 0.52 | 7.09E-06 | 0.0002529 |
| *bus-8* | 1620.32 | 288.17 | -2.33 | 0.45 | 1.62E-07 | 1.12E-05 |
| *elo-9* | 362.95 | 60.41 | -2.34 | 0.54 | 1.59E-05 | 0.0004679 |
| *K08E7.5* | 1960.51 | 332.54 | -2.34 | 0.52 | 5.83E-06 | 0.0002192 |
| *ngn-1* | 49.93 | 5.90 | -2.34 | 0.77 | 0.002561 | 0.0252696 |
| *C18H7.11* | 614.54 | 99.04 | -2.34 | 0.57 | 4.41E-05 | 0.0010633 |
| *Y46D2A.2* | 1081.47 | 172.48 | -2.34 | 0.58 | 5.97E-05 | 0.0013294 |
| *grd-1* | 717.93 | 120.60 | -2.35 | 0.52 | 5.74E-06 | 0.0002176 |
| *mec-12* | 3669.18 | 611.78 | -2.35 | 0.52 | 6.75E-06 | 0.0002441 |
| *C06G4.6* | 105.73 | 16.19 | -2.35 | 0.61 | 0.000126 | 0.0024503 |
| *cln-3.1* | 248.10 | 41.19 | -2.36 | 0.53 | 7.32E-06 | 0.0002598 |
| *C04G6.7* | 23.88 | 1.99 | -2.36 | 0.86 | 0.005841 | 0.045726 |
| *F56C3.8* | 248.23 | 38.88 | -2.36 | 0.59 | 6.07E-05 | 0.0013483 |
| *pcp-3* | 8104.90 | 1251.35 | -2.36 | 0.60 | 8.95E-05 | 0.001861 |
| *Y52B11B.1* | 25.00 | 2.37 | -2.36 | 0.83 | 0.004462 | 0.0377115 |
| *Y71G12B.25* | 3613.49 | 632.83 | -2.36 | 0.43 | 5.16E-08 | 4.28E-06 |
| *F58G6.7* | 881.06 | 144.08 | -2.36 | 0.54 | 1.02E-05 | 0.0003356 |
| *F54D10.8* | 193.76 | 28.02 | -2.36 | 0.65 | 0.000304 | 0.0049348 |
| *pept-1* | 7620.44 | 1066.16 | -2.36 | 0.68 | 0.000507 | 0.0072163 |
| *bli-1* | 14106.81 | 2313.11 | -2.37 | 0.53 | 8.62E-06 | 0.0002971 |
| *Y55D5A.4* | 179.91 | 27.66 | -2.37 | 0.60 | 7.81E-05 | 0.0016559 |
| *C40H1.8* | 294.76 | 45.64 | -2.37 | 0.59 | 6.32E-05 | 0.001389 |
| *ucr-2.2* | 7726.04 | 1278.88 | -2.37 | 0.51 | 3.68E-06 | 0.0001501 |
| *col-88* | 12747.58 | 2061.56 | -2.37 | 0.54 | 1.21E-05 | 0.0003784 |
| *mlt-11* | 12744.00 | 2155.57 | -2.37 | 0.48 | 7.54E-07 | 4.14E-05 |
| *sago-2* | 430.17 | 67.85 | -2.38 | 0.57 | 2.93E-05 | 0.0007607 |
| *pcp-4* | 3590.69 | 601.48 | -2.38 | 0.49 | 1.14E-06 | 5.76E-05 |
| *M01H9.5* | 43.37 | 5.06 | -2.38 | 0.77 | 0.001906 | 0.0202227 |
| *bli-2* | 7884.87 | 1254.22 | -2.38 | 0.55 | 1.70E-05 | 0.0004936 |
| *Y54G2A.76* | 231.73 | 34.77 | -2.38 | 0.61 | 9.50E-05 | 0.0019407 |
| *Y54G2A.45* | 1408.52 | 232.45 | -2.39 | 0.50 | 1.99E-06 | 9.03E-05 |
| *nas-8* | 206.14 | 30.73 | -2.39 | 0.61 | 0.000102 | 0.0020596 |
| *nep-12* | 148.31 | 22.30 | -2.39 | 0.60 | 7.48E-05 | 0.0015972 |
| *lec-9* | 18837.37 | 2992.47 | -2.39 | 0.55 | 1.14E-05 | 0.0003647 |
| *B0334.6* | 64.44 | 8.69 | -2.39 | 0.69 | 0.00049 | 0.0070632 |
| *scav-4* | 897.08 | 131.89 | -2.40 | 0.62 | 0.000117 | 0.0023071 |
| *swip-10* | 182.29 | 28.87 | -2.40 | 0.54 | 1.05E-05 | 0.0003427 |
| *aat-1* | 660.37 | 104.83 | -2.40 | 0.54 | 9.82E-06 | 0.0003263 |
| *mlt-8* | 5515.79 | 940.80 | -2.40 | 0.43 | 2.98E-08 | 2.73E-06 |
| *M02D8.5* | 345.24 | 54.85 | -2.40 | 0.54 | 7.88E-06 | 0.0002747 |
| *alh-13* | 4483.30 | 762.98 | -2.40 | 0.44 | 3.76E-08 | 3.36E-06 |
| *tba-4* | 8227.48 | 1313.34 | -2.40 | 0.53 | 5.51E-06 | 0.0002109 |
| *pud-3* | 154.31 | 16.60 | -2.40 | 0.79 | 0.002317 | 0.0235501 |
| *M04C9.3* | 229.79 | 35.47 | -2.41 | 0.56 | 1.91E-05 | 0.0005383 |
| *cyp-13A12* | 90.61 | 12.92 | -2.41 | 0.64 | 0.000158 | 0.0029264 |
| *col-68* | 248.63 | 38.65 | -2.41 | 0.55 | 1.18E-05 | 0.0003738 |
| *Y73E7A.8* | 832.32 | 131.08 | -2.42 | 0.53 | 5.10E-06 | 0.0001966 |
| *dlhd-1* | 674.70 | 107.43 | -2.43 | 0.51 | 1.83E-06 | 8.51E-05 |
| *F44E7.4* | 3919.96 | 650.85 | -2.43 | 0.44 | 3.95E-08 | 3.51E-06 |
| *F58G6.9* | 551.89 | 78.18 | -2.44 | 0.63 | 0.000101 | 0.002049 |
| *M153.1* | 2842.76 | 459.65 | -2.44 | 0.48 | 3.01E-07 | 1.91E-05 |
| *gpa-6* | 36.00 | 4.02 | -2.44 | 0.76 | 0.001313 | 0.0153044 |

| | | | | | | |
|---|---|---|---|---|---|---|
| *F53C11.1* | 584.45 | 89.41 | -2.44 | 0.55 | 7.75E-06 | 0.000271 |
| *col-170* | 3636.85 | 585.94 | -2.44 | 0.48 | 2.85E-07 | 1.82E-05 |
| *vha-12* | 43190.86 | 6903.24 | -2.44 | 0.49 | 4.88E-07 | 2.87E-05 |
| *F13B6.2* | 999.82 | 157.21 | -2.45 | 0.51 | 1.30E-06 | 6.52E-05 |
| *trx-3* | 1215.22 | 162.84 | -2.45 | 0.66 | 0.000202 | 0.0035654 |
| *mel-32* | 27227.60 | 4318.04 | -2.45 | 0.49 | 4.88E-07 | 2.87E-05 |
| *T22B7.7* | 1830.02 | 279.89 | -2.46 | 0.53 | 4.01E-06 | 0.0001615 |
| *ver-4* | 136.08 | 19.36 | -2.46 | 0.61 | 5.26E-05 | 0.0012176 |
| *col-145* | 36222.88 | 5577.06 | -2.46 | 0.52 | 2.50E-06 | 0.0001096 |
| *col-157* | 10954.34 | 1749.62 | -2.46 | 0.47 | 1.74E-07 | 1.19E-05 |
| *R07B1.5* | 149.30 | 21.06 | -2.46 | 0.61 | 5.93E-05 | 0.0013244 |
| *Y39B6A.21* | 348.61 | 52.63 | -2.46 | 0.54 | 6.16E-06 | 0.0002284 |
| *F46F2.3* | 10274.10 | 1432.36 | -2.46 | 0.62 | 7.79E-05 | 0.0016531 |
| *F56D3.1* | 8308.66 | 1354.39 | -2.46 | 0.43 | 1.19E-08 | 1.21E-06 |
| *cky-1* | 20.67 | 0.77 | -2.46 | 0.90 | 0.006436 | 0.048978 |
| *lpr-6* | 1441.27 | 227.76 | -2.46 | 0.48 | 2.63E-07 | 1.71E-05 |
| *R07B1.7* | 36.93 | 3.97 | -2.47 | 0.76 | 0.001247 | 0.0146864 |
| *B0334.13* | 129.09 | 17.22 | -2.47 | 0.65 | 0.000152 | 0.002833 |
| *C17B7.12* | 37.55 | 4.06 | -2.47 | 0.76 | 0.001154 | 0.0138468 |
| *F26G1.11* | 81.35 | 10.95 | -2.47 | 0.64 | 0.000125 | 0.0024333 |
| *col-156* | 1864.42 | 287.06 | -2.48 | 0.50 | 8.24E-07 | 4.40E-05 |
| *T09D3.8* | 76.64 | 10.30 | -2.48 | 0.64 | 0.000104 | 0.0020802 |
| *R07B1.13* | 418.53 | 65.44 | -2.48 | 0.48 | 2.03E-07 | 1.37E-05 |
| *lagr-1* | 79.58 | 9.60 | -2.49 | 0.71 | 0.000442 | 0.0065566 |
| *F57H12.6* | 848.15 | 117.78 | -2.49 | 0.61 | 4.24E-05 | 0.0010361 |
| *T16G12.1* | 3438.41 | 532.42 | -2.49 | 0.48 | 1.62E-07 | 1.12E-05 |
| *M02G9.1* | 403.45 | 7.92 | -2.50 | 0.91 | 0.00636 | 0.0484703 |
| *W07A12.8* | 22.18 | 1.57 | -2.50 | 0.85 | 0.003429 | 0.0311625 |
| *clec-242* | 52.36 | 5.82 | -2.50 | 0.74 | 0.000686 | 0.0092376 |
| *dpy-3* | 2146.25 | 329.24 | -2.50 | 0.48 | 1.97E-07 | 1.33E-05 |
| *C30G12.1* | 62.02 | 5.73 | -2.50 | 0.80 | 0.00185 | 0.0197894 |
| *linc-61* | 38.64 | 4.06 | -2.50 | 0.76 | 0.000958 | 0.0119711 |
| *ZK742.3* | 157.10 | 21.67 | -2.51 | 0.60 | 2.87E-05 | 0.000749 |
| *cpg-9* | 25723.42 | 3747.53 | -2.51 | 0.54 | 3.96E-06 | 0.0001603 |
| *T06A4.3* | 1086.14 | 159.82 | -2.51 | 0.53 | 2.30E-06 | 0.0001021 |
| *F41E6.11* | 858.64 | 12.65 | -2.51 | 0.92 | 0.006125 | 0.0472898 |
| *col-110* | 3216.85 | 431.19 | -2.51 | 0.62 | 5.58E-05 | 0.0012805 |
| *msp-74* | 57.09 | 6.62 | -2.51 | 0.71 | 0.000426 | 0.0063619 |
| *dpy-8* | 3278.93 | 503.18 | -2.51 | 0.47 | 9.32E-08 | 7.12E-06 |
| *Y64H9A.2* | 249.71 | 36.62 | -2.52 | 0.53 | 1.85E-06 | 8.54E-05 |
| *dhs-26* | 53.38 | 6.29 | -2.52 | 0.70 | 0.00033 | 0.005249 |
| *clec-80* | 1748.64 | 263.18 | -2.54 | 0.47 | 7.28E-08 | 5.70E-06 |
| *col-79* | 2006.66 | 274.76 | -2.54 | 0.58 | 1.30E-05 | 0.0004016 |
| *wrt-6* | 6242.01 | 874.10 | -2.54 | 0.56 | 5.55E-06 | 0.0002113 |
| *ugt-12* | 864.51 | 130.42 | -2.54 | 0.46 | 4.41E-08 | 3.80E-06 |
| *cyn-6* | 1478.38 | 217.58 | -2.54 | 0.50 | 3.67E-07 | 2.26E-05 |
| *hmit-1.2* | 398.37 | 57.08 | -2.54 | 0.53 | 1.80E-06 | 8.41E-05 |
| *drd-10* | 14.75 | 0.41 | -2.54 | 0.90 | 0.00492 | 0.0405058 |
| *cpt-5* | 1118.49 | 136.64 | -2.55 | 0.67 | 0.000147 | 0.0027505 |
| *sec-61.B* | 3322.85 | 479.32 | -2.55 | 0.52 | 9.26E-07 | 4.91E-05 |
| *Y116F11B.2* | 35.38 | 3.30 | -2.55 | 0.78 | 0.001119 | 0.0135091 |
| *col-132* | 16.96 | 0.40 | -2.56 | 0.91 | 0.004771 | 0.0396257 |
| *Y50D7A.13* | 46.07 | 4.87 | -2.57 | 0.73 | 0.000481 | 0.0069645 |
| *sams-1* | 45228.06 | 6091.17 | -2.57 | 0.58 | 9.26E-06 | 0.0003119 |
| *T22D1.18* | 65.90 | 7.76 | -2.57 | 0.68 | 0.000156 | 0.0028879 |
| *T13C5.7* | 36.60 | 3.30 | -2.58 | 0.79 | 0.001062 | 0.0130301 |

| | | | | | | |
|---|---|---|---|---|---|---|
| *pgp-5* | 311.96 | 38.93 | -2.58 | 0.63 | 4.74E-05 | 0.0011207 |
| *C10C5.4* | 581.96 | 73.01 | -2.59 | 0.62 | 3.21E-05 | 0.0008221 |
| *hach-1* | 6900.19 | 986.02 | -2.59 | 0.49 | 1.01E-07 | 7.65E-06 |
| *F28A10.5* | 18.46 | 0.77 | -2.61 | 0.89 | 0.003337 | 0.030589 |
| *F53H8.3* | 620.86 | 81.05 | -2.61 | 0.58 | 5.84E-06 | 0.0002192 |
| *mltn-9* | 1353.45 | 183.15 | -2.61 | 0.53 | 1.00E-06 | 5.17E-05 |
| *hmit-1.1* | 4853.73 | 660.65 | -2.61 | 0.53 | 6.96E-07 | 3.88E-05 |
| *mam-3* | 300.40 | 41.74 | -2.62 | 0.50 | 1.65E-07 | 1.13E-05 |
| *sec-61.A* | 77748.29 | 10945.90 | -2.62 | 0.48 | 5.98E-08 | 4.83E-06 |
| *W03D8.11* | 85.08 | 9.80 | -2.62 | 0.67 | 8.74E-05 | 0.001825 |
| *Y43F8C.13* | 1909.78 | 266.03 | -2.62 | 0.49 | 1.15E-07 | 8.39E-06 |
| *F48G7.5* | 159.39 | 20.32 | -2.62 | 0.59 | 9.06E-06 | 0.0003079 |
| *dsc-4* | 5783.83 | 822.11 | -2.62 | 0.46 | 1.22E-08 | 1.23E-06 |
| *gcsh-1* | 1724.38 | 213.24 | -2.63 | 0.61 | 1.77E-05 | 0.0005075 |
| *R01E6.5* | 292.37 | 37.03 | -2.63 | 0.59 | 7.92E-06 | 0.0002754 |
| *gst-39* | 914.74 | 114.19 | -2.63 | 0.60 | 1.15E-05 | 0.0003651 |
| *ora-1* | 146.09 | 13.90 | -2.63 | 0.76 | 0.0005 | 0.0071547 |
| *aat-4* | 343.24 | 45.22 | -2.63 | 0.55 | 1.42E-06 | 6.98E-05 |
| *col-161* | 20294.52 | 2754.63 | -2.63 | 0.51 | 2.68E-07 | 1.72E-05 |
| *Y39D8A.1* | 704.91 | 93.42 | -2.64 | 0.54 | 9.38E-07 | 4.94E-05 |
| *fat-7* | 3187.93 | 415.35 | -2.64 | 0.55 | 1.78E-06 | 8.34E-05 |
| *alh-5* | 511.29 | 66.28 | -2.64 | 0.56 | 2.15E-06 | 9.67E-05 |
| *M04C9.4* | 221.26 | 26.36 | -2.65 | 0.63 | 2.38E-05 | 0.0006358 |
| *sym-1* | 5444.25 | 755.99 | -2.65 | 0.46 | 8.87E-09 | 9.27E-07 |
| *Y60A3A.23* | 50.59 | 5.20 | -2.66 | 0.71 | 0.000179 | 0.0032469 |
| *C02E7.6* | 31135.51 | 4100.00 | -2.66 | 0.53 | 4.21E-07 | 2.51E-05 |
| *pqn-36* | 346.88 | 44.53 | -2.66 | 0.55 | 1.46E-06 | 7.10E-05 |
| *F25C8.1* | 18.94 | 0.80 | -2.67 | 0.88 | 0.002468 | 0.0245873 |
| *Y69A2AR.3* | 2121.85 | 283.60 | -2.67 | 0.50 | 8.99E-08 | 6.93E-06 |
| *ZC374.2* | 730.64 | 100.09 | -2.67 | 0.46 | 8.96E-09 | 9.31E-07 |
| *F35D2.2* | 57.24 | 6.00 | -2.67 | 0.69 | 0.000115 | 0.0022687 |
| *Y46G5A.29* | 1193.52 | 157.25 | -2.68 | 0.51 | 1.30E-07 | 9.33E-06 |
| *cdh-12* | 2879.42 | 375.49 | -2.68 | 0.52 | 2.57E-07 | 1.69E-05 |
| *C35B1.7* | 702.94 | 87.40 | -2.68 | 0.57 | 2.43E-06 | 0.0001074 |
| *Y41C4A.32* | 1408.49 | 152.69 | -2.68 | 0.67 | 6.94E-05 | 0.0014999 |
| *slc-36.4* | 488.65 | 58.31 | -2.68 | 0.60 | 8.44E-06 | 0.0002916 |
| *elo-5* | 9502.17 | 1242.32 | -2.68 | 0.51 | 1.47E-07 | 1.04E-05 |
| *F18C5.5* | 893.73 | 114.52 | -2.69 | 0.53 | 3.24E-07 | 2.04E-05 |
| *C35C5.10* | 1800.15 | 241.92 | -2.69 | 0.46 | 6.53E-09 | 7.12E-07 |
| *srx-58* | 165.37 | 20.59 | -2.70 | 0.55 | 9.73E-07 | 5.05E-05 |
| *grd-11* | 40.30 | 3.30 | -2.70 | 0.78 | 0.000545 | 0.0076127 |
| *nhr-114* | 3730.94 | 500.11 | -2.70 | 0.46 | 5.04E-09 | 5.74E-07 |
| *cht-4* | 924.06 | 105.94 | -2.70 | 0.62 | 1.53E-05 | 0.0004545 |
| *M195.2* | 362.00 | 43.83 | -2.70 | 0.58 | 2.91E-06 | 0.0001253 |
| *C33B4.2* | 52.44 | 4.96 | -2.70 | 0.73 | 0.000211 | 0.0036835 |
| *C54G4.4* | 112.44 | 11.95 | -2.70 | 0.67 | 5.90E-05 | 0.0013228 |
| *atic-1* | 3359.83 | 450.44 | -2.70 | 0.46 | 3.44E-09 | 4.04E-07 |
| *col-133* | 20745.07 | 2670.68 | -2.71 | 0.51 | 9.23E-08 | 7.08E-06 |
| *col-149* | 5936.48 | 734.03 | -2.71 | 0.55 | 7.43E-07 | 4.09E-05 |
| *Y37A1B.7* | 337.30 | 42.56 | -2.72 | 0.52 | 1.51E-07 | 1.05E-05 |
| *trap-1* | 39324.26 | 5093.91 | -2.72 | 0.49 | 2.04E-08 | 1.95E-06 |
| *ifp-1* | 3965.25 | 526.58 | -2.73 | 0.45 | 1.04E-09 | 1.37E-07 |
| *C30A5.10* | 50.36 | 3.17 | -2.73 | 0.84 | 0.001112 | 0.0134629 |
| *irg-3* | 1559.57 | 190.29 | -2.73 | 0.55 | 6.68E-07 | 3.73E-05 |
| *T13C5.9* | 67.99 | 6.52 | -2.73 | 0.71 | 0.000132 | 0.0025191 |
| *F21E9.2* | 17.52 | 0.40 | -2.73 | 0.90 | 0.002424 | 0.0242689 |

| | | | | | | |
|---|---|---|---|---|---|---|
| *T15B7.1* | 3398.79 | 418.52 | -2.74 | 0.53 | 2.66E-07 | 1.72E-05 |
| *mltn-13* | 106.58 | 11.65 | -2.74 | 0.63 | 1.45E-05 | 0.0004381 |
| *clec-246* | 19.39 | 0.80 | -2.74 | 0.87 | 0.001687 | 0.018429 |
| *ZK180.6* | 20901.05 | 2725.70 | -2.75 | 0.45 | 7.90E-10 | 1.08E-07 |
| *B0280.7* | 60.03 | 3.54 | -2.75 | 0.84 | 0.00111 | 0.0134629 |
| *abhd-3.2* | 935.80 | 104.93 | -2.75 | 0.61 | 6.56E-06 | 0.0002401 |
| *T24A6.20* | 948.27 | 114.76 | -2.75 | 0.54 | 3.19E-07 | 2.01E-05 |
| *E02C12.6* | 23.13 | 1.19 | -2.75 | 0.85 | 0.001278 | 0.0149895 |
| *F32B4.8* | 68.37 | 6.85 | -2.77 | 0.67 | 4.12E-05 | 0.0010104 |
| *oac-42* | 39.86 | 3.17 | -2.77 | 0.77 | 0.000333 | 0.0052827 |
| *wrt-10* | 2387.98 | 307.08 | -2.77 | 0.45 | 8.44E-10 | 1.14E-07 |
| *pks-1* | 368.16 | 45.57 | -2.77 | 0.50 | 3.42E-08 | 3.07E-06 |
| *Y27F2A.9* | 13.96 | 0.15 | -2.78 | 0.91 | 0.002371 | 0.0240057 |
| *C38C6.3* | 304.32 | 36.72 | -2.78 | 0.52 | 1.06E-07 | 7.96E-06 |
| *Y39B6A.9* | 255.62 | 27.49 | -2.78 | 0.63 | 8.80E-06 | 0.0003019 |
| *F01G10.9* | 744.53 | 90.75 | -2.79 | 0.50 | 2.53E-08 | 2.35E-06 |
| *C35A5.11* | 56.31 | 5.24 | -2.79 | 0.70 | 7.03E-05 | 0.0015138 |
| *F53B3.3* | 79.57 | 7.89 | -2.79 | 0.67 | 3.04E-05 | 0.0007842 |
| *F48G7.8* | 33.83 | 2.41 | -2.80 | 0.79 | 0.000415 | 0.0062747 |
| *T24C12.4* | 76.17 | 7.04 | -2.80 | 0.71 | 7.26E-05 | 0.0015557 |
| *ugt-65* | 280.08 | 29.63 | -2.80 | 0.62 | 7.09E-06 | 0.0002529 |
| *ugt-47* | 3314.78 | 411.71 | -2.81 | 0.45 | 5.61E-10 | 8.01E-08 |
| *twnk-1* | 35.04 | 2.33 | -2.81 | 0.81 | 0.000483 | 0.006975 |
| *pho-1* | 2707.29 | 321.34 | -2.81 | 0.51 | 3.37E-08 | 3.06E-06 |
| *C34E11.2* | 228.88 | 23.39 | -2.82 | 0.64 | 1.01E-05 | 0.0003314 |
| *Y48G8AL.12* | 13061.58 | 1517.75 | -2.83 | 0.52 | 4.61E-08 | 3.95E-06 |
| *C01F1.5* | 506.52 | 56.54 | -2.84 | 0.56 | 3.41E-07 | 2.12E-05 |
| *ltah-1.2* | 6361.39 | 703.46 | -2.84 | 0.56 | 4.12E-07 | 2.47E-05 |
| *ZK6.11* | 36688.87 | 4351.32 | -2.84 | 0.48 | 4.21E-09 | 4.83E-07 |
| *F13A7.12* | 114.49 | 12.11 | -2.84 | 0.60 | 1.85E-06 | 8.54E-05 |
| *sdz-24* | 9493.24 | 1101.49 | -2.84 | 0.51 | 2.48E-08 | 2.32E-06 |
| *T13C5.2* | 84.81 | 8.17 | -2.85 | 0.66 | 1.61E-05 | 0.0004712 |
| *R05A10.6* | 39.47 | 2.49 | -2.85 | 0.81 | 0.000413 | 0.0062638 |
| *ilys-5* | 27576.47 | 2917.39 | -2.86 | 0.59 | 1.33E-06 | 6.65E-05 |
| *lgc-27* | 168.43 | 17.90 | -2.86 | 0.58 | 9.43E-07 | 4.95E-05 |
| *C09B8.3* | 21.93 | 0.41 | -2.86 | 0.90 | 0.001469 | 0.0165989 |
| *ugt-21* | 318.56 | 33.16 | -2.86 | 0.60 | 1.86E-06 | 8.54E-05 |
| *lec-10* | 29557.79 | 3336.81 | -2.86 | 0.52 | 4.19E-08 | 3.68E-06 |
| *glf-1* | 1301.38 | 152.03 | -2.87 | 0.48 | 2.06E-09 | 2.56E-07 |
| *msra-1* | 4502.27 | 521.28 | -2.87 | 0.49 | 3.48E-09 | 4.06E-07 |
| *lin-42* | 18481.67 | 2216.64 | -2.88 | 0.43 | 3.04E-11 | 6.10E-09 |
| *nspb-4* | 32.59 | 2.02 | -2.89 | 0.80 | 0.000296 | 0.004823 |
| *lact-7* | 86.07 | 7.99 | -2.90 | 0.66 | 1.04E-05 | 0.0003406 |
| *acs-1* | 5326.31 | 620.72 | -2.90 | 0.45 | 1.82E-10 | 2.86E-08 |
| *gly-1* | 132.97 | 12.44 | -2.91 | 0.65 | 7.30E-06 | 0.0002596 |
| *slc-17.8* | 269.95 | 25.74 | -2.91 | 0.64 | 4.98E-06 | 0.0001931 |
| *vit-1* | 30340.00 | 3410.15 | -2.91 | 0.49 | 1.95E-09 | 2.44E-07 |
| *F21C10.9* | 1477.34 | 148.96 | -2.92 | 0.59 | 7.93E-07 | 4.31E-05 |
| *D1054.8* | 922.96 | 98.69 | -2.92 | 0.54 | 5.43E-08 | 4.46E-06 |
| *spds-1* | 1450.12 | 159.84 | -2.92 | 0.50 | 6.28E-09 | 6.98E-07 |
| *trap-2* | 25172.02 | 2727.65 | -2.93 | 0.51 | 1.27E-08 | 1.27E-06 |
| *str-7* | 27.32 | 1.22 | -2.94 | 0.84 | 0.000494 | 0.0071068 |
| *C35A5.3* | 238.36 | 24.64 | -2.95 | 0.55 | 6.44E-08 | 5.15E-06 |
| *lpr-4* | 3040.77 | 338.95 | -2.95 | 0.46 | 1.15E-10 | 1.98E-08 |
| *gst-28* | 1070.91 | 109.54 | -2.97 | 0.54 | 4.90E-08 | 4.13E-06 |
| *pho-11* | 43113.48 | 4247.51 | -2.97 | 0.58 | 2.65E-07 | 1.72E-05 |

| | | | | | | |
|---|---|---|---|---|---|---|
| *clec-258* | 39.47 | 2.10 | -2.97 | 0.81 | 0.00026 | 0.0043472 |
| *C04G6.10* | 18.70 | 0.15 | -2.98 | 0.91 | 0.001091 | 0.0133012 |
| *col-152* | 388.02 | 40.62 | -2.98 | 0.51 | 4.05E-09 | 4.68E-07 |
| *ZC266.1* | 170.02 | 16.24 | -2.99 | 0.59 | 3.34E-07 | 2.09E-05 |
| *Y16B4A.2* | 19831.26 | 2157.79 | -2.99 | 0.45 | 4.12E-11 | 8.16E-09 |
| *F35G2.5* | 151.76 | 13.88 | -3.01 | 0.61 | 7.57E-07 | 4.14E-05 |
| *C54F6.3* | 128.05 | 11.86 | -3.02 | 0.59 | 3.59E-07 | 2.22E-05 |
| *aqp-4* | 1590.35 | 159.35 | -3.02 | 0.52 | 7.29E-09 | 7.78E-07 |
| *phat-1* | 97.75 | 5.31 | -3.02 | 0.81 | 0.000188 | 0.003381 |
| *ZK1307.1* | 6978.34 | 743.14 | -3.02 | 0.45 | 1.65E-11 | 3.59E-09 |
| *R05A10.7* | 51.46 | 3.55 | -3.02 | 0.74 | 4.26E-05 | 0.0010394 |
| *D2024.4* | 33.96 | 0.77 | -3.04 | 0.89 | 0.000626 | 0.0085618 |
| *M03B6.1* | 186.39 | 14.69 | -3.04 | 0.69 | 9.48E-06 | 0.0003187 |
| *F11A5.5* | 116.23 | 10.22 | -3.04 | 0.61 | 7.19E-07 | 3.99E-05 |
| *klo-2* | 344.69 | 32.34 | -3.05 | 0.56 | 6.79E-08 | 5.37E-06 |
| *cln-3.3* | 549.00 | 52.81 | -3.05 | 0.54 | 1.84E-08 | 1.78E-06 |
| *grd-6* | 7223.44 | 682.12 | -3.06 | 0.55 | 2.99E-08 | 2.73E-06 |
| *ugt-22* | 1600.29 | 162.36 | -3.06 | 0.48 | 1.17E-10 | 1.98E-08 |
| *drd-5* | 1563.54 | 154.27 | -3.07 | 0.50 | 1.04E-09 | 1.37E-07 |
| *F26D11.2* | 72.65 | 5.20 | -3.08 | 0.71 | 1.45E-05 | 0.0004381 |
| *srd-7* | 35.89 | 1.57 | -3.08 | 0.83 | 0.000204 | 0.0035964 |
| *mboa-1* | 449.11 | 44.37 | -3.08 | 0.49 | 3.02E-10 | 4.61E-08 |
| *F07H5.13* | 141.44 | 12.61 | -3.08 | 0.58 | 1.29E-07 | 9.32E-06 |
| *C50F7.9* | 38.94 | 2.06 | -3.09 | 0.79 | 9.33E-05 | 0.0019176 |
| *clec-5* | 2715.98 | 245.45 | -3.09 | 0.57 | 5.16E-08 | 4.28E-06 |
| *F25D1.5* | 132.29 | 10.95 | -3.09 | 0.63 | 8.25E-07 | 4.40E-05 |
| *dct-18* | 21302.07 | 2056.13 | -3.10 | 0.50 | 6.09E-10 | 8.60E-08 |
| *R07B1.9* | 170.78 | 14.31 | -3.10 | 0.62 | 5.02E-07 | 2.92E-05 |
| *nspb-1* | 66.74 | 4.86 | -3.11 | 0.69 | 5.89E-06 | 0.00022 |
| *col-174* | 776.94 | 67.31 | -3.11 | 0.59 | 1.25E-07 | 9.02E-06 |
| *cut-3* | 68.30 | 4.40 | -3.13 | 0.73 | 1.94E-05 | 0.000544 |
| *Y43D4A.5* | 894.08 | 82.75 | -3.13 | 0.52 | 1.67E-09 | 2.10E-07 |
| *K10D11.5* | 609.49 | 56.47 | -3.13 | 0.52 | 1.37E-09 | 1.76E-07 |
| *Y47D7A.15* | 10537.20 | 914.59 | -3.13 | 0.57 | 4.86E-08 | 4.13E-06 |
| *T10E10.4* | 144.55 | 10.67 | -3.14 | 0.67 | 3.16E-06 | 0.000134 |
| *T23F6.5* | 182.83 | 16.13 | -3.14 | 0.55 | 1.30E-08 | 1.30E-06 |
| *clec-239* | 48.20 | 2.72 | -3.14 | 0.77 | 4.20E-05 | 0.0010299 |
| *C18E9.7* | 215.64 | 16.55 | -3.15 | 0.65 | 1.29E-06 | 6.47E-05 |
| *col-120* | 6214.98 | 581.91 | -3.15 | 0.49 | 1.43E-10 | 2.30E-08 |
| *mlt-10* | 5641.30 | 491.49 | -3.16 | 0.55 | 1.11E-08 | 1.14E-06 |
| *F17B5.8* | 121.25 | 8.87 | -3.17 | 0.66 | 1.71E-06 | 8.16E-05 |
| *lon-3* | 4268.22 | 398.84 | -3.18 | 0.47 | 8.56E-12 | 1.94E-09 |
| *pgp-13* | 354.30 | 30.39 | -3.19 | 0.55 | 5.98E-09 | 6.71E-07 |
| *folt-2* | 3916.49 | 282.85 | -3.21 | 0.66 | 9.91E-07 | 5.13E-05 |
| *cest-10* | 403.93 | 34.30 | -3.21 | 0.54 | 2.53E-09 | 3.04E-07 |
| *col-47* | 83.32 | 5.71 | -3.21 | 0.67 | 1.93E-06 | 8.83E-05 |
| *cyp-14A2* | 137.86 | 10.64 | -3.22 | 0.61 | 1.13E-07 | 8.27E-06 |
| *F38B6.4* | 2287.46 | 209.97 | -3.22 | 0.45 | 9.90E-13 | 2.92E-10 |
| *gst-26* | 2552.20 | 214.38 | -3.23 | 0.54 | 2.11E-09 | 2.59E-07 |
| *T22B2.6* | 60.62 | 2.71 | -3.23 | 0.80 | 5.80E-05 | 0.0013102 |
| *gst-4* | 698.46 | 61.29 | -3.23 | 0.49 | 5.16E-11 | 1.01E-08 |
| *phg-1* | 200.90 | 12.30 | -3.24 | 0.72 | 6.69E-06 | 0.0002425 |
| *lips-6* | 39.34 | 1.54 | -3.24 | 0.82 | 7.33E-05 | 0.0015687 |
| *Y102A11A.7* | 70.46 | 4.27 | -3.25 | 0.72 | 5.57E-06 | 0.0002113 |
| *twk-11* | 46.07 | 2.02 | -3.25 | 0.80 | 4.41E-05 | 0.0010633 |
| *K12B6.11* | 142.33 | 9.51 | -3.26 | 0.67 | 1.38E-06 | 6.85E-05 |

| | | | | | | |
|---|---|---|---|---|---|---|
| *F07C6.6* | 18.94 | 0.15 | -3.26 | 0.90 | 0.000307 | 0.0049722 |
| *dod-17* | 459.09 | 37.37 | -3.26 | 0.54 | 2.12E-09 | 2.59E-07 |
| *C32H11.6* | 30.20 | 0.40 | -3.28 | 0.89 | 0.000223 | 0.0038539 |
| *F31B9.4* | 46.40 | 2.02 | -3.28 | 0.79 | 3.53E-05 | 0.0008888 |
| *K02E11.10* | 8231.23 | 692.11 | -3.31 | 0.48 | 5.10E-12 | 1.24E-09 |
| *ZK218.1* | 38.02 | 1.58 | -3.31 | 0.79 | 3.16E-05 | 0.0008137 |
| *cest-28* | 47.97 | 2.41 | -3.31 | 0.75 | 1.10E-05 | 0.0003563 |
| *cutl-7* | 24.85 | 0.40 | -3.32 | 0.87 | 0.000149 | 0.0027778 |
| *fkb-5* | 3158.61 | 246.81 | -3.32 | 0.54 | 9.25E-10 | 1.24E-07 |
| *C04G6.13* | 24.21 | 0.40 | -3.33 | 0.87 | 0.000136 | 0.0025768 |
| *acdh-1* | 43863.42 | 3668.99 | -3.33 | 0.46 | 5.41E-13 | 1.65E-10 |
| *C48D5.3* | 28.58 | 0.40 | -3.34 | 0.88 | 0.000149 | 0.0027754 |
| *C34F6.1* | 254.12 | 13.29 | -3.34 | 0.74 | 6.92E-06 | 0.0002481 |
| *nspb-7* | 39.91 | 1.25 | -3.34 | 0.83 | 5.47E-05 | 0.0012579 |
| *C05C9.1* | 48.97 | 1.22 | -3.35 | 0.86 | 9.15E-05 | 0.001897 |
| *abu-4* | 46.93 | 1.92 | -3.35 | 0.79 | 2.30E-05 | 0.0006188 |
| *F38B6.2* | 37.16 | 0.41 | -3.37 | 0.89 | 0.000149 | 0.0027754 |
| *vit-4* | 23441.24 | 1438.42 | -3.38 | 0.67 | 4.45E-07 | 2.64E-05 |
| *dao-4* | 1726.24 | 138.11 | -3.39 | 0.47 | 3.93E-13 | 1.25E-10 |
| *R06C1.4* | 21997.01 | 1710.86 | -3.39 | 0.50 | 8.62E-12 | 1.94E-09 |
| *hrg-7* | 10009.61 | 768.97 | -3.39 | 0.51 | 2.06E-11 | 4.30E-09 |
| *ifc-1* | 3354.00 | 212.52 | -3.41 | 0.64 | 1.00E-07 | 7.63E-06 |
| *ZK662.2* | 384.85 | 23.07 | -3.42 | 0.66 | 2.61E-07 | 1.70E-05 |
| *asp-8* | 4829.34 | 320.49 | -3.42 | 0.61 | 1.78E-08 | 1.73E-06 |
| *F22E5.1* | 397.20 | 28.43 | -3.45 | 0.53 | 6.83E-11 | 1.26E-08 |
| *T06E4.12* | 1043.04 | 66.92 | -3.45 | 0.61 | 1.53E-08 | 1.50E-06 |
| *Y48E1B.8* | 492.22 | 28.12 | -3.47 | 0.67 | 2.09E-07 | 1.39E-05 |
| *ugt-53* | 325.22 | 22.61 | -3.47 | 0.54 | 1.24E-10 | 2.07E-08 |
| *lrx-1* | 85.64 | 3.97 | -3.48 | 0.74 | 2.25E-06 | 0.0001008 |
| *col-137* | 1543.41 | 104.15 | -3.48 | 0.56 | 4.99E-10 | 7.18E-08 |
| *T24F1.7* | 94.56 | 4.02 | -3.48 | 0.76 | 4.44E-06 | 0.0001757 |
| *elo-6* | 14477.32 | 1006.50 | -3.49 | 0.53 | 5.22E-11 | 1.01E-08 |
| *plpr-1* | 382.89 | 22.58 | -3.49 | 0.64 | 5.08E-08 | 4.26E-06 |
| *nas-14* | 153.31 | 6.11 | -3.49 | 0.78 | 6.60E-06 | 0.0002406 |
| *mfsd-11* | 316.09 | 21.38 | -3.49 | 0.55 | 1.80E-10 | 2.86E-08 |
| *R11G11.6* | 40.12 | 0.83 | -3.50 | 0.85 | 3.61E-05 | 0.0009079 |
| *K01D12.1* | 45.15 | 1.63 | -3.51 | 0.78 | 6.50E-06 | 0.0002391 |
| *abu-10* | 633.89 | 34.90 | -3.53 | 0.66 | 8.00E-08 | 6.20E-06 |
| *pqn-26* | 426.87 | 25.39 | -3.54 | 0.61 | 8.31E-09 | 8.81E-07 |
| *ech-1.1* | 532.87 | 35.77 | -3.55 | 0.52 | 5.40E-12 | 1.29E-09 |
| *lron-2* | 325.38 | 21.01 | -3.56 | 0.55 | 7.30E-11 | 1.30E-08 |
| *acox-1.4* | 1728.80 | 117.74 | -3.56 | 0.50 | 1.12E-12 | 3.23E-10 |
| *F26F12.4* | 72.22 | 3.14 | -3.57 | 0.73 | 9.60E-07 | 5.02E-05 |
| *C04G6.2* | 874.40 | 60.91 | -3.57 | 0.47 | 1.92E-14 | 7.72E-12 |
| *F26F12.8* | 268.36 | 17.00 | -3.58 | 0.55 | 6.98E-11 | 1.27E-08 |
| *ech-6* | 24168.91 | 1600.67 | -3.59 | 0.50 | 9.96E-13 | 2.92E-10 |
| *ZK470.6* | 83.63 | 1.95 | -3.61 | 0.84 | 1.65E-05 | 0.0004805 |
| *dhs-25* | 6707.54 | 411.87 | -3.61 | 0.55 | 6.48E-11 | 1.21E-08 |
| *act-5* | 80709.57 | 4922.08 | -3.62 | 0.56 | 8.39E-11 | 1.46E-08 |
| *C46H11.7* | 84.70 | 2.71 | -3.62 | 0.79 | 4.78E-06 | 0.0001868 |
| *F26C11.1* | 368.38 | 18.32 | -3.65 | 0.66 | 3.39E-08 | 3.07E-06 |
| *F10A3.4* | 4853.16 | 287.49 | -3.65 | 0.56 | 7.30E-11 | 1.30E-08 |
| *wrt-4* | 5389.89 | 325.14 | -3.66 | 0.54 | 8.37E-12 | 1.94E-09 |
| *pqn-29* | 56.70 | 0.78 | -3.70 | 0.86 | 1.74E-05 | 0.0005025 |
| *T05E12.6* | 9429.09 | 515.11 | -3.71 | 0.58 | 2.10E-10 | 3.27E-08 |
| *gst-10* | 5550.57 | 335.34 | -3.72 | 0.50 | 8.19E-14 | 2.91E-11 |

| | | | | | | |
|---|---|---|---|---|---|---|
| *grl-27* | 857.64 | 43.17 | -3.75 | 0.61 | 1.06E-09 | 1.39E-07 |
| *lgx-1* | 689.16 | 32.50 | -3.75 | 0.65 | 6.36E-09 | 6.98E-07 |
| *R08E5.3* | 3017.63 | 152.40 | -3.77 | 0.60 | 3.19E-10 | 4.82E-08 |
| *F20B10.3* | 100.86 | 3.86 | -3.78 | 0.71 | 1.06E-07 | 7.95E-06 |
| *clec-210* | 116.70 | 4.60 | -3.79 | 0.69 | 4.75E-08 | 4.05E-06 |
| *nas-1* | 70.55 | 2.31 | -3.80 | 0.75 | 3.38E-07 | 2.11E-05 |
| *F41G3.3* | 87.29 | 1.61 | -3.80 | 0.83 | 5.26E-06 | 0.0002019 |
| *col-148* | 36.09 | 0.40 | -3.81 | 0.85 | 7.42E-06 | 0.000262 |
| *F12E12.11* | 128.40 | 4.60 | -3.81 | 0.73 | 1.49E-07 | 1.04E-05 |
| *abu-1* | 845.01 | 35.13 | -3.81 | 0.68 | 2.14E-08 | 2.01E-06 |
| *Y47D3B.6* | 3954.77 | 224.39 | -3.81 | 0.49 | 8.34E-15 | 3.64E-12 |
| *dct-16* | 42765.74 | 2271.62 | -3.82 | 0.54 | 1.53E-12 | 4.23E-10 |
| *pgp-12* | 150.85 | 6.54 | -3.84 | 0.65 | 2.88E-09 | 3.44E-07 |
| *E01G6.1* | 326.41 | 13.77 | -3.85 | 0.66 | 5.11E-09 | 5.78E-07 |
| *pmp-5* | 5768.37 | 328.90 | -3.85 | 0.46 | 4.72E-17 | 3.27E-14 |
| *M02G9.2* | 212.46 | 8.93 | -3.89 | 0.64 | 1.35E-09 | 1.75E-07 |
| *pqn-63* | 1222.13 | 57.09 | -3.91 | 0.58 | 1.84E-11 | 3.96E-09 |
| *Y75B7AR.1* | 517.27 | 16.73 | -3.96 | 0.72 | 4.10E-08 | 3.62E-06 |
| *nep-22* | 8907.40 | 444.59 | -3.98 | 0.49 | 6.69E-16 | 3.52E-13 |
| *pqn-54* | 1776.98 | 66.81 | -4.02 | 0.65 | 6.33E-10 | 8.86E-08 |
| *nspe-2* | 58.13 | 0.40 | -4.03 | 0.86 | 2.86E-06 | 0.0001233 |
| *F35A5.4* | 188.18 | 5.06 | -4.05 | 0.75 | 5.82E-08 | 4.73E-06 |
| *Y71G12B.18* | 597.15 | 22.90 | -4.05 | 0.63 | 1.19E-10 | 1.99E-08 |
| *F59B10.3* | 184.47 | 6.54 | -4.08 | 0.65 | 3.48E-10 | 5.15E-08 |
| *C23H3.9* | 512.21 | 15.76 | -4.08 | 0.70 | 6.83E-09 | 7.39E-07 |
| *nspb-2* | 40.80 | 0.40 | -4.10 | 0.83 | 7.39E-07 | 4.09E-05 |
| *srh-237* | 294.10 | 11.67 | -4.10 | 0.59 | 2.68E-12 | 6.94E-10 |
| *dhs-23* | 84.69 | 1.96 | -4.17 | 0.74 | 1.89E-08 | 1.81E-06 |
| *F54B11.11* | 1278.02 | 56.53 | -4.17 | 0.48 | 1.82E-18 | 1.85E-15 |
| *hphd-1* | 18573.58 | 776.40 | -4.17 | 0.52 | 1.09E-15 | 5.37E-13 |
| *nspb-3* | 33.51 | 0.15 | -4.17 | 0.87 | 1.43E-06 | 6.98E-05 |
| *D2096.6* | 1411.14 | 58.33 | -4.19 | 0.52 | 5.49E-16 | 3.10E-13 |
| *C54D2.1* | 353.74 | 12.62 | -4.19 | 0.61 | 4.35E-12 | 1.07E-09 |
| *nhr-235* | 65.45 | 1.21 | -4.21 | 0.77 | 4.25E-08 | 3.71E-06 |
| *M03E7.4* | 144.24 | 3.55 | -4.21 | 0.73 | 6.36E-09 | 6.98E-07 |
| *Y48G8AL.16* | 216.48 | 7.99 | -4.22 | 0.57 | 1.18E-13 | 3.99E-11 |
| *F49C5.11* | 384.70 | 15.37 | -4.23 | 0.52 | 4.08E-16 | 2.40E-13 |
| *T06E4.14* | 1272.49 | 43.44 | -4.24 | 0.61 | 3.93E-12 | 9.98E-10 |
| *Y43F8B.3* | 638.58 | 23.91 | -4.27 | 0.54 | 5.03E-15 | 2.26E-12 |
| *cth-1* | 2673.07 | 91.06 | -4.33 | 0.58 | 5.96E-14 | 2.27E-11 |
| *nhr-234* | 267.42 | 9.05 | -4.35 | 0.56 | 1.10E-14 | 4.66E-12 |
| *abu-14* | 3720.94 | 129.33 | -4.35 | 0.55 | 2.94E-15 | 1.36E-12 |
| *nas-15* | 483.02 | 9.76 | -4.43 | 0.73 | 1.52E-09 | 1.93E-07 |
| *str-168* | 105.68 | 2.38 | -4.45 | 0.68 | 7.62E-11 | 1.34E-08 |
| *F07H5.8* | 500.95 | 13.35 | -4.47 | 0.64 | 1.88E-12 | 5.03E-10 |
| *T06E4.8* | 1120.50 | 32.24 | -4.49 | 0.60 | 6.31E-14 | 2.35E-11 |
| *lbp-8* | 41.29 | 0.15 | -4.50 | 0.85 | 1.19E-07 | 8.67E-06 |
| *C09B8.4* | 293.90 | 7.79 | -4.51 | 0.62 | 3.09E-13 | 1.00E-10 |
| *Y39B6A.1* | 87698.09 | 2644.81 | -4.52 | 0.56 | 8.67E-16 | 4.41E-13 |
| *pho-13* | 954.66 | 28.62 | -4.57 | 0.54 | 1.41E-17 | 1.08E-14 |
| *F21G4.3* | 81.07 | 1.17 | -4.58 | 0.75 | 7.71E-10 | 1.07E-07 |
| *pqn-2* | 775.78 | 19.34 | -4.60 | 0.62 | 1.12E-13 | 3.88E-11 |
| *pqn-74* | 4984.22 | 141.33 | -4.62 | 0.55 | 4.71E-17 | 3.27E-14 |
| *acox-1.2* | 4766.40 | 148.46 | -4.62 | 0.48 | 1.43E-21 | 2.18E-18 |
| *nspb-8* | 46.54 | 0.15 | -4.68 | 0.84 | 2.69E-08 | 2.49E-06 |
| *R03C1.1* | 1226.20 | 33.01 | -4.69 | 0.55 | 9.05E-18 | 7.67E-15 |

| | | | | | | |
|---|---|---|---|---|---|---|
| *abu-11* | 2284.01 | 59.52 | -4.70 | 0.56 | 5.53E-17 | 3.67E-14 |
| *K01D12.8* | 131.38 | 2.35 | -4.75 | 0.68 | 2.07E-12 | 5.45E-10 |
| *R02F11.1* | 1338.82 | 28.33 | -4.78 | 0.63 | 2.59E-14 | 1.01E-11 |
| *nspb-10* | 51.17 | 0.15 | -4.81 | 0.84 | 8.63E-09 | 9.08E-07 |
| *T25E4.1* | 714.93 | 16.20 | -4.86 | 0.57 | 1.20E-17 | 9.67E-15 |
| *R12A1.3* | 456.98 | 9.03 | -4.90 | 0.61 | 1.55E-15 | 7.38E-13 |
| *F49C5.12* | 58.48 | 0.15 | -4.95 | 0.83 | 2.41E-09 | 2.92E-07 |
| *H17B01.2* | 293.71 | 4.43 | -5.16 | 0.63 | 1.61E-16 | 9.85E-14 |
| *D1014.7* | 780.23 | 14.47 | -5.21 | 0.53 | 3.47E-23 | 6.63E-20 |
| *C27D9.2* | 93.79 | 0.40 | -5.28 | 0.77 | 5.77E-12 | 1.36E-09 |
| *E03H4.4* | 77.02 | 0.15 | -5.41 | 0.81 | 1.96E-11 | 4.16E-09 |
| *C30G12.2* | 6228.96 | 82.44 | -5.50 | 0.59 | 7.52E-21 | 9.56E-18 |
| *T06E4.9* | 1019.63 | 11.33 | -5.52 | 0.64 | 6.45E-18 | 5.79E-15 |
| *abu-7* | 1251.13 | 15.72 | -5.60 | 0.57 | 1.17E-22 | 1.99E-19 |
| *acdh-2* | 236.06 | 1.18 | -6.02 | 0.69 | 2.02E-18 | 1.93E-15 |
| *ctl-1* | 231.58 | 0.40 | -6.44 | 0.73 | 9.52E-19 | 1.04E-15 |
| *D1014.6* | 814.29 | 4.43 | -6.62 | 0.57 | 2.87E-31 | 7.31E-28 |
| *BE0003N10.6* | 4110.84 | 21.00 | -7.04 | 0.47 | 7.02E-50 | 1.07E-45 |
| *argk-1* | 1049.17 | 3.23 | -7.15 | 0.60 | 1.38E-32 | 4.21E-29 |
