## supplementary table 2 for "Genome-wide screen identifies curli amyloid fibril as a bacterial component promoting host neurodegeneration"

| Gene | BaseMean of aex-3p::α-syn(A53T) fed with K12 WT | BaseMean of aex-3p::α-syn(A53T) fed with K12 csgA(-) | log2Fold Change | lfcSE | p value | adjusted p value |
|---|---|---|---|---|---|---|
| *srh-237* | 15.00 | 449.62 | 3.88 | 0.45 | 3.01E-18 | 2.83E-14 |
| *Y39B6A.1* | 3342.17 | 55720.64 | 3.45 | 0.40 | 3.70E-18 | 2.83E-14 |
| *F54B11.11* | 71.87 | 1276.26 | 3.47 | 0.41 | 4.22E-17 | 2.14E-13 |
| *tts-1* | 41714.28 | 3694.91 | -3.06 | 0.37 | 1.93E-16 | 7.36E-13 |
| *hphd-1* | 988.61 | 9152.52 | 2.85 | 0.35 | 9.58E-16 | 2.92E-12 |
| *tbb-6* | 2211.91 | 214.57 | -2.93 | 0.38 | 6.11E-15 | 1.55E-11 |
| *pmp-5* | 419.63 | 4597.26 | 2.98 | 0.38 | 8.03E-15 | 1.75E-11 |
| *cnc-4* | 742.66 | 58.94 | -3.06 | 0.41 | 1.12E-13 | 2.13E-10 |
| *pho-13* | 36.81 | 682.28 | 3.28 | 0.46 | 9.42E-13 | 1.60E-09 |
| *C50F7.5* | 4408.63 | 288.80 | -3.07 | 0.46 | 2.86E-11 | 4.37E-08 |
| *lips-6* | 2.00 | 111.40 | 3.48 | 0.53 | 3.69E-11 | 5.11E-08 |
| *acdh-1* | 4690.89 | 32287.58 | 2.43 | 0.37 | 7.08E-11 | 9.00E-08 |
| *ech-6* | 2030.90 | 11292.48 | 2.21 | 0.35 | 1.53E-10 | 1.80E-07 |
| *F10A3.4* | 359.89 | 2868.29 | 2.54 | 0.40 | 2.81E-10 | 3.07E-07 |
| *Y53F4B.8* | 152.65 | 5.22 | -3.21 | 0.52 | 5.32E-10 | 5.41E-07 |
| *MTCE.7* | 12113.70 | 1707.73 | -2.42 | 0.39 | 7.96E-10 | 7.59E-07 |
| *dod-17* | 47.96 | 444.80 | 2.63 | 0.43 | 1.04E-09 | 9.29E-07 |
| *T05E12.6* | 639.45 | 7350.71 | 2.77 | 0.46 | 1.17E-09 | 9.93E-07 |
| *T12D8.5* | 2308.75 | 279.70 | -2.51 | 0.42 | 2.94E-09 | 2.36E-06 |
| *asp-8* | 402.87 | 2764.60 | 2.34 | 0.41 | 8.07E-09 | 5.35E-06 |
| *nep-22* | 570.75 | 4419.04 | 2.44 | 0.42 | 7.96E-09 | 5.35E-06 |
| *clec-210* | 5.74 | 171.16 | 3.06 | 0.53 | 7.09E-09 | 5.35E-06 |
| *Y38H8A.8* | 125.41 | 6.69 | -2.92 | 0.51 | 7.45E-09 | 5.35E-06 |
| *glf-1* | 191.03 | 1140.03 | 2.21 | 0.39 | 1.25E-08 | 7.94E-06 |
| *mboa-1* | 56.11 | 376.69 | 2.31 | 0.41 | 1.80E-08 | 1.10E-05 |
| *lpr-4* | 429.73 | 2389.36 | 2.14 | 0.38 | 1.89E-08 | 1.11E-05 |
| *rpr-1* | 81.42 | 1.93 | -3.02 | 0.54 | 2.18E-08 | 1.23E-05 |
| *K10D11.5* | 72.44 | 539.39 | 2.38 | 0.43 | 2.27E-08 | 1.24E-05 |
| *elo-6* | 1298.26 | 10029.31 | 2.40 | 0.44 | 3.79E-08 | 2.00E-05 |
| *acox-1.4* | 151.33 | 1041.85 | 2.30 | 0.42 | 5.16E-08 | 2.63E-05 |
| *wrt-10* | 388.63 | 2100.85 | 2.10 | 0.39 | 6.02E-08 | 2.96E-05 |
| *T27A1.2* | 170.81 | 13.38 | -2.64 | 0.50 | 9.96E-08 | 4.75E-05 |
| *fbxa-163* | 163.43 | 9.11 | -2.75 | 0.52 | 1.22E-07 | 5.62E-05 |
| *T13F3.6* | 206.06 | 1043.06 | 2.02 | 0.38 | 1.43E-07 | 6.40E-05 |
| *cyp-13A4* | 1.50 | 73.63 | 2.88 | 0.55 | 1.57E-07 | 6.83E-05 |
| *lec-10* | 4235.51 | 17187.82 | 1.81 | 0.35 | 1.76E-07 | 7.46E-05 |
| *acox-1.2* | 190.54 | 1114.22 | 2.14 | 0.41 | 2.21E-07 | 8.87E-05 |
| *dhc-4* | 551.53 | 82.24 | -2.24 | 0.43 | 2.19E-07 | 8.87E-05 |
| *ZK6.11* | 5533.69 | 22919.97 | 1.82 | 0.35 | 2.44E-07 | 9.56E-05 |
| *alh-5* | 83.11 | 498.01 | 2.15 | 0.42 | 2.77E-07 | 0.0001058 |
| *Y39B6A.25* | 243.39 | 30.91 | -2.34 | 0.46 | 2.89E-07 | 0.0001075 |
| *F01G10.9* | 115.38 | 559.95 | 1.97 | 0.38 | 3.04E-07 | 0.0001103 |
| *msra-1* | 663.66 | 5670.85 | 2.39 | 0.47 | 3.59E-07 | 0.0001273 |
| *T23F6.5* | 20.41 | 173.38 | 2.37 | 0.47 | 3.73E-07 | 0.0001293 |
| *C30G12.2* | 106.58 | 983.97 | 2.42 | 0.48 | 4.36E-07 | 0.0001478 |
| *Y16B4A.2* | 2746.93 | 11925.67 | 1.86 | 0.37 | 4.56E-07 | 0.0001513 |
| *acs-1* | 793.73 | 3712.62 | 1.92 | 0.38 | 5.11E-07 | 0.0001658 |
| *R03G8.6* | 77.91 | 449.08 | 2.10 | 0.42 | 5.53E-07 | 0.0001757 |
| *M163.11* | 88.23 | 1.92 | -2.76 | 0.55 | 6.23E-07 | 0.0001939 |
| *C40H1.8* | 56.84 | 432.10 | 2.29 | 0.46 | 6.62E-07 | 0.0002019 |
| *ifp-1* | 671.58 | 2848.77 | 1.83 | 0.37 | 7.86E-07 | 0.0002305 |

| | | | | | | |
|---|---|---|---|---|---|---|
| *mtl-1* | 2037.27 | 194.75 | -2.45 | 0.50 | 7.81E-07 | 0.0002305 |
| *clx-1* | 77.33 | 3.80 | -2.63 | 0.54 | 9.01E-07 | 0.0002594 |
| *ilys-3* | 1131.70 | 204.06 | -2.05 | 0.42 | 9.45E-07 | 0.0002669 |
| *pgp-1* | 328.45 | 1403.14 | 1.83 | 0.37 | 1.07E-06 | 0.0002978 |
| *C17H12.6* | 58.61 | 361.52 | 2.13 | 0.44 | 1.14E-06 | 0.0003093 |
| *Y38A10A.2* | 105.54 | 501.59 | 1.92 | 0.39 | 1.16E-06 | 0.0003096 |
| *Y94H6A.2* | 8.99 | 105.39 | 2.45 | 0.51 | 1.50E-06 | 0.000394 |
| *T23F6.3* | 455.23 | 102.90 | -1.85 | 0.39 | 1.98E-06 | 0.000513 |
| *C17H12.8* | 1081.47 | 3965.66 | 1.67 | 0.35 | 2.06E-06 | 0.0005241 |
| *cyp-14A5* | 512.70 | 102.18 | -1.95 | 0.41 | 2.36E-06 | 0.0005914 |
| *R03H10.6* | 245.41 | 31.33 | -2.25 | 0.48 | 2.41E-06 | 0.0005922 |
| *drd-5* | 194.55 | 805.67 | 1.78 | 0.38 | 2.65E-06 | 0.0006406 |
| *F55C10.4* | 10.71 | 102.18 | 2.33 | 0.50 | 2.74E-06 | 0.000652 |
| *C42D4.3* | 9552.66 | 2175.96 | -1.83 | 0.39 | 3.17E-06 | 0.000744 |
| *dao-4* | 174.56 | 769.23 | 1.83 | 0.39 | 3.34E-06 | 0.0007717 |
| *act-5* | 6330.10 | 30187.04 | 1.89 | 0.41 | 3.88E-06 | 0.0008829 |
| *clec-80* | 334.51 | 1247.19 | 1.67 | 0.36 | 4.04E-06 | 0.0009067 |
| *Y46G5A.38* | 25.45 | 188.78 | 2.19 | 0.48 | 4.31E-06 | 0.0009535 |
| *Y48A6B.8* | 165.88 | 14.89 | -2.37 | 0.52 | 4.51E-06 | 0.0009748 |
| *clec-218* | 182.25 | 925.49 | 1.94 | 0.42 | 4.60E-06 | 0.0009748 |
| *Y58A7A.3* | 895.03 | 205.75 | -1.81 | 0.40 | 4.55E-06 | 0.0009748 |
| *pgp-13* | 38.97 | 261.69 | 2.13 | 0.47 | 4.74E-06 | 0.0009905 |
| *cln-3.3* | 67.82 | 389.38 | 2.02 | 0.45 | 5.71E-06 | 0.0011613 |
| *F47B8.5* | 17.43 | 131.11 | 2.18 | 0.48 | 5.66E-06 | 0.0011613 |
| *idh-1* | 2740.69 | 9677.56 | 1.61 | 0.36 | 6.42E-06 | 0.0012895 |
| *grd-1* | 153.11 | 612.50 | 1.73 | 0.38 | 7.21E-06 | 0.0014097 |
| *F20D1.3* | 11069.65 | 2914.83 | -1.68 | 0.37 | 7.18E-06 | 0.0014097 |
| *tos-1* | 20194.97 | 3952.56 | -1.93 | 0.43 | 7.41E-06 | 0.0014302 |
| *nas-37* | 400.32 | 1452.95 | 1.64 | 0.37 | 7.55E-06 | 0.0014322 |
| *anr-32* | 70.68 | 3.35 | -2.45 | 0.55 | 7.60E-06 | 0.0014322 |
| *C15F1.2* | 89.52 | 413.55 | 1.85 | 0.41 | 8.15E-06 | 0.001517 |
| *pcp-4* | 763.12 | 2628.04 | 1.58 | 0.36 | 8.34E-06 | 0.0015332 |
| *str-168* | 3.03 | 56.56 | 2.42 | 0.54 | 8.52E-06 | 0.0015436 |
| *R155.4* | 44.24 | 0.95 | -2.48 | 0.56 | 8.60E-06 | 0.0015436 |
| *clec-5* | 307.38 | 1327.35 | 1.79 | 0.40 | 9.26E-06 | 0.0016423 |
| *wrt-4* | 419.94 | 2345.05 | 1.98 | 0.45 | 9.72E-06 | 0.0016974 |
| *bus-1* | 36.44 | 212.92 | 2.01 | 0.45 | 9.79E-06 | 0.0016974 |
| *D1065.3* | 158.21 | 24.34 | -2.07 | 0.47 | 1.16E-05 | 0.0019846 |
| *sym-1* | 959.27 | 4024.59 | 1.76 | 0.40 | 1.23E-05 | 0.0020816 |
| *R08E5.3* | 197.74 | 1548.81 | 2.16 | 0.50 | 1.30E-05 | 0.0021771 |
| *Y71G12B.33* | 96.35 | 11.94 | -2.17 | 0.50 | 1.32E-05 | 0.0021845 |
| *mfsd-11* | 27.44 | 175.86 | 2.05 | 0.47 | 1.53E-05 | 0.0025096 |
| *elo-5* | 1598.91 | 7505.69 | 1.84 | 0.43 | 1.62E-05 | 0.0025978 |
| *bus-8* | 364.29 | 1301.31 | 1.60 | 0.37 | 1.62E-05 | 0.0025978 |
| *C45B2.8* | 68.30 | 5.74 | -2.28 | 0.53 | 1.68E-05 | 0.0026758 |
| *cyp-13A9* | 0.19 | 41.22 | 2.41 | 0.56 | 1.76E-05 | 0.0027617 |
| *abf-2* | 123.40 | 18.25 | -2.06 | 0.48 | 1.87E-05 | 0.0029066 |
| *dct-16* | 2837.92 | 16263.06 | 1.97 | 0.46 | 2.00E-05 | 0.00308 |
| *Y53F4B.24* | 303.51 | 52.32 | -1.97 | 0.46 | 2.06E-05 | 0.0031397 |
| *ant-1.4* | 397.30 | 100.77 | -1.69 | 0.40 | 2.09E-05 | 0.0031609 |
| *tth-1* | 4363.61 | 1240.54 | -1.58 | 0.37 | 2.12E-05 | 0.0031772 |
| *ugt-63* | 83.83 | 347.62 | 1.73 | 0.41 | 2.22E-05 | 0.0032859 |
| *Y75B7B.2* | 42.26 | 1.43 | -2.36 | 0.56 | 2.28E-05 | 0.0033413 |
| *mel-32* | 5471.66 | 18227.33 | 1.53 | 0.36 | 2.41E-05 | 0.0035011 |
| *F38B6.4* | 268.01 | 1049.30 | 1.68 | 0.40 | 2.43E-05 | 0.0035011 |
| *dlhd-1* | 138.00 | 616.26 | 1.78 | 0.42 | 2.52E-05 | 0.0035601 |

| | | | | | | |
|---|---|---|---|---|---|---|
| *Y54G2A.40* | 209.70 | 21.54 | -2.20 | 0.52 | 2.51E-05 | 0.0035601 |
| *F13A7.12* | 15.31 | 107.59 | 2.06 | 0.49 | 2.56E-05 | 0.0035807 |
| *acox-1.3* | 55.12 | 261.91 | 1.83 | 0.44 | 2.68E-05 | 0.0036868 |
| *sulp-6* | 212.85 | 37.06 | -1.96 | 0.47 | 2.66E-05 | 0.0036868 |
| *F48F5.2* | 194.42 | 31.81 | -1.99 | 0.47 | 2.82E-05 | 0.0037771 |
| *C01F1.5* | 70.83 | 336.63 | 1.83 | 0.44 | 2.80E-05 | 0.0037771 |
| *Y40B10A.4* | 68.17 | 5.71 | -2.24 | 0.53 | 2.77E-05 | 0.0037771 |
| *ZK973.8* | 213.41 | 47.33 | -1.79 | 0.43 | 2.89E-05 | 0.0038287 |
| *K02E11.10* | 887.02 | 3305.37 | 1.63 | 0.39 | 2.93E-05 | 0.0038496 |
| *nhx-2* | 571.87 | 1882.19 | 1.51 | 0.36 | 3.05E-05 | 0.0039715 |
| *W03G1.2* | 189.92 | 39.14 | -1.84 | 0.44 | 3.15E-05 | 0.0040725 |
| *fbxa-125* | 97.62 | 13.86 | -2.05 | 0.49 | 3.20E-05 | 0.0041054 |
| *dsc-4* | 1045.71 | 3499.14 | 1.53 | 0.37 | 3.27E-05 | 0.0041592 |
| *ZC266.1* | 20.70 | 123.63 | 1.96 | 0.47 | 3.46E-05 | 0.0043582 |
| *C35B1.7* | 112.03 | 492.68 | 1.76 | 0.43 | 3.84E-05 | 0.0047989 |
| *nnt-1* | 1637.14 | 322.83 | -1.86 | 0.45 | 4.03E-05 | 0.0049962 |
| *mlt-10* | 618.70 | 2119.35 | 1.55 | 0.38 | 4.18E-05 | 0.0051303 |
| *irg-3* | 240.64 | 798.14 | 1.51 | 0.37 | 4.20E-05 | 0.0051303 |
| *ifc-1* | 276.58 | 1937.29 | 2.04 | 0.50 | 4.32E-05 | 0.0051849 |
| *clec-48* | 1344.88 | 4645.11 | 1.55 | 0.38 | 4.29E-05 | 0.0051849 |
| *F46A8.9* | 96.92 | 14.35 | -2.01 | 0.49 | 4.64E-05 | 0.0053575 |
| *Y37A1B.7* | 53.52 | 258.18 | 1.82 | 0.45 | 4.61E-05 | 0.0053575 |
| *B0222.5* | 2390.26 | 599.12 | -1.68 | 0.41 | 4.66E-05 | 0.0053575 |
| *R02F2.5* | 77.19 | 9.07 | -2.10 | 0.52 | 4.67E-05 | 0.0053575 |
| *T15B7.1* | 524.01 | 2001.50 | 1.64 | 0.40 | 4.54E-05 | 0.0053575 |
| *Y75B7AR.1* | 21.47 | 261.01 | 2.21 | 0.54 | 4.50E-05 | 0.0053575 |
| *sago-2* | 87.58 | 454.32 | 1.87 | 0.46 | 4.73E-05 | 0.0053896 |
| *ZK1251.5* | 153.29 | 18.93 | -2.08 | 0.51 | 4.91E-05 | 0.005549 |
| *odc-1* | 5444.40 | 944.73 | -1.93 | 0.48 | 4.96E-05 | 0.0055634 |
| *endu-2* | 5321.55 | 1730.21 | -1.44 | 0.35 | 5.08E-05 | 0.0056613 |
| *pyk-2* | 342.14 | 1085.88 | 1.47 | 0.36 | 5.29E-05 | 0.00582 |
| *K08D10.18* | 89.85 | 11.91 | -2.05 | 0.51 | 5.30E-05 | 0.00582 |
| *lon-3* | 504.14 | 1580.44 | 1.46 | 0.36 | 5.38E-05 | 0.0058663 |
| *egas-3* | 95.53 | 14.39 | -1.99 | 0.49 | 5.99E-05 | 0.0064662 |
| *wago-10* | 393.12 | 83.55 | -1.79 | 0.45 | 6.02E-05 | 0.0064662 |
| *T26C5.2* | 190.72 | 820.24 | 1.72 | 0.43 | 6.35E-05 | 0.0067774 |
| *nhr-114* | 640.00 | 2559.79 | 1.67 | 0.42 | 6.45E-05 | 0.0068379 |
| *col-51* | 45.77 | 0.96 | -2.23 | 0.56 | 6.92E-05 | 0.0072355 |
| *Y51H4A.13* | 217.76 | 48.12 | -1.76 | 0.44 | 6.92E-05 | 0.0072355 |
| *H17B01.2* | 5.61 | 61.44 | 2.13 | 0.54 | 7.08E-05 | 0.0072936 |
| *cnc-11* | 42.89 | 1.43 | -2.22 | 0.56 | 7.04E-05 | 0.0072936 |
| *dct-18* | 2606.56 | 7854.42 | 1.41 | 0.36 | 7.20E-05 | 0.0073728 |
| *mtrr-1* | 1269.69 | 4293.87 | 1.52 | 0.38 | 7.54E-05 | 0.0076516 |
| *Y54G2A.57* | 38.02 | 0.48 | -2.22 | 0.56 | 7.57E-05 | 0.0076516 |
| *dpy-10* | 329.60 | 1198.39 | 1.58 | 0.40 | 8.18E-05 | 0.008209 |
| *cln-3.1* | 52.47 | 223.39 | 1.71 | 0.43 | 8.30E-05 | 0.0082813 |
| *C44B9.2* | 387.04 | 90.30 | -1.71 | 0.44 | 8.47E-05 | 0.008395 |
| *gst-26* | 275.23 | 1105.11 | 1.66 | 0.42 | 8.70E-05 | 0.0083981 |
| *cdr-4* | 4511.16 | 829.84 | -1.86 | 0.48 | 8.66E-05 | 0.0083981 |
| *lpr-6* | 286.86 | 1055.69 | 1.59 | 0.40 | 8.59E-05 | 0.0083981 |
| *ZK507.1* | 125.41 | 24.40 | -1.83 | 0.47 | 8.61E-05 | 0.0083981 |
| *C18D11.6* | 141.10 | 7.11 | -2.18 | 0.56 | 9.21E-05 | 0.0088394 |
| *F15H10.8* | 75.78 | 307.88 | 1.66 | 0.43 | 9.81E-05 | 0.0092631 |
| *mltn-1* | 183.11 | 619.73 | 1.51 | 0.39 | 9.84E-05 | 0.0092631 |
| *lin-42* | 2801.70 | 8121.32 | 1.37 | 0.35 | 9.73E-05 | 0.0092631 |
| *C38C3.8* | 87.28 | 11.90 | -1.99 | 0.51 | 9.97E-05 | 0.0093345 |

| | | | | | | |
|---|---|---|---|---|---|---|
| *R06C1.4* | 2157.72 | 6367.19 | 1.38 | 0.36 | 0.000101 | 0.0094197 |
| *R06B10.1* | 230.93 | 41.24 | -1.87 | 0.48 | 0.000102 | 0.0094505 |
| *M03B6.3* | 118.62 | 495.33 | 1.68 | 0.43 | 0.000103 | 0.0094826 |
| *F35D2.1* | 23.79 | 135.89 | 1.88 | 0.48 | 0.000109 | 0.0099495 |
| *D1086.2* | 181.17 | 39.90 | -1.74 | 0.45 | 0.000114 | 0.0101989 |
| *F55G11.2* | 26.30 | 166.48 | 1.92 | 0.50 | 0.000113 | 0.0101989 |
| *sdz-24* | 1415.04 | 6428.19 | 1.74 | 0.45 | 0.000113 | 0.0101989 |
| *C17B7.5* | 78.16 | 309.07 | 1.64 | 0.42 | 0.000115 | 0.010214 |
| *cutl-28* | 45.17 | 198.58 | 1.71 | 0.44 | 0.000115 | 0.010214 |
| *W01F3.2* | 2570.74 | 817.44 | -1.44 | 0.37 | 0.000117 | 0.0103326 |
| *ZK507.3* | 140.79 | 29.60 | -1.76 | 0.46 | 0.000125 | 0.0109345 |
| *C32F10.4* | 4373.35 | 1481.30 | -1.38 | 0.36 | 0.000126 | 0.0110146 |
| *hsp-12.6* | 206.78 | 42.28 | -1.78 | 0.47 | 0.000132 | 0.0114098 |
| *spe-47* | 190.76 | 41.40 | -1.74 | 0.46 | 0.000132 | 0.0114098 |
| *argk-1* | 4.10 | 51.44 | 2.09 | 0.55 | 0.000135 | 0.0115385 |
| *enu-3.5* | 742.06 | 198.34 | -1.59 | 0.42 | 0.000141 | 0.0120144 |
| *cpb-2* | 260.26 | 69.21 | -1.59 | 0.42 | 0.000143 | 0.0120981 |
| *Y43D4A.5* | 104.19 | 388.50 | 1.58 | 0.42 | 0.000153 | 0.0129246 |
| *mltn-9* | 231.16 | 816.49 | 1.54 | 0.41 | 0.000157 | 0.0131563 |
| *F27E5.3* | 245.77 | 61.46 | -1.63 | 0.43 | 0.000159 | 0.0132198 |
| *Y46G5A.29* | 197.64 | 668.80 | 1.50 | 0.40 | 0.000162 | 0.0134117 |
| *C45G7.4* | 89.02 | 310.57 | 1.52 | 0.40 | 0.000164 | 0.0134438 |
| *vrp-1* | 373.50 | 1404.05 | 1.59 | 0.42 | 0.000163 | 0.0134438 |
| *Y1A5A.1* | 499.68 | 157.75 | -1.44 | 0.38 | 0.000167 | 0.0136066 |
| *Y64H9A.2* | 46.15 | 190.60 | 1.65 | 0.44 | 0.000168 | 0.0136132 |
| *C01G5.3* | 220.95 | 47.99 | -1.73 | 0.46 | 0.00017 | 0.0137222 |
| *ugt-47* | 521.45 | 1512.07 | 1.36 | 0.36 | 0.000171 | 0.0137578 |
| *B0554.4* | 14.05 | 102.44 | 1.95 | 0.52 | 0.000175 | 0.0139438 |
| *F13B6.3* | 1166.13 | 3448.05 | 1.37 | 0.37 | 0.000182 | 0.014425 |
| *F59H6.5* | 29.88 | 0.48 | -2.10 | 0.56 | 0.000182 | 0.014425 |
| *F26A3.5* | 360.60 | 100.54 | -1.54 | 0.41 | 0.000193 | 0.0151773 |
| *B0393.4* | 374.89 | 86.84 | -1.68 | 0.45 | 0.000194 | 0.0151963 |
| *Y37F4.15* | 42.55 | 2.88 | -2.06 | 0.55 | 0.000197 | 0.0153031 |
| *mec-7* | 554.54 | 2039.37 | 1.56 | 0.42 | 0.000201 | 0.0155975 |
| *swip-10* | 36.40 | 158.25 | 1.68 | 0.45 | 0.000205 | 0.0157727 |
| *myo-3* | 20135.47 | 7139.42 | -1.32 | 0.36 | 0.000208 | 0.0158533 |
| *R07B1.13* | 83.11 | 292.22 | 1.52 | 0.41 | 0.000208 | 0.0158533 |
| *W05H9.1* | 13037.21 | 3342.30 | -1.60 | 0.43 | 0.000218 | 0.0165121 |
| *F21C10.9* | 192.70 | 1074.55 | 1.82 | 0.49 | 0.000221 | 0.0166999 |
| *ptr-4* | 564.22 | 1771.51 | 1.42 | 0.39 | 0.000224 | 0.0167505 |
| *F53H4.2* | 336.09 | 1221.37 | 1.55 | 0.42 | 0.000224 | 0.0167505 |
| *dpy-8* | 632.94 | 2164.86 | 1.50 | 0.41 | 0.000227 | 0.0168738 |
| *C49F5.9* | 12.26 | 78.78 | 1.88 | 0.51 | 0.000228 | 0.0168738 |
| *W10G11.3* | 19.80 | 120.50 | 1.86 | 0.50 | 0.000233 | 0.0171641 |
| *W02D7.11* | 115.45 | 8.09 | -2.04 | 0.56 | 0.000238 | 0.0174338 |
| *grl-4* | 6838.22 | 2346.51 | -1.35 | 0.37 | 0.00024 | 0.0175013 |
| *T02E1.7* | 286.34 | 80.12 | -1.53 | 0.42 | 0.000242 | 0.0175013 |
| *C18H7.11* | 123.68 | 490.26 | 1.61 | 0.44 | 0.000241 | 0.0175013 |
| *oac-1* | 36.14 | 162.34 | 1.69 | 0.46 | 0.000247 | 0.0175282 |
| *K10G4.3* | 560.29 | 107.20 | -1.78 | 0.49 | 0.000247 | 0.0175282 |
| *cest-28* | 3.05 | 41.66 | 2.03 | 0.55 | 0.000245 | 0.0175282 |
| *T17H7.1* | 2.53 | 82.79 | 2.06 | 0.56 | 0.000247 | 0.0175282 |
| *drd-10* | 0.51 | 30.29 | 2.05 | 0.56 | 0.000251 | 0.017732 |
| *lpr-1* | 48.40 | 187.68 | 1.59 | 0.44 | 0.000268 | 0.0187843 |
| *atic-1* | 569.00 | 1647.05 | 1.34 | 0.37 | 0.000268 | 0.0187843 |
| *klo-2* | 40.67 | 205.71 | 1.76 | 0.48 | 0.000278 | 0.0193428 |

| | | | | | | |
|---|---|---|---|---|---|---|
| *C54F6.15* | 26.63 | 0.18 | -2.02 | 0.56 | 0.000279 | 0.0193778 |
| *C23H3.9* | 20.32 | 195.18 | 1.97 | 0.54 | 0.000282 | 0.0194354 |
| *cyn-17* | 6.56 | 83.95 | 2.01 | 0.55 | 0.000289 | 0.0198376 |
| *T11F8.4* | 147.56 | 29.46 | -1.75 | 0.48 | 0.000295 | 0.0201435 |
| *Y39D8A.1* | 120.43 | 501.15 | 1.64 | 0.45 | 0.000296 | 0.0201435 |
| *dpy-9* | 327.99 | 996.25 | 1.39 | 0.38 | 0.000304 | 0.0206266 |
| *pho-1* | 413.23 | 1449.04 | 1.51 | 0.42 | 0.000309 | 0.0206512 |
| *cest-10* | 42.97 | 190.44 | 1.67 | 0.46 | 0.000308 | 0.0206512 |
| *Y58A7A.4* | 73.76 | 6.72 | -1.98 | 0.55 | 0.000307 | 0.0206512 |
| *R08C7.5* | 283.31 | 83.10 | -1.48 | 0.41 | 0.000314 | 0.020939 |
| *snf-4* | 78.00 | 7.57 | -1.97 | 0.55 | 0.000319 | 0.0211742 |
| *mam-3* | 53.10 | 198.70 | 1.55 | 0.43 | 0.00033 | 0.0216129 |
| *nduo-4* | 2665.76 | 7652.07 | 1.33 | 0.37 | 0.000328 | 0.0216129 |
| *W05H9.3* | 6115.60 | 2323.56 | -1.25 | 0.35 | 0.000329 | 0.0216129 |
| *M02H5.8* | 508.77 | 1659.19 | 1.45 | 0.40 | 0.000336 | 0.0218993 |
| *gcy-17* | 31.87 | 0.95 | -2.01 | 0.56 | 0.000342 | 0.0222232 |
| *elo-9* | 77.66 | 308.54 | 1.60 | 0.45 | 0.000348 | 0.0223023 |
| *ZK896.5* | 155.49 | 490.96 | 1.42 | 0.40 | 0.000345 | 0.0223023 |
| *ttr-30* | 1104.36 | 322.93 | -1.48 | 0.41 | 0.000347 | 0.0223023 |
| *wht-5* | 245.03 | 67.20 | -1.53 | 0.43 | 0.000352 | 0.0224852 |
| *oac-36* | 5.17 | 112.06 | 2.00 | 0.56 | 0.00036 | 0.0226664 |
| *ZK1053.6* | 88.97 | 16.72 | -1.76 | 0.49 | 0.000358 | 0.0226664 |
| *cbs-2* | 155.67 | 37.18 | -1.63 | 0.46 | 0.00036 | 0.0226664 |
| *Y39H10A.1* | 84.29 | 12.88 | -1.85 | 0.52 | 0.000361 | 0.0226664 |
| *C27D9.2* | 0.51 | 32.91 | 1.99 | 0.56 | 0.000367 | 0.0229472 |
| *enu-3.1* | 1172.51 | 384.46 | -1.38 | 0.39 | 0.000376 | 0.0234266 |
| *rrf-2* | 457.77 | 123.38 | -1.54 | 0.43 | 0.000387 | 0.0239751 |
| *mec-12* | 771.79 | 2241.55 | 1.34 | 0.38 | 0.000395 | 0.0244188 |
| *oac-14* | 430.44 | 119.79 | -1.52 | 0.43 | 0.000397 | 0.0244212 |
| *lpr-5* | 776.27 | 2591.14 | 1.46 | 0.41 | 0.000403 | 0.024701 |
| *kin-31* | 109.50 | 22.40 | -1.71 | 0.48 | 0.00041 | 0.0249018 |
| *dhcr-24* | 365.79 | 1008.62 | 1.29 | 0.36 | 0.000409 | 0.0249018 |
| *papl-1* | 261.84 | 850.66 | 1.43 | 0.41 | 0.000417 | 0.0252198 |
| *dpy-2* | 442.52 | 1327.80 | 1.37 | 0.39 | 0.000419 | 0.0252421 |
| *C36H8.1* | 1206.52 | 400.64 | -1.37 | 0.39 | 0.000425 | 0.025516 |
| *pgp-12* | 8.41 | 72.87 | 1.90 | 0.54 | 0.000434 | 0.025795 |
| *mam-2* | 134.62 | 406.55 | 1.37 | 0.39 | 0.000435 | 0.025795 |
| *F18C5.5* | 143.20 | 510.89 | 1.51 | 0.43 | 0.000432 | 0.025795 |
| *aat-4* | 58.17 | 242.50 | 1.62 | 0.46 | 0.000443 | 0.0258201 |
| *F35C11.2* | 171.98 | 44.85 | -1.56 | 0.44 | 0.00044 | 0.0258201 |
| *F55F3.4* | 18.25 | 95.77 | 1.74 | 0.50 | 0.000438 | 0.0258201 |
| *W08G11.1* | 67.83 | 246.74 | 1.52 | 0.43 | 0.000438 | 0.0258201 |
| *hpo-15* | 5694.68 | 1600.61 | -1.51 | 0.43 | 0.000443 | 0.0258201 |
| *F17B5.8* | 11.42 | 83.25 | 1.86 | 0.53 | 0.000449 | 0.0260586 |
| *Y47D7A.6* | 31.09 | 189.62 | 1.80 | 0.52 | 0.000475 | 0.0274727 |
| *C15C6.1* | 205.80 | 631.79 | 1.38 | 0.40 | 0.000483 | 0.0277949 |
| *F56A4.3* | 67.95 | 273.90 | 1.59 | 0.46 | 0.000492 | 0.0281931 |
| *faah-3* | 1162.06 | 413.09 | -1.30 | 0.37 | 0.000496 | 0.0283338 |
| *clec-97* | 77.23 | 349.38 | 1.66 | 0.48 | 0.000503 | 0.0286535 |
| *F23H12.5* | 487.17 | 1512.69 | 1.39 | 0.40 | 0.00051 | 0.0289451 |
| *vit-1* | 4271.97 | 14212.11 | 1.45 | 0.42 | 0.000514 | 0.0290279 |
| *ncr-2* | 177.13 | 47.22 | -1.54 | 0.44 | 0.000523 | 0.0294208 |
| *F32A11.3* | 2212.02 | 775.00 | -1.31 | 0.38 | 0.000525 | 0.0294208 |
| *Y32F6A.6* | 259.93 | 78.31 | -1.44 | 0.42 | 0.00053 | 0.0296004 |
| *C49G7.10* | 675.36 | 162.61 | -1.60 | 0.46 | 0.000534 | 0.0297423 |
| *F35E2.5* | 213.25 | 41.17 | -1.72 | 0.50 | 0.000546 | 0.0302934 |

| | | | | | | |
|---|---|---|---|---|---|---|
| *F36H12.14* | 148.73 | 36.71 | -1.58 | 0.46 | 0.00056 | 0.0309492 |
| *gska-3* | 396.85 | 133.34 | -1.35 | 0.39 | 0.000562 | 0.0309542 |
| *clec-76* | 16.90 | 98.33 | 1.77 | 0.51 | 0.000571 | 0.031348 |
| *alh-13* | 971.66 | 2659.36 | 1.27 | 0.37 | 0.000604 | 0.0330353 |
| *dpy-3* | 413.45 | 1323.69 | 1.41 | 0.41 | 0.000613 | 0.0333604 |
| *F13B6.2* | 201.28 | 623.92 | 1.38 | 0.40 | 0.000614 | 0.0333604 |
| *lpr-3* | 1685.24 | 5659.25 | 1.45 | 0.42 | 0.000619 | 0.0334991 |
| *poml-2* | 44.67 | 197.64 | 1.63 | 0.48 | 0.000624 | 0.0336671 |
| *col-39* | 12348.41 | 4806.34 | -1.21 | 0.35 | 0.000637 | 0.0340791 |
| *C53A3.1* | 4.04 | 44.67 | 1.89 | 0.55 | 0.000636 | 0.0340791 |
| *T19C9.8* | 48.97 | 193.12 | 1.56 | 0.46 | 0.000653 | 0.0348377 |
| *col-179* | 2522.07 | 7950.09 | 1.39 | 0.41 | 0.000659 | 0.0350422 |
| *R07B1.5* | 26.38 | 124.08 | 1.66 | 0.49 | 0.000672 | 0.0355755 |
| *set-9* | 2417.57 | 718.39 | -1.45 | 0.43 | 0.000679 | 0.0355755 |
| *hacd-1* | 2551.86 | 7163.47 | 1.29 | 0.38 | 0.000677 | 0.0355755 |
| *lgc-27* | 22.77 | 107.16 | 1.66 | 0.49 | 0.000676 | 0.0355755 |
| *cnnm-5* | 80.74 | 11.85 | -1.80 | 0.53 | 0.000682 | 0.0356095 |
| *dpy-7* | 472.76 | 1506.05 | 1.40 | 0.41 | 0.00069 | 0.0358018 |
| *bgnt-1.8* | 2.06 | 40.69 | 1.90 | 0.56 | 0.000688 | 0.0358018 |
| *Y110A2AL.2* | 33.91 | 1.42 | -1.90 | 0.56 | 0.000698 | 0.0361161 |
| *Y53F4B.11* | 143.20 | 37.83 | -1.53 | 0.45 | 0.000701 | 0.0361294 |
| *fkb-5* | 318.74 | 1177.30 | 1.51 | 0.45 | 0.000713 | 0.0364176 |
| *Y105E8A.13* | 79.43 | 254.66 | 1.40 | 0.41 | 0.000714 | 0.0364176 |
| *Y49F6B.8* | 264.16 | 72.26 | -1.50 | 0.44 | 0.000709 | 0.0364176 |
| *Y43F8C.13* | 338.10 | 896.53 | 1.24 | 0.37 | 0.000743 | 0.0377713 |
| *Y43F4A.1* | 215.80 | 663.72 | 1.37 | 0.41 | 0.00076 | 0.038362 |
| *F26D11.2* | 6.60 | 51.06 | 1.82 | 0.54 | 0.000758 | 0.038362 |
| *bah-1* | 110.43 | 459.38 | 1.58 | 0.47 | 0.000762 | 0.038362 |
| *C18E9.8* | 284.66 | 80.35 | -1.48 | 0.44 | 0.00077 | 0.0386652 |
| *acs-15* | 209.10 | 49.82 | -1.59 | 0.47 | 0.000774 | 0.0387242 |
| *col-36* | 275.74 | 41.79 | -1.78 | 0.53 | 0.000816 | 0.0406974 |
| *F36G9.13* | 127.08 | 20.82 | -1.75 | 0.52 | 0.000822 | 0.0408357 |
| *Y24D9B.1* | 243.29 | 63.16 | -1.53 | 0.46 | 0.000825 | 0.040846 |
| *cpt-6* | 149.91 | 430.12 | 1.30 | 0.39 | 0.000854 | 0.0421439 |
| *ent-7* | 78.65 | 244.61 | 1.37 | 0.41 | 0.000862 | 0.0424117 |
| *T06C10.3* | 210.41 | 64.49 | -1.41 | 0.42 | 0.000886 | 0.0434476 |
| *cdh-7* | 172.09 | 567.93 | 1.42 | 0.43 | 0.000898 | 0.0436217 |
| *bcat-1* | 3415.05 | 9611.44 | 1.29 | 0.39 | 0.000901 | 0.0436217 |
| *irld-57* | 46.24 | 4.75 | -1.83 | 0.55 | 0.000897 | 0.0436217 |
| *M60.4* | 9863.47 | 3410.01 | -1.31 | 0.39 | 0.000894 | 0.0436217 |
| *linc-22* | 542.64 | 194.04 | -1.28 | 0.39 | 0.000908 | 0.0438197 |
| *clec-47* | 346.88 | 1574.21 | 1.62 | 0.49 | 0.000918 | 0.0441618 |
| *B0207.1* | 207.58 | 53.68 | -1.53 | 0.46 | 0.000927 | 0.0444891 |
| *hil-3* | 719.30 | 1818.54 | 1.18 | 0.36 | 0.000943 | 0.0448936 |
| *F32B4.8* | 8.67 | 58.13 | 1.76 | 0.53 | 0.000947 | 0.0448936 |
| *Y39G8C.2* | 89.82 | 15.66 | -1.72 | 0.52 | 0.000942 | 0.0448936 |
| *ZK1010.5* | 1036.82 | 309.83 | -1.43 | 0.43 | 0.000947 | 0.0448936 |
| *R05A10.6* | 3.11 | 38.81 | 1.84 | 0.56 | 0.000959 | 0.045282 |
| *F14F7.4* | 197.83 | 58.68 | -1.43 | 0.43 | 0.000968 | 0.0455703 |
| *F28F8.7* | 254.71 | 82.79 | -1.36 | 0.41 | 0.00099 | 0.0464661 |
| *lbp-8* | 0.19 | 23.02 | 1.82 | 0.55 | 0.000994 | 0.0465296 |
| *C38D4.7* | 103.72 | 24.34 | -1.58 | 0.48 | 0.001005 | 0.0468986 |
| *hmit-1.1* | 834.55 | 2102.25 | 1.18 | 0.36 | 0.001011 | 0.0470229 |
| *Y66D12A.11* | 358.36 | 112.79 | -1.38 | 0.42 | 0.00104 | 0.0479186 |
| *hach-1* | 1245.64 | 3077.38 | 1.16 | 0.35 | 0.001038 | 0.0479186 |
| *F56D2.5* | 411.47 | 123.33 | -1.42 | 0.43 | 0.001038 | 0.0479186 |

| | | | | | | |
|---|---|---|---|---|---|---|
| *C09B9.7* | 155.80 | 36.08 | -1.58 | 0.48 | 0.001048 | 0.0481567 |
| *mab-7* | 203.23 | 553.60 | 1.25 | 0.38 | 0.001055 | 0.0483358 |
| *sod-5* | 69.05 | 7.69 | -1.80 | 0.55 | 0.001062 | 0.0485158 |
