## Supplementary material for "Genome-wide screen identifies curli amyloid fibril as a bacterial component promoting host neurodegeneration": key resources table

| REAGENT or RESOURCE | SOURCE | IDENTIFIER |
|---|---|---|
| **Antibodies** | | |
| alpha Synuclein Monoclonal Antibody (Syn 211) | Thermo Fisher | Cat # 32-8100; RRID:AB_2533094 |
| Mouse monoclonal [LB 509] to Alpha-synuclein, 100ug | Abcam | Cat #ab27766; RRID:AB_727020 |
| Monoclonal ANTI-FLAG® M2 antibody produced in mouse 1 mg/mL | Sigma -Aldrich | Cat #F1804; RRID:AB_262044 |
| ANTI-FLAG® antibody produced in rabbit | Merck | Cat #F7425; RRID:AB_439687 |
| Rhodamine (TRITC) AffiniPure Goat Anti-Rabbit IgG (H+L) | Jackson ImmunoResearch | Cat #111-025-003; RRID:AB_2337926 |
| Alexa Fluor® 488 AffiniPure Goat Anti-Mouse IgG (H+L) | Jackson ImmunoResearch | Cat #115-545-003; RRID:AB_2338840 |
| Alexa Fluor® 647 AffiniPure Donkey Anti-Rabbit IgG (H+L) | Jackson ImmunoResearch | Cat #711-605-152; RRID:AB_2492288 |
| Rhodamine (TRITC) AffiniPure Donkey Anti-Rabbit IgG (H+L) | Jackson ImmunoResearch | Cat #711-025-152; RRID:AB_2340588 |
| Alexa Fluor® 488 AffiniPure Goat Anti-Rabbit IgG (H+L) | Jackson ImmunoResearch | Cat #111-545-003; RRID:AB_2338046 |
| Peroxidase AffiniPure Goat Anti-Rabbit IgG (H+L) | Jackson ImmunoResearch | Cat #111-035-003; RRID:AB_2313567 |
| Anti-Mouse IgG (whole molecule)–Peroxidase antibody produced in rabbit | Sigma -Aldrich | Cat #A9044; RRID:AB_258431 |
| GAPDH Loading Control Monoclonal Antibody (GA1R) | Thermo Fisher | Cat # MA5-15738; RRID:AB_10977387 |
| **Bacterial and Virus Strains** | | |
| UTI2 | From Dr. Aixin Yan | N/A |
| Escherichia coli K-12 Keio Knockout Collection | Dharmocon | Cat# OEC4988 |
| Escherichia coli OP50 | CGC | N/A |
| E. coli Keio Knockout parent strain BW25113 | Dharmocon | Cat# OEC5042 |
| **Chemicals, Peptides, and Recombinant Proteins** | | |
| N'-QYGGNA-C' | Ontores Biotechnology, Shanghai, China | N/A |
| N'-QYGGNN-C' | Ontores Biotechnology, Shanghai, China | N/A |
| Rhodamine B-QYGGNN | Ontores Biotechnology, Shanghai, China | N/A |
| Ubiquitin E1 Inhibitor, PYR-41 | Sigma -Aldrich | Cat# 662105 |
| Bortezomib, 5mg - CAS 179324-69-7 | Sigma -Aldrich | Cat# 5043140001 |
| InSolution MG-132 in EtOH, ≥95% by HPLC - Calbiochem | Sigma -Aldrich | Cat# 474787 |
| (−)-Epigallocatechin gallate | Sigma -Aldrich | Cat# PHR1333 |
| **Critical Commercial Assays** | | |
| XFe24 FLUXPAK MINI (includes 6 XFe24 sensor cartridges, 10 XF24 cell culture microplates, and 1 bottle of Seahorse XF Calibrant Solution 500mL) (D1) | Agilent Technologies | Cat# 102342-100 |
| Agilent Seahorse XFe24 Analyzer | Agilent Technologies | RRID:SCR_019539 |

| | | |
|---|---|---|
| Cell Counting Kit 8 (CCK8) | Abcam | Cat# ab228554 |
| **Deposited data** | | |
| Raw image data for western blot | This paper and Mendeley data | https://data.mendeley.com/datasets/vyc23scp8p/1 |
| RNA-seq data | This paper and GEO | https://www.ncbi.nlm.nih.gov/geo/query/acc.cgi?acc=GSE169204 |
| **Experimental Models: Cell Lines** | | |
| Human: SH-SY5Y line | ATCC® | CRL-2266™; RRID:CVCL_0019 |
| **Experimental Models: Organisms/Strains** | | |
| Caenorhabditis elegans N2 | CGC | RRID:SCR_007341 |
| UM10 *unkIs7[aex-3p::α-syn(A53T), dat-1p::gfp]* | This paper | N/A |
| UM6 *unkIs9 [dat-1p::α-syn(A53T), dat-1p::gfp]* | This paper | N/A |
| **Oligonucleotides** | | |
| Primers for qPCR, see Table S3 | This paper | N/A |
| Homolgous arms for inserting 3xFLAG tag in K12 and UTI2 E. coli, see Table S3 | This paper | N/A |
| Homolgous arms for deleting csgA in UTI2 E. coli, see Table S3 | This paper | N/A |
| **Recombinant DNA** | | |
| pHM6- alphasynuclein-WT | Addgene | Cat# 40824 |
| pHM6- alphasynuclein-A53T | Addgene | Cat# 40825 |
| EGFP-alphasynuclein-WT | Addgene | Cat# 40822 |
| EGFP- alphasynuclein-A53T | Addgene | Cat# 40823 |
| **Software and Algorithms** | | |
| Leica Application Suite X (3.7.0.20979) software | Leica | https://www.leica-microsystems.com/products/microscope-software/p/leica-las-x-ls/ |
| THUNDER Imaging Systems | Leica | https://www.leica-microsystems.com/products/thunder-imaging-systems/ |
| Graphpad Prism 8.0 | Graphpad | https://www.graphpad.com/scientific-software/prism/ |
| ImageJ | Schneider et al., 2012 | https://imagej.nih.gov/ij/ |
